# Insights into the phylogeny and enigmatic mitochondrial biology of eustigmatophyte algae from over 50 newly sequenced organellar genomes

**DOI:** 10.64898/2026.03.28.715030

**Authors:** Michal Richtář, Eliška Klapuchová, Tatiana Yurchenko, Karen P. Fawley, Marvin W. Fawley, Dovilė Barcytė, Karin Jaške, Benjamin M. Wolf, Fay-Wei Li, Tereza Ševčíková, Marek Eliáš

## Abstract

Organellar genomes are both a resource for reconstructing organismal phylogenies and interesting subjects for evolutionary studies. Herein, we focused on the organellar genomes of eustigmatophytes (eustigs), a class of the algal phylum Ochrophyta with a growing biotechnological potential, and massively expanded the existing limited sample by 51 new organellar genomes. Analyses of this large dataset provided a robustly resolved eustig phylogeny and important insights into the evolution of unique features of eustig organellar genomes. Eustig plastomes are rather stable in terms of the gene content, with only minor differences stemming from differential gene loss and rare lineage-specific gain. In contrast, eustig mitogenomes contain a very stable core of conserved genes accompanied by a broadly varying shell comprising “accessory” genes. Notably, the new data illuminated the origin of two mitochondrial genes previously deemed eustig-specific, namely *orfX* and *orfY* that were found to have evolved, respectively, by *rps4* duplication and extreme divergence of an *rps1* ortholog. Most interestingly, we identified five previously unrecognized orthogroups of mysterious mitochondrial *orfs* that are patchily distributed across eustigs yet likely evolved in the ancestor of this class. These *orfs* have no discernible homologs outside eustigmatophytes but are predicted to encode multipass membrane proteins with a soluble C-terminal domain. Our results also revise some of the previous conclusions regarding the mitochondrial translation in eustigs and suggest the recruitment of a group of unusual tRNAs for a translation-independent function in the genus *Vischeria*. Our study thus provides a glimpse into a “dark matter” of mitochondrial biology in eustigmatophytes.

**Significance statement:** Organellar genomes have been characterized for only a small subset of members of Eustigmatophyceae, a class of biotechnologically attractive coccoid microalgae within the phylum Ochrophyta. In this study, we present >50 newly determined organellar genome sequences from diverse eustigmatophytes, filling most major gaps in sampling across the eustigmatophyte phylogenetic tree and more than doubling the number of species represented. This greatly expanded dataset was analyzed to provide numerous new insights into phylogenetic relationships within the class, variation in organellar gene content, and the evolutionary processes and trends shaping eustigmatophyte organellar genomes. Most notably, our work reveals a surprisingly rich and evolutionarily dynamic repertoire of eustigmatophyte-specific mitochondrial genes, pointing to a functional “dark matter” within eustigmatophyte mitochondria.

## Introduction

Despite a large number of plastid and mitochondrial genome sequences in databases, organellar genomes remain an attractive sequencing target. Firstly, they provide a wealth of data for inferring organismal phylogeny, being instrumental in reconstructing the “backbone” phylogeny of particular groups (Lemieux et al. 2014; Fučíková et al. 2019) as well as in recognizing novel lineages at a high taxonomic level (Leliaert et al. 2016; Barcytė et al. 2021; Jamy et al. 2025). Secondly, the sampling of various groups, such as diverse little-known protist taxa, is still far from satisfying. Thirdly, analyses of newly determined organellar sequences do not cease to yield discoveries of unprecedented features or new extremes, including new variants of the genetic code (Lukeš et al. 2025), outrageous proliferation of introns or repeats (Muñoz-Gómez et al. 2017; Pánek et al. 2022), incredibly low GC content values (Nguyen et al. 2020), or acquisition of lineage-specific genes and other types of genetic elements (Milner et al. 2021; Barcytė et al. 2024). Finally, improvements in the organellar genome sampling may be critical for functional interpretations of the genome sequences, such as identification of evolutionarily constrained regions indicative of their functional significance or distinguishing lineage-specific new organellar genes from non-functional open-reading frames (ORFs) that have arisen randomly or from degenerated insertions of various parasitic genetic elements.

In this study we present the main results of our taxonomically comprehensive exploration of organellar genomes of the algal class Eustigmatophyceae. Eustigmatophytes, or eustigs for short, are unicellular coccoid algae that constitute a separate, well-defined lineage of a broad algal assembly called the Ochrophyta (Eliáš et al. 2017). As such, they are distant relatives of more familiar groups, including diatoms, brown algae, and golden algae. Eustigs have become one of the most extensively studied microalgal groups, owing particularly to the fact that they are considered a highly promising resource for biotechnological applications (Stoyneva-Gärtner et al. 2019). Work in the past decade has resulted in a major overhaul of our view of eustig phylogenetic diversity. Based mainly on initial taxonomic studies in the 1970’s and 1980’s, eustigs were considered to be an inconspicuous species-poor group, but this perspective has changed with recent renewed culturing efforts and environmental DNA surveys, uncovering brand new diversity in the group (Wolf et al. 2018; Fawley et al. 2019; Fawley et al. 2021; Barcytė et al. 2022; Přibyl and Procházková 2022; Alexandre et al. 2025), and with reassessment of previously described taxa originally classified in other algal classes but eventually recognized as eustigs upon closer investigation (Fawley and Fawley 2017; Amaral et al. 2020; Amaral et al. 2021). However, this vastly expanded concept of the Eustigmatophyceae is still not properly reflected by the formal classification of the group.

In phylogenetic terms, the group embraces two deeply diverged major monophyletic groups, the order Eustigmatales and the clade *Goniochloridales* (described under the PhyloCode and for technical reasons not yet formalized under the International Code of Nomenclature for algae, fungi, and plants). These two order-level clades are further divided into multiple lineages. Thus, the principal lineages of Eustigmatales are the families Monodopsidaceae, Chlorobotryaceae (previously referred to as the “Eustigmataceae group”), and Neomonodontaceae (originally introduced with the incorrect spelling “Neomonodaceae”), plus the genus *Paraeustigmatos* (Fawley et al. 2014; Fawley et al. 2019; Amaral et al. 2020; Barcytė et al. 2022). In *Goniochloridales*, the genus *Pseudostaurastrum* and three clades informally denoted IIa, IIb and IIc were recognized (Fawley et al. 2014; Přibyl and Procházková 2022). However, the branching order in many areas of the eustigmatophyte phylogeny remains unresolved, and some of the recently characterized isolates do not fit neatly into any of the presently delimited clades. Furthermore, 18S rRNA and *rbcL* gene sequences obtained from PCR-based environmental surveys suggest the existence of additional eustig lineages (both within the confines of the two order-level clades as well as outside them) that lack cultured representatives (Villanueva et al. 2014; Fawley et al. 2021).

Although the Eustigmatophyceae are no longer thought to be a narrow taxon, the attention paid by researchers to eustigs has been extremely biased towards a single lineage represented by the closely related genera *Nannochloropsis* and *Microchloropsis* (Fawley et al. 2015). This lineage is in many respects rather unrepresentative of eustigmatophytes as a whole, also because it thrives primarily in marine environments, whereas eustigs as a group are ancestrally and predominantly freshwater algae (Eliáš et al. 2017). Genomics is particularly instrumental in illuminating eustig biology, unsurprisingly first targeting *Nanno-*/*Microchloropsis* (Pan et al. 2011; Radakovits et al. 2012; Vieler et al. 2012; Wang et al. 2014), but recently also members of the genera *Vischeria* (syn. *Eustigmatos*; Kryvenda et al. 2018) and *Monodopsis* (Yang et al. 2021; Gao et al. 2023). Analyses of the eustig nuclear genome features and gene repertoire have provided numerous fundamental insights into the cell biology, biochemistry, physiology, and evolution of these algae. For instance, survey sequencing of the genome of *Trachydiscus minutus* led to the discovery of a new genus of endosymbiotic bacteria, *Candidatus* Phycorickettsia, that was found in multiple distantly related eustigs (but noticeably not in *Nanno-*/*Microchloropsis*) and seems to be specifically associated with this algal group (Yurchenko et al. 2018).

However, the scope of nuclear genome exploration in eustigs lags behind the extent of organellar genome investigation. Thanks to an effort of multiple labs including ours, plastid genome (plastome) sequences have been determined not only for most of the *Nanno-*/*Microchloropsis* species (Wei et al. 2013), but also for representatives of most other major eustig lineages (Ševčíková et al. 2015; Yurchenko et al. 2016; Huang et al. 2019a; Ševčíková et al. 2019; Barcytė et al. 2022). Mitochondrial genomes (mitogenomes) have been much less extensively explored in eustigs; in addition to the *Nanno-*/*Microchloropsis* species (Wei et al. 2013), full mitogenome sequences have been reported for several strains of *Vischeria* and single representatives of two other genera (Ševčíková et al. 2016; Huang et al. 2019b; Luo et al. 2023).

Comparative analyses have painted an overall picture of the diversity and evolution of organellar genomes in eustigs, and unveiled a number of interesting aspects, including novel eustig-specific organellar genes of unknown function or unique modifications to the structure of standard genes and proteins encoded by them. For example, analysis of the plastomes of several eustigs from the families Monodopsidaceae and Chlorobotryaceae revealed the existence of a previously unknown six-gene cluster, a putative operon denoted *ebo* (Yurchenko et al. 2016; Barcytė et al. 2022), shared with diverse bacteria and acquired by eustigs from a *Phycorickettsia*-related bacterium (Yurchenko et al. 2018). Only later did the broad functional significance of the *ebo* operon and its actual biochemical role begin to be investigated (Klicki et al. 2018; Zhou et al. 2022; Tanoeyadi et al. 2024).

Despite the recent progress in eustig organellar genomics, numerous important unresolved questions and significant sampling gaps remain. Here, we report on a massive expansion of the number of completely determined eustig organellar genome sequences compared to previous analyses. We exploited these data to obtain the most taxonomically comprehensive multigene phylogenies of eustigmatophytes by far, robustly resolving the relationships of all major lineages. Backed by this new phylogenetic framework, we updated the picture of the evolutionary changes of the plastid gene content in eustigs. Our main focus, however, was on the much less explored eustig mitogenomes, resulting in a fundamental update of our decade-old study (Ševčíková et al. 2016). Our findings underscore surprising evolutionary malleability of eustig mitogenomes and unveil the existence of unknown biological processes mediated by eustig-specific mitochondrial genes.

## Results and Discussion

### Expanding the eustigmatophyte sequence data resources with 20 new plastid and 31 new mitochondrial genomes

Guided by taxonomically comprehensive phylogenies of 18S rRNA and *rbcL* genes (Figs. S1 and S2) and considering the existing set of eustig organellar genomes, we selected 16 eustig strains as new targets for generating Illumina reads from DNA isolated from the respective cultures (Table S1). Light microscopy images of ten newly sequenced strains that are not previously morphologically documented in the literature are provided in Fig. 1 for reference. Complete sequences of plastomes and mitogenomes were obtained by extracting the corresponding contigs from assemblies of the sequence data followed by validation of the completeness of the respective genome or manual assembly of a complete sequence from multiple contigs (see Materials and Methods for details). We obtained both types of organellar genomes for all these strains. In the case of the strain *Pseudotetraëdriella kamillae* SAG 2056, sequencing of the original material obtained from the culture collection revealed that the culture is dominated by a member of the eustig genus *Vischeria*. We assembled both organellar genomes of this contaminant and included it in our analyses as *Vischeria* sp. ex *P. kamillae* SAG 2056. Authentic *P. kamillae* organellar genome sequences were then obtained from a newly received SAG 2056 culture. Additional previously unreported complete organellar genome sequences were obtained by exploiting existing eustig sequencing data (Ševčíková et al. 2019; Yang et al. 2021; Barcytė et al. 2022). Together with already published sequences and considering only a single representative strain for each species of the genera *Nannochloropsis* and *Microchloropsis*, our final dataset comprised 42 different eustig strains for which both the plastome and mitogenome sequences have been determined (details provided in Tables S2 and S3, respectively).

**Fig. 1.**
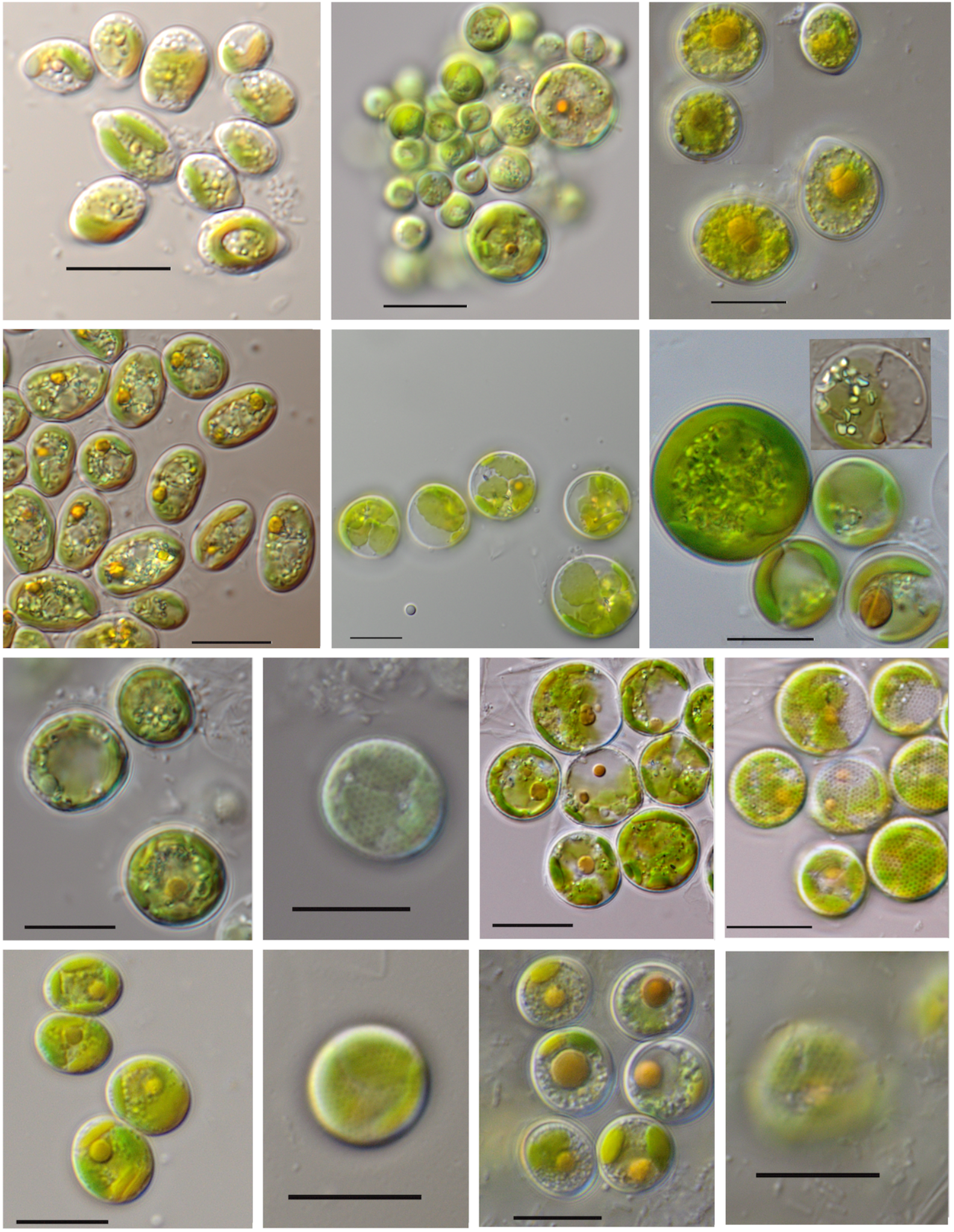
Light microscopy images of a selection of eustigmatophyte strains subjected to genome sequencing in this study. (A) Monodopsidaceae sp. WarPS-5; (B) Chlorobotryaceae sp. WTwin 8/9 T-6m6.8; (C) Neomonodontaceae sp. BogD 8/9 T-6m; (D) Chlorobotryaceae sp. BogD 8/9 T-7m; (E) Goniochloridales sp. WTwin 8/9 T-15m6.8; (F) Goniochloridales sp. BogD 8/9 T-2w (vegetative cells; image at 100× in right corner showing v-shaped crystal on red-orange globule), (G) Goniochloridales sp. WTwin 8/9 T-7m6.8 vegetative cells and (H) cells with sculpting; (I) Goniochloridales sp. Chic 4/9 P-4w (2 images)-vegetative cells and (J) cells with sculpting; (K) Goniochloridales sp. UP3 5/31-12m vegetative cells and (L) cell with sculpting; (M) WTwin 8/9T-12m6.8-vegetative cells and (N) cell with sculpting. Scale bar: 10 μm.

The basic characteristics of all the organellar genomes analyzed here for the first time, including the size, GC content, and the number of annotated genes (Tables 1, S2, S3, and S4), fit into the range defined by previous investigations of eustig organellar genomes (Ševčíková et al. 2016; Ševčíková et al. 2019). All the genomes are circular-mapping and all the plastid genomes exhibit the canonical organization with inverted repeats, comprising the rDNA region plus a varying set of other genes, separated by the short and long single copy regions (Green 2011; Datasets S1 and S2). While mitochondria or plastids of certain algal groups exhibit non-standard genetic code variants (e.g., Turmel et al. 2019; Barcytė et al. 2025), no indications for codon reassignments were detected in any of the eustig organellar genomes analyzed, so the standard translation table was employed to annotate the whole organellar genome set (in some cases allowing for an initiation codon other than AUG, as is common in organelles). As demonstrated by previous investigations of a smaller eustig organellar sample (e.g. Ševčíková et al. 2016; Ševčíková et al. 2019), the gene sets in both plastid and mitochondrial genomes varied across different eustig lineages, and the salient aspects of this variation are discussed below.

**Table 1.**
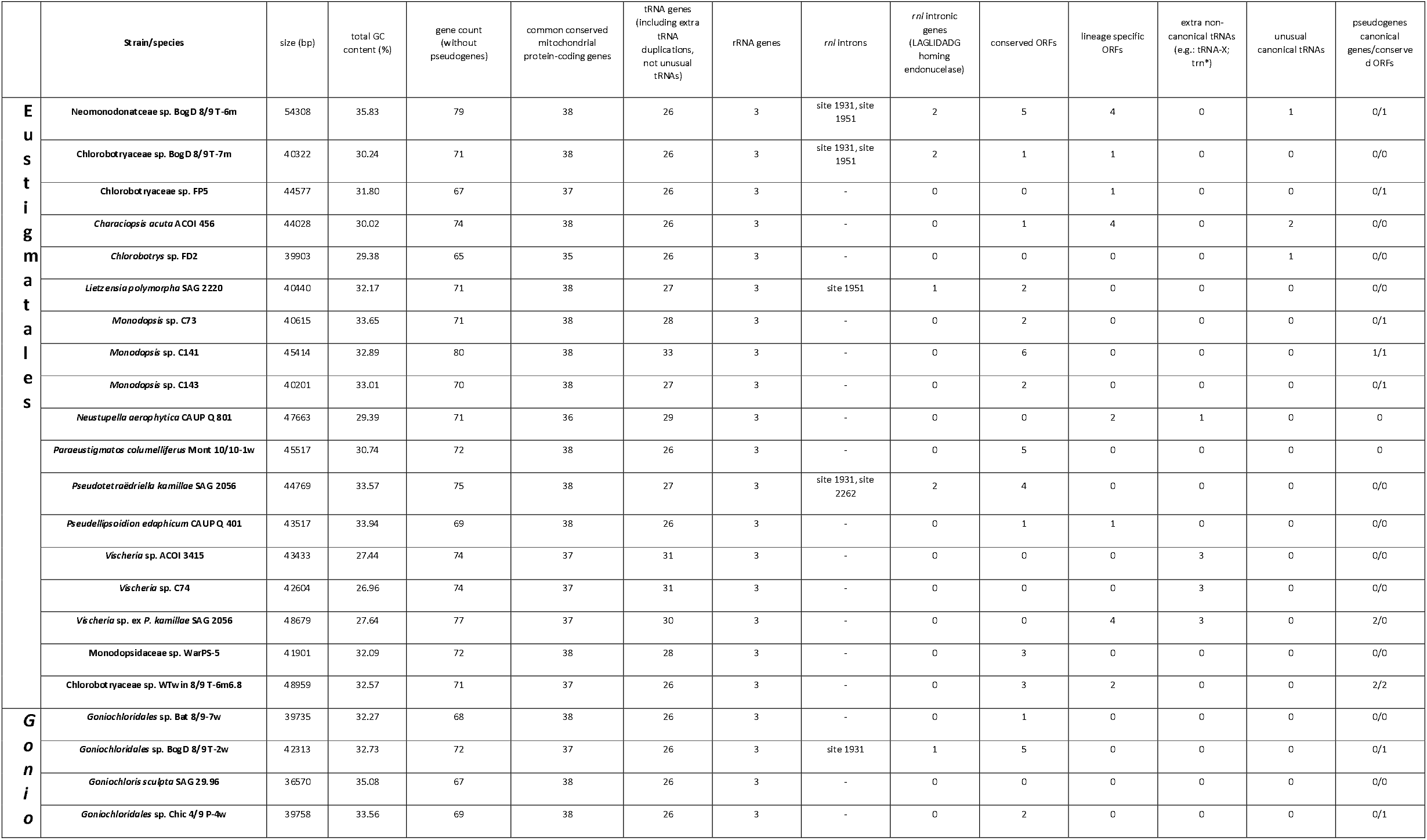
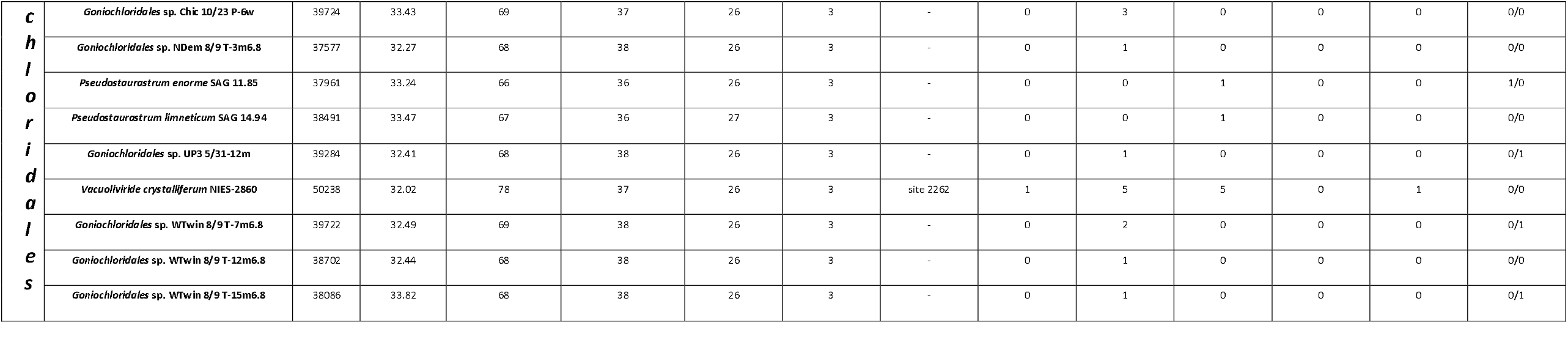
Basic characteristics of the newly sequenced mitochondrial genomes in Eustigmatales and *Goniochloridales*.

For reference we obtained updated 18S rRNA and *rbcL* phylogenies of the class Eustigmatophyceae (Figs. S1 and S2), for which we utilized the newly generated genome sequence data further supplemented by a separately determined 18S rRNA gene sequence from a *Goniochloridales* strain (F5a 4/24-2w) not targeted for genome sequencing. Our sequencing effort has filled major gaps in the sampling of the eustig phylogenetic diversity, with most major lineages of the class as defined by 18S rRNA and *rbcL* phylogenies represented by at least one species or strain in the present set. Filling in the remaining major sampling gaps, such as several *Goniochloridales* strains from the ACOI culture collection that form relatively long branches in the 18S rRNA tree and the eustig clades currently represented solely by environmental DNA sequences (Fig. S1; Fawley et al. 2021; Barcytė et al. 2022), is an obvious target for future research.

Both 18S rRNA and *rbcL* gene phylogenies consistently indicate the existence of two major monophyletic subgroups of the family Monodopsidaceae, one combining the genus *Monodopsis* with the species *Pseudotetraëdriella kamillae* and the strain WarPS-5, and the other comprised of the genera *Nannochloropsis* and *Microchloropsis* (Figs. S1 and S2). This division is fully supported by organellar phylogenomic analyses (see below) and members of each subgroup share specific traits, so it is convenient to recognize the subgroups as formally named taxa. We therefore describe these subgroups as two tribes, Monodopsideae and Nannochloropsideae, respectively (the formal description is provided below). Furthermore, we reassign the unidentified *Goniochloridales* strain NDem 8/9 T-3m6.8 and two related strains from clade IIa into a newly delimited informal group denoted clade IId. This is to account for their deep split from clade IIa members in the organellar phylogenomic and 18S rRNA trees, and even the lack of clustering with clade IIa members in the *rbcL* phylogeny (Fig. S2).

### Comprehensive phylogeny of eustigmatophytes established by phylogenomic analyses of organellar genome data

The previous study into the eustig phylogeny based on a concatenated dataset of 69 plastid-encoded proteins yielded an essentially fully resolved tree (Barcytė et al. 2022), contrasting with the much less resolved (albeit better sampled) 18S rRNA and *rbcL* phylogenies (Figs. S1 and S2). It was thus natural to exploit the newly established sequence resources to obtain an updated plastid phylogenomic tree of eustigmatophytes. A supermatrix built by concatenation of trimmed alignments of 69 conserved plastome-encoded proteins and comprising 15,541 aligned amino acid positions was analyzed in IQ-TREE 3 (Wong et al. 2025) using the maximum likelihood (ML) method and the site-heterogeneous model LG+C60+G+F.

The resulting tree (Fig. 2) provides an excellent resolution of the eustig phylogeny and supports all the previously recognized order- and family-level clades and their branching order as recovered by the previous much less comprehensive analysis (Barcytė et al. 2022). Our results confirm the deep divide of the family Chlorobotryaceae into the so-called clade Ia and another clade comprising all other family members. The branching order of *Lietzensia polymorpha, Chlorobotrys* sp. FD2, and a subclade combining *Vischeria* spp. with *Neustupella aerophytica* is not fully resolved and differs from that obtained in the previous plastome-based multigene analysis (Barcytė et al. 2022), but it is notable that the new topology is more congruent with the phylogeny based on mitogenome-encoded proteins (see below). The division of Monodopsidaceae into two subclades equal to the newly distinguished tribes Monodopsideae and Nannochloropsideae is fully supported. In *Goniochloridales* the deepest split separates *Goniochloris sculpta*, a sole representative of the previously unsampled clade IIb, from other *Goniochloridales* subgroups. The two species of the genus *Pseudostaurastrum* that are also a new addition to the analysis branch together as expected and most likely form a sister group of clade IIc. The unidentified strain NDem 8/9 T-3m6.8, representing the newly recognized clade IId (see above), is positioned as a deeply separated sister lineage of clade IIa as in previous analyses (Ševčíková et al. 2019; Barcytė et al. 2022). Our analysis also recapitulates the previously observed sisterhood of eustigs and an uncultivated marine alga represented by the plastome TARA_CHLORO_00525 reconstructed from TARA Oceans metagenomic data (Jamy et al. 2025). The identity of TARA_CHLORO_00525 is unknown, but it likely represents one of the marine ochrophyte (MOCH) lineages known from environmental 18S rRNA gene clones (Massana et al. 2014; Terpis et al. 2025).

**Fig. 2.**
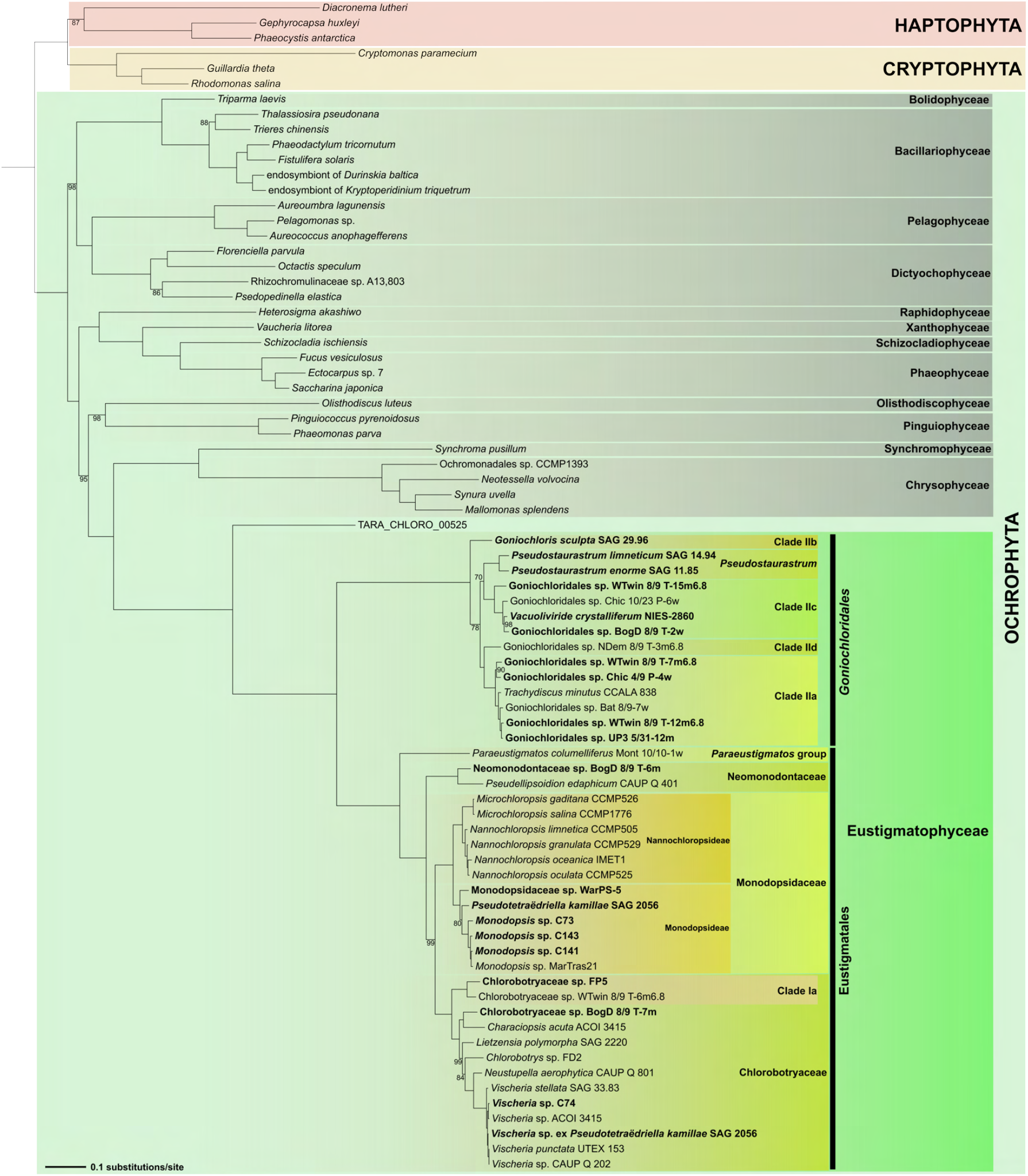
Maximum likelihood phylogeny of ochrophytes derived from a concatenated dataset of 69 plastome-encoded proteins. The tree was inferred from a supermatrix of 15,541 amino acid positions with IQ-TREE v3.0.1 and the LG+C60+F+G model. Sequences from haptophytes and cryptophytes were included as an outgroup. Branch support values (ultrafast bootstraps calculated from 10,000 replicates) are indicated only if lower than 100. Newly sequenced or assembled plastid genomes are highlighted in bold. Accession numbers of all plastome sequences included in the analysis are provided in Table S2.

In contrast to plastome sequences, the utility of mitochondrial genome data for the reconstruction of the eustigmatophyte phylogeny has not been properly tested before. The previous analyses (Ševčíková et al. 2016; Di Franco et al. 2022) were limited by both a poor taxon sampling and the reliance on site-homogeneous substitution models that may not adequately capture the complexity of the sequence evolution of mitochondrial genes. Indeed, the results of these previous analyses were strikingly incongruent with the eustig phylogeny as established based on other markers including plastid genes, in particular regarding the position of the *Vischeria* lineage unexpectedly placed sister to all other eustigs. We hypothesized that this unexpected result may be an artefact stemming from the *Vischeria* mitochondrial gene sequences being noticeably more divergent than those from other eustigs sampled at that time, and wondered if a new analysis, benefiting from the vastly improved representation the eustig diversity and employing a more sophisticated methodology of phylogenetic inference, would provide a different outcome. To this end, we exploited conserved mitochondrial genes from 42 eustigmatophytes and 30 other stramenopiles, built a supermatrix comprising 5,463 reliably aligned amino acid positions derived from 26 genes, and inferred a ML tree by using the site-heterogeneous LG+C60+L+G model.

The topology of the new mitogenome-based tree is indeed much more congruent with the results of the plastome-based analyses (Fig. S3). Crucially, the split of the Eustigmatophyceae into the monophyletic Eustigmatales and *Goniochloridales* is recovered in the mitogenome-based tree, with the two principal clades receiving very high and full statistical support, respectively. The topology within Eustigmatales follows the grouping of the taxa into three families and the stand-alone genus *Paraeustigmatos* as established by single-gene and plastid phylogenomic analyses, but the branching order of the four lineages differs, with Neomonodontaceae forming a moderately supported sister lineage to *Paraeustigmatos* rather than being sister to Chlorobotryaceae and Monodopsidaceae combined. The branching order within Chlorobotryaceae also shows some differences from the plastome phylogenomic analysis, particularly regarding the position of *Lietzensia polymorpha*. Additional topological differences between the trees inferred from the different organellar genomes concern the branching order among the closely related species/strains of the genera *Vischeria* and *Monodopsis*. Regarding the relationships within *Goniochloridale*s, the monophyly of the clades IIa and IIc and of the genus *Pseudostaurastrum* is preserved in the mitogenome-based analysis, but the branching order of these clades and the other (single-taxon) lineages differs from the plastid phylogeny.

In general, the partial incongruence of the plastome- and mitogenome-based inferences of the eustig phylogeny concerns branches that are not fully supported in one or both analyses, and may have various causes, both technical (insufficient signal in the data used, especially in the small mitochondrial dataset; model misspecification) and biological (truly incongruent evolutionary histories due to introgression or incomplete lineage sorting). Still another question is the different position of Eustigmatophyceae among ochrophytes in the plastome- and mitogenome-based phylogenies, which was noticed before by Di Franco et al. (2022). In addition, previous analyses have also documented a strong incongruence in the position of eustigs among ochrophyte lineages when the results of plastome-based analyses are compared with those of phylogenomic analyses employing nuclear genes (Di Franco et al. 2022; Cho et al. 2024; Terpis et al. 2025). At this stage, we refrain from suggesting the reasons for these discrepancies and admit that their resolution will require additional data, including comprehensive sequence data from various uncultivated MOCH lineages. Employment of the recently developed computationally demanding complex substitution models, as demonstrated, e.g., by the recent plastome phylogenomics study by Jamy et al. (2025), is also warranted. For the purpose of interpreting the history of changes of the gene content of eustig organellar genomes, we cautiously use as the reference tree a strict consensus of the plastome- and mitogenome-based topologies of the eustig subtree. However, we note that allowing multifurcations fortunately has little impact on the specific conclusions drawn.

### An updated view of the evolution of the plastid gene content in eustigmatophytes

Annotation of the newly obtained plastome sequences and their comparison with the previously analyzed ones revealed that eustigmatophyte plastomes are generally very stable in terms of their gene content (Table S2). The relatively minor differences reflect differential and sometimes recurrent loss of some of the genes in different eustig lineages, with a minimal contribution of gene gain (Fig. 3). Notably, the newly sequenced plastomes did not reveal any gene gains in addition to those described before (Ševčíková et al. 2019; Barcytė et al. 2022). The new data also cemented the notion that the *ebo* operon was acquired by the common ancestor of Chlorobotryaceae and Monodopsidaceae, with subsequent recurrent loss in both lineages. We see no obvious correlation between the distribution of the *ebo* operon and the specific features or ecology of different eustigs, leaving the biological role of the *ebo* operon in eustigmatophytes as enigmatic as before.

**Fig. 3.**
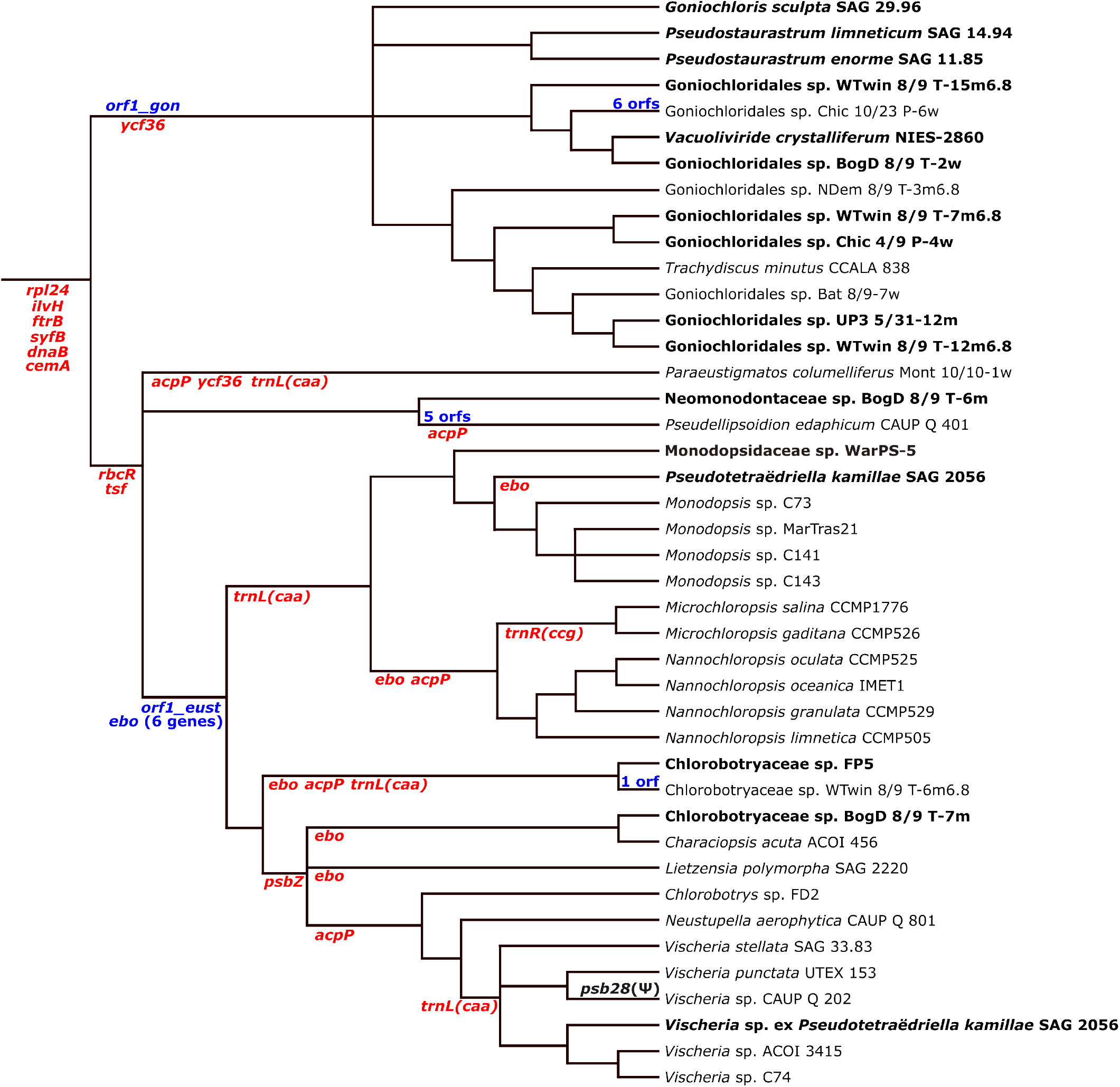
Major events in the evolution of the gene content of eustigmatophyte plastomes. The tree topology displayed is a consensus of the trees reconstructed by phylogenomic analyses of plastome- and mitogenome-encoded proteins (see Figs. 1 and 2, respectively). Gene loss (red) and gain (blue) events are mapped onto the phylogeny based on parsimony reasoning, i.e. minimizing the number of events needed to explain the occurrence of the genes in extant taxa. The six genes mapped as lost before the radiation of eustig lineages correspond to those that are present in at least some plastomes of the well-characterized closely related group, i.e. Chrysophyceae, yet are not found in any eustig plastome. One of them (*dnaB*) was undoubtedly lost in the eustig stem lineage, but the other five are also missing in the incomplete plastome sequence TARA_CHLORO_00525 representing an uncultivated sister group of Eustigmatophyceae (not included in the figure; see the main text). Hence, their loss may have occurred before the split of these two lineages. The Greek letter Ψ at the gene psb28 in one of the Vischeria strains indicates the gene sequence is present in the plastome but presumably pseudogenized by a frame-shift mutation.

Apart from the *ebo* operon, the eustigmatophyte plastid genomes have been previously shown to contain other genes that seem to represent gains specific for the whole group or its particular lineages. One of the genes, found in all eustig plastid genomes sequenced at that time, was denoted *ycf95* (Ševčíková et al. 2019). The expanded sampling reported here documents this gene to be ubiquitous in eustigs, which suggests that it has functional significance. We leveraged on the expanded sampling of different Ycf95 sequences to illuminate its origin, and as detailed in Note S1, we obtained compelling hints suggesting that it is most likely an extremely divergent evolutionary derivative of the common plastidial gene *ycf35*. Consistent with this notion, *ycf35* is readily identifiable in the aforementioned plastome from the uncultivated marine relative of eustigmatophytes (TARA_CHLORO_00525). Hence, instead of getting lost, *ycf35* was transmogrified into *ycf95* in the eustig stem lineage.

Two other plastid genes, denoted *orf1_eust* and *orf1_gon*, were previously found to be conserved in a subset of Eustigmatales and in *Goniochloridales*, respectively, yet lacked any discernible homology in other organisms (Ševčíková et al. 2019). Our expanded sampling showed that *orf1_eust* is present in all members of the families Chlorobotryaceae and Monodopsidaceae analyzed, but it is missing from the two more basally branching lineages of the Eustigmatales (Table S2). These results solidify the notion that *orf1_eust* emerged within Eustigmatales after the divergence of the two basal-most lineages, and once established it remained an essential part of the plastid gene set. Similarly, *orf1_gon* proved to be ubiquitous in *Goniochloridales* (Table S2; Dataset S1). Unfortunately, no new insights into the origin, and hence a possible function, of *orf1_eust* and *orf1_gon* could be gleaned, and the same negative conclusion pertains to the non-conserved orfs annotated in some of the previously analyzed eustig plastomes (for further details see Note S1).

The newly obtained plastid genome data also offer an opportunity to re-evaluate the previously recorded notable features concerning individual plastid genes in different eustig lineages. For example, *Pseudellipsoidion edaphicum* CAUP Q 401 was found to have the gene *sufB* divided into two parts, which presumably are translated independently to reconstitute the “complete” SufB protein by a non-covalent interaction of a short separate N-terminal domain with the main part of the protein (Ševčíková et al. 2019). Sequencing the plastid genome of a second member of Neomonodontaceae unveiled an intact *sufB* gene in this alga, indicating that the *sufB* split is specific for *P. edaphicum* CAUP Q 401 and presumably its hitherto unsampled specific relatives within the Neomonodontaceae family. The putative pseudogenization of *psb28* previously reported for *Vischeria* sp. CAUP Q 202 (Yurchenko et al. 2016) remains restricted to this particular strain, as the gene is apparently intact in plastid genomes of the five additional *Vischeria* strains analyzed here.

The *psb28* pseudogenization might be viewed as the first step towards the loss of the gene in the future descendants of *Vischeria* sp. CAUP Q 202. Indeed, gene loss has sculpted the gene content of eustig plastomes on a broader scale. Four genes (*rpl24, ilvH, ftrB, syfB*) were previously inferred to have been lost in the eustig stem lineage (Ševčíková et al. 2019). None of these genes is found in any of the newly sequenced eustig plastomes, but they are also missing from the metagenome-derived plastome sequence TARA_CHLORO_00525. Hence, their loss might have happened before the emergence of the eustig stem lineage, but this provisional conclusion is awaiting confirmation after complete plastome sequences of the uncultivated marine sister group of eustigmatophytes become available. On the other hand, the TARA_CHLORO_00525 plastome possesses the *dnaB* gene absent from eustig plastomes, which implies its loss in the eustig stem lineage (Fig. 3). This is a revision of the previous evolutionary interpretation placing the *dnaB* loss before the divergence of eustigs and chrysophytes (Ševčíková et al. 2019), consistent also with the encounter of the gene in some of the more recently sequenced chrysophyte plastomes (Kim et al. 2019). The *ycf35* gene, also previously hypothesized to have been lost before the eustig-chrysophyte divergence (Ševčíková et al. 2019), now appears to have persisted in eustigs as the gene annotated as *ycf95* (see above).

The previously reconstructed history of plastid gene loss in eustigs pointed to unique loss events specific for particular eustig clades, as well as to recurrent loss of certain genes in different eustig lineages (Ševčíková et al. 2019). With the expanded sampling we here provide a more comprehensive picture of these loss events (Fig. 3). Perhaps most notable is the finding that no gene loss specific for a particular lineage of *Goniochloridales* was detected despite the increase in the number of plastid genomes from four to fourteen. Thus, except for the emergence of the six non-conserved *orfs* in the strain Chic 10/23 P-6w (Ševčíková et al. 2019), the plastid gene repertoire is constant across the whole *Goniochloridales* radiation and differs from the ancestral eustigmatophyte plastid gene set only due to loss of *ycf36* and gain of *orf1_gon* in the *Goniochloridales* stem lineage. The evolutionary stasis of the plastome in *Goniochloridales* contrasts with a more dynamic plastome evolution in Eustigmatales. The expanded plastome sampling reinforced the previous inference that the loss of both the *rbcR* and *tsf* genes is synapomorphic for Eustigmatales lineages, whereas additional losses occurred only in some of the lineages. Some of the losses are unique, i.e. restricted to a single lineage, but the genes *acpP* and *trnL*(*caa*), as well as the whole *ebo* operon, were lost at least four or five times independently.

We previously analyzed evolutionary adaptations potentially compensating for the loss of particular plastid genes in eustigs, with the most notable outcome being the realization that the recurrent loss of the *acpP* gene is apparently enabled by the existence of a nuclear gene for a plastid-targeted acyl carrier protein that was acquired by the eustigmatophyte common ancestor from the *Phycorickettsia* lineage (Ševčíková et al. 2019). However, in some other cases no putative compensatory solutions that would explain the respective losses could be determined. One of these cases was the *tsf* gene, which encodes a homolog of the bacterial EF-Ts protein, i.e. a regulator (guanine nucleotide exchange factor) of the protein EF-Tu functioning as an essential elongation factor in plastidial translation (Tiller and Bock 2014). Its absence in Eustigmatales (Fig. 3) is thus surprising. Here we demonstrate that all Eustigmatales possess two different nuclear *tsf* homologs, one encoding a protein with a predicted mitochondrial targeting signal and the second coding for a product with an N-terminal extension with features characteristic for the bi-partite plastid-targeting presequence characteristic for ochrophyte algae (Gruber et al. 2025). In contrast, *Goniochloridales* exhibit only one nuclear *tsf* copy that corresponds to the mitochondrial version (Table S5). Phylogenetic analysis indicates that all the eustig nuclear *tsf* genes are monophyletic and apparently correspond to a common eukaryotic gene encoding the mitochondrial EF-Ts protein (Fig. S4). Therefore, the *tsf* gene was most likely duplicated in the common Eustigmatales ancestor and one copy became neofunctionalized, i.e. was retargeted to the plastid, allowing for the loss of the plastidial *tsf* gene. Recruiting a protein originally serving in the mitochondrion for a plastidial function is not unprecedented, as exemplified by the plant plastidial RNA polymerase NEP originated from a duplicated copy of a mitochondrial RNA polymerase (Liere et al. 2011), or the organellar genome-replicating DNA polymerase POP originally serving the mitochondrion only but dually targeted also into the plastid in most plant and algal taxa (Harada et al. 2024).

### Identification of the eustigmatophyte orfX and orfY as genes for mitoribosomal proteins

Despite their smaller size compared to the plastid genomes, our analyses demonstrate that the eustigmatophyte mitogenomes have experienced a more eventful evolutionary history including gene loss, gain, duplications, and fusions. Some of the events were noted in the previous detailed analysis of a smaller number of eustigmatophyte mitogenomes (Ševčíková et al. 2016). The enlarged sample of mitogenomes analyzed in this study provides a fuller and finer picture (Figs. 4 and 5). For example, the previously reported unique loss of the *tatA* gene in a *Vischeria* species can now be mapped to a deeper internal branch within Chlorobotryaceae, indicating a simplification of the mitochondrial TAT translocase in the whole broader descendant clade comprising at least three genera.

**Fig. 4.**
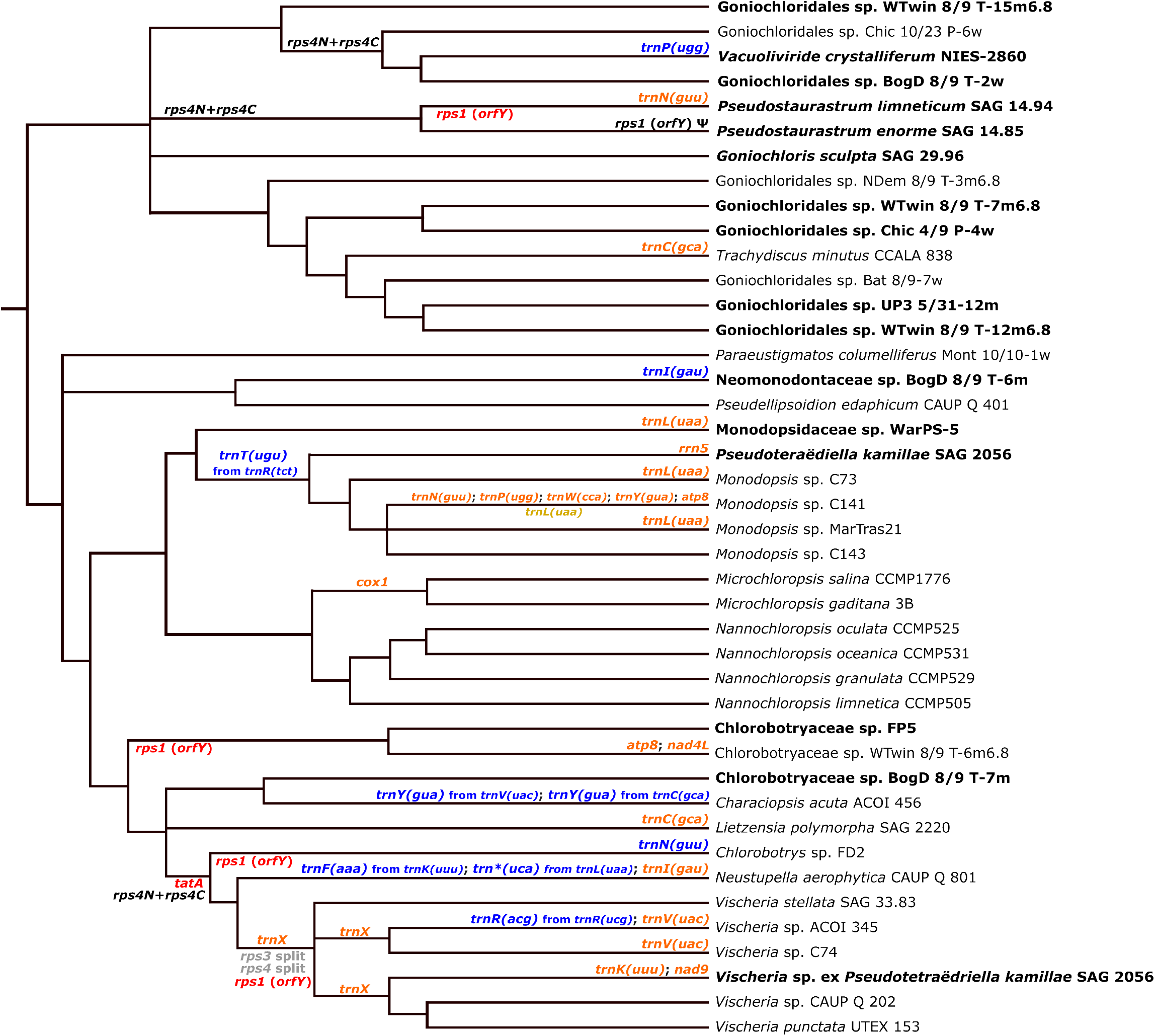
Major events in the evolution of the gene content of eustigmatophyte mitogenomes. The tree topology displayed is a consensus of the trees reconstructed by phylogenomic analyses of plastome- and mitogenome-encoded proteins (see Figs. 2 and S3, respectively). The evolutionary journey started with the ancestral eustig mitogenome comprising at least 73 genes (Table S3). All events inferred to have changed this ancestral state are mapped onto the phylogeny based on parsimony reasoning, i.e. minimizing the number of events needed to explain the occurrence of the genes in extant taxa. These events consider gene loss (red), gain (blue), duplication (orange), and triplication (gold). Gene fusion (black and the “plus” sign) and split (grey) concern only the *rps4* and the *rps3* gene, respectively. The Greek letter Ψ indicates putative pseudogenization. Note that for the sake of clarity gains and losses of patchily distributed eustig-specific non-standard *orfs* (*orfN* to *orfW* plus *orfZ*) are not included in this scheme but are displayed separately as Fig. 5.

**Fig. 5.**
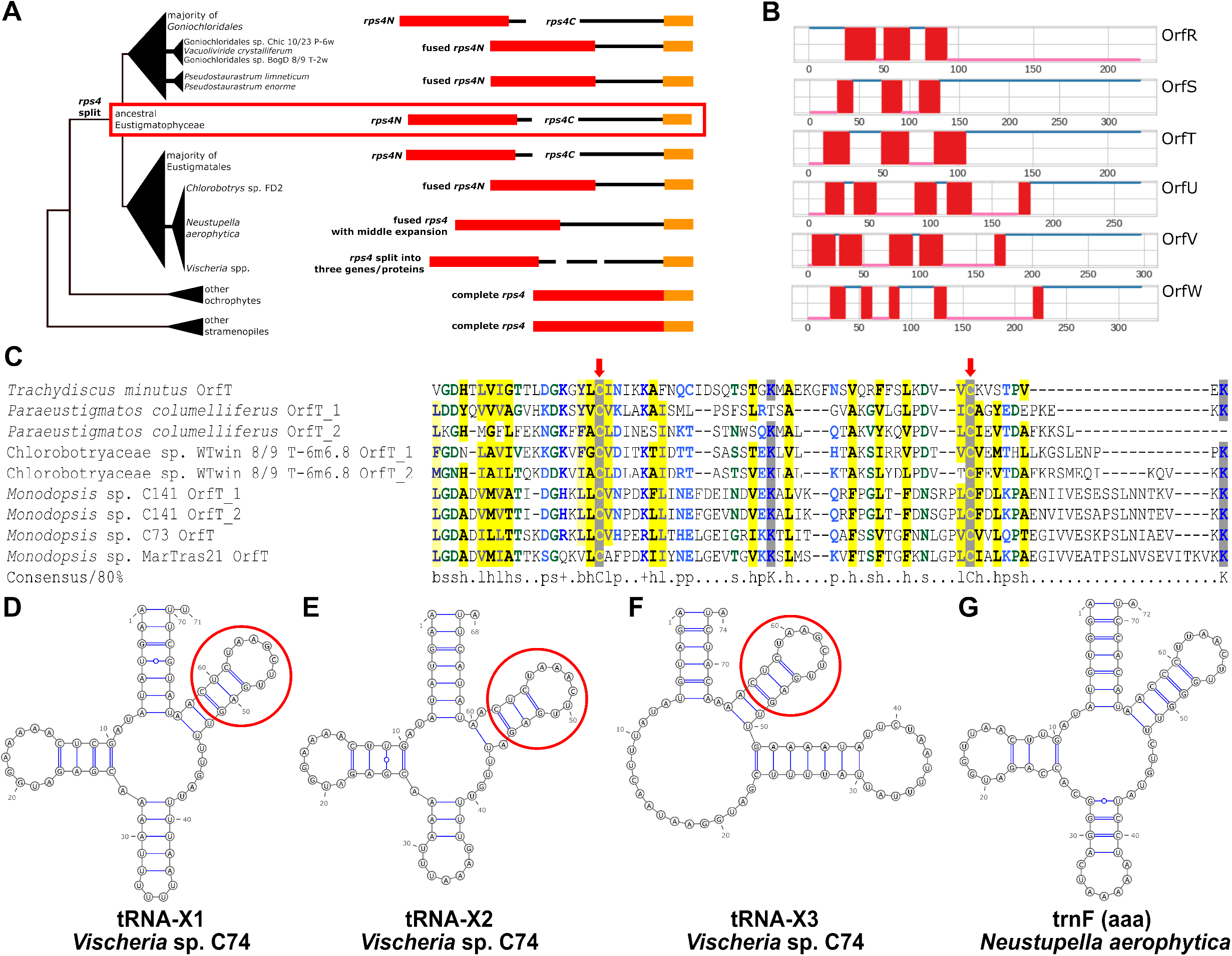
Occurrence and putative evolutionary history of the conserved non-standard *orfs* (*orfN* to *orfW* plus *orfZ*) in eustigmatophyte mitogenomes. The specific combination of the *orfs* in individual extant species/strains is indicated by the respectively colored circles at terminal branches. Origins (the presence in the common eustigmatophyte ancestor or lineage-specific gains), losses (crossed circles), and pseudogenizations (the Greek letter Ψ inside the circle) of the *orfs* as inferred from their occurrence in extant eustigs (without assuming the possibility of HGT) are mapped onto the consensus phylogeny of the group. The pseudogenes range from cases of a recent disruption of the coding sequence (by in-frame termination codons or frameshifts) to only short fragments left. For simplicity, duplication events that likely explain the presence of multiple copies of *orfT, orfV* and *orfW* in some eustigs (indicated by numbers inside the respective circles at the terminal branches), and potential subsequent differential losses of paralogs are not mapped, as their accurate inference is difficult owing to low resolution and perhaps limited accuracy of phylogenetic analyses of the *orfs*. The figure also omits cases of pseudogenes co-existing in the same mitogenome with an intact copy of the respective *orf*.

An important update also concerns the occurrence of introns in eustig mitogenomes, previously restricted to a group IIA intron in the *cox1* gene in *Nannochloropsis oculata* CCMP 525 (Starkenburg et al. 2014; Ševčíková et al. 2016). The newly sequenced mitogenomes now uncovered six taxa from multiple major eustig lineages whose *rnl* gene (specifying 23S rRNA) is colonized by one or two group I introns (Table 1). This amounts to nine introns occupying three different conserved positions in the *rnl* gene, referred to as sites 1931, 1951, and 2262 following the standard nomenclature based on the *rnl* gene of *Escherichia coli* as a reference. All nine introns encode a homing endonuclease of the LAGLIDADG family, i.e. enzymes mediating mobility of the introns encoding them (Liu et al. 2026). The intron-containing eustigs are generally not related to each other, and the pattern of the occurrence of particular introns does not generally reflect the relative phylogenetic distance of the taxa containing them. Hence, each of the nine identified introns may potentially represent a separate acquisition event (as indicated in Fig. 4), unless multiple secondary losses of the introns are considered. It is notable that none of the non-eustig ochrophyte mitogenomes included in our phylogenomic analysis has an intron at any of the three *rnl* sites, and the homing endonucleases encoded by the eustig *rnl* introns typically retrieved as best BLASTP hits proteins encoded by introns in green algal organellar genomes. The eustig *rnl* introns thus might have been acquired from green algal sources, but further analyses would have to be carried out to clarify the actual direction of transfer and also to test for the possibility that the introns moved directly between different eustig lineages.

An important outcome of the improved eustig mitogenome sampling is the clarification of the identity of two mitochondrial genes previously found to be conserved across eustigs and annotated *orfX* and *orfY* for the lack of discernible homologs in other organisms (Ševčíková et al. 2016). In the case of *orfX* we noticed that the gene occurs in all eustig mitogenomes sequenced so far and is virtually always located directly downstream of the *rps4* gene, which encodes a component of the ribosomal small subunit (the S4 protein). The exceptions are the genus *Vischeria*, in which the two genes are separated by a short *orf* (see below), and four other separate eustig lineages, in which the open reading frames of *rps4* and *orfX* genes are continuous, i.e. presumably encoding a single fused protein (Table S3). These findings suggested that OrfX is functionally linked to the mitoribosomal S4 protein. We then used the multiple available OrfX protein sequences to construct a representative profile HMM and employ it in a HHpred search, which matched the C-terminal region of OrfX to the C-terminal region of the S4 protein from organisms outside eustigmatophytes, although with low statistical significance (Fig. S5A). Analogous investigation of the protein sequences encoded by the eustig *rps4* gene revealed that they match most of the length of standard S4 proteins, except for their C-terminal region (Fig. S5B).

Based on these observations, we propose that tandem duplication of the *rps4* gene occurred in a eustig ancestor, followed by subfunctionalization of the two copies (Fig. 6A). One of the copies, now reannotated as *rps4N*, has diverged in the very 3’ region, resulting in the divergence of the C-terminal part of the encoded protein from the standard S4 homologs. The other copy, originally called *orfX* but now reannotated as *rps4C*, has complementarily preserved the 3’ region, so that the encoded protein exhibits a C-terminal segment homologous to the C-terminal part of the standard S4 protein. The *rps4C* gene has diverged extremely along the rest of its span, resulting in a “novel” protein sequence that is relatively conserved among eustigs but with no discernible homology outside the group. As detailed in Note S2, the *rps4N* and *rps4C* gene must have fused multiple time independently, with evidence for splitting secondarily in the genus *Vischeria*.

**Fig. 6.**
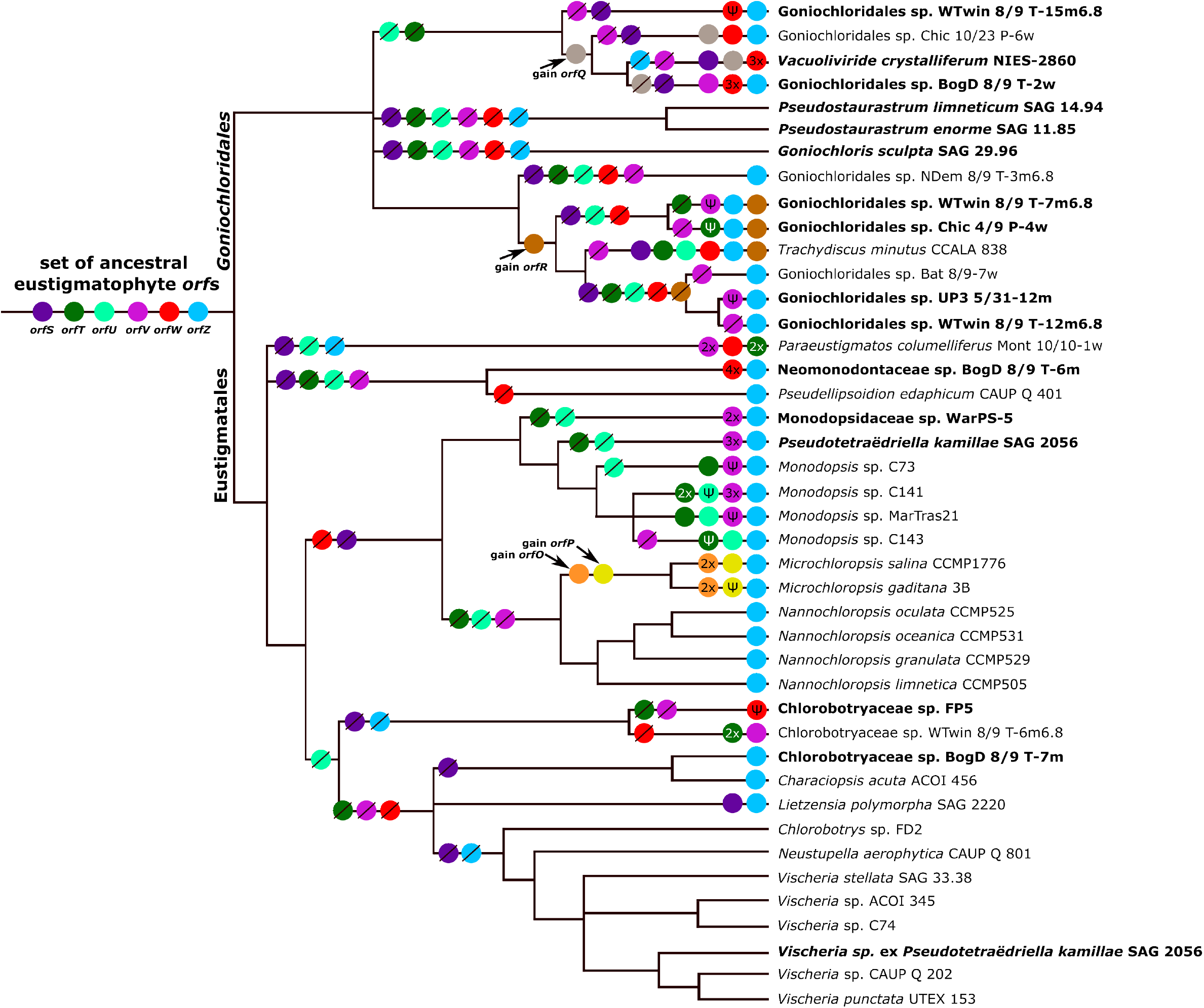
Unique features of selected eustigmatophyte mitochondrial genes. (A) Schematic representation of the reorganized *rps4* gene in eustigmatophytes. Generally (and ancestrally), the gene is split into two parts, one (now annotated as *rps4N*) matching a nearly full standard S4 protein except for the very C-terminal part. The other, previously denoted *orfX* but now reannotated as *rps4C*, consists mostly of a novel sequence (black line) except for the C-terminal region (red rectangle) that matches the C-terminal region of standard S4 proteins. In a few unrelated eustigs *rps4N* and *rps4C* have independently fused to a single ORF. The middle part of the fused gene expanded significantly in *Neustupella aerophytica* and the whole process culminated in genus *Vischeria* where the *rps4* gene is divided into three separate ORFs (*rps4N, rps4M* and *rps4C*). Intact *rps4* gene structure in other stramenopiles is shown as a reference. (B) The presence of predicted transmembrane helices in the N-terminal regions of representative proteins encoded by the conserved non-standard mitochondrial genes *orfR* to *orfW*. Proteins from *T. minutus* are shown, except for OrfV (missing in this species), which comes from *Pseudotetraëdriella kamillae*. A similar architecture is generally exhibited by homologs from other eustigs (the full set included in Dataset S3). (C) Multiple sequence alignment of the conserved C-terminal soluble domain of OrfT proteins. Note the two absolutely conserved cysteine residues (red arrows). (D-G) Predicted secondary structures of selected non-standard eustigmatophyte tRNAs evolutionarily derived from a duplicated *trnK(ttt)* gene. Panels D to F show all three such tRNAs, denoted tRNA-X, from *Vischeria* sp. C074 modeled using Rfam (D and E) or RNAfold (F). Structures A and B display slightly atypical tRNA configurations (mainly in the anticodon arm and loop) but still resembling standard tRNA structure, whereas the secondary structure of tRNA-X3 (panel E) is extremely modified. All three tRNAs differ substantially from standard tRNAs in their anticodon arm (and usually along other parts of the sequence, see Fig. S10), which may indicate that they no longer play a role in translation. In contrast, the T-arm (including the T-loop) is highly conserved in all these tRNAs (red circle), pointing to its specific functional significance. Panel G shows the structure (modeled by Rfam) of the putative product of a novel tRNA gene from a close relative of *Vischeria* that is annotated as *trnF(aaa)* (the amino acid specificity solely reflects the anticodon sequence) but also clearly evolved from a *trnK(ttt)* copy, potentially as a result of a gene duplication event preceding the *Neustupella-Vischeria* split (see main text and Fig. S11).

While the ubiquity of *orfX* (=*rps4C*) in eustig mitogenomes indicates that this gene encodes an essential component of the molecular machinery of the organelle, the *orfY* gene was previously found secondarily missing from *Vischeria* sp. CAUP Q 202 (Ševčíková et al. 2016). We now understand that *orfY* was lost in the common ancestor of the genus *Vischeria* and independently in three other eustig lineages. Furthermore, the 5’ end of *orfY* in *Pseudostaurastrum enorme* is divergent and truncated, suggesting that the gene is degrading and potentially non-functional (Fig. 4; Table S3). Following the procedure that illuminated the origin of *orfX*, we found evidence for *orfY* most likely representing an unrecognized divergent homolog of one of the standard mitochondrial genes, namely *rps1*. Firstly, a HHpred search with a comprehensive alignment of *orfY*-encoded proteins as a query retrieved only two hits, both with low probability values / high e-values, yet both notably representing the ribosomal protein S1 (Fig. S6). Secondly, although modeling the proteins encoded by *orfY* from different eustigs using AlphaFold 3 generally resulted in low-confidence prediction, the few predicted with more robust statistics consistently retrieved bacterial or mitochondrial S1 proteins as best hits in Foldseek searches, often with convincing statistical support (probability values up to 1.0, E-values as low as 6.98e-8; Table S6). A survey of the mitochondrial gene complement in eukaryotes had indicated a patchy distribution of the *rps1* gene, with homologs detected only in a few distantly related protist lineages (Butenko et al. 2024), whereas among all Stramenopiles the mitochondrial *rps1* has been recorded only in pelagophytes (Ševčíková et al. 2016; Sibbald et al. 2021). The identification of the eustig *orfY* as *rps1* suggests that unrecognized divergent *rps1* orthologs may occur more commonly in mitogenomes of other ochrophytes or eukaryotes in general.

One more mitochondrial gene of an unknown origin and function, annotated as *orfZ*, was previously detected in eustigs except for a *Vischeria* species (Ševčíková et al. 2016). Our present sampling indicates six independent *orfZ* losses (three times in both the Eustigmatales and the *Goniochloridales*; Fig. 5; Table S3). In contrast to *orfX* and *orfY, orfZ* proved recalcitrant to our attempts to decipher the origin using HHpred searches or structural modeling with AlphaFold 3 followed by a Foldseek search, as these approaches failed to reveal any significant *orfZ* homologs. Nevertheless, the presence of two predicted transmembrane (TM) helices in *orfZ* products was previously reported by Ševčíková et al. (2016), and confirmed here to be a consistent feature of the whole ensemble of homologs (Fig. S7). This observation suggests that it is unlikely *orfZ* would represent any of the two standard mitochondrial genes known to occur in at least some ochrophytes yet apparently missing from eustigs, i.e. *rpl10* and *rpl31* (see the data presented in Barcytė et al. 2024), as the ribosomal proteins encoded by these are not expected to have transmembrane helices.

### A surprisingly diverse set of patchily distributed but potentially ancestral eustigmatophyte mitochondrial orfs

Two of the previously analyzed eustig mitogenomes were found to contain additional non-standard *orf*s, but these were considered to be taxon-specific features given the lack of discernible homologs in other organisms including the eustigs available for comparison (Ševčíková et al. 2016). Strikingly, our analyses of the expanded set of eustig mitogenomes revealed that some of them do have homologs in other eustigs, sometimes even distant relatives spanning the Eustigmatales-*Goniochloridales* split. To better understand the space of eustig-specific mitochondrial *orfs*, we gathered the corresponding putative protein sequences from all eustigs analyzed and clustered them based on pairwise sequence similarity using CLANS (Fig. 7). Twenty-nine *orfs* were classified as genome-specific singletons, as they did not join any cluster, and two clusters each corresponded to a pair of loci from the same mitogenome, most parsimoniously interpreted as a genome-specific singleton duplicated in the history of the given species/strain. These putative genes were annotated as “*orfXXX*” with the “*XXX*” referring to the number of encoded amino acids (Table S7). The other mitochondrial non-standard *orf*s were grouped into nine clusters (other than those corresponding to *orfX* to *orfZ*), with each cluster comprising sequences from multiple eustig species/strains. Two of the clusters were loosely connected in the CLANS analysis, but as their taxonomic composition overlapped rather than being complementary, we considered them to represent separate yet potentially homologous genes. Hence, we annotated the non-standard *orfs* encoding members of the nine clusters as *orfO* to *orfW* (Table S6).

**Fig. 7.**
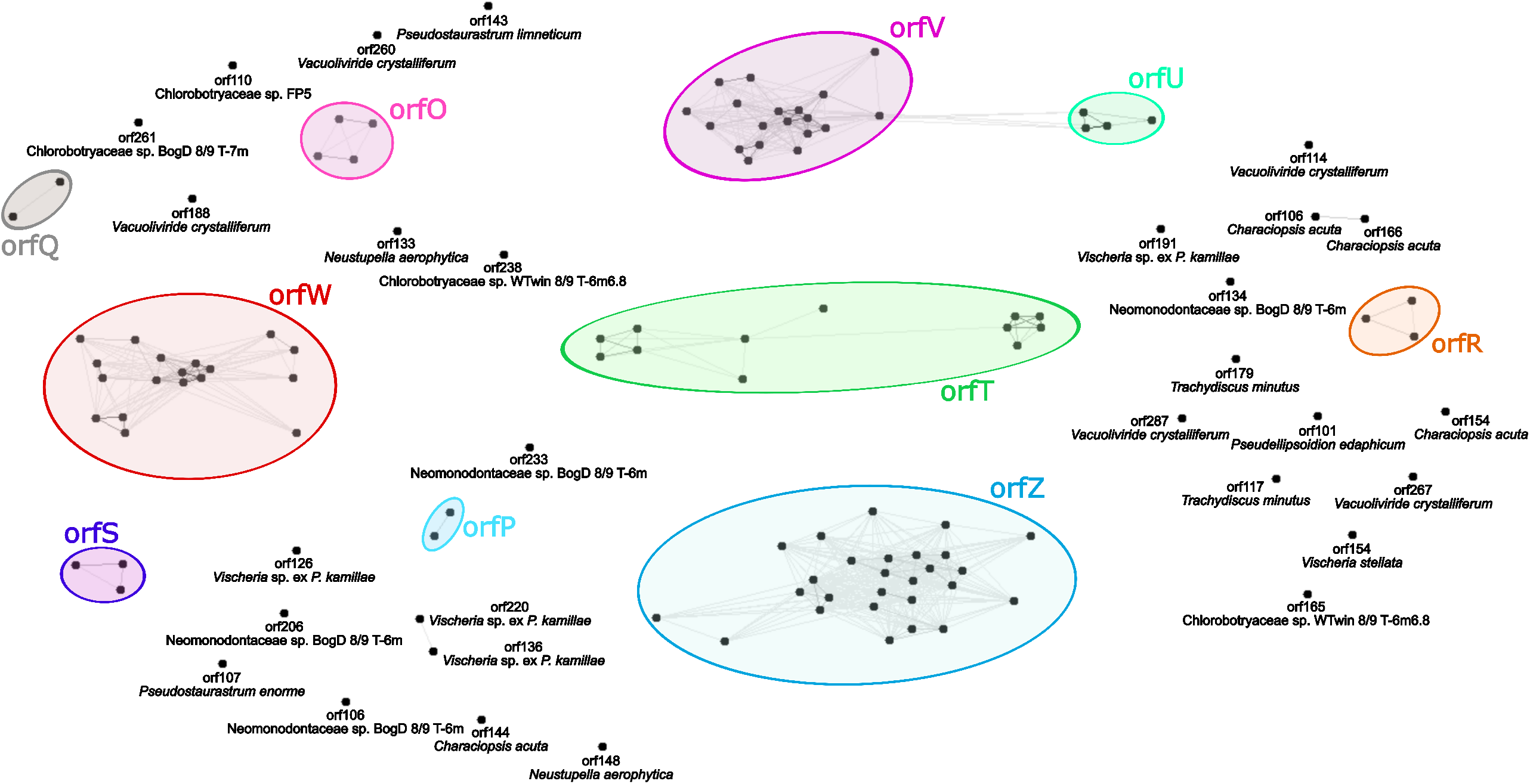
Clustering analysis performed with the MPI Bioinformatics Toolkit CLANS (for details see Material and Methods) showing eleven clusters of *orfs* present in eustigmatophyte mitogenomes (in colors) and other *orfs* without any homologs in other eustigmatophytes (black). Colors correspond to those used in the Fig. 5.

We then looked for possible homology of all the putative products of non-standard *orfs* to other proteins using HHpred searches against available collections of profile HMMs and by BLASTP searches against a collection of all eustig mitogenome-encoded proteins, with the complete results listed in Tables S6 and S7. The conceptual translations of a few singleton *orfs* exhibited short regions that were highly similar to some of the standard mitochondrial proteins or that retrieved these proteins as weakly supported homologs. These *orfs* thus most likely evolved from duplicated copies of standard genes or fragments thereof. No homology to other proteins was discernible for products of virtually all other singleton *orfs*, making their origin elusive, but it is notable that some of them contain predicted TM helices (Table S7). The only exception was an *orf* identified as an apparently non-functional fragment of a phage-derived RNA polymerase gene, most likely representing a chance insertion of a broader DNA region into the *V. crystalliferum* mitogenome (Fig. 8A; for further details see Note S3).

**Fig. 8.**
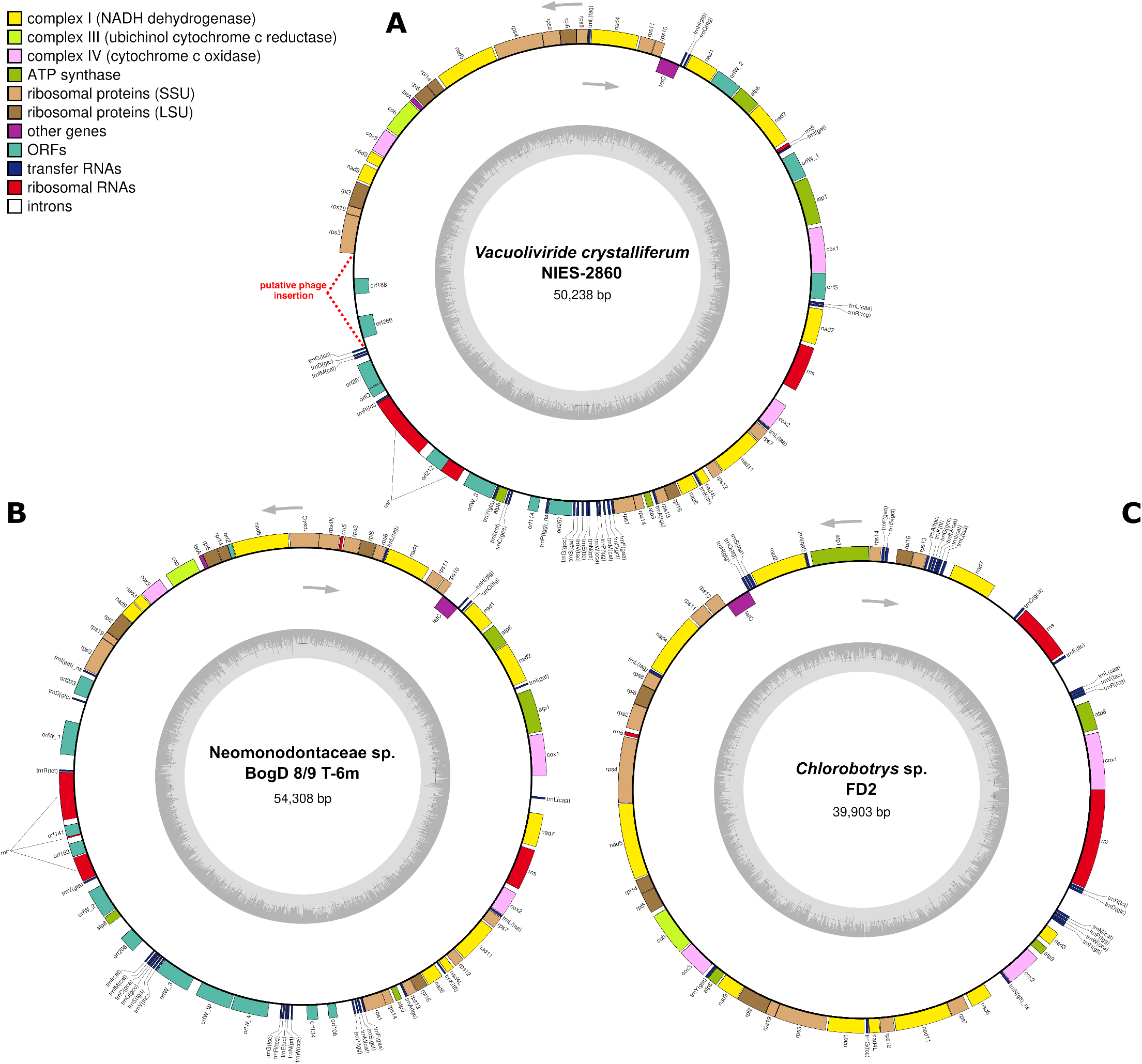
Mitochondrial genome maps for selected eustig representatives showing unusual genome organization, namely the presence of an insertion putatively derived from a phage in Vacuoliviride *crystalliferum* (A), a high number of unique *orfs* in Neomonodontaceae sp. BogD 8/9 T-6m (B), and the complete lack of eustig-specific non-standard *orfs* in *Chlorobotrys* sp. FD2 (C). Note that in addition to *orf260* identified as a divergent fragment of a phage-type RNA polymerase gene, the putative phage-derived insertion in the *V. crystalliferum* mitogenome also potentially includes the adjacent *orf188*, which is likewise located on the opposite strand compared to virtually all other genes in the genome.

The availability of multiple homologous sequences representing the *orfO* to *orfW* genes enabled us to use profile HMMs constructed from multiple protein sequence alignments, rather than individual sequences, as queries for HHpred searches. Despite the presumed higher sensitivity of such searches, no convincing candidate homologs were retrieved for any of the queries. No credible results regarding possible homology of the proteins of the OrfO-to-OrfW series to other proteins were obtained even by modeling their structure and querying the models with Foldseek (Table S6). The origins of these *orfs* are, therefore, mysterious. Four of them, specifically *orfO* to *orfR*, are each restricted to a different narrow eustig clade, but *orfS* to *orfW* each occur in at least one representative of both the Eustigmatales and the *Goniochloridales*. To further test for possible homology between any of the non-standard conserved *orfs*, especially whether any of the *orfO* to *orfR* genes are in fact divergent offshoots of any of the broadly occurring *orfS* to *orfW* genes, we used HHpred and multiple protein sequence alignments of representatives of each of the ten clusters to perform all possible pairwise profile HMM-HMM comparisons. However, besides additional support for the relationships between *orfU* and *orfV* (Fig. S8) already suggested by the CLANS analysis (Fig. 7), we did not detect any convincing evidence for additional relationships. Given the evidence in hand, the *orfO* to *orfR* group represents *de novo* genes each evolved in the common ancestor of a particular narrow eustig clade, whereas *orfS* to *orfW* are candidates for five different genes present already in the last eustigmatophyte common ancestor (Fig. 5).

Although we could not infer the origin and function of OrfO to OrfW proteins, some interesting insights could be obtained by considering their generic sequence features and evolutionary dynamics; details are provided in Note S4. Most notably, these proteins have predicted TM helices, and the number and location of the helices is generally conserved across the particular group of homologs (Fig. 6B; Dataset S3; Table S6). Hence, we propose they are embedded in the inner mitochondrial membrane. The OrfR to OrfW proteins are furthermore characterized by putative C-terminal soluble domains (different between different Orf proteins) that protrude to the matrix or intermembrane space, but no specific clues could be gleaned regarding their biochemical activities. Nevertheless, in the case of OrfT proteins, we noticed a conspicuous sequence pattern in their C-terminal domain, which contains two absolutely conserved cysteine residues (Fig. 6C). We speculate that these residues form a disulfide bridge and/or are implicated in reversible redox regulation of the OrfT activity.

The pattern of occurrence of *orfR* in the *Goniochloridales* clade IIa members implies its loss from one of the subclades, whereas the presumed presence of *orfS* to *orfW* in the mitogenome of the last eustigmatophyte common ancestor implies multiple independent losses of each of the genes in different eustig lineages, from at least nine for *orfW* to minimally 15 for *orfV* (Fig. 5). The scenario of recurrent loss is supported by the observation that for *orfT, orfU, orfV*, and *orfW*, apparently non-functional fragments or pseudogenes are found in some eustig mitogenomes as vestiges of the past presence of complete versions of these genes (Fig. 5; Table S6). The ancestral status is less certain for *orfS* and *orfU*, which exhibit a highly restricted distribution, each *orf* being present in a single taxon of Eustigmatales (different for each *orf*), whereas in *Goniochloridales* they are present only in *Trachydiscus minutus* (both) and *Vacuoliviride crystalliferum* (*orfS* only). Furthermore, except for a recently pseudogenized *orfU* in *Monodopsis* sp. C141 due to a frameshift mutation, no remnants of these genes were found in other eustig mitogenomes. Combined with the high number of independent losses required by the vertical inheritance scenario (14 for *orfS* and 13 for *orfU*), alternative explanations assuming HGT between the mitogenomes or independent acquisition by HGT from a common non-eustig source cannot be dismissed. Furthermore, given the evidence for *orfU* being related to *orfV*, it is possible that *orfU* evolved in either the *Trachydiscus* or the *Monodopsis* lineage by high sequence divergence of a copy of *orfV* and was then transferred by HGT to the other lineage.

The shared general architecture of OrfR to OrfW proteins suggests that they may have all evolved from the same common ancestral gene by rounds of duplication and extreme sequence divergence. Given the details provided above, the emergence of *orfS* to *orfW* would precede the Eustigmatales-*Goniochloridales* split, whereas *orfR* either evolved by an additional duplication/divergence event later within *Goniochloridales* or may be as ancient as *orfS* to *orfW*, but missing from Eustigmatales secondarily. Regardless, the evolutionary dynamics of *orfR* to *orfW* is truly unique and differs from the evolutionary pattern exhibited by the standard mitochondrial protein-coding genes, as is underscored by additional observations. One feature pertinent to all of *orfR* to *orfW* is the apparently much higher rate of sequence evolution of the *orfs*, manifested by the orthologous sequences exhibiting a much lower sequence similarity compared to the standard genes. For example, when the proteins encoded by the mitogenomes of two related *Monodopsis* species were compared, virtually all pairs of standard mitochondrial proteins exhibited mutual sequence identity exceeding 90%, whereas for the pair of OrfT orthologs encode by the same mitogenomes this value was only 80%. Another striking aspect concerns *orfT, orfV*, and *orfW*, which tend to be present in multiple copies in the same mitogenome (Fig. 8B; Table S6). While some of the copies are just shorter or longer fragments of their full-length versions, two, three, or even four apparently intact copies may co-exist in the same organism. This feature further deepens the distinction of these *orfs* from standard mitochondrial protein-coding genes, whose functional duplications (i.e. co-occurrence of multiple copies without obvious signs of degeneration of any of them) is very rare in eustigs; the only exception being two copies of *cox1* in *Microchloropsis* spp. While this may mean that the accumulation of *orfT, orfV*, and *orfW* copies in the same mitogenome is driven by positive selection, an alternative explanation is that standard genes are more sensitive to gene dosage, which constrains their duplication.

To better understand the origin of the *orf* copies, we performed phylogenetic analyses of *orfT, orfV*, and *orfW* genes (Fig. S9). In some cases (e.g., both *orfT* versions and all three *orfV* versions in *Monodopsis* sp. C141), the copies clearly stem from duplication of the *orf* in a particular terminal eustig lineage, but in other cases the co-occurring versions do not branch together. This is best exemplified by *orfW* in *Goniochloridales* sp. BogD 8/9 T-2w, with one copy being closely related with strong support to homologs from other clade IIc members, whereas the other two copies branch together but far away from all other *orfW* genes from clade II representatives. At face value the trees inferred for the proteins encoded by *orfT, orfV*, and *orfW* would imply highly complex evolutionary histories of the genes including deep internal paralogy combined with massive differential loss of the paralogs, and possibly even dissemination by HGT as discussed in relation to *orfS* and *orfU*. Perhaps more realistically, the incongruence of the topologies with the species tree, which is generally poorly supported by the bootstrap analysis, may stem from a combination of stochastic errors and rapid and/or uneven sequence evolution of the *orfs*. Regardless of these uncertainties, the wide phylogenetic span of *orfS* to *orfW* suggests that these *orfs* possess (*bona fide*) biological functions, as holds also for the similarly enigmatic (yet much more broadly occurring) *orfZ*. Nevertheless, as documented by the existence of eustig mitogenomes completely devoid of these *orfs* (Figs. 5; Fig. 8C; Table S3), these functions are only accessory to the processes operating in eustig mitochondria.

### A refined picture of unusual aspects of the mitochondrial translation in eustigmatophytes

The split of the original *rps4* gene discussed above is just one of the evolutionary modifications of the eustig mitochondrial translation apparatus; a few others were pointed to by the previous taxonomically limited analysis (Ševčíková et al. 2016). One concerned the *rps3* gene in *Vischeria* sp. CAUP Q 202, which is interrupted by an in-frame termination codon UAA in a region corresponding to a poorly conserved internal part of the Rps3 protein. It was previously hypothesized that the codon is decoded by an unusual *Vischeria*-specific tRNA, denoted tRNA-X and characterized by an expanded anticodon loop, which enables the translation of both parts of the *rps3* gene as a single continuous coding sequence (Ševčíková et al. 2016). With a mitogenome sequence from five additional *Vischeria* species (strains), we could revisit this hypothesis. Three additional species exhibit the *rps3* gene interrupted with an in-frame termination codon (UAA or UAG) like *Vischeria* sp. CAUP Q 202. However, in *Vischeria* sp. ACOI 3415 and *Vischeria* sp. C074, the coding sequence corresponding to the full-length *rps3* gene is divided into two parts in different reading frames and furthermore separated by a region including two tRNA genes (Dataset S2). The latter two *Vischeria* strains are closely related according to our organellar genome phylogenies (Figs. 2 and S3), and their *rps3* organization may reflect a next evolutionary step after an in-frame termination codon interrupted the gene in the *Vischeria* stem lineage. These findings thus support an alternative previously discussed scenario (see Ševčíková et al. 2016) that the *rps3* gene in *Vischeria* is truly disrupted to make two separately translated parts encoding the N-terminal and the C-terminal domain of the ribosomal S3 protein as separate polypeptides. *Vischeria* thus parallels some other eukaryotes with a similar split of the mitochondrial *rps3* gene (Swart et al. 2012; Fu et al. 2014).

The revised interpretation of the interrupted *Vischeria rps3* gene begs the question about the actual functionality of the unusual “tRNA-X”. As detailed in Note S5, the expanded taxon sampling provided interesting insights regarding this molecular item, renamed tRNA-X1, as well as other related tRNA-like loci in the genus *Vischeria*, including one previously reported as “tRNA-Leu2” (Ševčíková et al. 2016) but now renamed tRNA-X2. The tRNA-X1 and tRNA-X2 loci are present in mitogenomes of all and most, respectively, *Vischeria* species and are predicted to specify tRNA-like molecules that vary highly in the anticodon arm regions but whose other parts are remarkably conserved within each group of orthologous loci (Fig. 6D-E; Fig. S10). In addition, we newly identified a third locus present in all *Vischeria* spp., denoted tRNA-X3, with the putative RNA products generally departing even more from the conventional tRNA structure (Fig. 5F; Fig. S10). The three unusual tRNA-like loci share the same origin and evolved by gene duplications from the commonly occurring *trnK*(*uuu*) gene (Fig. S11). They seem to be restricted to the genus *Vischeria*, although it is notable that the closely related eustig *Neustupella aerophytica* exhibits a single unusual *trnK*(*uuu*)-derived paralog that might evolutionarily correspond to the tRNA-X1/-X2/-X3 group (Fig. 6G; Fig. S11). The evolutionary conservation of the three tRNA-like loci indicates they are kept by selection and hence functional in some way, but the high variation in the anticodon arm, which in some cases is essentially completely missing, indicates the respective RNA molecules do not serve in codon decoding in the ribosomal A site. What might be the alternative molecular function of the products of these loci is presently unclear, but tRNAs or tRNA-like molecules specialized for participating in biosynthetic processes or having regulatory roles have been reported before (Su et al. 2020).

Other “extranumerary” tRNA genes exist in some eustig mitogenomes, in most cases readily explained as having emerged via duplication of a standard gene followed by sequence divergence of one of the copies (Table S3), but the functionality of most of them is unclear (see Note S5 for further details). An obvious exception concerns a tRNA gene first encountered in the mitogenome of *Monodopsis* sp. MarTras21, recognized as a divergent copy of the *trnR*(*ucu*) gene and, given its anticodon mutated to UGU, proposed to specify a tRNA decoding threonine codons (Ševčíková et al. 2016). Our expanded sampling revealed the presence of the putatively neofunctionalized *trnR*(*ucu*) copy in all other representatives of the tribe Monodopsideae. Furthermore, the newly evolved *trnR*(*ucu*) paralog exhibits absolute sequence identity between *Monodopsis* spp. and the WarPS-5 strain, while the gene in *Pseudotetraëdriella kamillae* differs only at four positions, despite a considerable depth of divergence of these taxa apparent from the comparison of other mitochondrial (and plastid) genes (Fig. S3). Hence, the gene seems to be evolving under a constraint on its sequence, consistent with the notion that it specifies a functional product, presumably a tRNA charged by threonine, which is also consistent with the fact that tRNA exhibits the identity elements recognized by threonyl-tRNA synthetase (as defined by Giegé and Eriani 2023). This is remarkable, as a gene for a tRNA cognate for threonine codons is absent from all other stramenopile mitogenomes investigated so far and is thought to have been lost in the stramenopile stem lineage (Ševčíková et al. 2016; Sibbald et al. 2021), to re-evolve only much later in the ancestor of the tribe Monodopsideae.

The previous comparative study of eustig mitochondrial genomes (Ševčíková et al., 2016) concluded that two separate lineages, represented by the tribe Nannochloropsideae and *Trachydiscus minutus*, use only UAG and UAA as termination codons. In accord with this, all eustigs were found to possess the nucleus-encoded mitochondrial release factor recognizing the termination codon pair UAG and UAA, i.e. mtRF1 (more accurately called mtRF1a; Duarte et al. 2012), whereas the paralogous mtRF2 (more accurately, mtRF2a) recognizing the pair UAA and UGA was reported to be absent from Nannochloropsideae and *T. minutus* (Ševčíková et al., 2016). We revisited the occurrence of the UGA codon in eustig mitogenomes and the mitochondrial RFs by exploiting the expanded sample of eustig mitogenomes and nuclear genome data. Strikingly, we found out that *Nannochloropsis oculata* in fact does use UGA as a stop codon, albeit very rarely (Table S8). In addition, we were able to identify an mtRF2a gene in the nuclear genome of *N. oculata* as well as other representatives of the Nannochloropsideae and of *T. minutus*. In reality, we could confirm that all eustigs with the nuclear genome or transcriptome data of a sufficient quality possess the whole set of organellar release factors including mtRF2a (Fig. S12). In spite of this, some lineages including *Microchloropsis* spp. and *T. minutus* really do not use UGA in their mitochondrial genes, as reported before (Table S8).

The retention of mtRF2a yet dispensing with UGA as a stop codon in mitochondria is not unprecedented and has been reported from several unrelated eukaryotes (Duarte et al. 2012). While the reasons are not known, one possible explanation is that in parallel to its role in translation termination, mtRF2a has other important functions. Indeed, the bacterial RF2 is involved in post peptidyl transfer quality control (post PT QC) during translation (Petropoulos et al. 2014), and mtRF2a might function analogously. Acquisition of a novel function by a “redundant” mtRF2a is, however, also imaginable, at least in some cases. Our phylogenetic analysis of eustig organellar release factors revealed that mtRF2a sequences from the genus *Microchloropsis* are extremely divergent (Fig. S12) and exhibit unusual mutations in functionally critical regions (Fig. S13). Specifically, their codon-binding region as well as the region normally containing the catalytic motif GGQ essential for the peptidyl-tRNA hydrolase activity (Duarte et al. 2012) are highly modified by multiple substitutions of otherwise invariant positions and even deletions that are expected to drastically modify the geometry of the regions and abrogate the catalytic activity of the protein. Despite its unusual sequence features, the mtRF2a gene is still transcribed in both *Microchloropsis* spp. (Table S8), indicating its functionality, although the role it plays is certainly not peptide release in translation termination. Interestingly, for some reason sequences of the equivalent plastidial version of the release factor, i.e. pRF2, in *Microchloropsis* are also more divergent compared to pRF2 from other eustigs (Fig. S12). Nevertheless, the protein most likely functions normally, as UGA is clearly employed (however rarely) as a termination codon by *Microchloropsis* plastomes (Table S8).

## Conclusions

Our work has further improved the knowledge of the genomic diversity and phylogenetic relationships of eustigmatophytes, an algal lineage that used to be neglected but lately has attracted substantial attention by both biologists interested in fundamental questions of algal life as well as the community of biotechnology-oriented researchers. Our results demonstrate that both plastid and mitochondrial genomes are highly informative on the cladogenesis of eustigmatophyte algae. With the current sequencing technologies, it is generally straightforward and affordable to generate organellar genome sequences from any newly isolated eustig, and we propose phylogenies based on both organellar genomes should become the main reference for delimiting eustig taxa. Besides implications for the eustig biology at the organismal level, our survey of the eustig organellar genomes underscores various facets of the organellar evolution and functioning *per se*. Perhaps most remarkable is the identification of several mitochondrial *orfs* that are patchily distributed across eustigmatophytes yet seem to be ancestral for the whole group or its major clades. What is the exact origin of these *orfs*? Why should a highly recurrent loss affect them? Why do these *orfs* occur in multiple different copies in the same genome? The most burning question then is what is the function of the proteins encoded by these *orfs*. While the answer is presently not in reach, our data point to the existence of an unknown “mitochondrial biology” in many eustigmatophytes, loosely paralleling the previous discovery of a similarly enigmatic biological process underpinned by the *ebo* operon in plastids of some eustigs. In summary, the organellar biology of eustigmatophytes defined a very attractive research frontier that calls for the development of reverse-genetic methods capable of targeting the organellar genomes.

## Appendix

Formal description of new taxa

Tribe Monodopsideae M.Eliáš, trib. nov.

Type genus: *Monodopsis* D.J.Hibberd, 1981

Diagnosis: The new tribe is created for a monophyletic group of eustigmatophyte algae containing the genus *Monodopsis* and its specific relatives, but excluding the genera *Nannochloropsis* D.J.Hibberd, 1981 and *Microchloropsis* M.W.Fawley, I.Jameson & K.P.Fawley, 2015. Members of the group share unique genomic features, for example a newly evolved tRNA gene in their mitochondrial genome decoding codons for threonine. Remark: Apart from *Monodopsis*, the tribe includes the genus *Pseudotetraëdriella* E.Hegewald, 2007 and presumably other genera, pending the clarification of the genus-level classification of organisms that are assigned to the tribe based on molecular phylogenetic evidence (see Fig. S1), such as the strain WarPS-5, the alga referred to as “*Monodus* sp. NIES-3918” (GenBank LC129528.1), or the uncultured organism documented by the environmental DNA sequence referred to as “eustigmatophyte clone PRS2_3E_43” (GenBank GQ330585.1).

Tribe Nannochloropsideae M.Eliáš, trib. nov.

Type genus: *Nannochloropsis* D.J.Hibberd, 1981

Diagnosis: The new tribe is created for a monophyletic group of eustigmatophyte algae containing the genus *Nannochloropsis* and its specific relatives, but excluding the genera *Monodopsis* D.J.Hibberd, 1981 and *Pseudotetraëdriella* E.Hegewald, 2007. Members of the group are distinguished from other eustigmatophytes including their sister group (the tribe Monodopsideae) by having transitioned from freshwater to the marine environment (except for possibly secondarily freshwater *Nannochloropsis limnetica*) and by exhibiting various simplifications at the molecular level, such as the loss of the mevalonate pathway for the synthesis of isoprenoid precursors (Yang et al. 2021) or of the proteins constituting the characteristic stramenopile tripartite flagellar hairs (Barcytė et al. 2022).

Remark: Besides *Nannochloropsis*, the tribe includes the genus *Microchloropsis* M.W.Fawley, I.Jameson & K.P.Fawley, 2015.

## Material and Methods

### Algal cultures, light microscopy, DNA isolation and sequencing

Cultivation of 16 newly sequenced strains was described previously or they were obtained from culture collections (details provided in Table S1). The nine unidentified strains previously reported by Fawley et al. (2021) were grown at 20⍰°C using liquid WH+ medium (Fawley et al. 2013) with a 14⍰h:10⍰h light:dark cycle under cool white fluorescent lights at 50⍰µmol m^−2^ s^−1^. The strain denoted Chlorobotryaceae sp. FP5 (now lost) was grown as described before (Wolf et al. 2018). DNA from these strains was isolated using the procedure of Fawley and Fawley (2004) with minor modifications (Ševčíková et al. 2019). The strains obtained from the Culture Collection of Algae at the University of Göttingen, Germany (SAG; https://sagdb.uni-goettingen.de/) or the NIES collection (https://mcc.nbrp.jp/) were cultivated in liquid Bold’s Basal Medium under continuous illumination (irradiance of approximately 40–50 µmol photons m^−2^ s^−1^) at a constant temperature of ~20⍰°C. Genomic DNA was extracted using a modified version of the Dellaporta et al. (1983) protocol, incorporating additional RNase and proteinase K treatment steps. Differential interference contrast (DIC) microscopy used a Nikon E600 microscope (Nikon, Melville, New York, USA) equipped with 60× (NA 1.4) and 100× (NA 1.4) objectives. Digital images were captured with an Olympus SC180 camera system (Olympus, Waltham, Massachusetts, USA) with CellSens imaging software. Images were finished with paint.NET 5.1.9. Library preparation and sequencing were performed by Macrogen Europe using the TruSeq Nano DNA Kit (insert size: 350 bp) and the Illumina NovaSeq 6000 platform. The 18S rRNA gene of the strain F5a 4/24-2w was amplified by PCR using the primers 18L-X and NS1-X with PCR conditions described in Fawley and Fawley (2004) and sequenced as described in Alexandre et al. (2025). The sequence was deposited at GenBank with the accession number PX677469.

### Assembly of plastid and mitochondrial genome sequences

The newly obtained Illumina reads were trimmed by Trimmomatic (Bolger et al. 2014) and assembled using SPAdes (Bankevich et al. 2012; for program versions used for particular strains see Table S1). Generation of whole genome assemblies from the additional 16 eustig strains, whose plastome and/or mitogenome sequences have not been reported yet and that were also included in this study, was described elsewhere (Ševčíková et al. 2019; Yang et al. 2021; Barcytė et al. 2022). Note that we do not separately report on the mitogenome sequence of *Lietzensia polymorpha* SAG 2217, as it proved to be identical to the mitogenome sequence of the strain *L. polymorpha* SAG 2220, in analogy to the previously noticed identical plastome sequences of these two strains (Barcytė et al. 2022). Conserved proteins encoded by previously published eustig organellar genome sequences were used as queries in TBLASTN searches (Altschul et al. 1997) of the new or previously described whole genome assemblies to identify scaffolds corresponding to the respective organellar genomes. For plastomes, three separate scaffolds were typically found, with their coverage and termini overlaps consistent with the conventional tetrapartite architecture of a circular-mapping genome with short and long single-copy regions separated by inverted repeats. Full plastome sequences thus could be readily reconstructed. In a few other cases manual curation of the assembled scaffolds was required, in all cases resulting in an assembly with the same conventional sequence organization. Mitogenome sequences were also readily assembled, in most cases being represented in the whole genome by a single scaffold with a short region of sequence identity at both termini indicating the circular-mapping nature of the genome sequence. Manual curation of the few cases that have the mitogenome split in the initial assembly into two or more separate scaffolds always resulted in an analogous circular-mapping sequence with no evidence for longer internal (direct or inverted) repeats. The assembled sequences were further validated by inspecting alignments of sequencing reads mapped to the consensus assembled sequences using Tablet (version 1.21.02.08; Milne et al. 2013).

### Organellar genome annotation and gene identification

FACIL (Dutilh et al. 2011) was used to assess possible codon reassignments in organelles of the eustigs investigated. In a few cases FACIL suggested a possible alteration of the genetic code, but manual investigations of all these candidates did not support the putative reassignments. Hence, the genetic code 1 (the standard code; following the NCBI nomenclature) and 11 (“Bacterial, Archaeal and Plant plastid code”, differing from the standard code only by a wider set of initiation codons) were employed for annotation of the mitochondrial and plastid genomes, respectively. Initial organellar genome annotation was obtained by using MFannot (Lang et al. 2023). The MFannot outputs were manually curated to correctly determine gene and coding sequence boundaries, verify and edit gene names, and to fill in missing open reading frames shorter than 100 nucleotides that proved to be conserved in at least one other eustig organellar genome. Initiation codons other than AUG were considered as the beginning of the coding sequence in both plastomes and mitogenomes if strongly supported by high conservation of the encoded N-terminal region of the given protein apparent in multiple sequence alignments of eustig homologs. BLASTP searches against NCBI protein sequence databases were used as the default method of homology detection for protein-coding genes. For genes specific for eustigs or their particular subgroups and lacking non-eustigmatophyte homologs discernible with BLAST, more sensitive methods were employed. These included HHpred searches against collections of profile HMMs as provided by the HHpred server (https://toolkit.tuebingen.mpg.de/tools/hhpred; Zimmermann et al. 2018). Custom multiple sequence alignments created with MAFFT version 7 (Katoh et al. 2019) or individual sequences were used as the HHpred queries. In some instances, protein structure models were built using AlphaFold2 accessed via ColabFold v1.5.5 (Mirdita et al. 2022), or AlphaFold 3 (Abramson et al. 2024) accessed via the dedicated AlphaFold server (https://alphafoldserver.com). The highest-scoring model was then compared to the suite of experimentally determined and predicted 3D structures as offered by the Foldseek Search Server (van Kempen et al. 2024).

The *rrn5* gene was identified and annotated with Infernal version 1.1.2 (Nawrocki and Eddy 2013) utilizing the conservative covariance model defined by Valach et al. (2014). The initial annotation of tRNA genes provided by MFannot was refined by assigning the specific identity to the three genes in each organellar genome annotated as *trnM*(*cat*), which correspond to the elongator tRNA^Met^, the initiator tRNA^fMet^, and the lysidinylated tRNA^Ile^ that decodes the AUA codon. These three tRNA species were distinguished by sequence comparison to tRNA genes in previously annotated eustig organellar genomes, in the case of mitochondrial sequences further verified by phylogenetic analysis (see below). Dedicated analyses were performed for a group of non-standard tRNA genes in mitogenomes of the genus *Vischeria*, including manual identification of respective homologs in those genomes where the region was not annotated as tRNA gene by MFannot. For each organellar genome a curated Feature Table (a five-column tab-delimited table) was generated and converted to a GenBank file using tbl2asn version 25.8 (NCBI Resource Coordinators 2018). GenBank files were fed into OGDRAW (Greiner et al. 2019) to generate gene maps of the organellar genomes. All newly assembled organellar genome sequences were deposited at GenBank (National Center for Biotechnology Information, NCBI); accession numbers of all organellar genome sequences analyzed in this study are listed in Tables S2 and S3.

### Single-gene phylogenetic analyses

The datasets for phylogenetic analyses of the 18S rRNA and *rbcL* genes were created by expanding the previously analyzed datasets (Barcytė et al. 2022) with additional eustig sequences published in the meantime, extracted from the newly generated genome assemblies, or newly determined by PCR and Sanger sequencing (strain F5a 4/24-2w; see above). Nucleotide sequences were aligned utilizing MAFFT v7.525 and the resulting alignments were trimmed with trimAl v1.5.rev0 (-gt 0.9 option; Capellla-Gutiérrez et al. 2009). Maximum likelihood phylogenies were inferred using IQ-TREE v2.0.3 (Minh et al. 2020) and the substitution models TN+F+I+G4 and GTR+F+I+G4 (selected by ModelFinder; Kalyaanamoorthy et al. 2017) for the 18S rRNA and the *rbcL* gene dataset, respectively. Node support was assessed with 10,000 ultrafast bootstrap replicates (option -B 10000; Hoang et al. 2018). The phylogenetic analysis of eustig mitochondrial tRNA genes was done using the ETE3 3.1.3 pipeline (Huerta-Cepas et al. 2016) as implemented on the GenomeNet (https://www.genome.jp/tools/ete/). The sequences were aligned using MAFFT v6.861b with the default options and a tree was inferred using IQ-TREE 1.5.5 (Nguyen et al. 2015) with TESTNEWONLY for model finder (TVM+R7 selected by the program) and 1000 replicates for SH-aLRT.

Nuclear genome-encoded proteins of special interest, namely homologs of the EF-Ts protein and of organelle-targeted release factors were gathered from eustig nuclear genome and transcriptome data or protein sequence annotations derived from them (accession numbers in Tables S5 and S8). Candidate sequences were identified with BLAST and when needed, coding sequences were reconstructed by manually predicting exon-intron structures of the respective gene. Conceptual translations of coding sequences were combined with existing protein sequence predictions and the two sequence sets (EF-Ts and release factors) were supplemented by homologs from reference non-eustigmatophyte organisms (accession numbers in Table S8). The sequences were aligned with MAFFT v7.310 (--auto --reorder --anysymbol options) and the alignments were trimmed with trimAl v1.4.rev15 (-gt 0.2 option). Phylogenetic trees were inferred using IQ-TREE v2.2.2.7 with the substitution models Q.pfam+R5 (EF-Ts) or Q.insect+F+R6 (release factors) selected by ModelFinder. Branch support was assessed by calculating 1,000 ultrafast bootstrap replicates. For presentation purposes the alignment of mtRF2a sequences was visualized in AliView v1.28 (Larsson 2014) and adjusted in Inkscape v1.2.2 (www.inkscape.org).

Phylogenetic analyses of the mitogenome-encoded OrfT, OrfV, and OrfW proteins were performed using IQ-TREE based on multiple sequence alignments obtained with MAFFT and trimmed with trimAl (the gappyout mode). The substitution model was selected automatically for each dataset and the tree topology was assessed by non-parametric bootstraps (100 replicates). Pseudogenized versions of *orfT, orfV*, and *orfW* existing only as short fragments were excluded from the analysis. Amino acid sequences corresponding to (longer) pseudogenes were conceptually translated with in-frame termination codons ignored. In the case of pseudogenes affected by frame-shift mutations, the coding sequence was (tentatively) reconstituted to yield a single continuous open reading frame. All phylogenetic trees presented in this paper were visualized in iTOL v7 (Letunic and Bork 2024) and graphically processed in Inkscape.

### Organellar phylogenomic analyses

For the phylogenetic analysis utilizing plastome sequences, the set of 69 conserved plastome encoded proteins used in a previous study (Barcytė et al. 2022) was updated and expanded by the corresponding sequences from the newly sequenced or assembled eustig plastomes plus sequences from the incomplete plastid MAG TARA_CHLORO_00525 reported by Jamy et al. (2025). The full list of plastome sequences included in the phylogenetic analysis with all relevant details is provided in Table S10. For the analyses based on mitogenome sequences, a set of 26 mitogenome-encoded proteins (their list is apparent from Table S3) was selected based on preliminary analyses and the respective orthologs were gathered from the previously reported and newly generated eustig mitogenome sequences supplemented by orthologs from other ochrophyte and non-ochrophyte representatives (the full list of mitogenomes included in the analysis is provided in Table S11). For *N. oculata* we used the mitogenome sequence KJ410688.1, as we found that the sequence in the RefSeq record NC_022257.1 is most likely chimeric (see Note S6 for details). The selection of the outgroup taxa was guided by sampling across as many major (class-level) lineages as possible but avoiding sequences that were found to form very long branches. Mitogenome sequences with no available annotation were annotated using MFannot and the newly predicted protein sequences were integrated into our dataset. The final manually curated sets of orthologous protein sequences were aligned with MAFFT v7.526 (-auto option). The alignments were trimmed with BMGE v1.12 with the default option (Criscuolo and Gribaldo 2010) and concatenated (separately for plastome- and mitogenome-encoded proteins) with Goalign v0.4.0 (Lemoine and Gascuel 2021) to obtain the final supermatrices. Maximum likelihood phylogenetic analyses were carried out with IQ-TREE multicore version 3.0.1 utilizing the substitution model LG+C60+F+G. Branch support was evaluated with 10,000 ultrafast bootstrap replicates.

### Additional analyses of protein and tRNA sequences

Protein sequences encoded by the mitochondrial *orfs* were gathered from all eustig mitogenomes analyzed in this study and clustered with CLANS 29.05.2012 (Frickey and Lupas 2004), using a matrix of pairwise sequence similarities generated by the CLANS web-utility in the MPI Bioinformatics Toolkit (https://toolkit.tuebingen.mpg.de/tools/clans) with the default setting (BLOSUM62, e-value of 1e-3). The occurrence of transmembrane domains in these proteins was assessed using TMHMM 2.0

(https://services.healthtech.dtu.dk/services/TMHMM-2.0/; Krogh et al. 2001) and DeepTMHMM - 1.0 (https://services.healthtech.dtu.dk/services/DeepTMHMM-1.0/; Hallgren et al. 2022). The presence of specific N-terminal organelle-targeting presequences in the nucleus-encoded EF-Ts homologs was assessed by using TargetP 2.0 (https://services.healthtech.dtu.dk/services/TargetP-2.0/; Almagro Armenteros et al. 2019). Secondary structures of tRNA molecules were predicted using Rfam (https://rfam.org/; Ontiveros-Palacios et al. 2025) or (in the cases of highly divergent sequences when Rfam did not recognize them as tRNA genes) with RNAfold WebServer (http://rna.tbi.univie.ac.at/cgi-bin/RNAWebSuite/RNAfold.cgi; Gruber et al. 2008). The predicted models were visualized and adjusted with VARNA version 3-93 (Darty et al. 2009).

## Supplementary Material

Supplementary material is available at Genome Biology and Evolution online.

## Acknowledgements

We thank John Cawley for initial analyses of some of the mitochondrial genome sequences reported in this study.

## Author contributions

M.R. performed most of the bioinformatic and phylogenetic analyses, assisted by T.Y. and T.Š, prepared most figures and tables, and drafted parts of the manuscript. E.K. performed the analyses of codon usage and release factors and prepared some of the tables and figures. K.P.F and M.W.F prepared Fig. 1, and along with K.J. and B.M.W grew algal cultures and isolated DNA for sequencing. D.B. and F.-W.L generated some of the sequence data and assemblies. M.E. conceived the research, secured funding, supervised M.R. and E.K., contributed to bioinformatic analyses, and drafted most of the manuscript.

## Data Availability

The newly sequenced or assembled plastid and mitochondrial genome sequences were submitted to GenBank with accession numbers PX978874-PX978889, PZ011442-PZ011445, PZ106320-PZ106330, PZ231586-PZ231598, PZ250447-PZ250453. Raw sequencing reads were deposited as NCBI BioProject PRJNA1425954. Draft genome sequence assemblies, sequence alignments, and original tree files are available from a Figshare repository (doi:10.6084/m9.figshare.31535176).

