## Supplementary notes and figures for "Insights into the phylogeny and enigmatic mitochondrial biology of eustigmatophyte algae from over 50 newly sequenced organellar genomes"

**Supplementary notes S1 to S6**

**Supplementary figures S1 to S14**

**Note S1.** Details on the eustig plastome-specific genes

To illuminate the possible role of *ycf95* we built a multiple sequence alignment of all available Ycf95 protein sequences and queried a database of profile HMMs with the highly sensitive homology detection tool HHpred, with no credible hits identified. However, as Ycf95 sequences from Eustigmatales are markedly divergent from those from *Goniochloridales*, we repeated the search with alignments restricted to sequences from one or the other eustig subgroup. No significant hits were retrieved with either of the alignments, but we noticed that with the *Goniochloridales* alignment as a query, the first hit was potentially homologous (though with an unconvincing probability value of 78.33) to the conserved domain of unknown function DUF1257. Strikingly, this domain corresponds to a functionally uncharacterized protein encoded by the plastid gene *ycf35*, which is common in plastomes of various ochrophytes, but is apparently missing in eustigs (Table S2). We also noticed that Ycf95 proteins exhibit absolute conservation of the first two amino acid residues directly downstream from the N-terminal methionine. This characteristic "MSH" motif is found at the N-terminus of essentially all Ycf35 proteins we examined in our study. Notably, this N-terminal region was not included in the alignment between HMMs of Ycf95 and Ycf35 generated by HHpred and did not contribute to evaluation of possible homology of these proteins by the program, providing an independent line of evidence linking the two proteins. Furthermore, although the absence of *ycf35* in chrysophytes seemed to support the interpretation that the gene was lost before the chrysophyte-eustig split (Ševčíková et al. 2019), we now found *ycf35* in the TARA_CHLORO_00525 plastome. We tried to further test the putative relationship of *ycf95* to *ycf35* by predicting the tertiary structure of *ycf95* gene products using AlphaFold 3 and using the predicted structures as queries for Foldseek searches against databases of other (experimentally determined or predicted) structures, but the quality scores of the predictions were generally very low and no convincing hits were retrieved. In contrast, the AlphaFold 3 model of the protein encoded by *ycf35* in the TARA_CHLORO_00525 plastome exhibited a much higher confidence and did retrieve predicted models of Ycf35 proteins from plastids and Cyanobacteria in a Foldseek search (Fig. S14). Hence, the evolutionary link between *ycf95* and *ycf35* remains to be corroborated, but the most parsimonious interpretation of our observations is that *ycf95* is an extremely divergent evolutionary descendant of the ancestral plastidial *ycf35* gene.

Two genes, denoted *orf1_eust* and *orf1_gon*, were previously found to be conserved in a subset of Eustigmatales and in *Goniochloridales*, respectively, yet lacked any discernible homology in other organisms (Ševčíková et al. 2019). Our expanded sampling showed that *orf1_eust* is present in all members of the families Chlorobotryaceae and Monodopsidaceae analyzed, but it is missing from the two more basally branching lineages of the Eustigmatales, i.e. Neomonodontaceae and *Paraeustigmatos columelliferus* (Table S2). These results solidify the notion that *orf1_eust* emerged within Eustigmatales after the divergence of the two basal-most lineages, and once established it remained an essential part of the plastid gene set. Similarly, *orf1_gon* proved to be ubiquitous in *Goniochloridales* (Table S2; Dataset S1). Analogously to *ycf95*, we tested the possibility that the expanded sampling and the progress in protein structure modelling will help illuminate the origin and function of *orf1_eust* and *orf1_gon*. HHpred analysis did not reveal significant similarity to any known protein for any of them. Modelling the structures of *orf1_eust* and *orf1_gon* protein products with AlphaFold2 or the newer AlphaFold 3 resulted in low-confidence predictions, and subsequent Foldseek search yielded no significant hits for the predicted 3D structures of any of the proteins. Hence, the origin and function of *orf1_eust* and *orf1_gon* remain elusive.

Plastid genomes of three strains, WTwin 8/9 T-6m6.8, Chic 10/23 P-6w, and *Pseudellipsoidion edaphicum* CAUP Q 401, were each previously shown to harbor one to multiple ORFs longer than the arbitrary threshold, but had no discernible homology to genes in other eustigs or other organisms, and hence annotated as *orfs* (Ševčíková et al. 2019; Barcytė et al. 2022). These *orfs* remain true orphans with elusive origin and uncertain functionality despite the acquisition of plastid genome data from close relatives of each strain, i.e. a second representative of clade Ia (a relative of WTwin 8/9 T-6m6.8), several additional members of the clade IIc (related to Chic 10/23 P-6w), and a second representative of Neomonodontaceae (a relative to *P. edaphicum* CAUP Q 401). Interestingly, none of the 20 eustig plastomes newly characterized in this study contain any new lineage-specific *orf* (other than *orf1_eust* and *orf1_gon* mentioned above). This indicates that expansions of the plastid gene complement by gaining new genes are rare events in eustig evolution.

**Note S2.** Details on the history of *rps4N* and *rps4C* fusions in eustig mitogenomes

The presumed physical proximity of the proteins encoded by *rps4N* and *rps4C* is consistent with the existence of their fused version (annotated simply as *rps4*) that occurs in a subclade of the *Goniochloridales* clade IIc (*Vacuoliviride crystalliferum* and the strains BogD 8/9 T-2w and Chic 10/23 P-6w), *Pseudostaurastrum* spp., *Chlorobotrys* sp. FD2, and *N. aerophytica*. This patchy distribution of the fused version indicates multiple independent origins, but the precise number of these events is uncertain due to the specific situation in a subclade of Chlorobotryaceae comprised of the genus *Vischeria*, having *rps4N* and *rsp4C* as separate genes, together with *N. aerophytica* and *Chlorobotrys* sp. FD2. The latter two taxa form a paraphyletic grade at the base of *Vischeria*, which may imply that the fused *rps4* version in the *Chlorobotrys* sp. FD2 and *N. aerophytica* evolved independently in their lineages. However, a scenario assuming a single fusion event in the ancestor of the whole subclade with a secondary split of the gene in *Vischeria* is also conceivable and in fact supported by considering the following additional observations. First, the fused *rps4* gene in *N. aerophytica* is unique (even compared to the one from *Chlorobotrys* sp. FD2) by encoding a protein in which the halves corresponding to *rps4N* and *rsp4C* are separated by an expanded middle lysine-enriched region. Second, in all *Vischeria* species/strains the *orf* separating the *rps4N* and *rps4C* genes encodes a putative short protein whose major part very well aligns with the unique middle region of the expanded *N. aerophytica* S4 protein, indicating homology (Fig. S5C). Hence, the most likely interpretation is that the fused *rps4* gene was modified in the common ancestor of *N. aerophytica* and *Vischeria* by an internal expansion and subsequently split in the lineage leading to the genus *Vischeria* into three separate ORFs. Hence, we annotated the *Vischeria*-specific short *orf* between *rps4N* and *rps4C* accordingly as *rps4M* ("M" from "middle") and posit that it encodes a part of the *Vischeria* mitoribosome.

**Note S3.** Details on the putative phage-derived DNA insertion in the mitogenome of *Vacuoliviride crystalliferum*

A HHpred search with the conceptual translation of orf260 from the *V. crystalliferum* mitogenome retrieved a high-confidence hit (probability 100, E-value of 4.3e-43) to the Pfam family PF00940 representing DNA-dependent RNA polymerases typified by the proteins from T3/T7 (T-odd) phages. In addition to phage proteins, PF00940 includes the nucleus-encoded mitochondrial RNA polymerase (POLRMT) transcribing mitogenomes of most eukaryotes and polymerases encoded by diverse previously identified mitochondrial plasmids (Shutt and Gray 2006). The *orf260* sequence corresponds only to a fragment of the full RNA polymerase, indicating it is non-functional. It is also not similar to the POLRMT, so *orf260* is unlikely to represent a DNA transfer from the nuclear to the mitochondrial genome. Inspired by the previous discovery of a novel mitochondrial plasmid in the non-photosynthetic ochrophyte *Leucomyxa plasmidifera* (Barcytė et al. 2024), we searched with the *orf260*-encoded protein sequence against all available full eustig genome assemblies, including the one from *V. crystalliferum,* to check for the possible existence of a mitochondrial plasmid encoding a related RNA polymerase, but we did not find any candidate. Hence, *V. crystalliferum orf260* most likely represents a chance insertion of a fragment of a DNA phage. There is another singleton (*orf188*) downstream of *orf260* in the *V. crystalliferum* mitogenome, and it is notable that these two *orfs* are both located on the opposite strand than virtually all other genes (see Fig. 8A and the discussion below of the conserved organization of eustig mitogenome), which supports the notion of an exogenous, single-insertion origin of this whole region. Altogether, there is presently no direct evidence for the functionality of any of the singleton *orfs* and most, if not all, may be just accidental outcomes of neutral evolutionary processes affecting eustig mitogenomes.

**Note S4.** Details on the occurrence and features of conserved patchily distributed eustigmatophyte-specific mitochondrial *orfs*.

The *orfO* gene is restricted to the two closely related species of the genus *Monodopsis* and occurs in two paralogous copies positioned directly downstream of the two copies of *cox1* that uniquely occur in *Microchloropsis* spp., indicating duplication of the whole *cox1-orfO* block (Dataset S4). Both OrfO paralogs are short proteins (<100 amino acid residues) with two TM helices occupying the opposite ends of the protein, and highly conserved between the two species but diverged from each other. The *orfP* gene, also restricted to *Microchloropsis* spp., is positioned between the duplicated *cox1-orfO* blocks. The *orfP* version occurring in *M. salina* encodes a protein similar in size and architecture to OrfO, but potential homology between OrfP and OrfO could not be confirmed by sequence comparisons. The *orfP* locus in *M. gaditana* is very similar to that of *M. salina* but it is interrupted by an in-frame termination codon. As a result, only the part starting with an internal AUG codon downstream of the stop codon was annotated as the locus NagaMp0004 by Starkenburg et al. (2014). This indicates recent pseudogenization of *orfP* in *M. gaditana*, or that it is only an accidental non-translated ORF even in *M. salina*. The *orfQ* gene occurs only in two members of the *Goniochloridales* clade IIc and again encodes a short protein with two TM helices at the ends, but in this case the divergence between the two orthologs is more pronounced and their positions in the two mitogenomes differ (Dataset S4).

The *orfR* gene is also taxonomically restricted, found only in a few members of the *Goniochloridales* clade IIa, but the OrfR proteins are >200 amino acid residues long and thus much longer than proteins of the OrfO to OrfQ set, being instead comparable to proteins encoded by the more broadly occurring *orfS* to *orfW* genes. Furthermore, all these proteins exhibit a similar general architecture, with all their multiple (three to five) TM helices located in the N-terminal region followed by a prolonged C-terminal region presumably folding into a soluble domain. The few exceptions that depart from this architecture, i.e. proteins with the N-terminal transmembrane or the C-terminal soluble domain truncated or missing, potentially all represent degraded versions of the respective *orfs* (Table S6). This may also be the case of the *orfS* locus in *Lietzensia polymorpha*, i.e. the only representative of *orfS* in Eustigmatales encountered in our mitogenome dataset, which differs from its *Goniochloridales* homologs by lacking the C-terminal soluble domain. However, the presence of a relatively long ORF directly downstream from the *L. polymorpha orfS* but in a different reading frame suggests that the original gene was longer but has been disrupted by a frameshift mutation, although the potential C-terminal extension is admittedly not particularly similar to the C-terminal domain of the proteins encoded by the two *Goniochloridales orfS* genes.

**Note S5.** Details on unusual tRNA loci in eustigmatophyte mitogenomes

The gene corresponding to tRNA-X in *Vischeria* sp. CAUP Q 202 was previously denoted "*trnX(uuaa)*", with "uuaa" referring to the putative expanded anticodon region (Ševčíková et al. 2016). The five additional *Vischeria* mitogenomes included in the main analyses presented in this study, as well as the additional mitogenome of *Vischeria* cf. *polyphem* strain CAUP H4302 (GenBank accession number MK170182.1), all contain an orthologous locus found at the same conserved position between the genes *rpl16* and *nad4L* (Dataset S4) and predicted to specify tRNA-like molecules that vary highly in the anticodon arm regions while showing little sequence variability in the other parts. Specifically, the expanded anticodon loop in *Vischeria* sp. CAUP Q 202 is an exception rather than a rule, whereas some of the newly analyzed *Vischeria* species/strains have the loop contracted, with the extreme encountered in *Vischeria* sp. ex *P. kamillae* having the anticodon stem truncated and the anticodon loop missing completely (Fig. 6D and Fig. S8). Another unusual mitochondrial tRNA, denoted tRNA-Leu2 for the presence of a predicted anticodon UAA matching leucine codons, was previously identified in *Vischeria* sp. CAUP Q 202 and shown to be related to tRNA-X, together forming a divergent offshoot of tRNA-Lys specified by the *trnK(uuu)* gene (Ševčíková et al. 2016). As with tRNA-X, the tRNA-Leu2 gene has orthologs in the other *Vischeria* species/strains, with the exception of *Vischeria stellata*. The respective locus is again positionally conserved, lying between the genes *rns* and *atp1*; only in *Vischeria* sp. ex *P. kamillae* the gene order is changed due to a three-gene block moved from elsewhere and inserted between the tRNA gene and *atp1* (Dataset S4). Analogously to tRNA-X, the tRNA-like molecules specified by the tRNA-Leu2-related loci vary primarily in the anticodon arm, with *Vischeria* sp. C074, *Vischeria* sp. ACOI 3415, and *Vischeria* cf. *polyphem* strain CAUP H4302 exhibiting only a truncated anticodon stem without an anticodon loop (Fig. 6E and Fig. S10). As the extended analysis raises serious doubts on the involvement of the respective RNA molecules in the process of translation (see main text), the title "tRNA-Leu2" is potentially misleading and we have renamed this tRNA as tRNA-X2, whereas the original closely related tRNA-X becomes "tRNA-X1".

Remarkably, when analyzing these unusual tRNA genes we found that all seven *Vischeria* species/strains exhibit yet another homolog, whose presence in the *Vischeria* sp. CAUP Q 202 mitogenome was previously missed, as the locus in this particular *Vischeria* member is not recognized by the tRNA-predicting algorithm built in MFannot. This third non-standard tRNA-like locus, naturally denoted tRNA-X3, occupies the position between the genes *atp9* and *rnl* (Dataset S4), is specifically related to the *Vischeria*-specific tRNA-X1 and tRNA-X2 (Fig. S11), and also specifies tRNA-like molecules that primarily vary in the anticodon arm region, with *Vischeria* sp. ex *P. kamillae* completely lacking it (Fig. 6F and Fig. S10). Hence, we assume that like specific tRNA-X1 and tRNA-X2, tRNA-X3 does not decode codons in the standard process of translation, although the persistence of these tRNA-like genes across the radiation of the genus *Vischeria* indicates that they are kept by selection. The exceptional absence of a tRNA-X2 gene in the *V. stellata* mitogenome is genuine and even traces of it were not found in the mitogenome (the region between the *rns* and *atp1* genes is very short). The mitogenome-based phylogeny favors an internal position of *V. stellata* in the *Vischeria* clade (Fig. S3), which would imply tRNA-X2 loss in this species. However, the plastome-based phylogeny places *V. stellata* sister to other *Vischeria* members (Fig. 2), which would be compatible with a scenario assuming the origin of tRNA-X2 (as a duplication of the pre-existing tRNA-X1 locus) only after the *V. stellata* lineage had diverged.

*Neustupella aerophytica*, a eustig most closely related to *Vischeria*, exhibits a non-standard tRNA locus clearly originated via duplication of the standard *trnK(uuu)* gene like the *Vischeria* tRNA-X1/2/3 clade (Fig. 6G). According to its anticodon sequence (AAA) the given *N. aerophytica* tRNA molecule would decode the phenylalanine codon UUU, but the *N. aerophytica* mitogenome at the same time contains the standard (and very different) *trnF(gaa)* gene specifying a tRNA that is the obvious default decoder of phenylalanine codons. Whether the product of the *trnK(uuu)*-derived "*trnF(aaa)*" locus in *N. aerophytica* is charged with phenylalanine and functions in translation in parallel to the standard $tRNA^{Phe}_{GAA}$ remains to be determined. Alternatively, it is conceivable (although not directly supported by the phylogenetic analysis of tRNA genes) that the common ancestor of the *Vischeria*/*Neustupella* clade duplicated the *trnK(uuu)* locus and exapted one of the copies for a new function that does not rely on the anticodon (or the anticodon arm as a whole). What this function might be, and why the neofunctionalized tRNA gene was further duplicated or triplicated in the *Vischeria* lineage is presently unclear.

Analogous to "*trnF(aaa)*" locus in *N. aerophytica*, there are additional "extranumerary" tRNA genes in some eustig mitogenomes that have emerged as divergent paralogs of standard tRNA genes (Table S3). In some cases, the duplication was followed by mutation of the anticodon sequence, potentially resulting in "tRNA gene recruitment" (Wang et al. 2011) or "tRNA remolding" (Sahyoun et al. 2015), i.e. adopting a new amino acid specificity and the corresponding decoding capacity by a pre-existing tRNA due to mutations of its gene. However, it is possible that most of these cases represent non-functional copies (pseudogenes), as a gene for the standard tRNA decoding the same codon(s) has been retained by the respective mitogenomes. This is most likely also the case of a tRNA locus in *N. aerophytica* that was annotated by MFannot as a "suppressor" tRNA gene with its putative product carrying the anticodon UCA cognate to the termination codon UGA. However, there is no indication for UGA encoding an amino acid in any *N. aerophytica* mitochondrial gene and the locus most likely evolved by extreme sequence divergence of the original *trnL(taa)* gene (whereas its new functional copy is found elsewhere in the genome).

**Note S6.** Identification of the KC568460.1/NC_022257.1 sequence as a likely chimera.

During our analyses we noticed that two independently determined mitogenome sequences for *Nannochloropsis oculata* CCMP 525, i.e. GenBank records KC568460.1 and KJ410688.1 reported by Wei et al. (2013) and Starkenburg et al. (2014), respectively, are not identical. The genome sequences share blocks of sequence identity separated by regions where the sequences substantially differ, with KJ410688.1 including an extra region making it longer by >2,000 bp than KC568460.1. Investigating this unexpected difference further we found that the sequence KC568460.1 exhibits regions of identity or near identity with mitogenomes from other *Nannochloropsis* (and *Microchloropsis*) species sequenced in the same study (Wei et al. 2013), and these regions correspond to the regions where KC568460.1 and KJ410688.1 differ. Hence, it seems that KC568460.1, which was chosen as the reference sequence for *N. oculata* (RefSeq record

NC_022257.1) is artificial (chimeric), and the sequence KJ410688.1 reported by Starkenburg et al. (2014) is the more credible version; we thus used KJ410688.1 rather than KC568460.1/NC_022257.1 in our analyses.

**References to supplementary notes**


Alkatib S, et al. The contributions of wobbling and superwobbling to the reading of the genetic code. PLoS Genet. 2012:8:e1003076. https://10.1371/journal.pgen.1003076.

Sahyoun AH, et al. Towards a comprehensive picture of alloacceptor tRNA remolding in metazoan mitochondrial genomes. Nucleic Acids Res. 2015:43:8044–8056. https://doi.org/10.1093/nar/gkv746.

Ševčíková T, et al. A comparative analysis of mitochondrial genomes in eustigmatophyte algae. Genome Biol Evol. 2016:8:705–722. https://doi.org/10.1093/gbe/evw027.

Shutt TE, Gray MW. Bacteriophage origins of mitochondrial replication and transcription proteins. Trends Genet. 2006:22:90–95. https://doi.org/10.1016/j.tig.2005.11.007.

Starkenburg SR, et al. A pangenomic analysis of the *Nannochloropsis* organellar genomes reveals novel genetic variations in key metabolic genes. BMC Genomics. 2014:15:212. https://10.1186/1471-2164-15-212.

Wang X, Lavrov DV. Gene recruitment – A common mechanism in the evolution of transfer RNA gene families. Gene. 2011:475:22–29. https://doi.org/10.1016/j.gene.2010.12.009.

Wei L, et al. Nannochloropsis plastid and mitochondrial phylogenomes reveal organelle diversification mechanism and intragenus phylotyping strategy in microalgae. BMC Genomics. 2013:14:534. https://doi.org/10.1186/1471-2164-14-534.

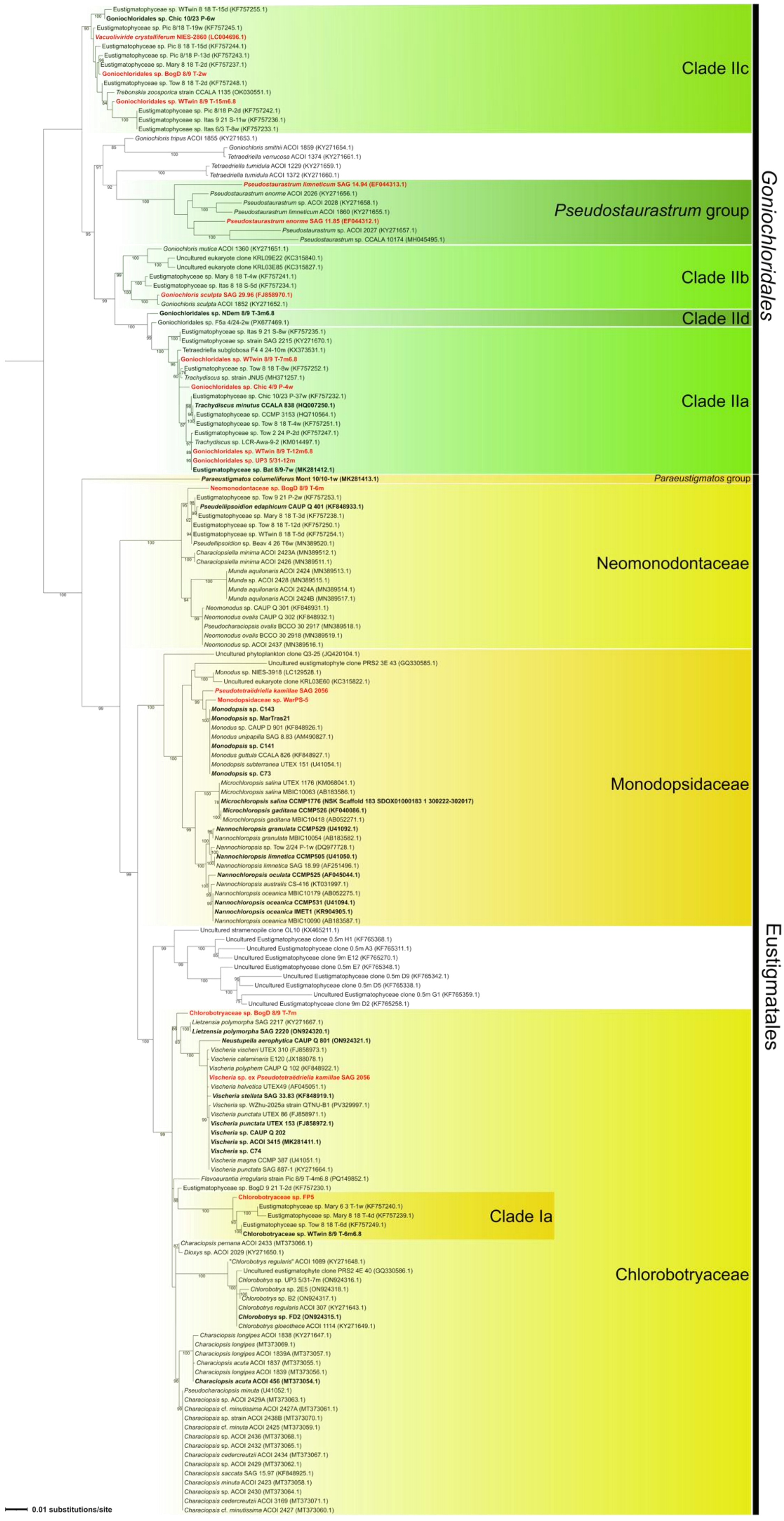


**Fig. S1.** ML phylogenetic tree of eustigmatophytes based on the 18S rRNA gene sequences (1,065 aligned nucleotide positions). The tree was inferred with the substitution model TN+F+I+G4. Numbers at branches correspond to UFB values (shown when ≥50). Strains analyzed and included in plastid and mitochondrial trees are highlighted in bold, newly sequenced strains in this study are highlighted in red.

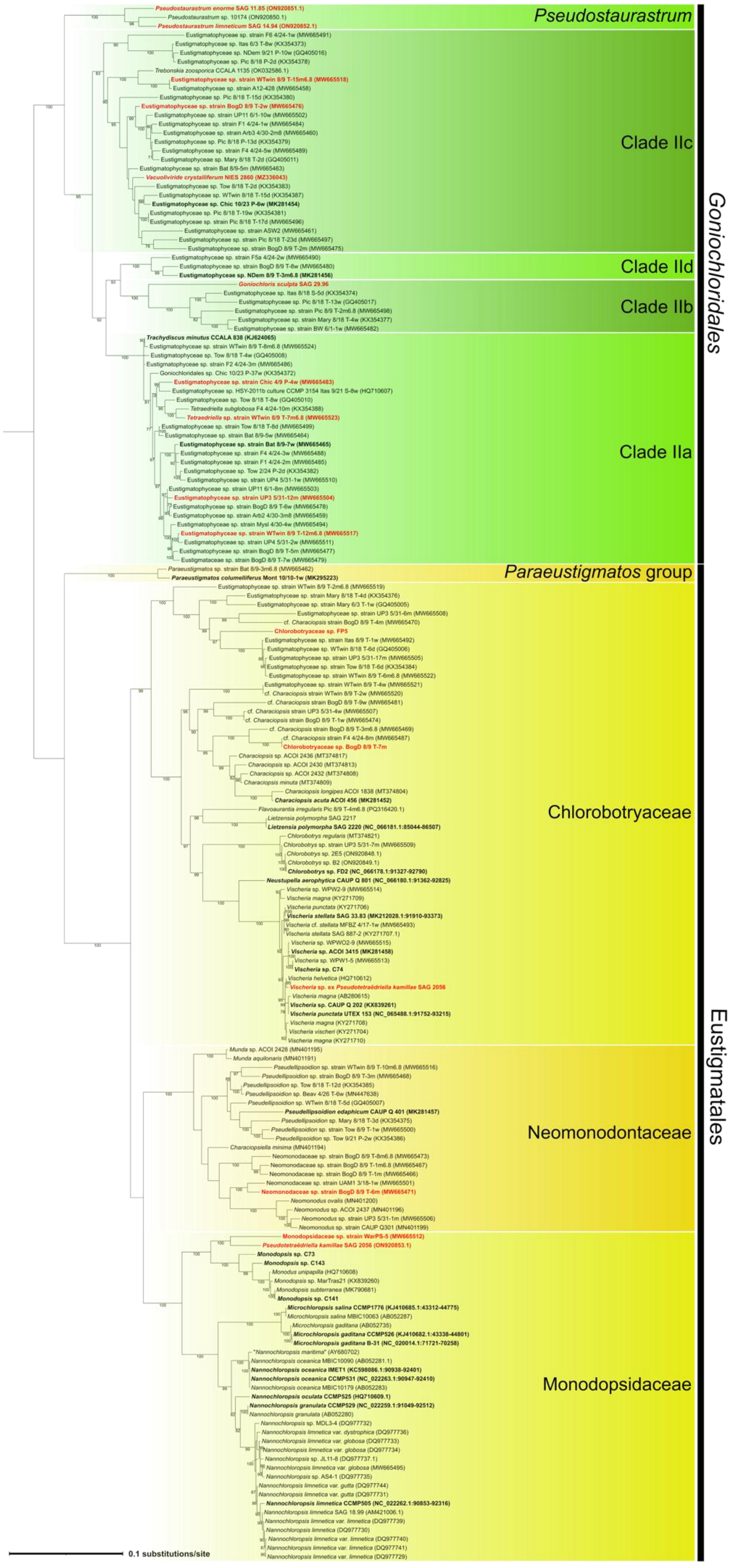


**Fig. S2.** ML phylogenetic tree of eustigmatophytes based on the *rbcL* gene sequences (1,516 aligned nucleotide positions). The tree was inferred with the substitution model GTR+F+I+G4. Numbers at branches correspond to UFB values (shown when ≥50). Strains analyzed and included in plastid and mitochondrial trees are highlighted in bold, newly sequenced strains in this study are highlighted in red.

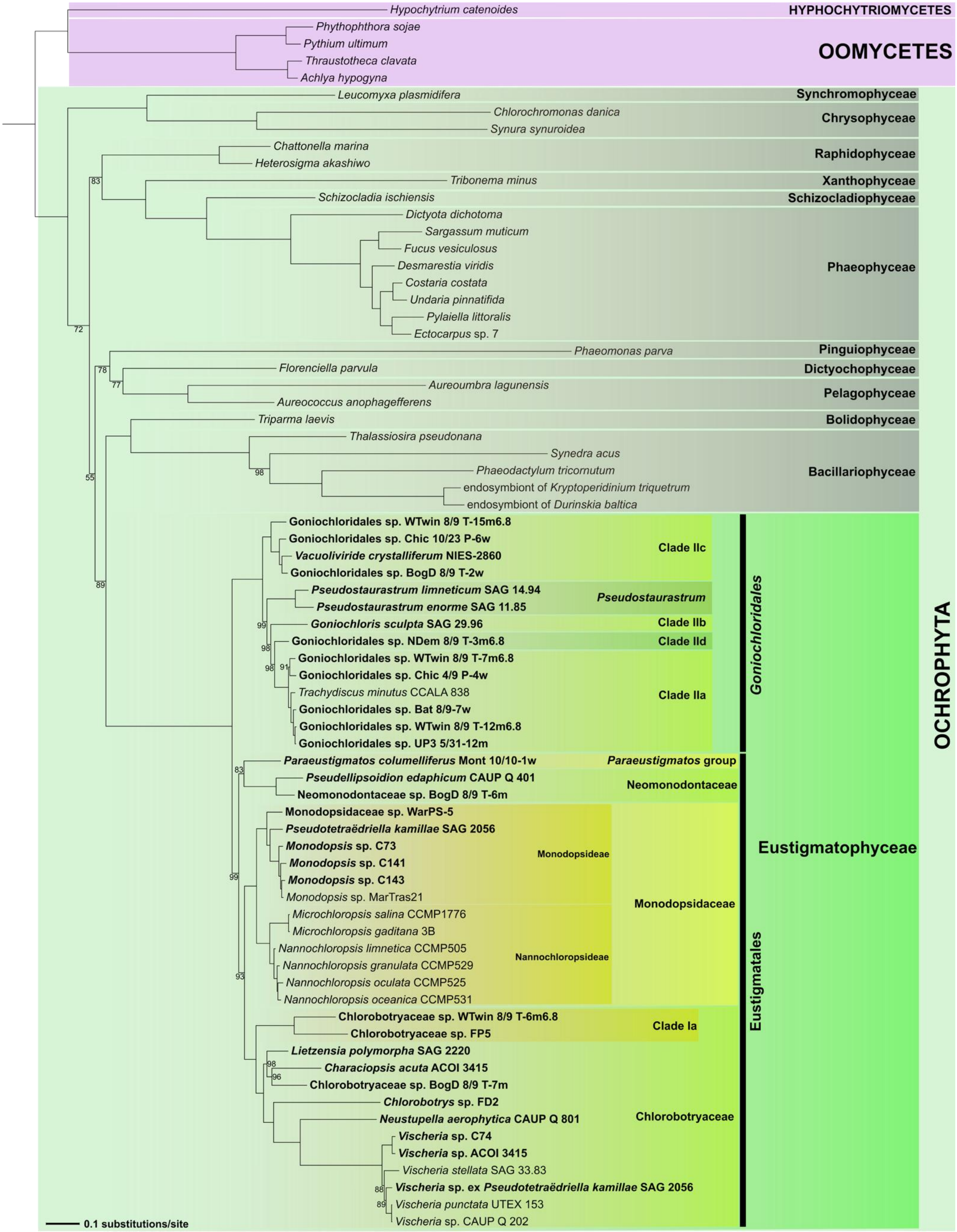


**Fig. S3.** Maximum likelihood phylogeny of ochrophytes derived from a concatenated dataset of 26 mitogenome-encoded proteins. The tree was inferred from a supermatrix of 5,463 amino acid positions with IQ-TREE v3.0.1 and the LG+C60+F+G model. Sequences from Pseudofungi (Oomycetes and Hyphochytriomycetes) were included as an outgroup. Branch support values (ultrafast bootstraps calculated from 10,000 replicates) are indicated only if lower than 100. Newly sequenced or assembled mitochondrial genomes are highlighted in bold. Accession numbers of all mitogenome sequences included in the analysis are provided in Table S3.

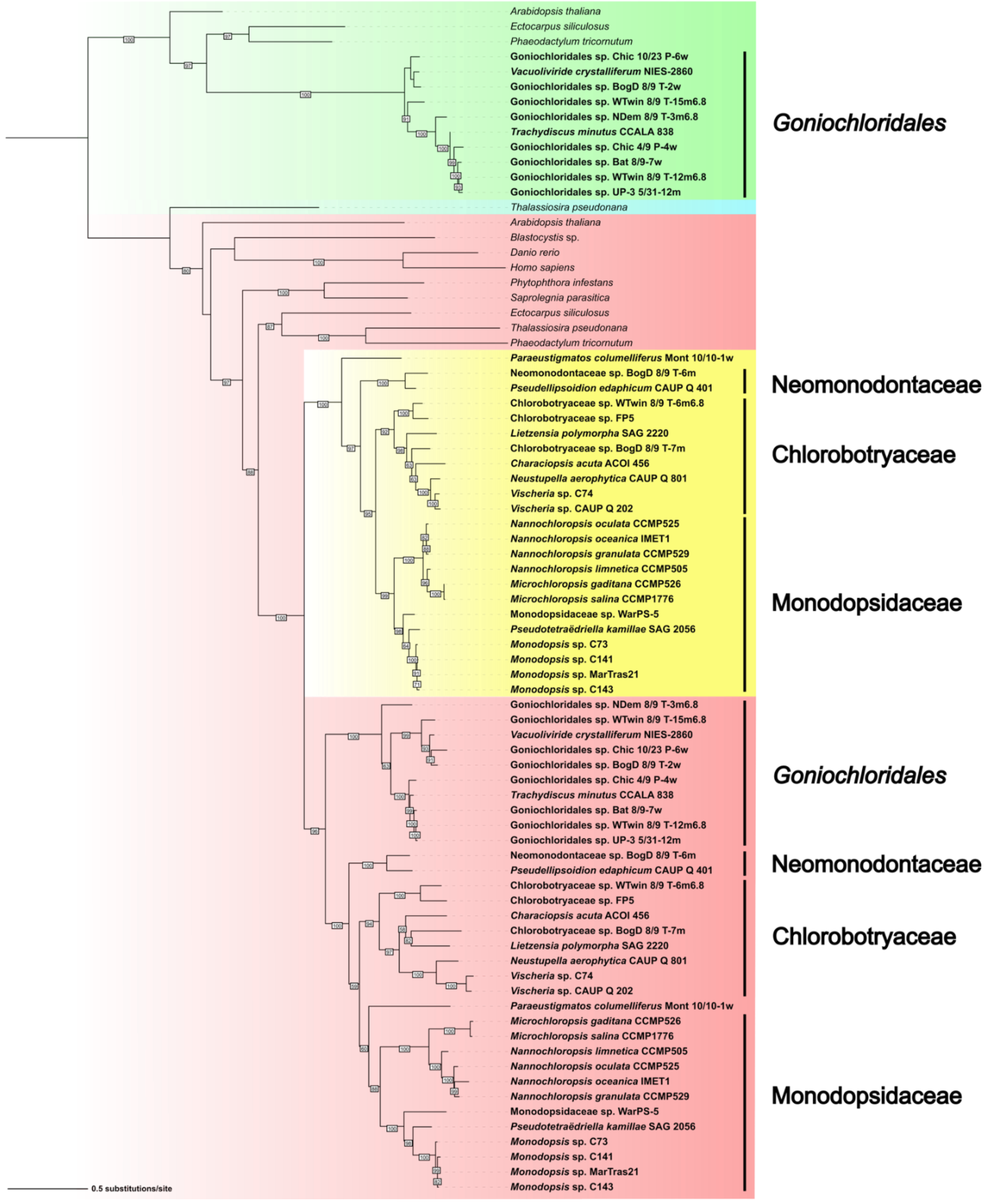


**Fig. S4.** Phylogeny of plastidial and mitochondrial elongation factor Ts (EF-Ts). Standard plastidial EF-Ts proteins (plastome-encoded in eustigs and other ochrophytes) are highlighted in green; note that in eustigs this version exists only in *Goniochloridales*. The nucleus-encoded mitochondrion-targeted version is highlighted in red, whereas the nested clade of proteins from Eustigmatales representing a newly evolved paralog with changed targeting presequences predicted to localize the protein into the plastid is highlighted in yellow. The sequence highlighted in turquoise is a nucleus-encoded plastid-targeted EF-Ts version found in a diatom lineage (in the tree exemplified by *Thalassiosira pseudonana*) that lack the plastome-encoded version (like Eustigmatales) and have replaced it with a gene most likely acquired from a bacterial source, as judged from a BLASTP search. At face value, the topology of the inferred tree suggests duplication of the gene encoding the mitochondrial EF-Ts in the eustigmatophyte stem lineage with the subsequent loss of one of the variants in the *Goniochloridales* stem lineage, but it seems more likely (also considering the limited length of the EF-Ts protein, and hence the amount of phylogenetic signal in its sequence), that the topology does not fully recapitulate the evolution of the gene family and the gene duplicated only in the Eustigmatales stem lineage.

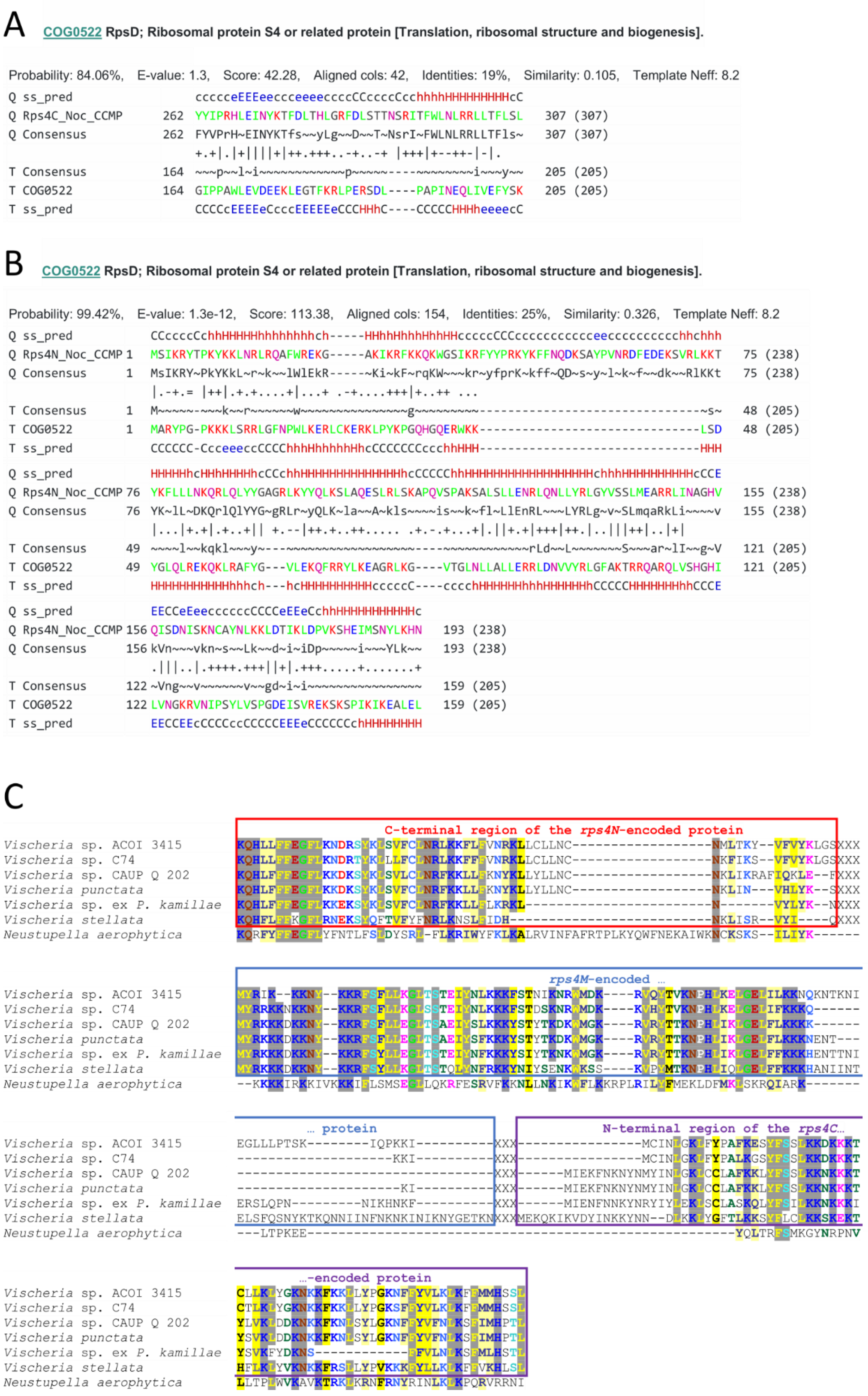


**Fig. S5.** Modifications of the mitochondrial *rps4* gene in eustigmatophytes. (A) and (B) Outputs of HHpred searches showing matches of profile HMMs constructed from multiple alignments of the eustigmatophyte proteins encoded by the *rps4C* (previously *orfX*) and *rps4N* (previously *rps4*) genes, respectively, to the profile HMM representing the COG family RpsD, i.e. the reference eubacterial *rsp4* equivalent. Note that the eustig Rps4C and Rps4N proteins match complementary regions of the RpsD protein. (C) Evidence for homology of the middle part of the fused *rps4* gene in *Neustupella aerophytica* and the short *Vischeria*-specific *orf,* annotated as *rps4M*, in between *rps4N* and *rps4C* genes. For the purpose of the presentation, concatenated protein sequences encoded by the *Vischeria* spp. *rps4N*, *rps4M*, and *rps4C* (separated by "XXX") were aligned to the protein sequence encoded by the (fused) *N. aerophytica rps4* gene (only the middle part of the full alignment is included in the figure).

# A

**8A22_Ba bS1m; mitoribosome, ribosome, mitochondria, polytomella, fragmentation, rrna, evolution, translation; HET: K, ATP, MG; 2.91A {Polytomella magna}**

Probability: 67.42%,   E-value: 120,   Score: 27.05,   Aligned cols: 118,   Identities: 13%,   Similarity: 0.045,   Template Neff: 8.8

```
Q ss_pred            CcccccCceEEEEEEeCCcCceEEEEecccceeeehHHHHHhhcHHhceeccCC---------CCCCCCceEEEEEEeeeC
Q orfY_Noce      21  PKPLKSQNIYEAFVRSRKKKEVDLEFREMLIYKTEQQFNCLKRLENRFDIYGK---------ALPNEGQVYTFRFKLFDN   91 (228)
Q Consensus      21  Pk~lk~~~iy~~~~~s~~Kkev~lE~~~~~~~~~~~~~~~lL~~lENRFwlY~~---------~~~~~~q~ysfcFks~D~   91 (228)
                     +..++.+.+..+.+.......+-+++ +....-.-..-++ ..-++... .....++...+.....|.
T Consensus      76  ~~~~~~G~~v~g~V~~v~~~gv~Vdi-g~~~~g~i~~~el-----~~~~~~~~~~~~~~~~~~~~~~vGd~i~v~V~~i~~   149 (242)
T 8A22_Ba        76  TMPILKGQQIQAKVIQKTDEVMLLDP-GFYNLSEVPVNYL-----TTAHIVRKVDDSPRENLYDVRPGDVVKVLVDDVYT   149 (242)
T ss_pred            CCCCCCCCEEEEEEEEEeCCEEEEEe-CCCcEEEEEHHHh-----chheeeccCCCCCccccccCCCCCEEEEEEEEeeC

Q ss_pred            CCCeEeeecc-----cccccchhHHHHHHHhcchhhhcCceeeehhheeeecCeEEEEecCCeEEE
Q orfY_Noce      92  TSKLVNFSDR-----KRRLQNISSYIYAYNSRRHILKNITKNIKSRLLCLTKGGFKISFFGQKVIL   152 (228)
Q Consensus      92  ~sk~~lFs~~-----kkkl~~~~S~~~~~~n~r~~~l~~~~~liKsRlL~LtKgGf~~s~~G~~a~l   152 (228)
                     ..+.+.++.+ ......|..+...+...+ .+.+++...+++|+...+-|..|++
T Consensus      150 ~~~~i~lS~~~~~~~~~~~~~~~~~~~~~~~~~g~--------~v~g~V~~~~~~G~~V~i~gi~g~l   207 (242)
T 8A22_Ba       150 PYGDMQLDVPQQDPRLILNQVWDELHLKMKKKE--------LVRGRILNECKSGYAVGVAGFVALL   207 (242)
T ss_pred            CCCcEEecccCChHHHHHHHHHHHHHhhhcCC--------eEEEEEeecCCEEEEEECCEEEEE
```

# B

**4Q7J_H 30S ribosomal protein S1; RNA polymerase, RNA binding motif, RNA dependent RNA polymerization, TRANSLATION-TRANSFERASE complex; HET: SO4; 2.9A {Escherichia coli}**

Probability: 51.23%,   E-value: 220,   Score: 24.61,   Aligned cols: 120,   Identities: 14%,   Similarity: 0.118,   Template Neff: 11.1

```
Q ss_pred            hhcCCcccccCceEEEEEEeCCcCceEEEEecccceeeehHHHHHhhcHHhceeccCCCCCCCCceEEEEEEeeeCCCCeE
Q orfY_Noce      17  SYFIPKPLKSQNIYEAFVRSRKKKEVDLEFREMLIYKTEQQFNCLKRLENRFDIYGKALPNEGQVYTFRFKLFDNTSKLV   96 (228)
Q Consensus      17  ~yFiPk~lk~~~iy~~~~~s~~Kkev~lE~~~~~~~~~~~~~~~lL~~lENRFwlY~~~~~~~~q~ysfcFks~D~~sk~~   96 (228)
                     +..-+..++.+.++.+++.+-.+..+.+++.+-..-.... +.--|-+.......++.........|...+..
T Consensus      11  ~~~~~~~~~~g~~v~g~V~~i~~~g~~v~~~~~~g~i~~--------~~~~~~~~~~~~vG~~i~~~V~~~~~~~~~~~   82 (281)
T 4Q7J_H        11  ESLKEIETRPGSIVRGVVVAIDKDVVLVDAGLKSESAIPA--------EQFKNAQGELEIQVGDEVDVALDAVEDGFGET   82 (281)
T ss_dssp            HHHHHHHHSCTTEECCEEEEECSSCEEEECSBSSCEEESC--------SSCTTTCSTTTCCTTCBCCEECCTTSSSSSST
T ss_pred            HhhhhcCCCCCCEEEEEEEEEcCCEEEEEcCCCceeEeeH--------HHHcccCCCcccCCCCEEEEEEEeeeCCCCeE

Q ss_pred            eeec-cccccchhHHHHHHHhcchhhhcCceeeehhheeeecCeEEEEecCCeEEE
Q orfY_Noce      97  NFSD-RKRRLQNISSYIYAYNSRRHILKNITKNIKSRLLCLTKGGFKISFFGQKVIL   152 (228)
Q Consensus      97  lFs~-~kkkl~~~~S~~~~~~n~r~~~l~~~~~liKsRlL~LtKgGf~~s~~G~~a~l   152 (228)
                     ..+. ....-..+......+...+ .+.+++......|.....-|..+++
T Consensus      83  ~ls~~~~~~~~~~~~~~~~~~~~~g~--------~v~~~V~~~~~~g~~v~~~~~~g~i   131 (281)
T 4Q7J_H        83  LLSREKAKRHEAWITLEKAYEDAE--------TVTGVINGKVKGGFTVELNGIRAFL   131 (281)
T ss_dssp            TCSSSHHHHHHHHHHHHHHHHHTCC--------CEEEEEEEECSSEEEEESSSCEEEE
T ss_pred            EEcHHHHHhHHHHHHHHHHHHHcCC--------cEEEEEeEECCEEEEEEECCEEEEe
```

# C

**4R71_E 30S ribosomal protein S1; OB FOLD, TRANSLATION, VIRAL PROTEIN-RIBOSOMAL PROTEIN complex; 3.21A {Escherichia coli}**

Probability: 26.15%,   E-value: 410,   Score: 20.33,   Aligned cols: 116,   Identities: 15%,   Similarity: 0.127,   Template Neff: 11.9

```
Q ss_pred            CcccccCceEEEEEEeCCcCceEEEEecccceeeehHHHHHhhcHHhceeccCCCCCCCCceEEEEEEeeeCCCCeEeeec
Q orfY_Noce      21  PKPLKSQNIYEAFVRSRKKKEVDLEFREMLIYKTEQQFNCLKRLENRFDIYGKALPNEGQVYTFRFKLFDNTSKLVNFSD   100 (228)
Q Consensus      21  Pk~lk~~~iy~~~~~s~~Kkev~lE~~~~~~~~~~~~~~~lL~~lENRFwlY~~~~~~~~q~ysfcFks~D~~sk~~lFs~   100 (228)
                     +..++.+.+..+.+..-....+.+++..-..-.... +.--|.+.......++.........|...+...++.
T Consensus      15  ~~~~~~g~~v~~~V~~~~~~~~~~~~v~~~~~~~~~~~~~~--------~~~~~~~~~~~~~~~g~~v~~~v~~~~~~~~~~~~~~~s~   86 (171)
T 4R71_E        15  EIETRPGSIVRGVVVAIDKDVVLVDAGLKSESAIPA--------EQFKNAQGELEIQVGDEVDVALDAVEDGFGETLLSR   86 (171)
T ss_dssp            HHTTSCTTEEEEEEEEECSSEEEEECSSSSCEEEEG--------GGGCCSSSCCCCCTTCEEEEECCCCC------CCTT
T ss_pred            hcCCCCCCEEEEEEEEEeCCEEEEEcCCCcEEEEEH--------HHHhccCCCccCCCCCEEEEEEEeeeCCCCeEEEeH

Q ss_pred            -cccccchhHHHHHHHhcchhhhcCceeeehhheeeecCeEEEEecCCeEEE
Q orfY_Noce     101  -RKRRLQNISSYIYAYNSRRHILKNITKNIKSRLLCLTKGGFKISFFGQKVIL   152 (228)
Q Consensus     101  -~kkkl~~~~S~~~~~~n~r~~~l~~~~~liKsRlL~LtKgGf~~s~~G~~a~l   152 (228)
                     ....-..+......+.... .+.+++....++|+...+-|..+++
T Consensus      87  ~~~~~~~~~~~~~~~~~~~~~~~g~--------~~~~~~v~~~~~~~~g~~v~~~~~~~~~~~   131 (171)
T 4R71_E        87  EKAKRHEAWITLEKAYEDAE--------TVTGVINGKVKGGFTVELNGIRAFL   131 (171)
T ss_dssp            TTTTCHHHHHHHHHHHHTTC--------EEEEEEEEECSSSEEEEETTEEEEE
T ss_pred            HHHHHhHHHHHHHHHHHHhCC--------CEEEEEEeeecCeEEEEECcEEEEe
```

**Fig. S6.** Evidence for the mitochondrial *orfY* gene being a divergent homolog of *rps1*. The figure shows pairwise alignments generated by a HHpred search against the profile HMM collection built for the sequences in the PDB database (PDB_mmCIF70_25_May), using as a query a profile HMM derived from a multiple alignment of *orfY*-encoded protein sequences. Only three hits were retrieved in the search, all corresponding to different representatives of the S1 protein, specifically the mitochondrial S1 from the green algal *Polytomella magna* (A) and two variants of the S1 protein from *Escherichia coli* (B and C). Note, however, the rather low probabilities and very high E-values of the hits, making the assignment of *orfY* as a *rps1* homolog tentative.

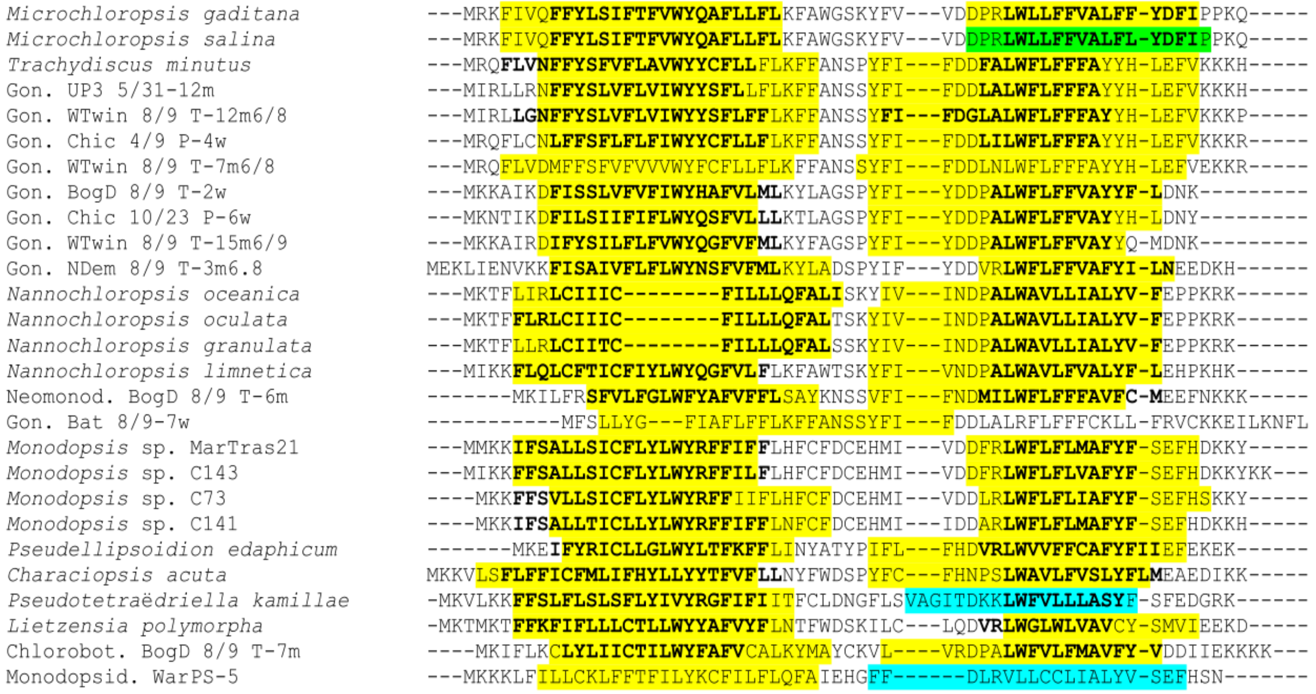


**Fig. S7.** Conservation of two transmembrane helices in *orfZ* gene products. The figure shows a multiple alignment of OrfZ sequences with the regions predicted by TMHMM 2.0 as exceeding the threshold for a TM helix highlighted in yellow. Regions with a predicted probability higher than 0.8 (yet beneath the threshold) are highlighted in green, whereas those with the probability from 0.2 to 0.8 are shown in blue. For comparison, TM helix predictions obtained using DeepTMHMM are indicated in boldface. Note that the OrfZ sequence from *Goniochloridales* sp. Bat 8/9-7w ("Gon. Bat 8/9-7w") is very divergent and may represent a degraded non-functional version of *orfZ*.

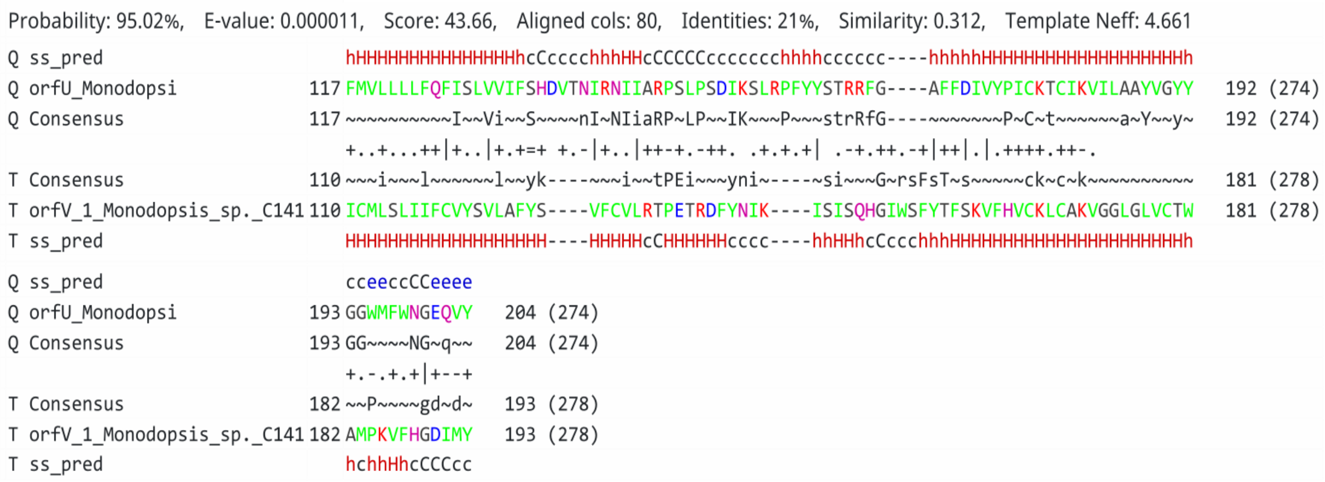


**Fig. S8.** Region of homology shared by OrfU and OrfV proteins, as detected by HHpred.

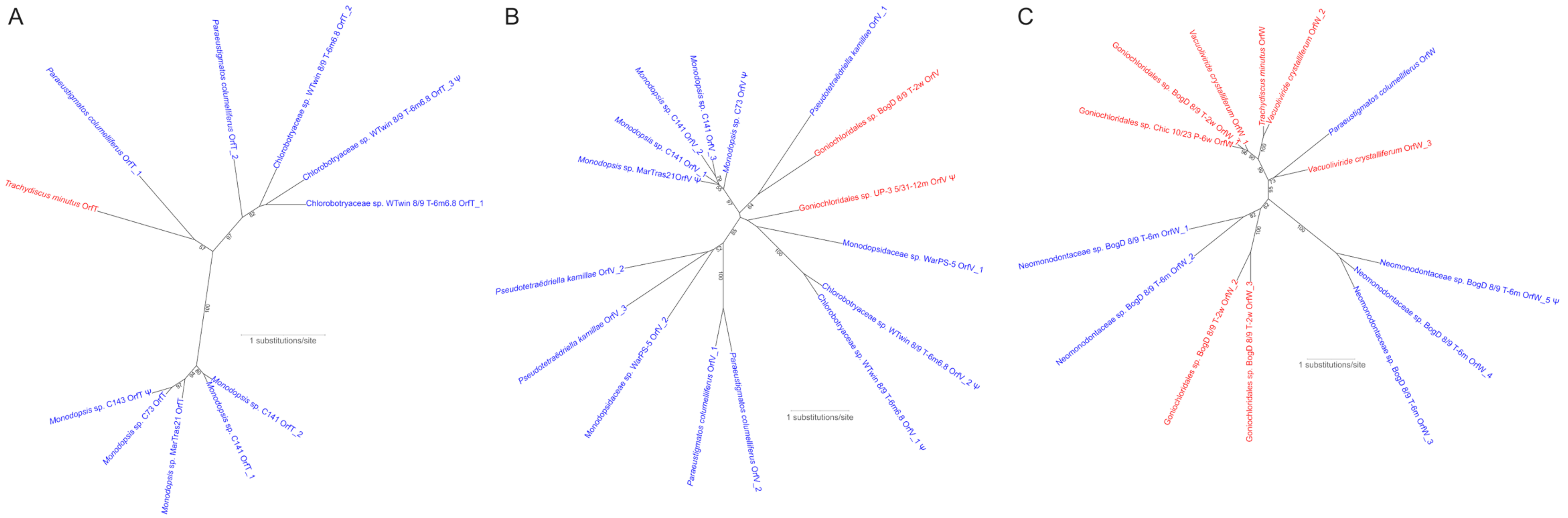


**Fig. S9.** Phylogenetic analysis of eustigmatophyte non-standard orfs that occur in multiple copies in particular mitogenomes. (A) OrfT, (B) OrfV, (C) OrfW. The trees were inferred with IQ-TREE from multiple sequence alignments comprising 202, 241, and 278 positions, respectively, using automatically selected substitutions models (JTT+F+I, mtInv+F+I+G4, and mtInv+F+G4, respectively). Non-parametric bootstrap values are shown when >50%. Sequences from Eustigmatales and *Goniochloridales* members are shown in blue and red, respectively. The Greek letter Ψ denotes obvious pseudogenes (only those representing a sufficiently complete sequence were included in the analysis). Note that the topology of the trees does not always recapitulate relationships of the organisms concerned.

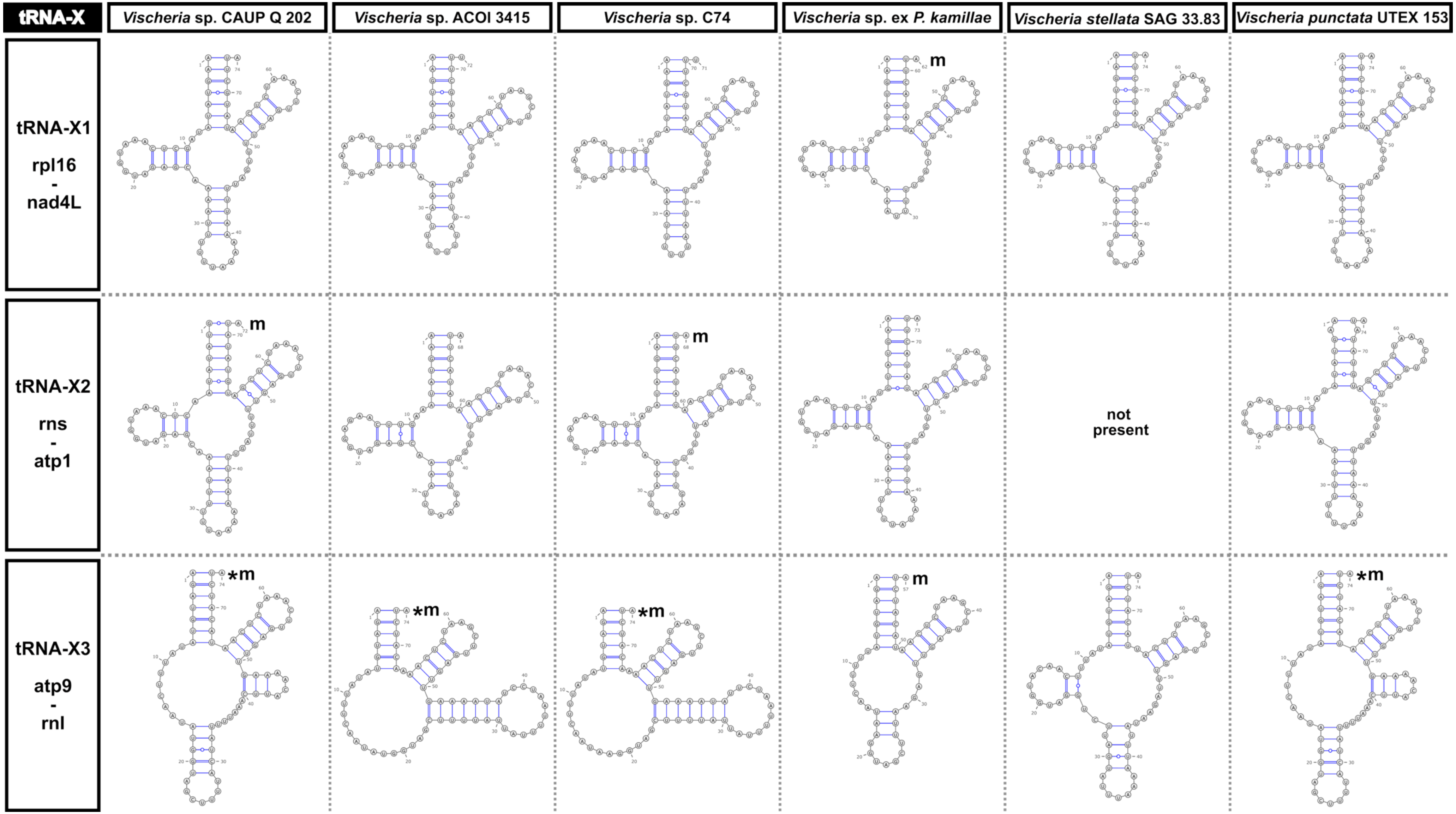


**Fig. S10.** Predicted secondary structure of tRNA-X1, tRNA-X2, and tRNA-X3 molecules specified by all *Vischeria* mitogenomes subjected to analysis. The tRNA structures marked by an asterisk were predicted by RNAfold, others by Rfam with default settings. Structures slightly manually modified compared to the original model obtained with Rfam or RNAfold are marked with “m”.

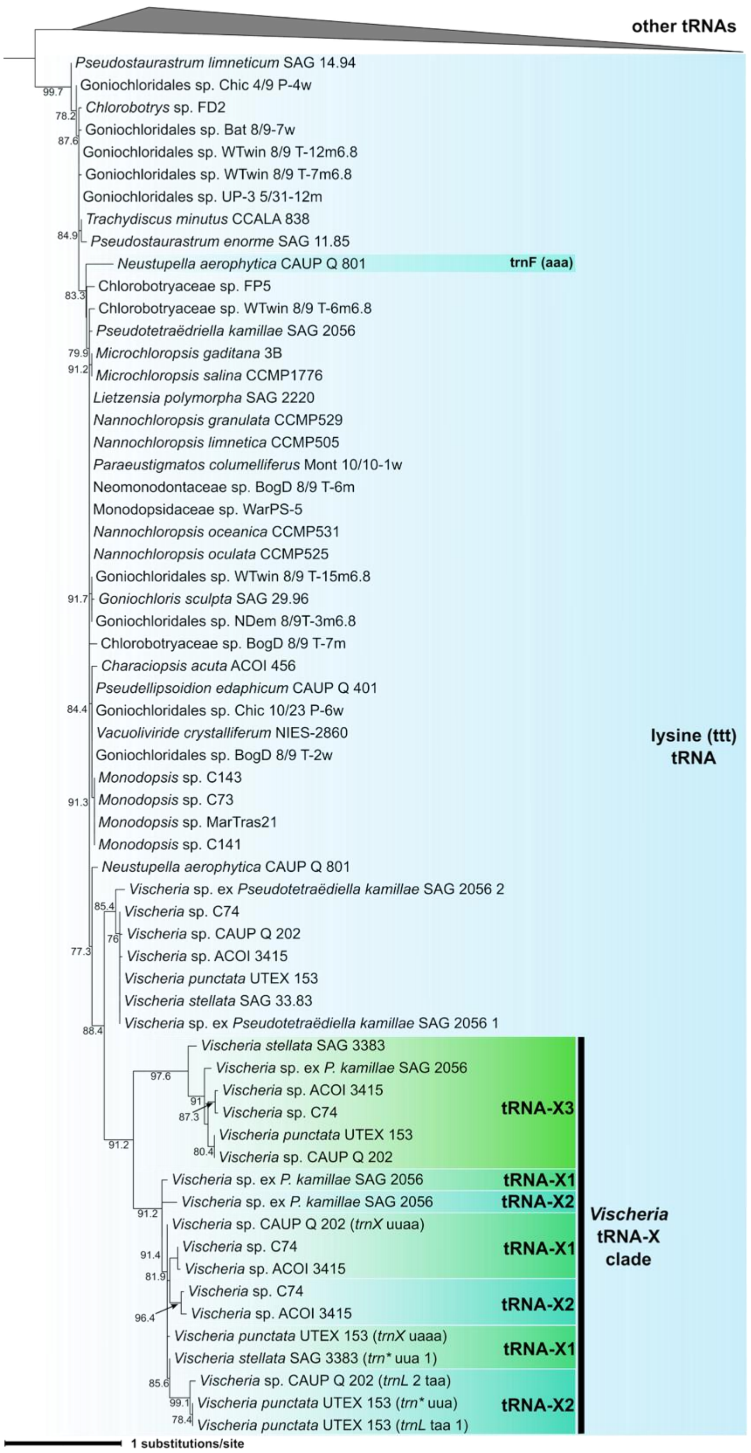


**Fig. S11.** Phylogenetic analysis of mitochondrial tRNA genes, focusing on the clade of *trnK(ttt)* genes and their non-standard evolutionary offshoots. The figure shows a simplified version of the full tree (available at Figshare, doi:10.6084/m9.figshare.31535176) inferred from a multiple alignment of all tRNA gene sequences specified by eustigmatophyte mitogenomes using IQ-TREE 1.5.5 (with the substitution model TVM+R6). Values at branches represent SH-aLRT support values (displayed when ≥75). Note that the clade dominated by eustig *trnK(ttt)* genes additionally contains a nested "*Vischeria* tRNA-X clade" that consists of more divergent non-canonical putative tRNA genes unique for species/strains of the genus Vischeria (different from the co-existing standard *trnK(ttt)* gene). The *Vischeria* tRNA-X clade itself is split into two subclades, one containing sequences of tRNA-X1 and tRNA-X2 genes intermingled (though their different identity is obvious from their differing conserved position in the *Vischeria* mitogenomes), and the other representing tRNA-X3 genes. In addition, the tree shows the existence of a non-standard evolutionary derivative of the *trnK(ttt)* gene found in the mitogenome of *Neustupella aerophytica*, labelled "trnF(aaa)" and potentially cognate to the UUU codon, although the amino acid specificity is inferred solely from the changed anticodon sequence. Note that *N. aerophytica* also possesses the standard *trnF(gaa)* gene that is expected to secure decoding of this codon.

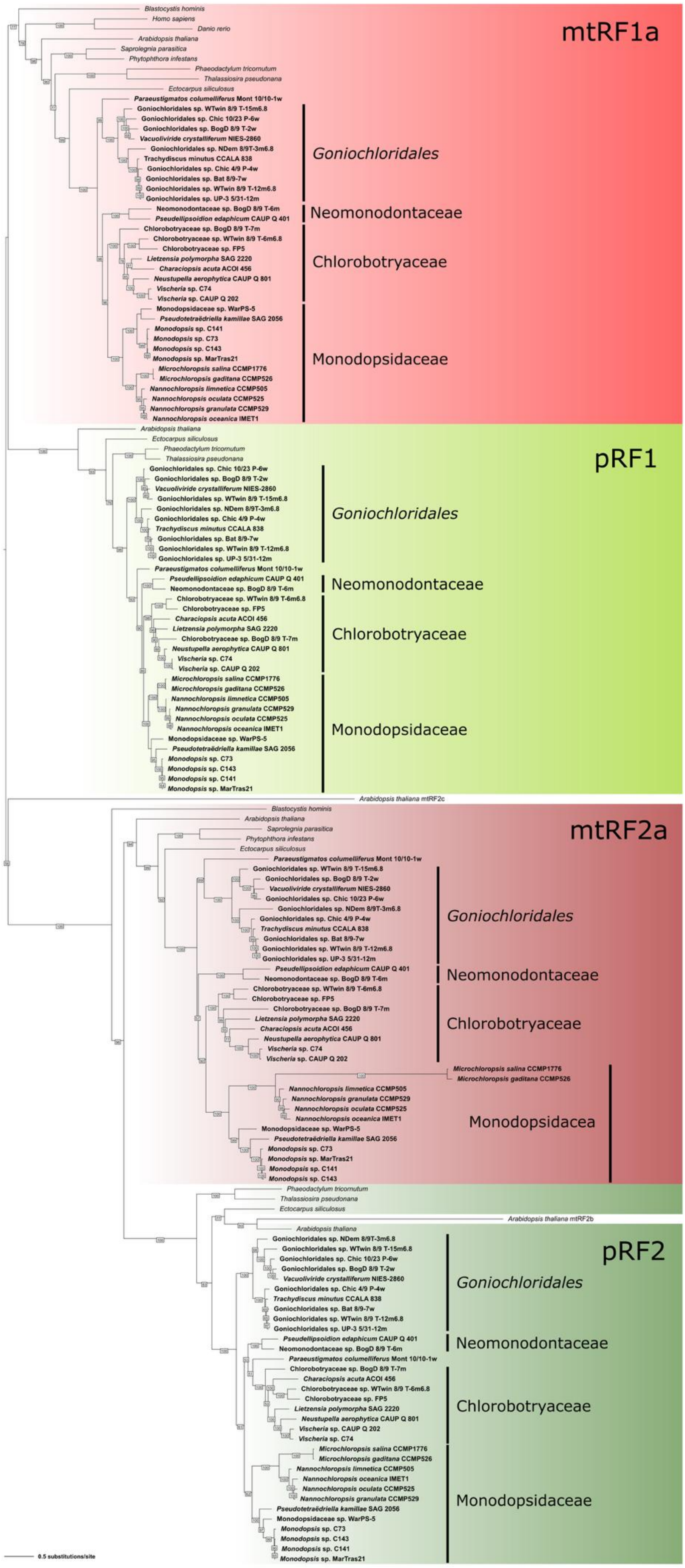


**Fig. S12.** ML phylogeny of organellar release factors. The tree is rooted between the two main RF groups (RF1 and RF2). The RF nomenclature is based on Duarte et al. (2012); each respective RF is marked by a colored box (mtRF1a – bright red, pRF1 – bright green, mtRF2a – dark red, pRF2 – dark green). Values at branches correspond to bootstrap values calculated from 1000 replicates (shown only when >50%). Eustigmatophyte species names are in bold.

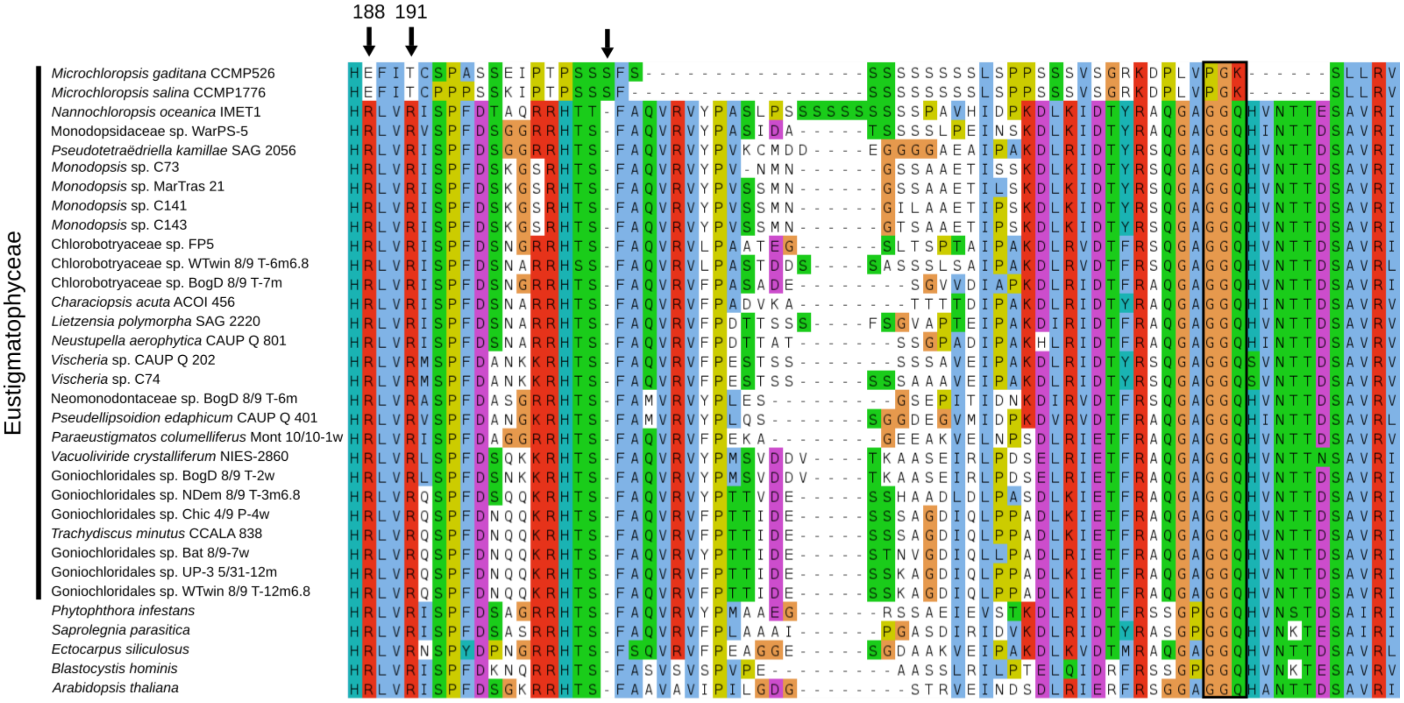


**Fig. S13.** Multiple alignment of mtRF2a sequences from eustigs and organisms used as an outgroup in the phylogeny displayed as Fig. S12. A region containing the anticodon recognition domain and the GGQ catalytic motif is shown. There are several mutations in mtRF2a from *Microchloropsis* spp. in positions that are highly conserved in other eustigs and are believed to be significant for changes in codon recognition (black arrows; the position numbering is based on the RF2 protein from *Thermus thermophilus;* for the discussion about the particular positions, see Žihala et al. (2020)). There is also an insertion next to the position 204 (non-numbered black arrow), which is considered to be crucial for the release factor to gain the ability to recognize the UUA codon. The GGQ catalytic motif is not present in *Microchloropsis* spp. (black rectangle), therefore their mtRF2a is unlikely to serve as a release factor in translation termination and its role remains unknown.

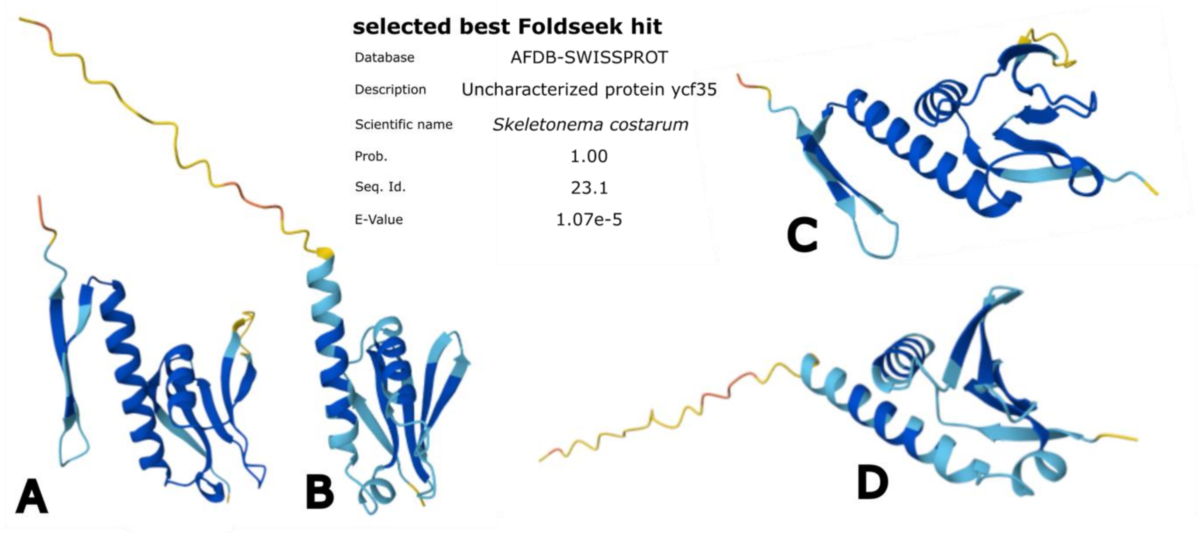


**Fig. S14.** Structural modeling supports the identification of a putative Ycf35 protein encoded by the plastid MAG TARA_CHLORO_00535. Querying the AlphaFold Protein Structure Database with the respective structural model generated by AlphaFold 3 retrieved known Ycf35 proteins as the best hits, here exemplified by the homolog from the diatom *Skeletonema costatum*. (A) and (C) show different views of the existing structural model of the *S. costatum* Ycf35 protein (note there is no experimentally determined structure of any Ycf35 protein available). For comparison, (B) and (D) show analogous views of a structural model of the putative Ycf35 encoded by TARA_CHLORO_00535. Note that the C-terminal region of the latter protein seems to stay unstructured, in contrast to the diatom homolog forming a beta strand pair in the corresponding region.
