## Supplementary Dataset S1 for "Insights into the phylogeny and enigmatic mitochondrial biology of eustigmatophyte algae from over 50 newly sequenced organellar genomes"

**Supplementary Dataset S1.** Gene maps of the newly generated eustigmatophyte plastid genomes. Individual genes are shown as blocks along the black line. Different gene types are highlighted in different colors (for details see the legends in lower left corner). Genes facing inwards are transcribed in the clockwise direction, genes facing outwards are transcribed in the counterclockwise direction. The inner circle shows two single copy regions and the inverted repeat (for the latter see also thickened line in outer black circle), and in grey the GC content (the fine inner line is 50%).

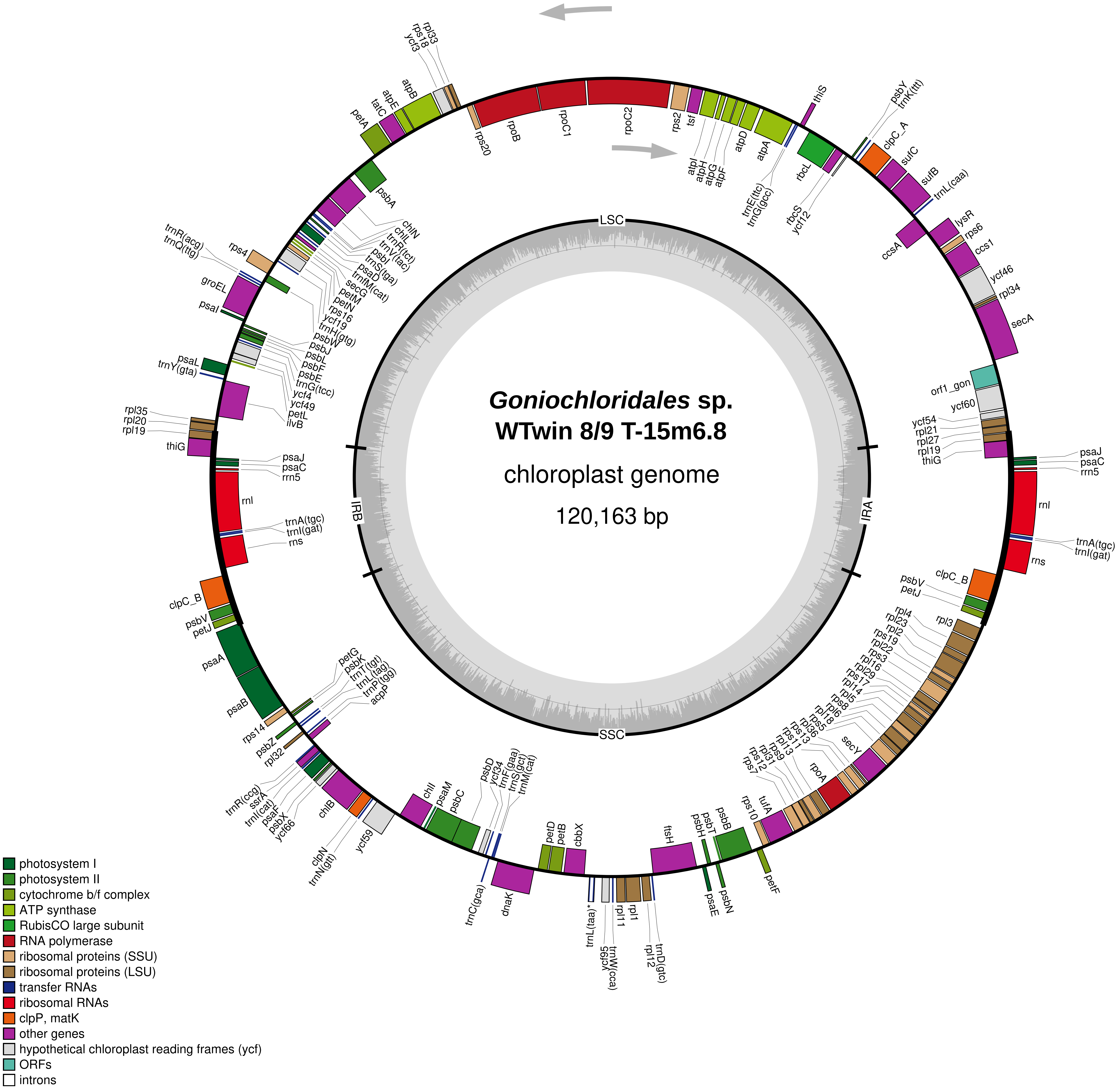

*Goniochloridales* sp.
WTwin 8/9 T-15m6.8
chloroplast genome
120,163 bp
LSC
IRA
SSC
IRB
rps20
rpoB
rpoC1
rpoC2
rps2
tsf
atpI
atpH
atpG
atpF
atpD
atpA
trnE(ttc)
trnG(gcc)
thiS
rbcL
rbcS
ycf12
psbY
trnK(ttt)
clpC_A
sufC
sufB
trnL(caa)
ccsA
lysR
rps6
ccs1
ycf46
rpl34
secA
orf1_gon
ycf60
ycf54
rpl21
rpl27
rpl19
thiG
psaJ
psaC
rrn5
rnl
trnA(tgc)
trnI(gat)
rns
clpC_B
psbV
petJ
rpl3
rpl4
rpl23
rpl2
rps19
rpl22
rps3
rpl16
rpl29
rps17
rpl14
rpl5
rps8
rpl6
rpl18
rps5
rpl36
rps13
rps11
rpl13
rps9
rpl31
rps12
rps7
secY
rpoA
rps10
tufA
petF
psbB
psbT
psbH
psbN
psaE
ftsH
rpl1
rpl11
rpl12
trnD(gtc)
trnW(cca)
ycf95
trnL(taa)*
cbbX
petB
petD
dnaK
trnC(gca)
psbD
ycf34
trnF(gaa)
trnS(gct)
trnM(cat)
psbC
psaM
chlI
ycf59
clpN
trnN(gtt)
chlB
ycf66
psbX
psaF
trnI(cat)
ssrA
trnR(ccg)
rpl32
psbZ
rps14
petG
psbK
trnT(tgt)
trnL(tag)
trnP(tgg)
acpP
psaB
psaA
petJ
psbV
clpC_B
rns
trnI(gat)
trnA(tgc)
rnl
rrn5
psaC
psaJ
thiG
rpl19
rpl20
rpl35
ilvB
petL
ycf49
ycf4
trnG(tcc)
psbE
psbF
psbL
psbJ
psbW
trnY(gta)
psaL
psaI
groEL
trnQ(ttg)
trnR(acg)
rps4
trnH(gtg)
ycf19
rps16
petN
petM
secG
trnfM(cat)
psaD
trnS(tga)
psbI
trnV(tac)
trnR(tct)
chlL
chlN
psbA
petA
tatC
atpE
atpB
ycf3
rps18
rpl33
photosystem I
photosystem II
cytochrome b/f complex
ATP synthase
RubisCO large subunit
RNA polymerase
ribosomal proteins (SSU)
ribosomal proteins (LSU)
transfer RNAs
ribosomal RNAs
clpP, matK
other genes
hypothetical chloroplast reading frames (ycf)
ORFs
introns

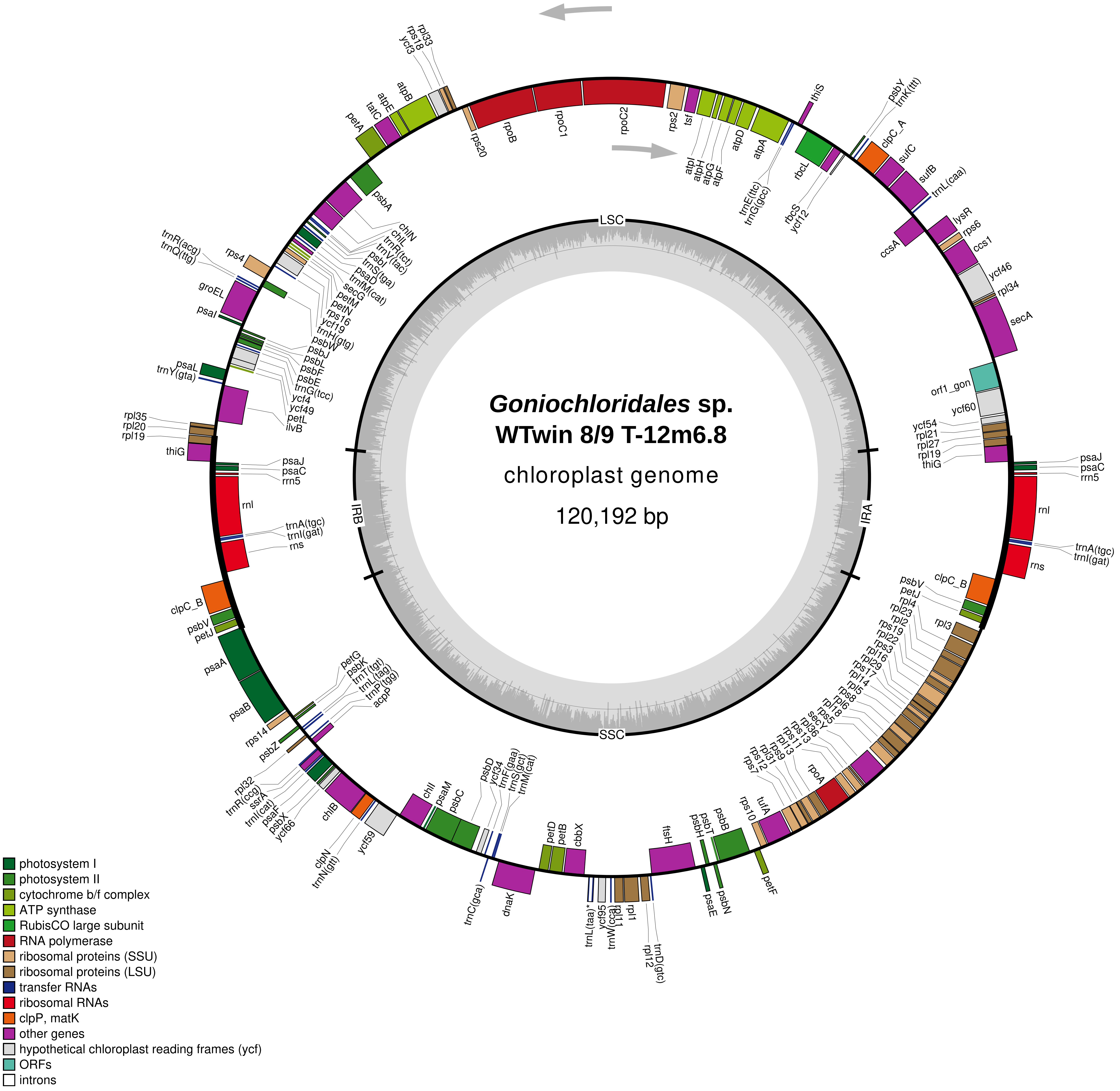

*Goniochloridales* sp.
WTwin 8/9 T-12m6.8
chloroplast genome
120,192 bp
LSC
IRA
SSC
IRB
rps20
rpoB
rpoC1
rpoC2
rps2
tsf
atpI
atpH
atpG
atpF
atpD
atpA
trnE(ttc)
trnG(gcc)
thiS
rbcL
rbcS
ycf12
psbY
trnK(ttt)
clpC_A
sufC
sufB
trnL(caa)
lysR
rps6
ccsA
ccs1
ycf46
rpl34
secA
orf1_gon
ycf60
ycf54
rpl21
rpl27
rpl19
thiG
psaJ
psaC
rrn5
rnl
trnA(tgc)
trnI(gat)
rns
clpC_B
psbV
petJ
rpl4
rpl23
rpl2
rps19
rpl22
rps3
rpl16
rpl29
rps17
rpl14
rpl5
rps8
rpl6
rpl18
rps5
secY
rpl36
rps13
rps11
rpl13
rps9
rpl31
rps12
rps7
rpoA
rps10
tufA
psbB
psbT
psbH
petF
psbN
psaE
ftsH
trnD(gtc)
rpl12
rpl1
rpl11
trnW(cca)
ycf95
trnL(taa)*
cbbX
petB
petD
dnaK
trnC(gca)
trnM(cat)
trnS(gct)
trnF(gaa)
ycf34
psbD
psbC
psaM
chlI
ycf59
trnN(gtt)
clpN
chlB
ycf66
psbX
psaF
trnI(cat)
ssrA
trnR(ccg)
rpl32
psbZ
rps14
psaB
psaA
acpP
trnP(tgg)
trnL(tag)
trnT(tgt)
psbK
petG
petJ
psbV
clpC_B
rns
trnI(gat)
trnA(tgc)
rnl
rrn5
psaC
psaJ
thiG
rpl19
rpl20
rpl35
ilvB
petL
ycf49
ycf4
trnG(tcc)
psbE
psbF
psbL
psbJ
psbW
trnH(gtg)
ycf19
rps16
petN
petM
secG
trnfM(cat)
psaD
trnS(tga)
psbI
trnV(tac)
trnR(tct)
chlL
chlN
psbA
psaL
trnY(gta)
psaI
groEL
trnQ(ttg)
trnR(acg)
rps4
petA
tatC
atpE
atpB
ycf3
rps18
rpl33
photosystem I
photosystem II
cytochrome b/f complex
ATP synthase
RubisCO large subunit
RNA polymerase
ribosomal proteins (SSU)
ribosomal proteins (LSU)
transfer RNAs
ribosomal RNAs
clpP, matK
other genes
hypothetical chloroplast reading frames (ycf)
ORFs
introns

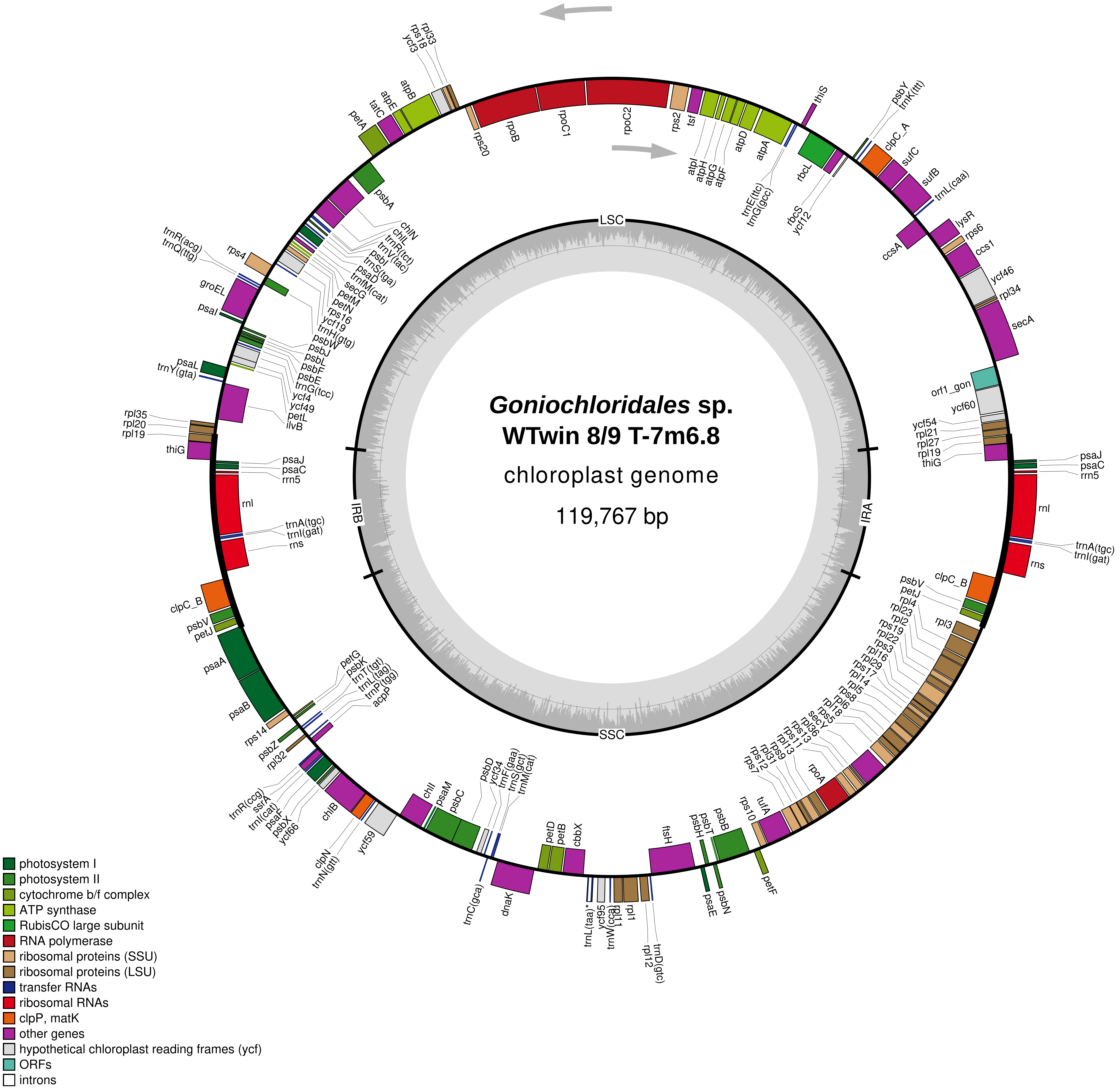

*Goniochloridales* sp.
WTwin 8/9 T-7m6.8
chloroplast genome
119,767 bp
LSC
IRA
SSC
IRB
rpl33
rps18
ycf3
atpB
atpE
tatC
petA
rps20
rpoB
rpoC1
rpoC2
rps2
tsf
atpI
atpH
atpG
atpF
atpD
atpA
trnE(ttc)
trnG(gcc)
thiS
rbcL
rbcS
ycf12
psbY
trnK(ttt)
clpC_A
sufC
sufB
trnL(caa)
lysR
rps6
ccsA
ccs1
ycf46
rpl34
secA
orf1_gon
ycf60
ycf54
rpl21
rpl27
rpl19
thiG
psaJ
psaC
rrn5
rnl
trnA(tgc)
trnI(gat)
rns
clpC_B
psbV
petJ
rpl4
rpl23
rpl2
rps19
rpl22
rps3
rpl16
rpl29
rps17
rpl14
rpl5
rps8
rpl6
rpl18
rps5
secY
rpl36
rps13
rps11
rpl13
rps9
rpl31
rps12
rps7
rpoA
rps10
tufA
petF
psbB
psbT
psbH
psbN
psaE
ftsH
trnD(gtc)
rpl12
rpl1
rpl11
trnW(cca)
ycf95
trnL(taa)*
cbbX
petB
petD
dnaK
trnC(gca)
trnM(cat)
trnS(gct)
trnF(gaa)
ycf34
psbD
psbC
psaM
chlI
ycf59
trnN(gtt)
clpN
chlB
ycf66
psbX
psaF
trnI(cat)
ssrA
trnR(ccg)
rpl32
psbZ
rps14
psaB
psaA
acpP
trnP(tgg)
trnL(tag)
trnT(tgt)
psbK
petG
ilvB
petL
ycf49
ycf4
trnG(tcc)
psbE
psbF
psbL
psbJ
psbW
trnH(gtg)
ycf19
rps16
petN
petM
secG
trnfM(cat)
psaD
trnS(tga)
psbI
trnV(tac)
trnR(tct)
chlL
chlN
psbA
rps4
trnR(acg)
trnQ(ttg)
groEL
psaI
psaL
trnY(gta)
rpl35
rpl20
photosystem I
photosystem II
cytochrome b/f complex
ATP synthase
RubisCO large subunit
RNA polymerase
ribosomal proteins (SSU)
ribosomal proteins (LSU)
transfer RNAs
ribosomal RNAs
clpP, matK
other genes
hypothetical chloroplast reading frames (ycf)
ORFs
introns

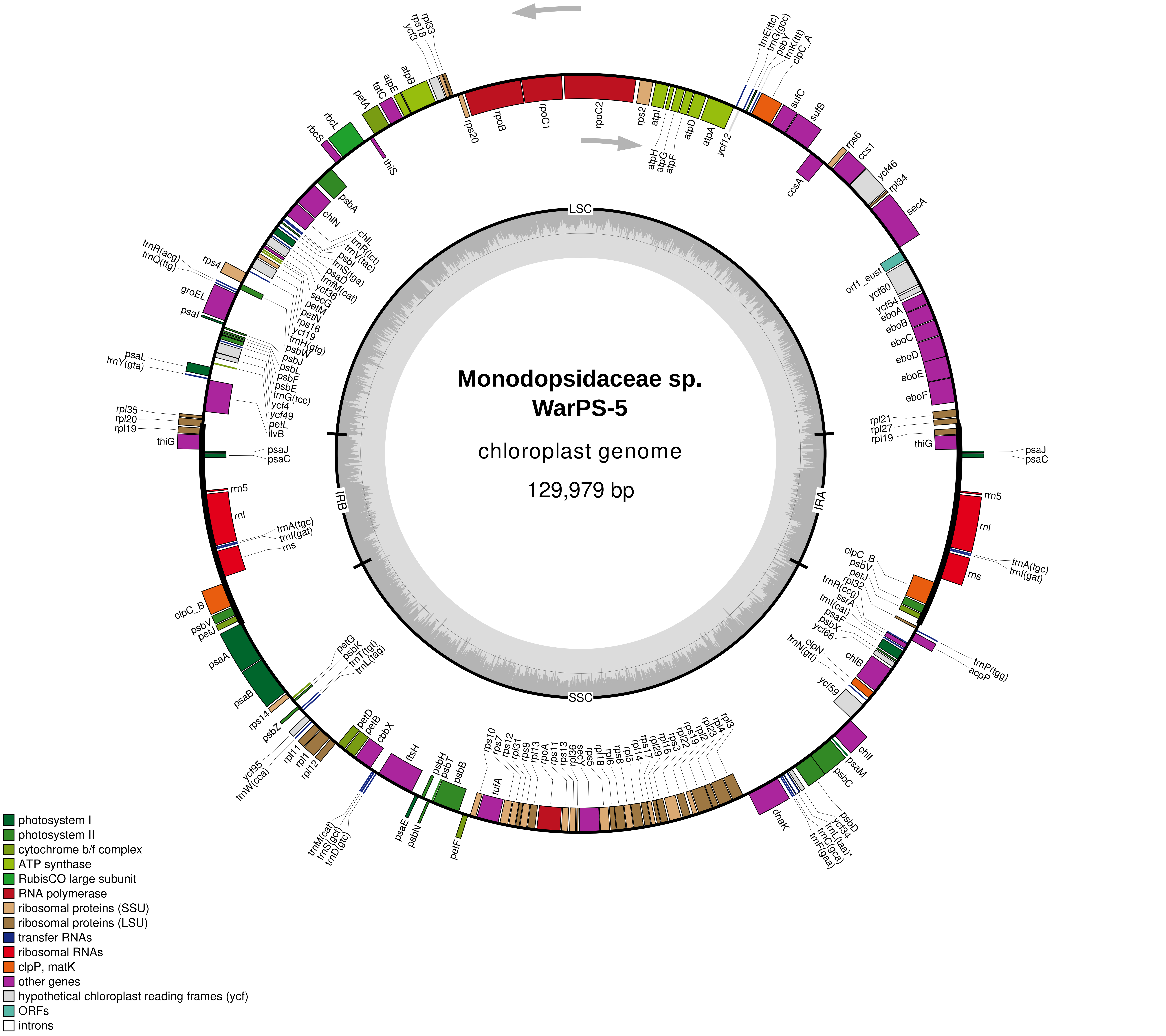
Monodopsidaceae sp. WarPS-5
chloroplast genome
129,979 bp
LSC
IRA
SSC
IRB
photosystem I
photosystem II
cytochrome b/f complex
ATP synthase
RubisCO large subunit
RNA polymerase
ribosomal proteins (SSU)
ribosomal proteins (LSU)
transfer RNAs
ribosomal RNAs
clpP, matK
other genes
hypothetical chloroplast reading frames (ycf)
ORFs
introns

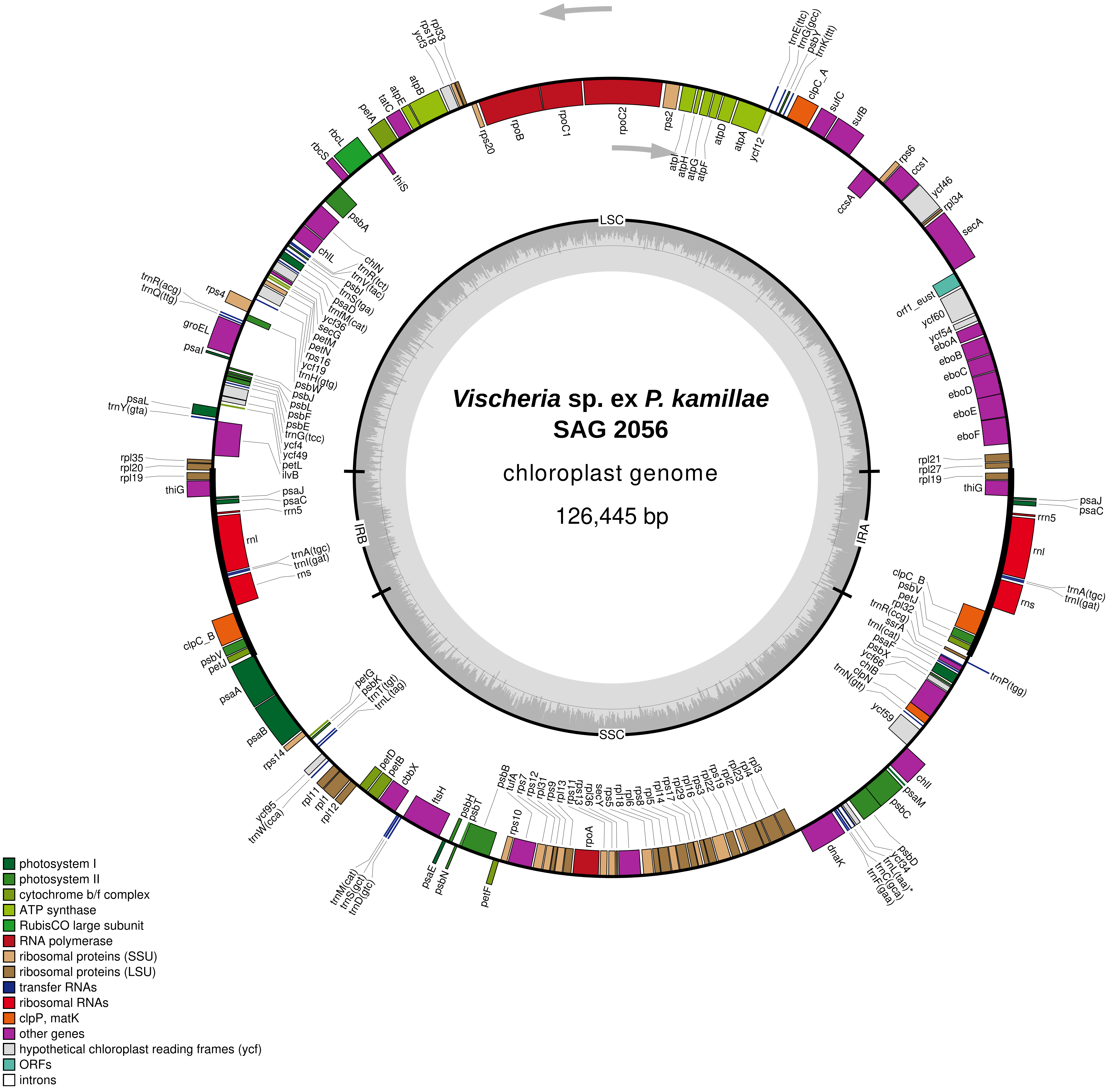

Vischeria sp. ex P. kamillae
SAG 2056
chloroplast genome
126,445 bp
LSC
IRA
SSC
IRB
rpl33
rps18
ycf3
atpB
atpE
tatC
petA
rbcL
rbcS
thiS
rps20
rpoB
rpoC1
rpoC2
rps2
atpI
atpH
atpG
atpF
atpD
atpA
ycf12
trnE(ttc)
trnG(gcc)
psbY
trnK(ttt)
clpC_A
sufC
sufB
rps6
ccs1
ccsA
ycf46
rpl34
secA
orf1_eust
ycf60
ycf54
eboA
eboB
eboC
eboD
eboE
eboF
rpl21
rpl27
rpl19
thiG
psaJ
psaC
rrn5
rnl
trnA(tgc)
trnI(gat)
rns
clpC_B
psbV
petJ
rpl32
trnR(ccg)
ssrA
trnI(cat)
psaF
psbX
ycf66
chlB
clpN
trnN(gtt)
trnP(tgg)
ycf59
chlI
psaM
psbC
psbD
ycf34
trnL(taa)*
trnC(gca)
trnF(gaa)
dnaK
rpl3
rpl4
rpl2
rpl23
rps19
rpl22
rps3
rpl16
rpl29
rps17
rpl14
rpl5
rps8
rpl6
rpl18
rps5
secY
rpl36
rps13
rps11
rpl13
rps9
rpl31
rps12
rps7
tufA
psbB
rps10
rpoA
petF
psbH
psbT
psbE
psbN
psaE
ftsH
trnM(cat)
trnS(gct)
trnD(gtc)
petD
petB
cbbX
rpl11
rpl1
rpl12
ycf95
trnW(cca)
rps14
psaB
psaA
petG
psbK
trnT(tgt)
trnL(tag)
psbA
chlL
chlN
trnR(tct)
trnV(tac)
psbI
trnS(tga)
psaD
trnfM(cat)
ycf36
secG
petM
petN
rps16
ycf19
trnH(gtg)
psbW
psbJ
psbL
psbF
psbE
trnG(tcc)
ycf4
ycf49
petL
ilvB
rps4
trnR(acg)
trnQ(ttg)
groEL
psaI
psaL
trnY(gta)
rpl35
rpl20
rpl19
thiG
psaJ
psaC
rrn5
rnl
trnA(tgc)
trnI(gat)
rns
clpC_B
psbV
petJ
photosystem I
photosystem II
cytochrome b/f complex
ATP synthase
RubisCO large subunit
RNA polymerase
ribosomal proteins (SSU)
ribosomal proteins (LSU)
transfer RNAs
ribosomal RNAs
clpP, matK
other genes
hypothetical chloroplast reading frames (ycf)
ORFs
introns

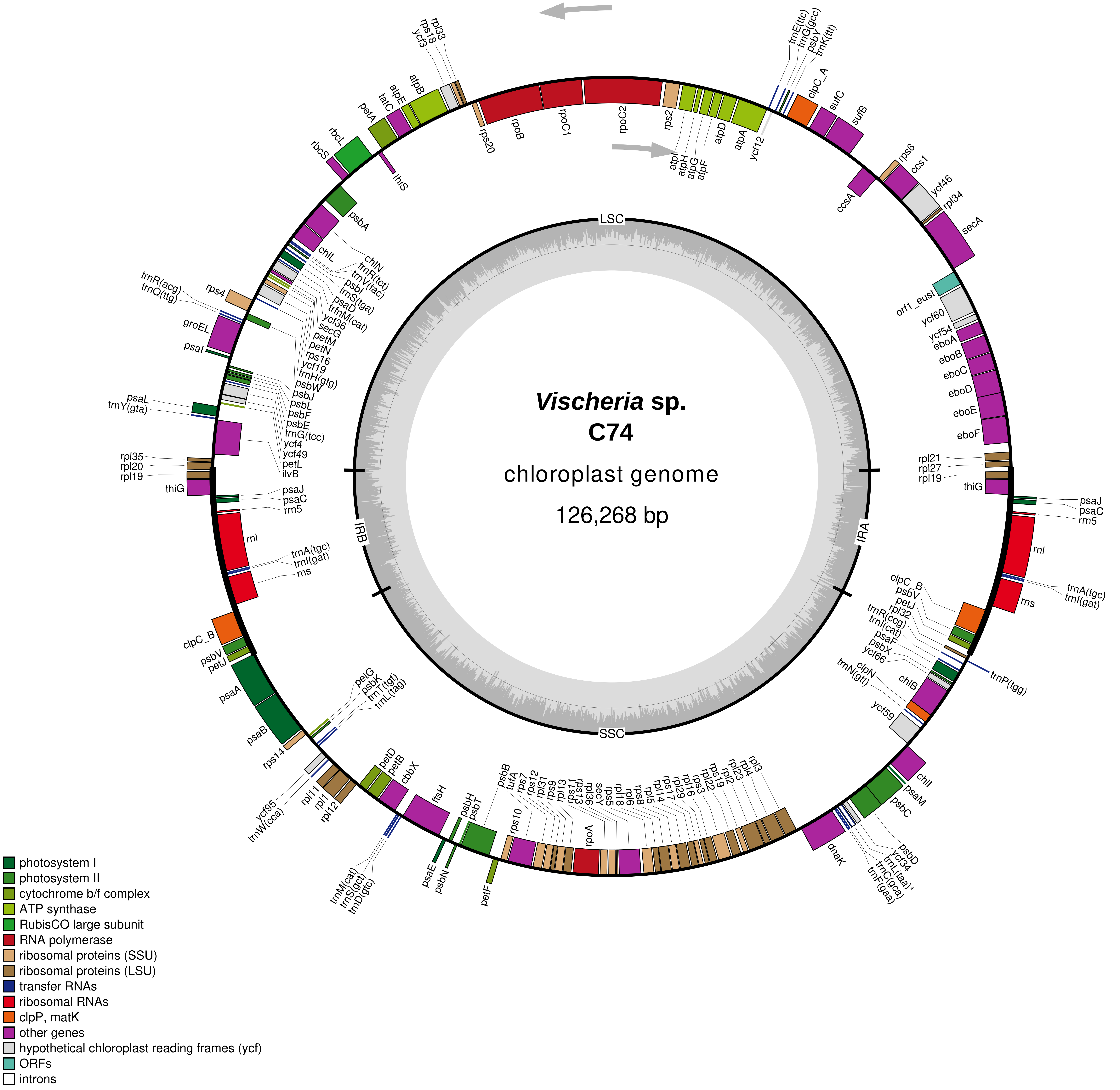

Vischeria sp.
C74
chloroplast genome
126,268 bp
LSC
IRA
SSC
IRB
rpl33
rps18
ycf3
atpB
atpE
tatC
petA
rbcL
rbcS
thiS
rps20
rpoB
rpoC1
rpoC2
rps2
atpI
atpH
atpG
atpF
atpD
atpA
ycf12
trnE(ttc)
trnG(gcc)
psbY
trnK(ttt)
clpC_A
sufC
sufB
rps6
ccs1
ccsA
ycf46
rpl34
secA
orf1_eust
ycf60
ycf54
eboA
eboB
eboC
eboD
eboE
eboF
rpl21
rpl27
rpl19
thiG
psaJ
psaC
rrn5
rnl
trnA(tgc)
trnI(gat)
rns
clpC_B
psbV
petJ
rpl32
trnR(ccg)
trnI(cat)
psaF
psbX
ycf66
chlB
clpN
trnN(gtt)
trnP(tgg)
ycf59
chlI
psaM
psbC
psbD
ycf34
trnL(taa)*
trnC(gca)
trnF(gaa)
dnaK
rpl3
rpl4
rpl23
rpl2
rps19
rpl22
rps3
rpl16
rpl29
rps17
rpl14
rpl5
rps8
rpl6
rpl18
rps5
secY
rpl36
rps13
rps11
rpl13
rps9
rpl31
rps12
rps7
tufA
psbB
rps10
rpoA
petF
psbH
psbT
psbE
psbN
psaE
ftsH
cbbX
petB
petD
trnM(cat)
trnS(gct)
trnD(gtc)
rpl11
rpl1
rpl12
ycf95
trnW(cca)
rps14
psaB
psaA
petG
psbK
trnT(tgt)
trnL(tag)
petJ
psbV
clpC_B
rns
trnI(gat)
trnA(tgc)
rnl
rrn5
psaC
psaJ
thiG
rpl19
rpl20
rpl35
psaL
trnY(gta)
ilvB
petL
ycf49
ycf4
trnG(tcc)
psbE
psbF
psbL
psbJ
psbW
trnH(gtg)
ycf19
rps16
petN
petM
secG
ycf36
trfnM(cat)
psaD
trnS(tga)
psbI
trnV(tac)
trnR(tct)
chlN
chlL
psbA
psaI
groEL
trnQ(ttg)
trnR(acg)
rps4
photosystem I
photosystem II
cytochrome b/f complex
ATP synthase
RubisCO large subunit
RNA polymerase
ribosomal proteins (SSU)
ribosomal proteins (LSU)
transfer RNAs
ribosomal RNAs
clpP, matK
other genes
hypothetical chloroplast reading frames (ycf)
ORFs
introns

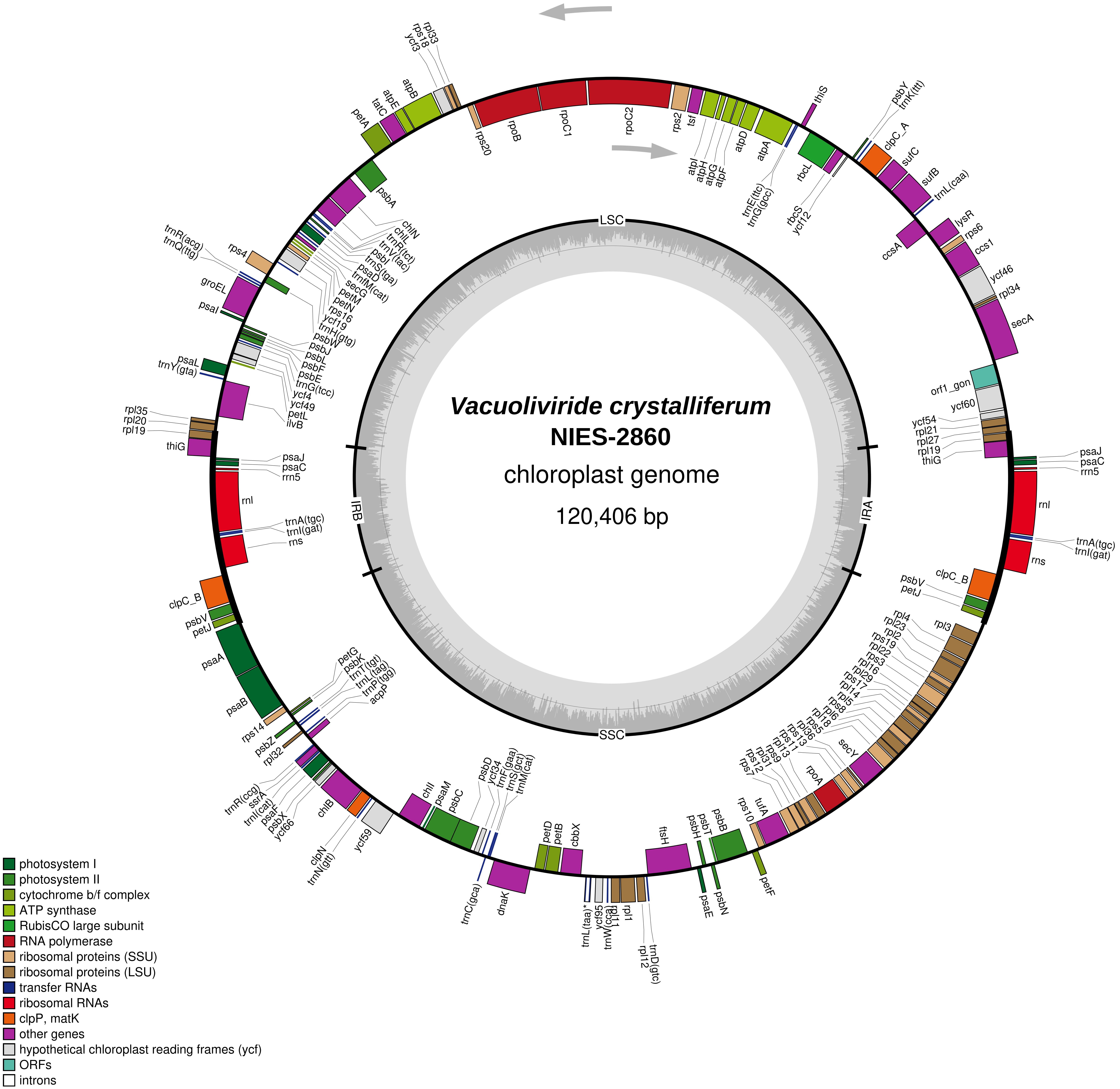

Vacuoliviride crystalliferum
NIES-2860
chloroplast genome
120,406 bp
LSC
IRA
SSC
IRB
rpl33
rps18
ycf3
atpB
atpE
tatC
petA
rps20
rpoB
rpoC1
rpoC2
rps2
tsf
atpI
atpH
atpG
atpF
atpD
atpA
trnE(ttc)
trnG(gcc)
thiS
rbcL
rbcS
ycf12
psbY
trnK(ttt)
clpC_A
sufC
sufB
trnL(caa)
lysR
rps6
ccsA
ccs1
ycf46
rpl34
secA
orf1_gon
ycf60
ycf54
rpl21
rpl27
rpl19
thiG
psaJ
psaC
rrn5
rnl
trnA(tgc)
trnI(gat)
rns
clpC_B
psbV
petJ
rpl3
rpl4
rpl23
rpl2
rps19
rpl22
rps3
rpl16
rpl29
rps17
rpl14
rpl5
rps8
rpl6
rpl18
rps5
secY
rpl36
rps13
rps11
rpoA
rps9
rpl31
rps12
rps7
rps10
tufA
petF
psbB
psbT
psbH
psbN
psaE
ftsH
rpl1
rpl11
rpl12
trnD(gtc)
trnW(cca)
ycf95
trnL(taa)*
cbbX
petB
petD
dnaK
trnC(gca)
psbD
ycf34
trnF(gaa)
trnS(gct)
trnM(cat)
psbC
psaM
chlI
ycf59
clpN
trnN(gtt)
chlB
ycf66
psbX
psaF
trnI(cat)
ssrA
trnR(ccg)
rpl32
psbZ
rps14
psaB
psaA
petG
psbK
trnT(tgt)
trnL(tag)
trnP(tgg)
acpP
rpl35
rpl20
thiG
psaL
trnY(gta)
ilvB
petL
ycf49
ycf4
trnG(tcc)
psbE
psbF
psbL
psbJ
psbW
trnH(gtg)
ycf19
rps16
petN
petM
secG
trnfM(cat)
psaD
trnS(tga)
psbI
trnV(tac)
trnR(tct)
chlL
chlN
psbA
rps4
trnR(acg)
trnQ(ttg)
groEL
psaI
photosystem I
photosystem II
cytochrome b/f complex
ATP synthase
RubisCO large subunit
RNA polymerase
ribosomal proteins (SSU)
ribosomal proteins (LSU)
transfer RNAs
ribosomal RNAs
clpP, matK
other genes
hypothetical chloroplast reading frames (ycf)
ORFs
introns

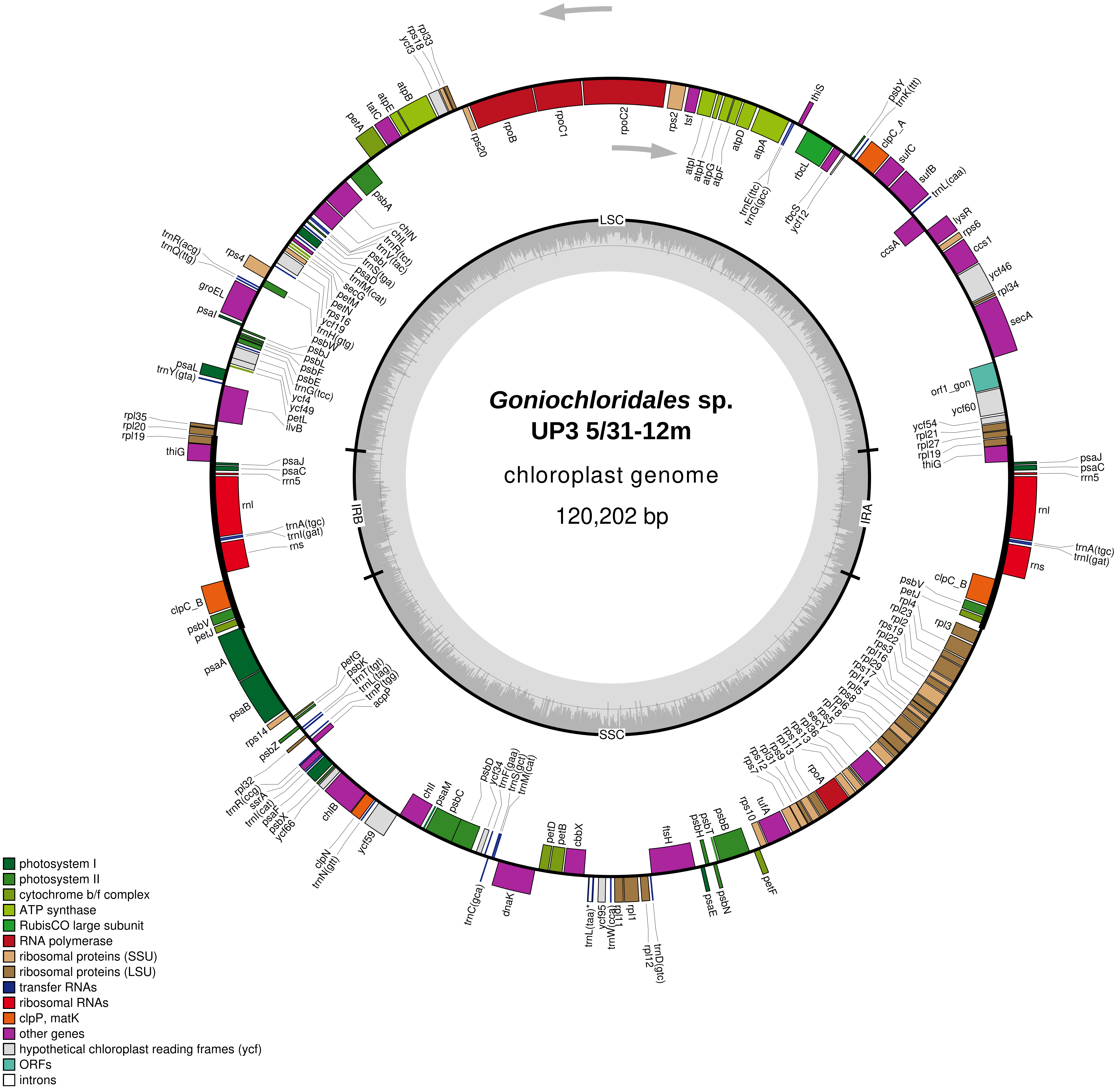

Goniochloridales sp.
UP3 5/31-12m
chloroplast genome
120,202 bp
LSC
IRA
SSC
IRB
rpl33
rps18
ycf3
atpB
atpE
tatC
petA
rps20
rpoB
rpoC1
rpoC2
rps2
tsf
atpI
atpH
atpG
atpF
atpD
atpA
trnE(ttc)
trnG(gcc)
thiS
rbcL
rbcS
ycf12
psbY
trnK(ttt)
clpC_A
sufC
sufB
trnL(caa)
lysR
rps6
ccsA
ccs1
ycf46
rpl34
secA
orf1_gon
ycf60
ycf54
rpl21
rpl27
rpl19
thiG
psaJ
psaC
rrn5
rnl
trnA(tgc)
trnI(gat)
rns
clpC_B
psbV
petJ
rpl4
rpl23
rpl2
rps19
rpl22
rps3
rpl16
rpl29
rps17
rpl14
rpl5
rps8
rpl6
rpl18
rps5
secY
rpl36
rps13
rps11
rpl13
rps9
rpl31
rps12
rps7
rpoA
rps10
tufA
psbB
psbT
psbH
psaE
psbN
petF
ftsH
trnD(gtc)
rpl12
rpl1
rpl11
trnW(cca)
ycf95
trnL(taa)*
cbbX
petB
petD
dnaK
trnC(gca)
psbD
ycf34
trnF(gaa)
trnS(gct)
trnM(cat)
psbC
psaM
chlI
ycf59
clpN
trnN(gtt)
chlB
ycf66
psbX
psaF
trnI(cat)
ssrA
trnR(ccg)
rpl32
psbZ
rps14
psaB
psaA
petG
psbK
trnT(tgt)
trnL(tag)
trnP(tgg)
acpP
petJ
psbV
clpC_B
rns
trnI(gat)
trnA(tgc)
rnl
rrn5
psaC
psaJ
thiG
rpl19
rpl20
rpl35
trnY(gta)
psaL
ilvB
petL
ycf49
ycf4
trnG(tcc)
psbE
psbF
psbL
psbJ
psbW
trnH(gtg)
ycf19
rps16
petN
petM
secG
trnfM(cat)
psaD
trnS(tga)
psbI
trnV(tac)
trnR(tct)
chlL
chlN
psbA
psaI
groEL
trnQ(ttg)
trnR(acg)
rps4
photosystem I
photosystem II
cytochrome b/f complex
ATP synthase
RubisCO large subunit
RNA polymerase
ribosomal proteins (SSU)
ribosomal proteins (LSU)
transfer RNAs
ribosomal RNAs
clpP, matK
other genes
hypothetical chloroplast reading frames (ycf)
ORFs
introns

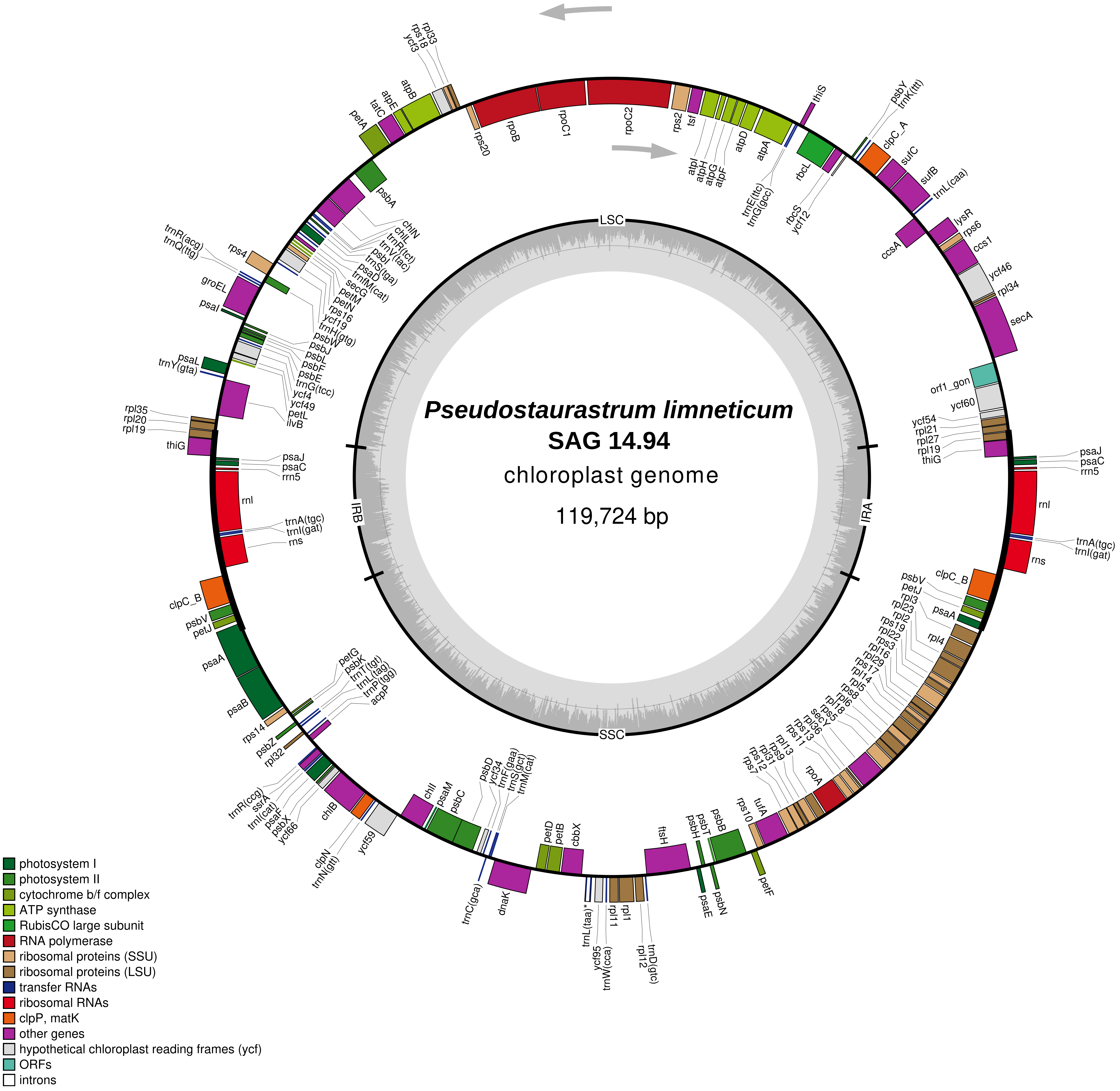

Pseudostaurastrum limneticum
SAG 14.94
chloroplast genome
119,724 bp
LSC
SSC
IRA
IRB
rpl33
rps18
ycf3
atpB
atpE
tatC
petA
rps20
rpoB
rpoC1
rpoC2
rps2
tsf
atpI
atpH
atpG
atpF
atpD
atpA
trnE(ttc)
trnG(gcc)
thiS
rbcL
rbcS
ycf12
psbY
trnK(ttt)
clpC_A
sufC
sufB
trnL(caa)
lysR
rps6
ccsA
ccs1
ycf46
rpl34
secA
psbA
chlN
chlL
trnR(tct)
trnV(tac)
psbI
trnS(tga)
psaD
trnfM(cat)
secG
petM
petN
rps16
ycf19
trnH(gtg)
psbW
psbJ
psbL
psbF
psbE
trnG(tcc)
ycf4
ycf49
petL
ilvB
trnR(acg)
trnQ(ttg)
rps4
groEL
psaI
psaL
trnY(gta)
rpl35
rpl20
rpl19
thiG
orf1_gon
ycf60
ycf54
rpl21
rpl27
rpl19
thiG
psaJ
psaC
rrn5
rnl
trnA(tgc)
trnI(gat)
rns
psaJ
psaC
rrn5
rnl
trnA(tgc)
trnI(gat)
rns
clpC_B
psbV
petJ
psaA
psaB
clpC_B
psbV
petJ
psaA
rpl3
rpl23
rpl2
rps19
rpl22
rps3
rpl16
rpl29
rps17
rpl14
rpl5
rps8
rpl6
rpl18
rps5
secY
rpl36
rps13
rps11
rpl13
rps9
rpl31
rps12
rps7
rpoA
rpl4
petG
psbK
trnT(tgt)
trnL(tag)
trnP(tgg)
acpP
rps14
psbZ
rpl32
trnR(ccg)
ssrA
trnI(cat)
psaF
psbX
ycf66
chlB
clpN
trnN(gtt)
ycf59
chlI
psaM
psbC
psbD
ycf34
trnF(gaa)
trnS(gct)
trnM(cat)
trnC(gca)
dnaK
petD
petB
cbbX
trnL(taa)*
ycf95
trnW(cca)
rpl11
rpl1
rpl12
trnD(gtc)
ftsH
psbH
psbT
psbB
psaE
psbN
rps10
tufA
petF
photosystem I
photosystem II
cytochrome b/f complex
ATP synthase
RubisCO large subunit
RNA polymerase
ribosomal proteins (SSU)
ribosomal proteins (LSU)
transfer RNAs
ribosomal RNAs
clpP, matK
other genes
hypothetical chloroplast reading frames (ycf)
ORFs
introns

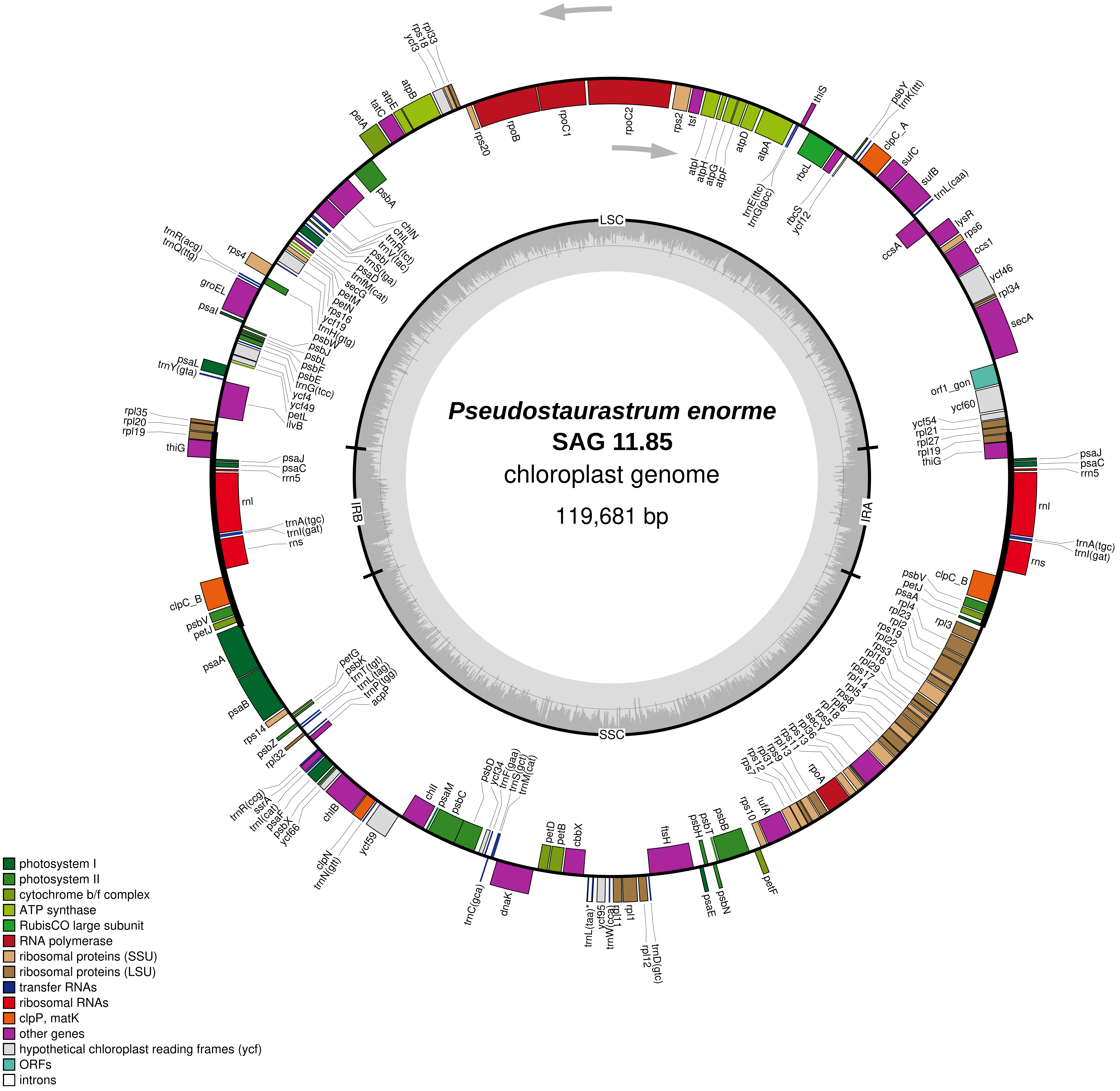

Pseudostaurastrum enorme
SAG 11.85
chloroplast genome
119,681 bp
LSC
IRA
SSC
IRB
rps20
rpoB
rpoC1
rpoC2
rps2
tsf
atpI
atpH
atpG
atpF
atpD
atpA
trnE(ttc)
trnG(gcc)
thiS
rbcL
rbcS
ycf12
psbY
trnK(ttt)
clpC_A
sufC
sufB
trnL(caa)
ccsA
lysR
rps6
ccs1
ycf46
rpl34
secA
orf1_gon
ycf60
ycf54
rpl21
rpl27
rpl19
thiG
psaJ
psaC
rrn5
rnl
trnA(tgc)
trnI(gat)
rns
clpC_B
psbV
petJ
psaA
rpl4
rpl23
rpl2
rps19
rpl22
rps3
rpl16
rpl29
rps17
rpl14
rpl5
rps8
rpl6
rpl18
rps5
secY
rpl36
rps13
rps11
rpl13
rps9
rpl31
rps12
rps7
rpoA
rps10
tufA
petF
psbB
psbT
psbH
psaE
psbN
ftsH
trnD(gtc)
rpl12
rpl1
rpl11
trnW(cca)
ycf95
trnL(taa)*
cbbX
petB
petD
dnaK
trnC(gca)
trnM(cat)
trnS(gct)
trnF(gaa)
ycf34
psbD
psbC
psaM
chlI
ycf59
trnN(gtt)
clpN
chlB
ycf66
psbX
psaF
trnI(cat)
ssrA
trnR(ccg)
rpl32
psbZ
rps14
psaB
acpP
trnP(tgg)
trnL(tag)
trnT(tgt)
psbK
petG
rpl35
rpl20
rpl19
thiG
ilvB
petL
ycf49
ycf4
trnG(tcc)
psbE
psbF
psbL
psbJ
psbW
trnH(gtg)
ycf19
rps16
petN
petM
secG
trnfM(cat)
psaD
trnS(tga)
psbI
trnV(tac)
trnR(tct)
chlL
chlN
psbA
petA
tatC
atpE
atpB
ycf3
rps18
rpl33
rps4
trnR(acg)
trnQ(ttg)
groEL
psaI
psaL
trnY(gta)
photosystem I
photosystem II
cytochrome b/f complex
ATP synthase
RubisCO large subunit
RNA polymerase
ribosomal proteins (SSU)
ribosomal proteins (LSU)
transfer RNAs
ribosomal RNAs
clpP, matK
other genes
hypothetical chloroplast reading frames (ycf)
ORFs
introns

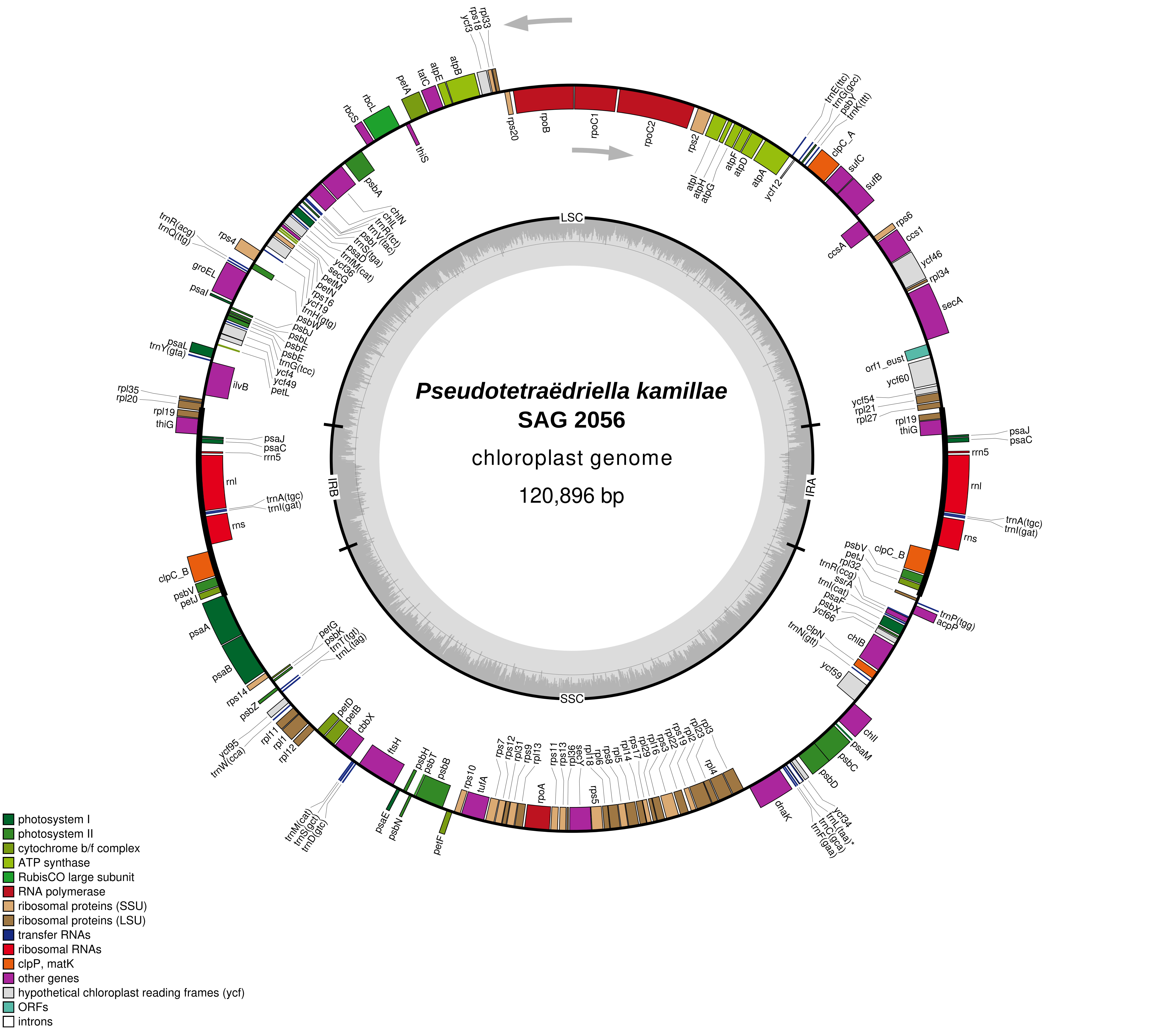

Pseudotetraëdriella kamillae
SAG 2056
chloroplast genome
120,896 bp
LSC
IRA
SSC
IRB
photosystem I
photosystem II
cytochrome b/f complex
ATP synthase
RubisCO large subunit
RNA polymerase
ribosomal proteins (SSU)
ribosomal proteins (LSU)
transfer RNAs
ribosomal RNAs
clpP, matK
other genes
hypothetical chloroplast reading frames (ycf)
ORFs
introns

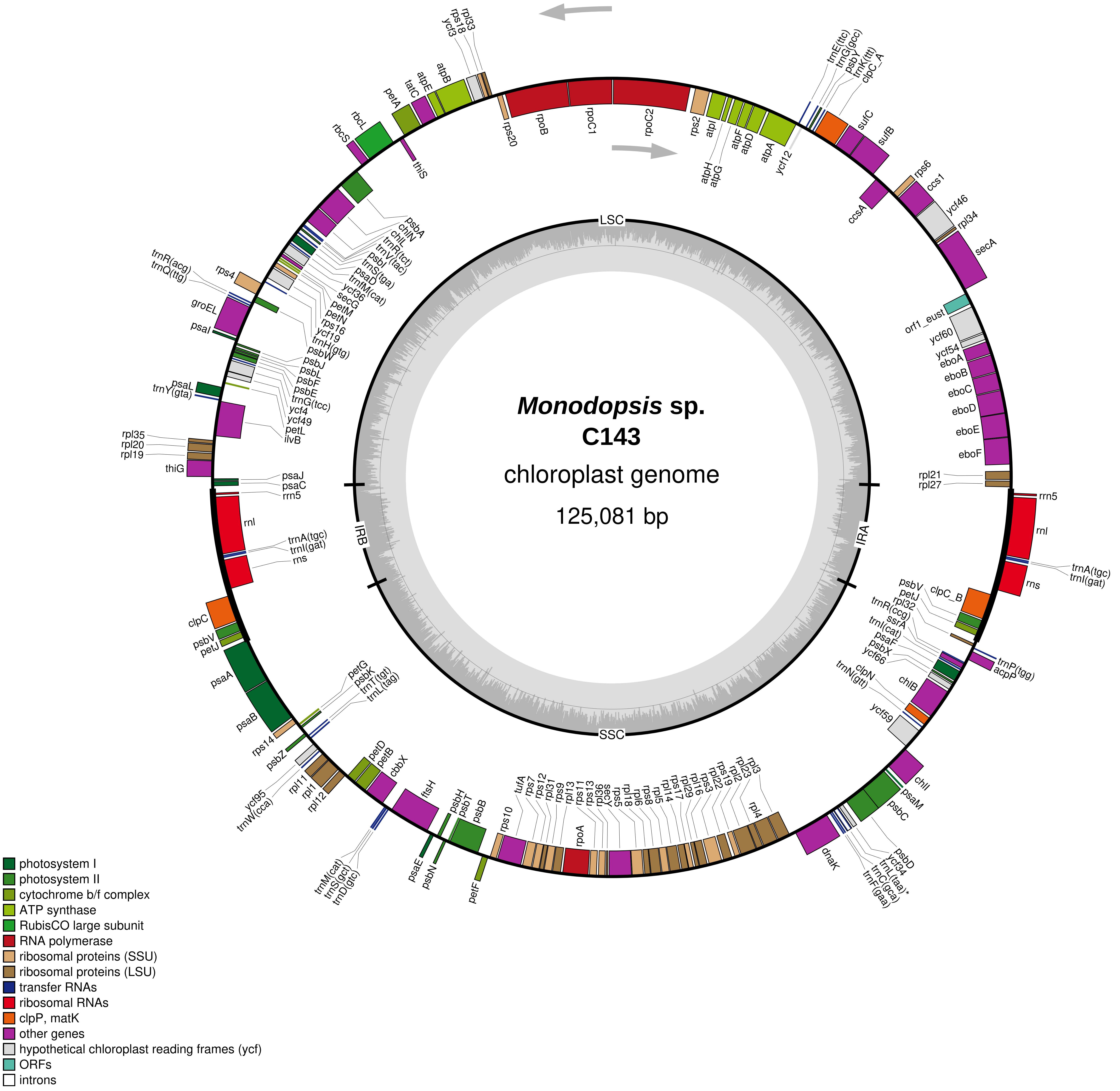

Monodopsis sp.
C143
chloroplast genome
125,081 bp
LSC
IRA
SSC
IRB
rpl33
rps18
ycf3
atpB
atpE
tatC
petA
rbcL
rbcS
thiS
rps20
rpoB
rpoC1
rpoC2
rps2
atpI
atpH
atpG
atpF
atpD
atpA
ycf12
trnE(ttc)
trnG(gcc)
psbY
trnK(ttt)
clpC_A
sufC
sufB
rps6
ccs1
ccsA
ycf46
rpl34
secA
orf1_eust
ycf60
ycf54
eboA
eboB
eboC
eboD
eboE
eboF
rpl21
rpl27
rrn5
rnl
trnA(tgc)
trnI(gat)
rns
psbV
petJ
clpC_B
rpl32
trnR(ccg)
ssrA
trnI(cat)
psaF
psbX
ycf66
trnP(tgg)
acpP
clpN
trnN(gtt)
chlB
ycf59
chlI
psaM
psbC
psbD
ycf34
trnL(taa)*
trnC(gca)
trnF(gaa)
dnaK
rpl3
rpl23
rpl2
rps19
rpl22
rps3
rpl16
rpl29
rps17
rpl14
rpl5
rps8
rpl6
rpl18
rps5
secY
rpl36
rps13
rps11
rpl13
rps9
rpl31
rps12
rps7
tufA
rpoA
rpl4
rps10
petF
psbB
psbT
psbH
psbN
psaE
ftsH
cbbX
petB
petD
trnM(cat)
trnS(gct)
trnD(gtc)
rpl11
rpl1
rpl12
ycf95
trnW(cca)
psbZ
rps14
psaB
psaA
petG
psbK
trnT(tgt)
trnL(tag)
psbV
petJ
clpC
rrn5
rnl
trnA(tgc)
trnI(gat)
rns
psaJ
psaC
thiG
rpl35
rpl20
rpl19
ilvB
psaL
trnY(gta)
petL
ycf49
ycf4
trnG(tcc)
psbE
psbF
psbL
psbJ
psbW
trnH(gtg)
ycf19
rps16
petN
petM
secG
ycf36
trnfM(cat)
psaD
trnS(tga)
psbI
trnV(tac)
trnR(tct)
chlL
chlN
psbA
psaI
groEL
rps4
trnR(acg)
trnQ(ttg)
photosystem I
photosystem II
cytochrome b/f complex
ATP synthase
RubisCO large subunit
RNA polymerase
ribosomal proteins (SSU)
ribosomal proteins (LSU)
transfer RNAs
ribosomal RNAs
clpP, matK
other genes
hypothetical chloroplast reading frames (ycf)
ORFs
introns

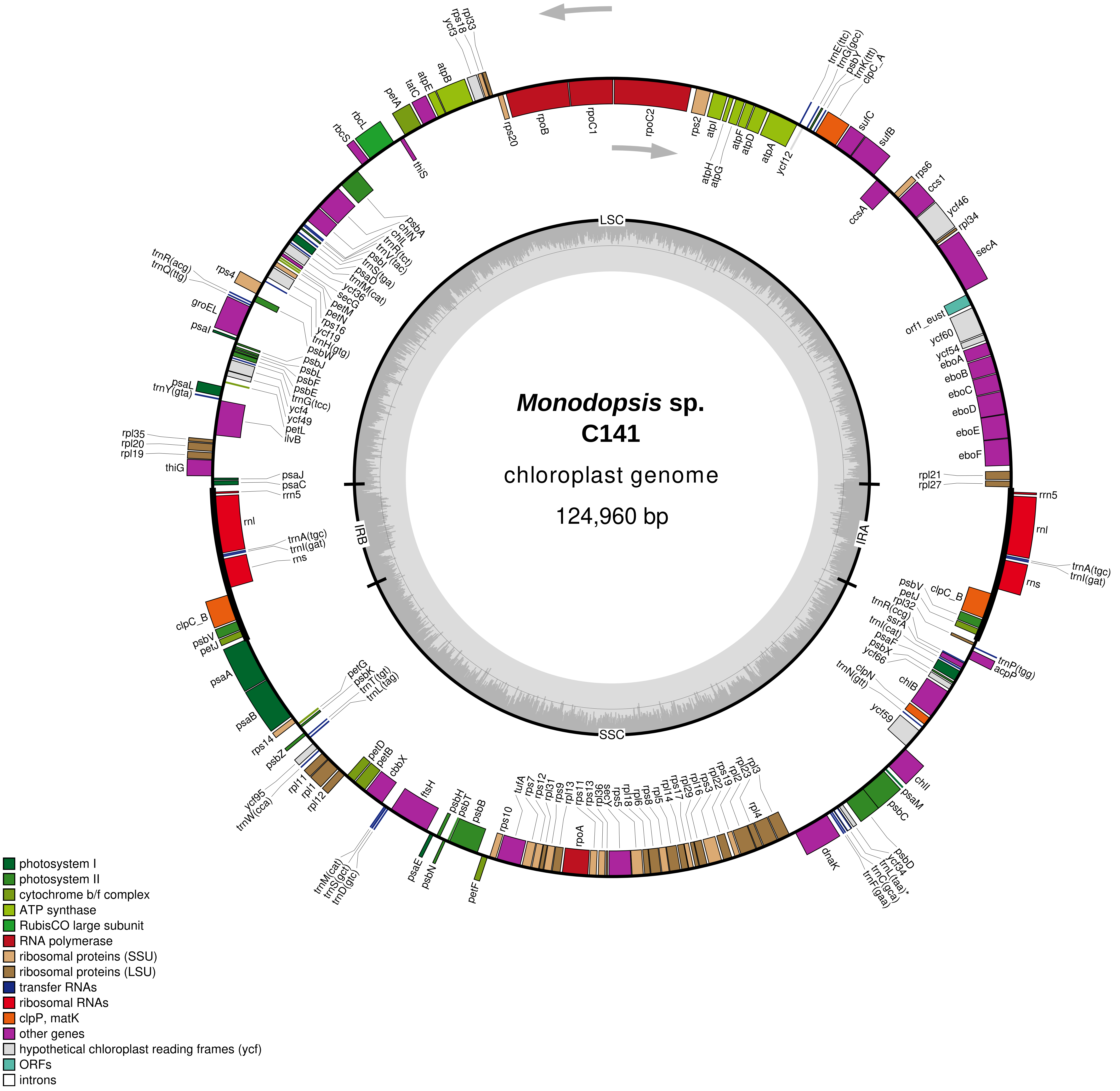

*Monodopsis* sp.
C141
chloroplast genome
124,960 bp
LSC
IRA
SSC
IRB
rpl33
rps18
ycf3
atpB
atpE
tatC
petA
rbcL
rbcS
thiS
rps20
rpoB
rpoC1
rpoC2
rps2
atpI
atpH
atpG
atpF
atpD
atpA
ycf12
trnE(ttc)
trnG(gcc)
psbY
trnK(ttt)
clpC_A
sufC
sufB
rps6
ccs1
ccsA
ycf46
rpl34
secA
orf1_eust
ycf60
ycf54
eboA
eboB
eboC
eboD
eboE
eboF
rpl21
rpl27
rrn5
rnl
trnA(tgc)
trnI(gat)
rns
psbV
petJ
rpl32
clpC_B
trnR(ccg)
ssrA
trnI(cat)
psaF
psbX
ycf66
trnP(tgg)
acpP
chlB
clpN
trnN(gtt)
ycf59
chlI
psaM
psbC
psbD
ycf34
trnL(taa)*
trnC(gca)
trnF(gaa)
dnaK
rpl3
rpl23
rpl2
rps19
rpl22
rps3
rpl16
rpl29
rps17
rpl14
rpl5
rps8
rpl6
rpl18
rps5
secY
rpl36
rps13
rps11
rpl13
rps9
rpl31
rps12
rps7
tufA
rpoA
rpl4
rps10
psbB
psbT
psbH
psbN
psaE
petF
ftsH
cbbX
petB
petD
trnM(cat)
trnS(gct)
trnD(gtc)
rpl11
rpl1
rpl12
ycf95
trnW(cca)
psbZ
rps14
petG
psbK
trnT(tgt)
trnL(tag)
psaB
psaA
petJ
psbV
clpC_B
rns
trnI(gat)
trnA(tgc)
rnl
rrn5
psaC
psaJ
thiG
rpl19
rpl20
rpl35
ilvB
petL
ycf49
ycf4
trnG(tcc)
psbE
psbF
psbL
psbJ
psbW
trnY(gta)
psaL
psaI
groEL
trnQ(ttg)
trnR(acg)
rps4
trnH(gtg)
ycf19
rps16
petN
petM
secG
ycf36
trnfM(cat)
psaD
trnS(tga)
psbI
trnV(tac)
trnR(tct)
chlL
chlN
psbA
photosystem I
photosystem II
cytochrome b/f complex
ATP synthase
RubisCO large subunit
RNA polymerase
ribosomal proteins (SSU)
ribosomal proteins (LSU)
transfer RNAs
ribosomal RNAs
clpP, matK
other genes
hypothetical chloroplast reading frames (ycf)
ORFs
introns

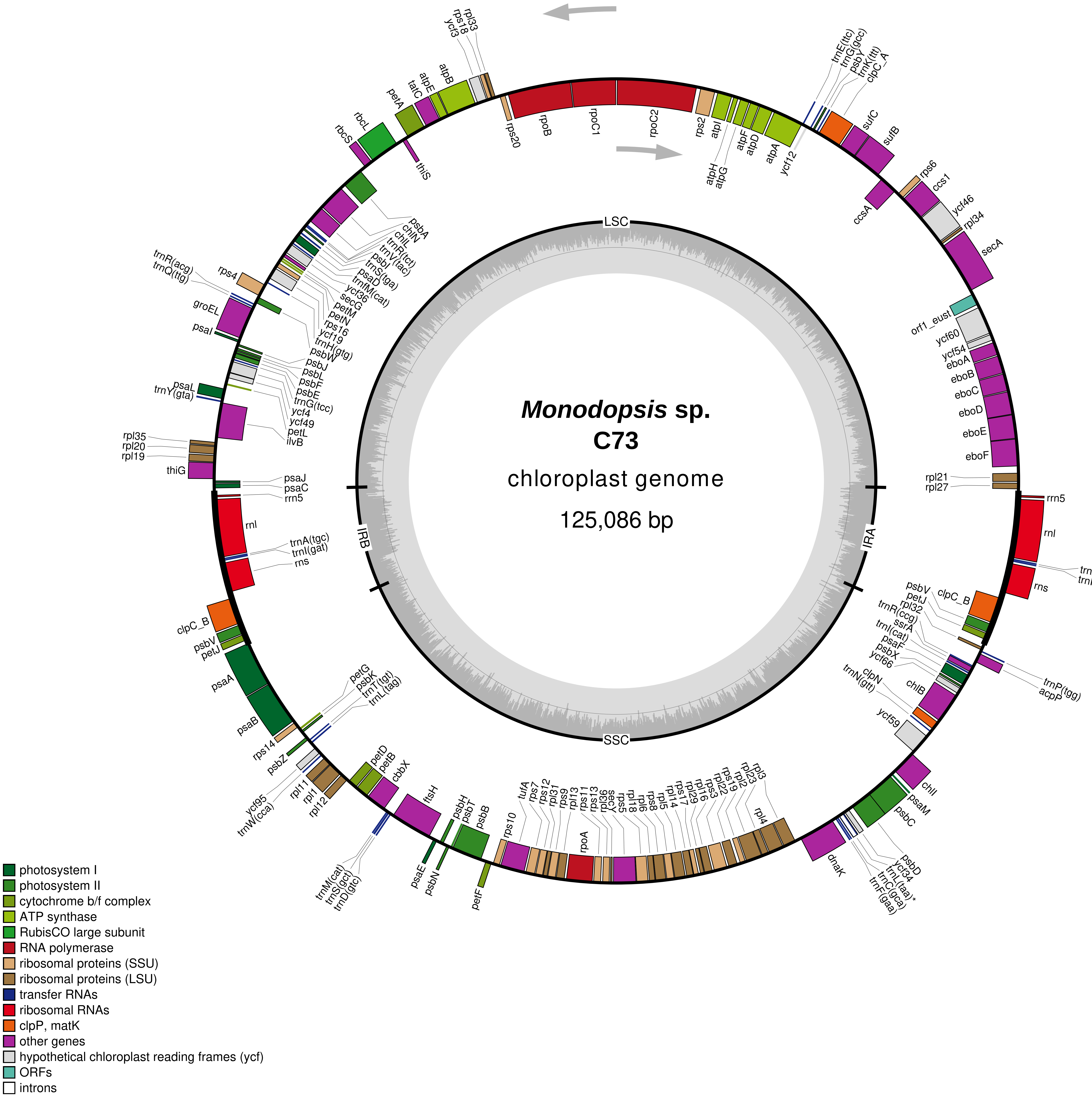
Monodopsis sp.
C73
chloroplast genome
125,086 bp
LSC
IRA
SSC
IRB
photosystem I
photosystem II
cytochrome b/f complex
ATP synthase
RubisCO large subunit
RNA polymerase
ribosomal proteins (SSU)
ribosomal proteins (LSU)
transfer RNAs
ribosomal RNAs
clpP, matK
other genes
hypothetical chloroplast reading frames (ycf)
ORFs
introns

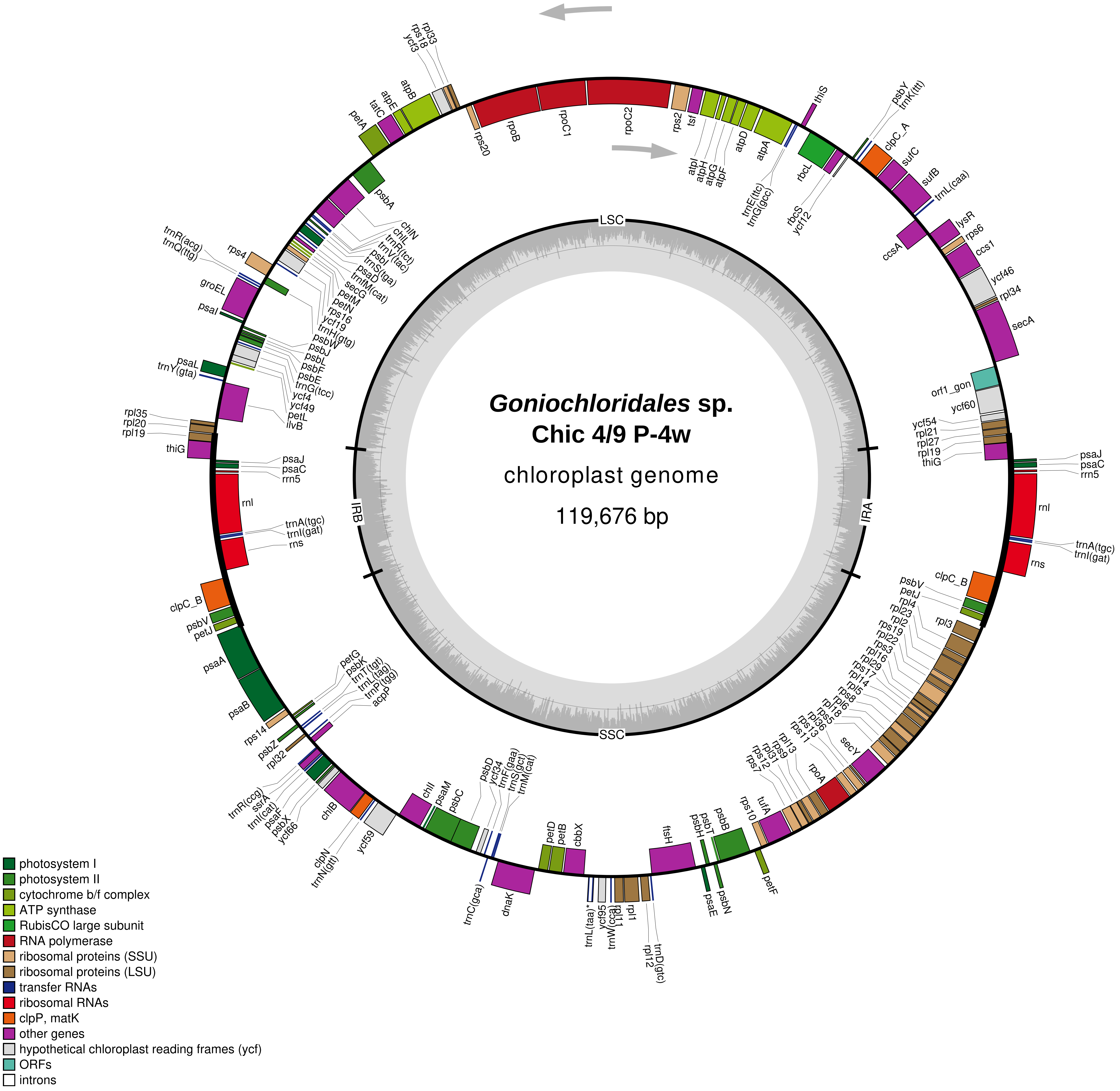
Goniochloridales sp.
Chic 4/9 P-4w
chloroplast genome
119,676 bp
LSC
IRA
SSC
IRB
rps20
rpoB
rpoC1
rpoC2
rps2
tsf
atpI
atpH
atpG
atpF
atpD
atpA
trnE(ttc)
trnG(gcc)
thiS
rbcL
rbcS
ycf12
psbY
trnK(ttt)
clpC_A
sufC
sufB
trnL(caa)
lysR
rps6
ccsA
ccs1
ycf46
rpl34
secA
orf1_gon
ycf60
ycf54
rpl21
rpl27
rpl19
thiG
psaJ
psaC
rrn5
rnl
trnA(tgc)
trnI(gat)
rns
clpC_B
psbV
petJ
rpl4
rpl23
rpl2
rps19
rpl22
rps3
rpl16
rpl29
rps17
rpl14
rpl5
rps8
rpl6
rpl18
rps5
rpl36
rps13
rps11
secY
rpoA
rpl13
rps9
rpl31
rps12
rps7
rps10
tufA
psbB
psbT
psbH
psaE
psbN
petF
ftsH
trnD(gtc)
rpl12
rpl1
rpl11
trnW(cca)
ycf95
trnL(taa)*
cbbX
petB
petD
dnaK
trnC(gca)
trnM(cat)
trnS(gct)
trnF(gaa)
ycf34
psbD
psbC
psaM
chlI
ycf59
trnN(gtt)
clpN
chlB
ycf66
psbX
psaF
trnI(cat)
ssrA
trnR(ccg)
rpl32
psbZ
rps14
acpP
trnP(tgg)
trnL(tag)
trnT(tgt)
psbK
petG
psaB
psaA
petJ
psbV
clpC_B
rns
trnI(gat)
trnA(tgc)
rnl
rrn5
psaC
psaJ
thiG
rpl19
rpl20
rpl35
ilvB
petL
ycf49
ycf4
trnG(tcc)
psbE
psbF
psbL
psbJ
psbW
trnY(gta)
psaL
psaI
groEL
trnQ(ttg)
trnR(acg)
rps4
trnH(gtg)
ycf19
rps16
petN
petM
secG
trnfM(cat)
psaD
trnS(tga)
psbI
trnV(tac)
trnR(tct)
chlL
chlN
psbA
petA
tatC
atpE
atpB
ycf3
rps18
rpl33
photosystem I
photosystem II
cytochrome b/f complex
ATP synthase
RubisCO large subunit
RNA polymerase
ribosomal proteins (SSU)
ribosomal proteins (LSU)
transfer RNAs
ribosomal RNAs
clpP, matK
other genes
hypothetical chloroplast reading frames (ycf)
ORFs
introns

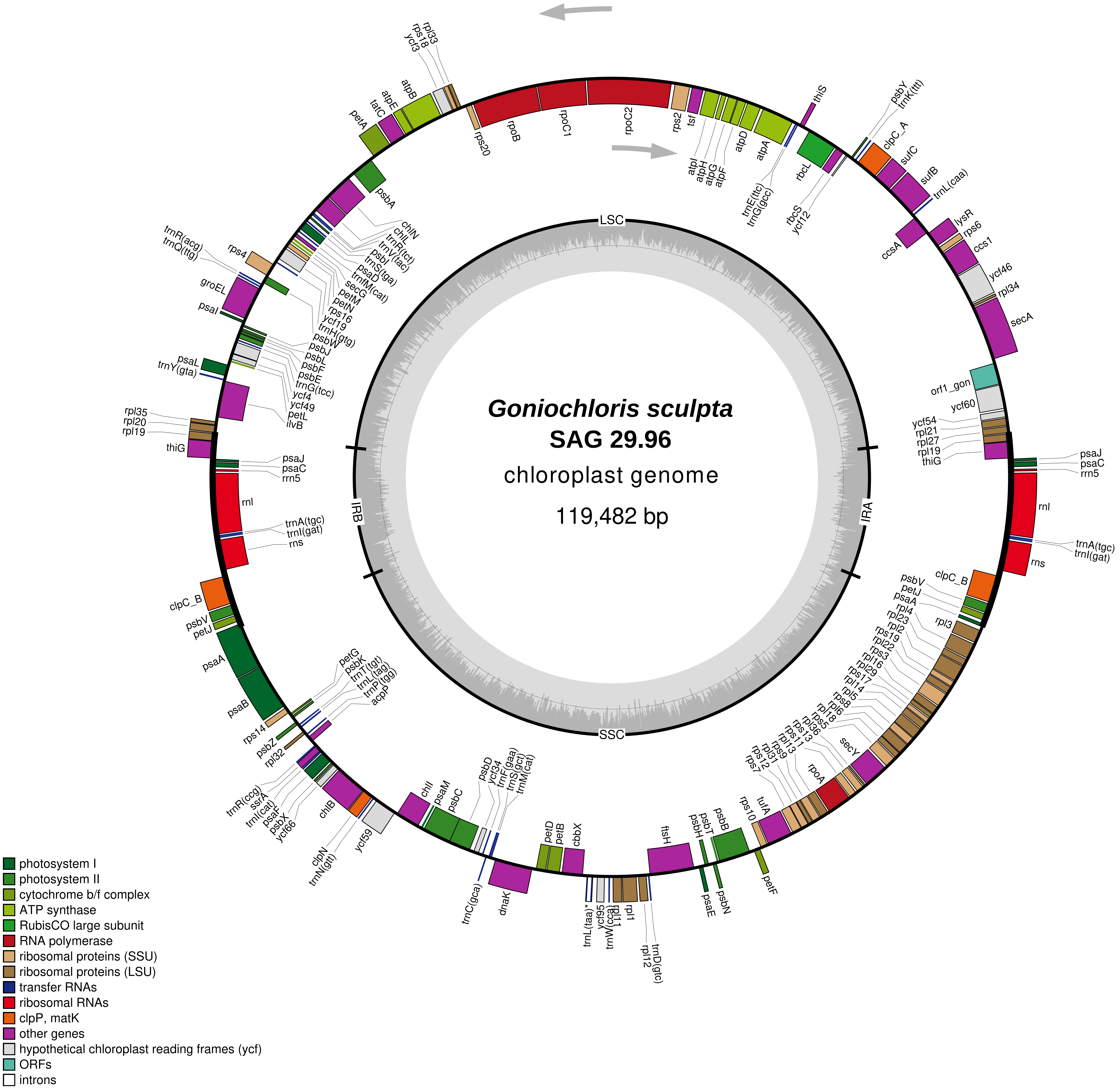
Goniochloris sculpta
SAG 29.96
chloroplast genome
119,482 bp
LSC
IRA
SSC
IRB
rps20
rpoB
rpoC1
rpoC2
rps2
tsf
atpI
atpH
atpG
atpF
atpD
atpA
trnE(ttc)
trnG(gcc)
thiS
rbcL
rbcS
ycf12
psbY
trnK(ttt)
clpC_A
sufC
sufB
trnL(caa)
ccsA
lysR
rps6
ccs1
ycf46
rpl34
secA
orf1_gon
ycf60
ycf54
rpl21
rpl27
rpl19
thiG
psaJ
psaC
rrn5
rnl
trnA(tgc)
trnI(gat)
rns
clpC_B
psbV
petJ
psaA
rpl4
rpl3
rpl23
rpl2
rps19
rpl22
rps3
rpl16
rpl29
rps17
rpl14
rpl5
rps8
rpl6
rpl18
rps5
rpl36
rps13
rps11
rpl13
rps9
rpl31
rps12
rps7
secY
rpoA
rps10
tufA
psbB
psbT
psbH
petF
psbN
psaE
ftsH
rpl1
rpl11
trnD(gtc)
rpl12
trnW(cca)
ycf95
trnL(taa)*
cbbX
petB
petD
dnaK
trnC(gca)
psbD
ycf34
trnF(gaa)
trnS(gct)
trnM(cat)
psbC
psaM
chlI
ycf59
clpN
trnN(gtt)
chlB
ycf66
psbX
psaF
trnI(cat)
ssrA
trnR(ccg)
rpl32
psbZ
rps14
psaB
petG
psbK
trnT(tgt)
trnL(tag)
trnP(tgg)
acpP
psaL
trnY(gta)
rpl35
rpl20
ilvB
petL
ycf49
ycf4
trnG(tcc)
psbE
psbF
psbL
psbJ
psbW
trnH(gtg)
ycf19
rps16
petN
petM
secG
trnfM(cat)
psaD
trnS(tga)
psbI
trnV(tac)
trnR(tct)
chlL
chlN
psbA
psaI
groEL
trnQ(ttg)
trnR(acg)
rps4
petA
tatC
atpE
atpB
ycf3
rps18
rpl33
photosystem I
photosystem II
cytochrome b/f complex
ATP synthase
RubisCO large subunit
RNA polymerase
ribosomal proteins (SSU)
ribosomal proteins (LSU)
transfer RNAs
ribosomal RNAs
clpP, matK
other genes
hypothetical chloroplast reading frames (ycf)
ORFs
introns

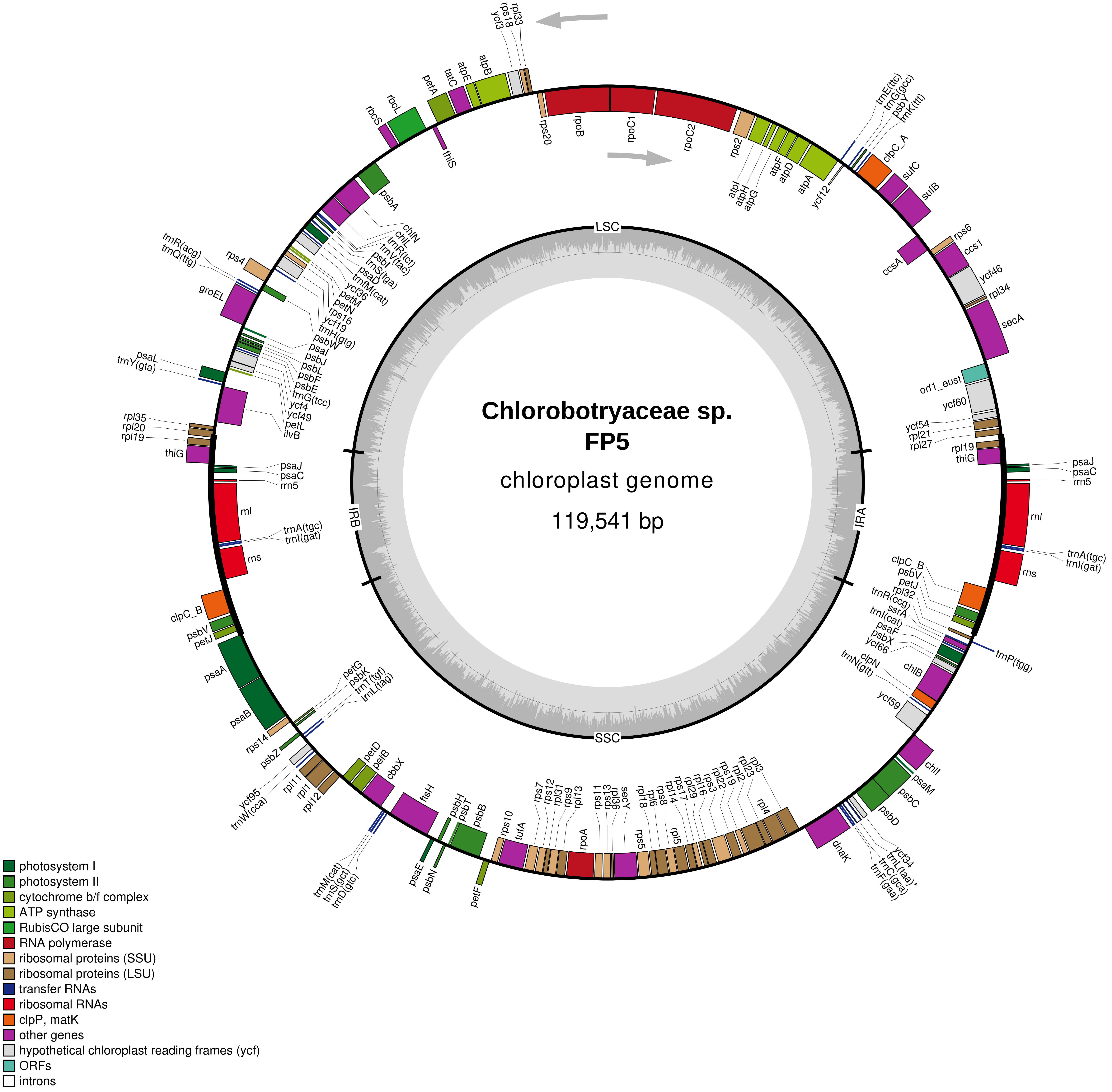

Chlorobotryaceae sp. FP5
chloroplast genome
119,541 bp
LSC
IRA
SSC
IRB
rpl33
rps18
ycf3
atpB
atpE
tatC
petA
rbcL
rbcS
thiS
rps20
rpoB
rpoC1
rpoC2
rps2
atpI
atpH
atpG
atpF
atpD
atpA
ycf12
trnE(ttc)
trnG(gcc)
psbY
trnK(ttt)
clpC_A
sufC
sufB
rps6
ccs1
ccsA
ycf46
rpl34
secA
orf1_eust
ycf60
ycf54
rpl21
rpl27
rpl19
thiG
psaJ
psaC
rrn5
rnl
trnA(tgc)
trnI(gat)
rns
clpC_B
psbV
petJ
rpl32
trnR(ccg)
ssrA
trnI(cat)
psaF
psbX
ycf66
chlB
clpN
trnN(gtt)
ycf59
trnP(tgg)
chlI
psaM
psbC
psbD
ycf34
trnL(taa)*
trnC(gca)
trnF(gaa)
dnaK
rpl3
rpl23
rpl2
rps19
rpl22
rps3
rpl16
rpl29
rps17
rpl14
rpl5
rps8
rpl6
rpl18
rps5
secY
rpl36
rps13
rps11
rpoA
rpl13
rps9
rpl31
rps12
rps7
tufA
rps10
psbB
psbT
psbH
psbN
psaE
petF
ftsH
cbbX
petB
petD
trnM(cat)
trnS(gct)
trnD(gtc)
rpl12
rpl1
rpl11
ycf95
trnW(cca)
psbZ
rps14
psaB
psaA
petG
psbK
trnT(tgt)
trnL(tag)
psbA
chlN
chlL
trnR(tct)
trnV(tac)
psbI
trnS(tga)
psaD
trnfM(cat)
ycf36
petM
petN
rps16
ycf19
trnH(gtg)
psbW
psaI
psbJ
psbL
psbF
psbE
trnG(tcc)
ycf4
ycf49
petL
ilvB
trnR(acg)
trnQ(ttg)
rps4
groEL
psaL
trnY(gta)
rpl35
rpl20
photosystem I
photosystem II
cytochrome b/f complex
ATP synthase
RubisCO large subunit
RNA polymerase
ribosomal proteins (SSU)
ribosomal proteins (LSU)
transfer RNAs
ribosomal RNAs
clpP, matK
other genes
hypothetical chloroplast reading frames (ycf)
ORFs
introns

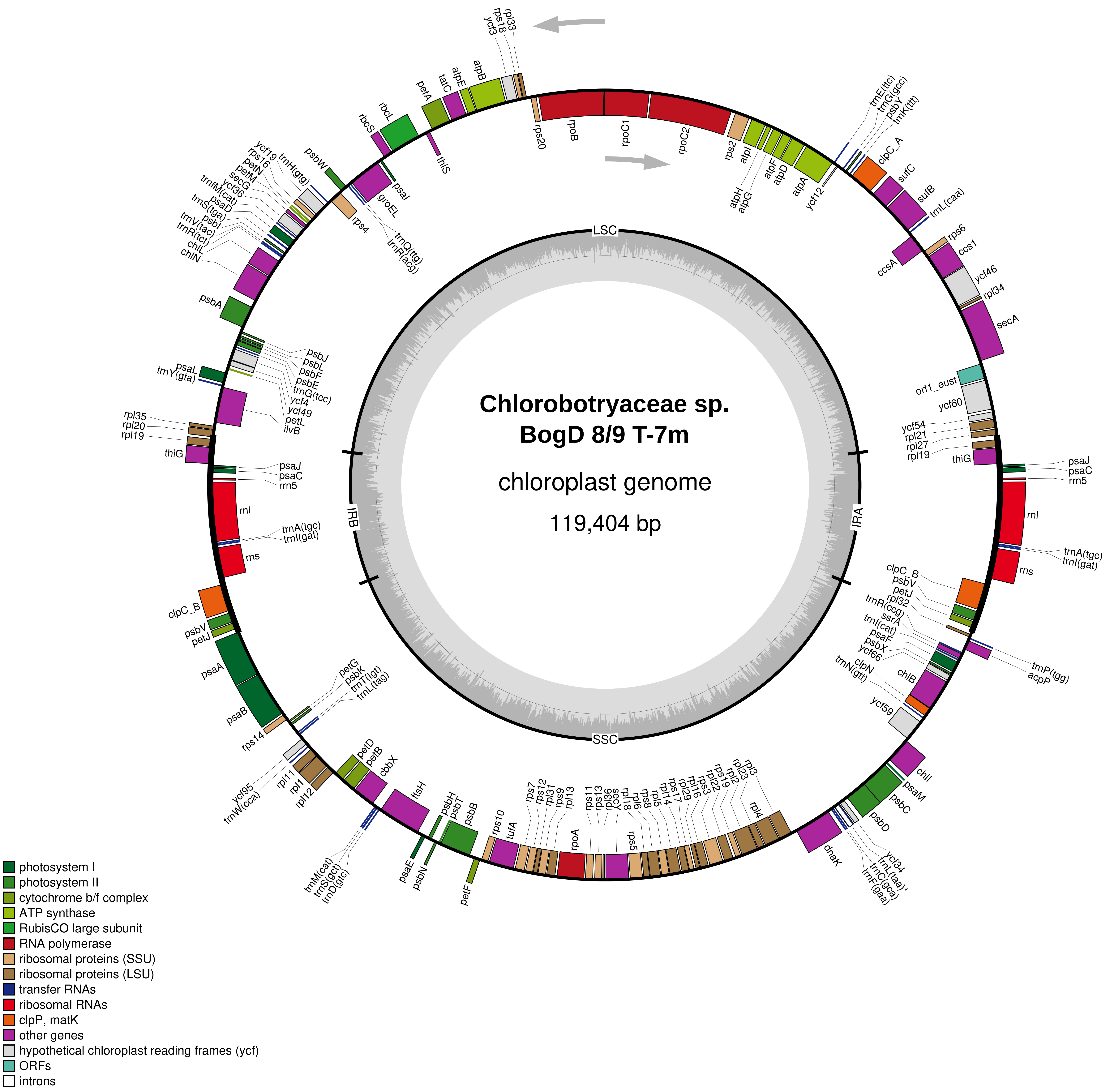
Chlorobotryaceae sp.
BogD 8/9 T-7m
chloroplast genome
119,404 bp
LSC
IRA
SSC
IRB
rpl33
rps18
ycf3
atpB
atpE
tatC
petA
rbcL
rbcS
thiS
psbW
psaI
groEL
trnQ(ttg)
trnR(acg)
rps4
ycf19
rps16
trnH(gtg)
petN
petM
secG
ycf36
trnM(cat)
psaD
trnS(tga)
psbI
trnV(tac)
trnR(tct)
chlL
chlN
psbA
psbJ
psbL
psbF
psbE
trnG(tcc)
ycf4
ycf49
petL
ilvB
psaL
trnY(gta)
rpl35
rpl20
rpl19
thiG
psaJ
psaC
rrn5
rnl
trnA(tgc)
trnI(gat)
rns
clpC_B
psbV
petJ
psaA
psaB
rps14
petG
psbK
trnT(tgt)
trnL(tag)
ycf95
trnW(cca)
rpl11
rpl1
rpl12
petD
petB
cbbX
ftsH
trnM(cat)
trnS(gct)
trnD(gtc)
psaE
psbN
psbH
psbT
psbB
petF
rps10
tufA
rps7
rps12
rpl31
rps9
rpl13
rpoA
rps11
rps13
rpl36
secY
rpl18
rpl6
rps8
rpl5
rpl14
rps17
rpl29
rpl16
rps3
rpl22
rps19
rpl2
rpl23
rpl3
rpl4
dnaK
ycf34
trnL(taa)*
trnC(gca)
trnF(gaa)
psbD
psbC
psaM
chlI
ycf59
trnN(gtt)
clpN
chlB
ycf66
psbX
psaF
trnI(cat)
ssrA
trnR(ccg)
rpl32
petJ
psbV
clpC_B
trnP(tgg)
acpP
rns
trnA(tgc)
trnI(gat)
rnl
psaJ
psaC
rrn5
thiG
rpl19
rpl27
rpl21
ycf54
ycf60
orf1_eust
secA
rpl34
ycf46
ccs1
rps6
ccsA
trnL(caa)
sufB
sufC
clpC_A
trnK(ttt)
psbY
trnG(gcc)
trnE(ttc)
ycf12
atpA
atpD
atpF
atpG
atpH
atpI
rps2
rpoC2
rpoC1
rpoB
rps20
photosystem I
photosystem II
cytochrome b/f complex
ATP synthase
RubisCO large subunit
RNA polymerase
ribosomal proteins (SSU)
ribosomal proteins (LSU)
transfer RNAs
ribosomal RNAs
clpP, matK
other genes
hypothetical chloroplast reading frames (ycf)
ORFs
introns

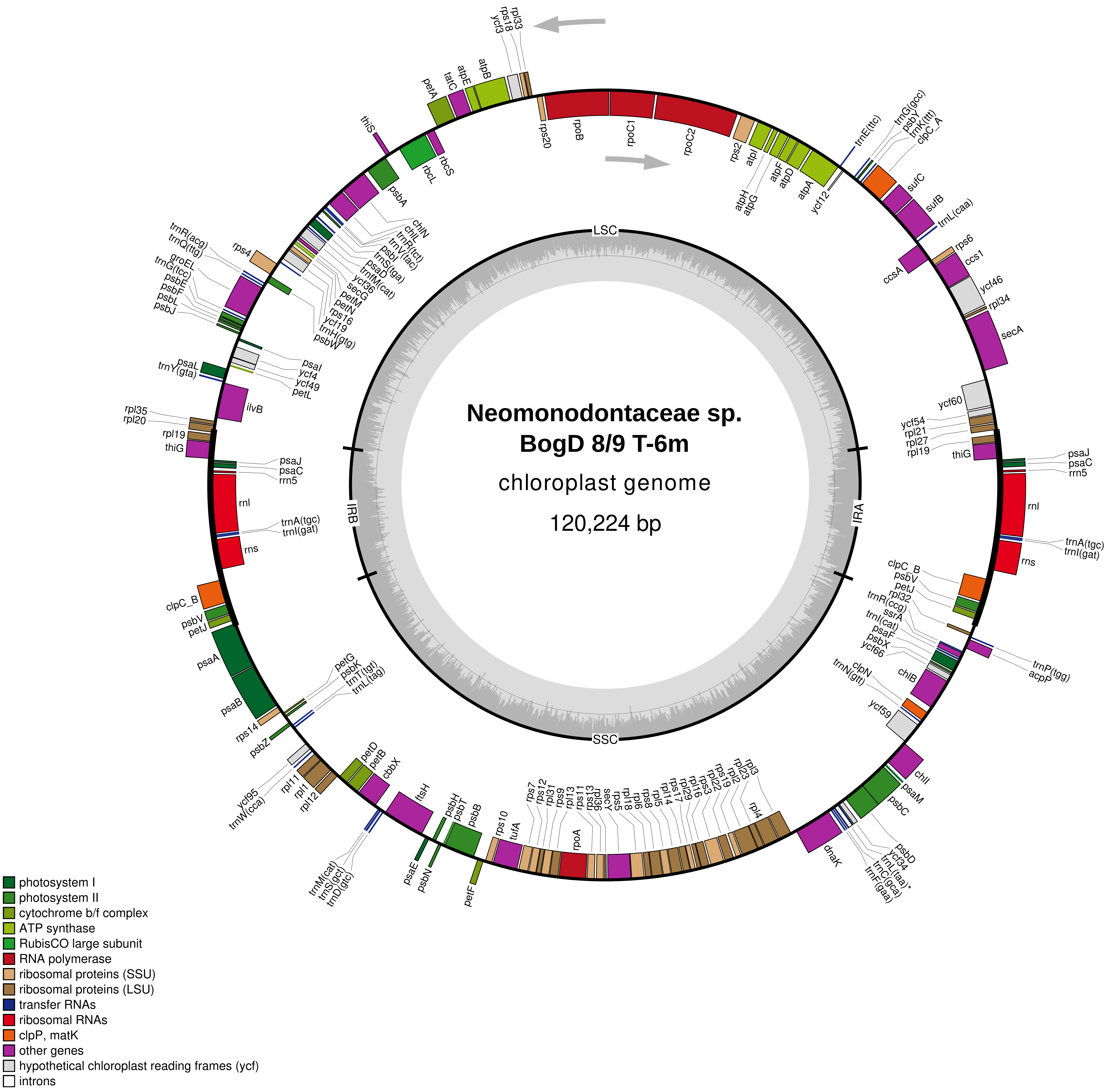

Neomonodontaceae sp.
BogD 8/9 T-6m
chloroplast genome
120,224 bp
LSC
IRA
SSC
IRB
rps20
rpoB
rpoC1
rpoC2
rps2
atpI
atpH
atpG
atpF
atpD
atpA
ycf12
trnE(ttc)
trnG(gcc)
psbY
trnK(ttt)
clpC_A
sufC
sufB
trnL(caa)
rps6
ccs1
ccsA
ycf46
rpl34
secA
ycf60
ycf54
rpl21
rpl27
rpl19
thiG
psaJ
psaC
rrn5
rnl
trnA(tgc)
trnI(gat)
rns
clpC_B
psbV
petJ
rpl32
trnR(ccg)
ssrA
trnI(cat)
psaF
psbX
ycf66
trnP(tgg)
acpP
chlB
clpN
trnN(gtt)
ycf59
chlI
psaM
psbC
psbD
ycf34
trnL(taa)*
trnC(gca)
trnF(gaa)
dnaK
rpl3
rpl23
rpl2
rps19
rpl22
rps3
rpl16
rpl29
rps17
rpl14
rpl5
rps8
rpl6
rpl18
rps5
secY
rpl36
rps13
rps11
rpoA
rpl13
rps9
rpl31
rps12
rps7
tufA
rps10
psbB
psbT
psbH
petF
psbN
psaE
ftsH
cbbX
petB
petD
trnM(cat)
trnS(gct)
trnD(gtc)
rpl12
rpl1
rpl11
ycf95
trnW(cca)
psbZ
rps14
petG
psbK
trnT(tgt)
trnL(tag)
psaB
psaA
ilvB
rpl35
rpl20
psaL
trnY(gta)
psaI
ycf4
ycf49
petL
trnR(acg)
trnQ(ttg)
groEL
trnG(tcc)
psbE
psbF
psbL
psbJ
rps4
psbW
trnH(gtg)
ycf19
rps16
petN
secG
ycf36
trnfM(cat)
psaD
trnS(tga)
psbI
trnV(tac)
trnP(tct)
chlL
chlN
psbA
thiS
rbcL
rbcS
petA
tatC
atpE
atpB
ycf3
rps18
rpl33
photosystem I
photosystem II
cytochrome b/f complex
ATP synthase
RubisCO large subunit
RNA polymerase
ribosomal proteins (SSU)
ribosomal proteins (LSU)
transfer RNAs
ribosomal RNAs
clpP, matK
other genes
hypothetical chloroplast reading frames (ycf)
introns

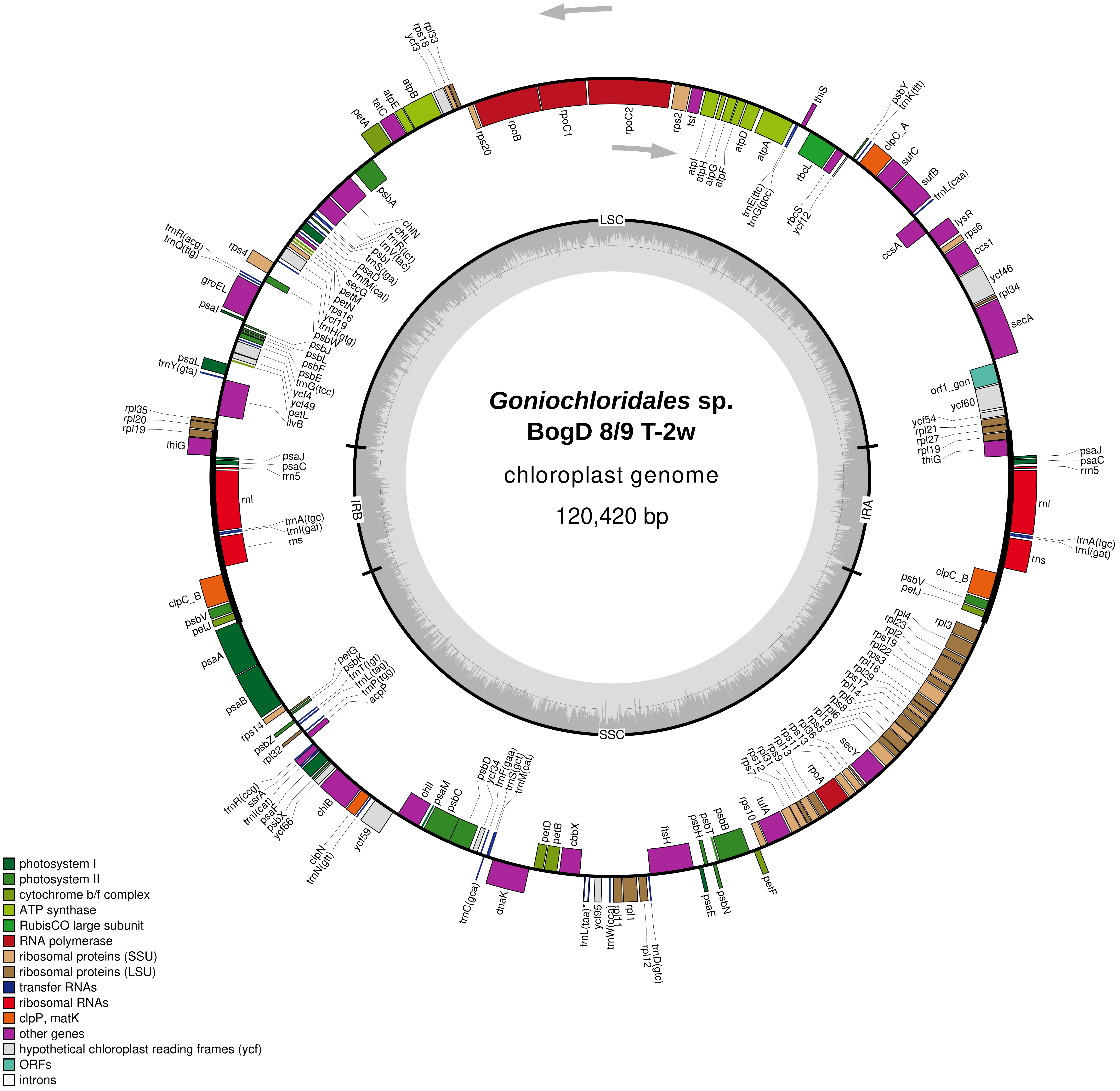

Goniochloridales sp.
BogD 8/9 T-2w
chloroplast genome
120,420 bp
LSC
IRA
SSC
IRB
rps20
rpoB
rpoC1
rpoC2
rps2
tsf
atpI
atpH
atpG
atpF
atpD
atpA
trnE(ttc)
trnG(gcc)
thiS
rbcL
rbcS
ycf12
psbY
trnK(ttt)
clpC_A
sufC
sufB
trnL(caa)
ccsA
lysR
rps6
ccs1
ycf46
rpl34
secA
orf1_gon
ycf60
ycf54
rpl21
rpl27
rpl19
thiG
psaJ
psaC
rrn5
rnl
trnA(tgc)
trnI(gat)
rns
clpC_B
psbV
petJ
rpl3
rpl4
rpl23
rpl2
rps19
rpl22
rps3
rpl16
rpl29
rps17
rpl14
rpl5
rps8
rpl6
rpl18
rps5
secY
rpl36
rps13
rps11
rpoA
rps9
rpl13
rpl31
rps12
rps7
tufA
rps10
psbB
psbT
psbH
petF
psbN
psaE
ftsH
trnD(gtc)
rpl12
rpl1
rpl11
trnW(cca)
ycf95
trnL(taa)*
cbbX
petB
petD
dnaK
trnC(gca)
trnM(cat)
trnS(gct)
trnF(gaa)
ycf34
psbD
psbC
psaM
chlI
ycf59
trnN(gtt)
clpN
chlB
ycf66
psbX
psaF
trnI(cat)
ssrA
trnR(ccg)
rpl32
psbZ
rps14
psaB
psaA
petG
psbK
trnT(tgt)
trnL(tag)
trnP(tgg)
acpP
petJ
psbV
clpC_B
rns
trnI(gat)
trnA(tgc)
rnl
rrn5
psaC
psaJ
thiG
rpl19
rpl20
rpl35
ilvB
petL
ycf49
ycf4
trnG(tcc)
psbE
psbF
psbL
psbJ
psbW
trnH(gtg)
ycf19
rps16
petN
petM
secG
trnfM(cat)
psaD
trnS(tga)
psbI
trnV(tac)
trnR(tct)
chlL
chlN
psbA
psaL
trnY(gta)
psaI
groEL
trnQ(ttg)
trnR(acg)
rps4
petA
tatC
atpE
atpB
ycf3
rps18
rpl33
photosystem I
photosystem II
cytochrome b/f complex
ATP synthase
RubisCO large subunit
RNA polymerase
ribosomal proteins (SSU)
ribosomal proteins (LSU)
transfer RNAs
ribosomal RNAs
clpP, matK
other genes
hypothetical chloroplast reading frames (ycf)
ORFs
introns
