## Supplementary Dataset S2 for "Insights into the phylogeny and enigmatic mitochondrial biology of eustigmatophyte algae from over 50 newly sequenced organellar genomes"

**Supplementary Dataset S2.** Gene maps of the newly generated eustigmatophyte mitochondrial genomes. The display convention is the same as in Dataset. S1.

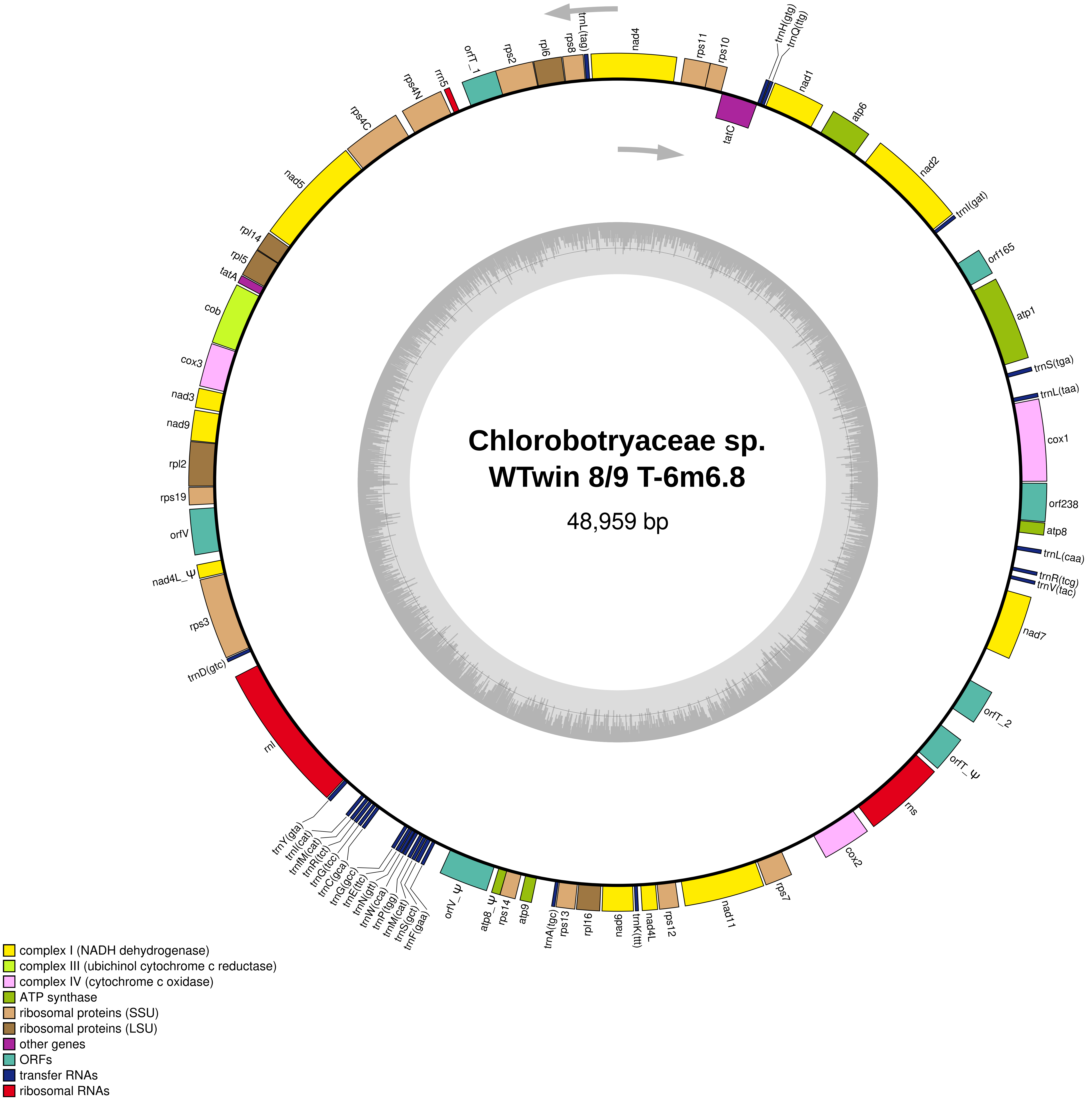
Chlorobotryaceae sp.
WTwin 8/9 T-6m6.8
48,959 bp
nad4
rps11
rps10
trnH(gtg)
trnQ(ttg)
nad1
atp6
nad2
trnI(gat)
orf165
atp1
trnS(tga)
trnL(taa)
cox1
orf238
atp8
trnL(caa)
trnR(tcg)
trnV(tac)
nad7
orfT_2
orfT_Ψ
rns
cox2
rps7
nad11
rps12
nad4L
trnK(ttt)
nad6
rpl16
rps13
trnA(tgc)
atp9
rps14
atp8_Ψ
orfV_Ψ
trnF(gaa)
trnS(gct)
trnM(cat)
trnP(tgg)
trnW(cca)
trnN(gtt)
trnE(ttc)
trnG(gcc)
trnC(gca)
trnG(tcc)
trnR(tct)
trnfM(cat)
trnI(cat)
trnY(gta)
rnl
trnD(gtc)
rps3
nad4L_Ψ
orfV
rps19
rpl2
nad9
nad3
cox3
cob
tatA
rpl5
rpl14
nad5
rps4C
rps4N
rrn5
orfT_1
rps2
rpl6
rps8
trnL(tag)
tatC
complex I (NADH dehydrogenase)
complex III (ubichinol cytochrome c reductase)
complex IV (cytochrome c oxidase)
ATP synthase
ribosomal proteins (SSU)
ribosomal proteins (LSU)
other genes
ORFs
transfer RNAs
ribosomal RNAs

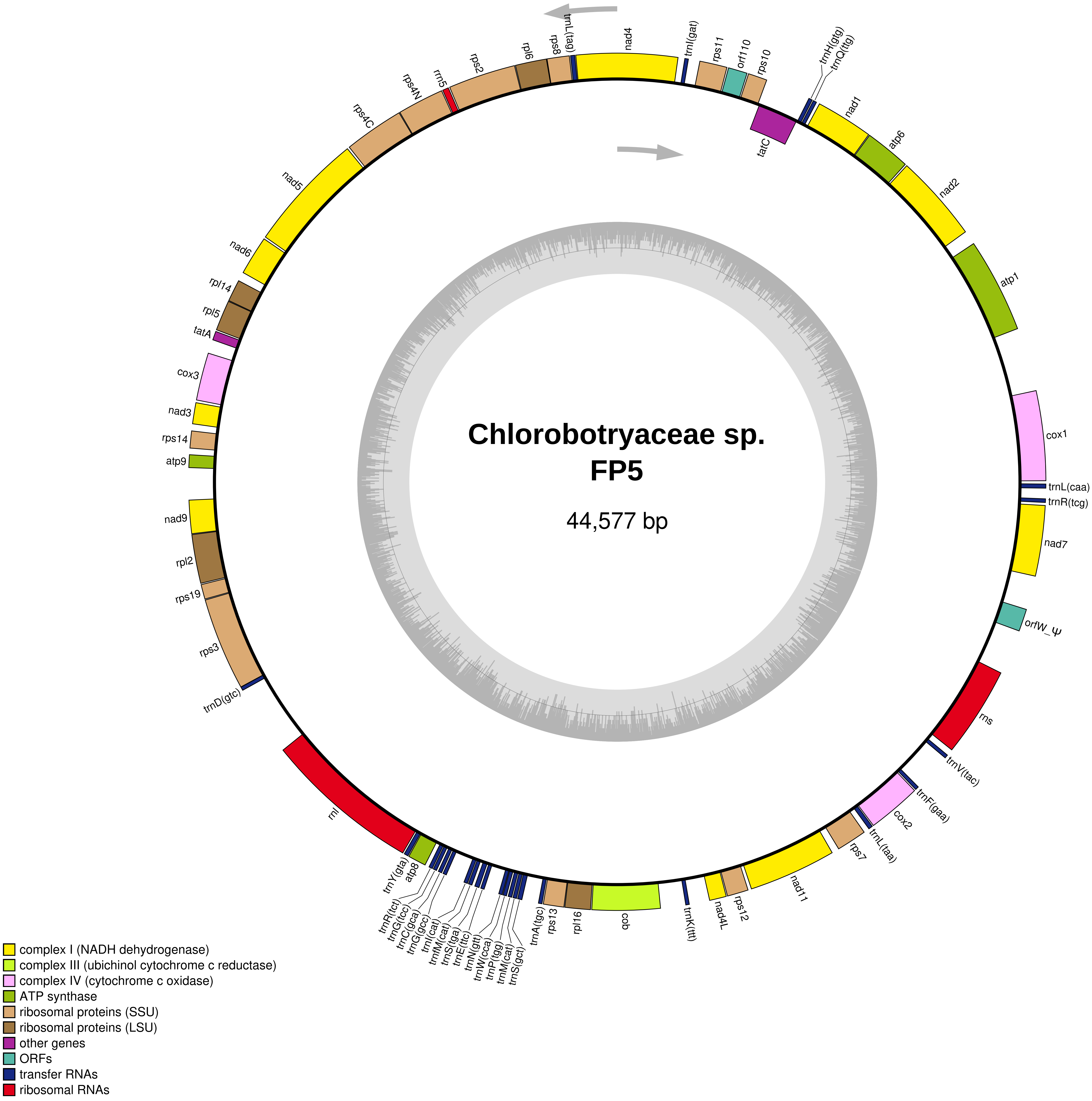
Chlorobotryaceae sp.
FP5
44,577 bp
nad4
trnL(tag)
rps8
rpl6
rps2
rrn5
rps4N
rps4C
nad5
nad6
rpl14
rpl5
tatA
cox3
nad3
rps14
atp9
nad9
rpl2
rps19
rps3
trnD(gtc)
rnl
trnY(gta)
atp8
trnR(tct)
trnG(tcc)
trnC(gca)
trnG(gcc)
trnI(cat)
trnfM(cat)
trnS(tga)
trnE(ttc)
trnN(gtt)
trnW(cca)
trnP(tgg)
trnM(cat)
trnS(gct)
trnA(tgc)
rps13
rpl16
cob
trnK(ttt)
nad4L
rps12
nad11
rps7
trnL(taa)
cox2
trnF(gaa)
trnV(tac)
rns
orfW_Ψ
nad7
trnR(tcg)
trnL(caa)
cox1
atp1
nad2
atp6
nad1
tatC
trnH(gtg)
trnQ(ttg)
rps10
orf110
rps11
trnI(gat)
complex I (NADH dehydrogenase)
complex III (ubichinol cytochrome c reductase)
complex IV (cytochrome c oxidase)
ATP synthase
ribosomal proteins (SSU)
ribosomal proteins (LSU)
other genes
ORFs
transfer RNAs
ribosomal RNAs

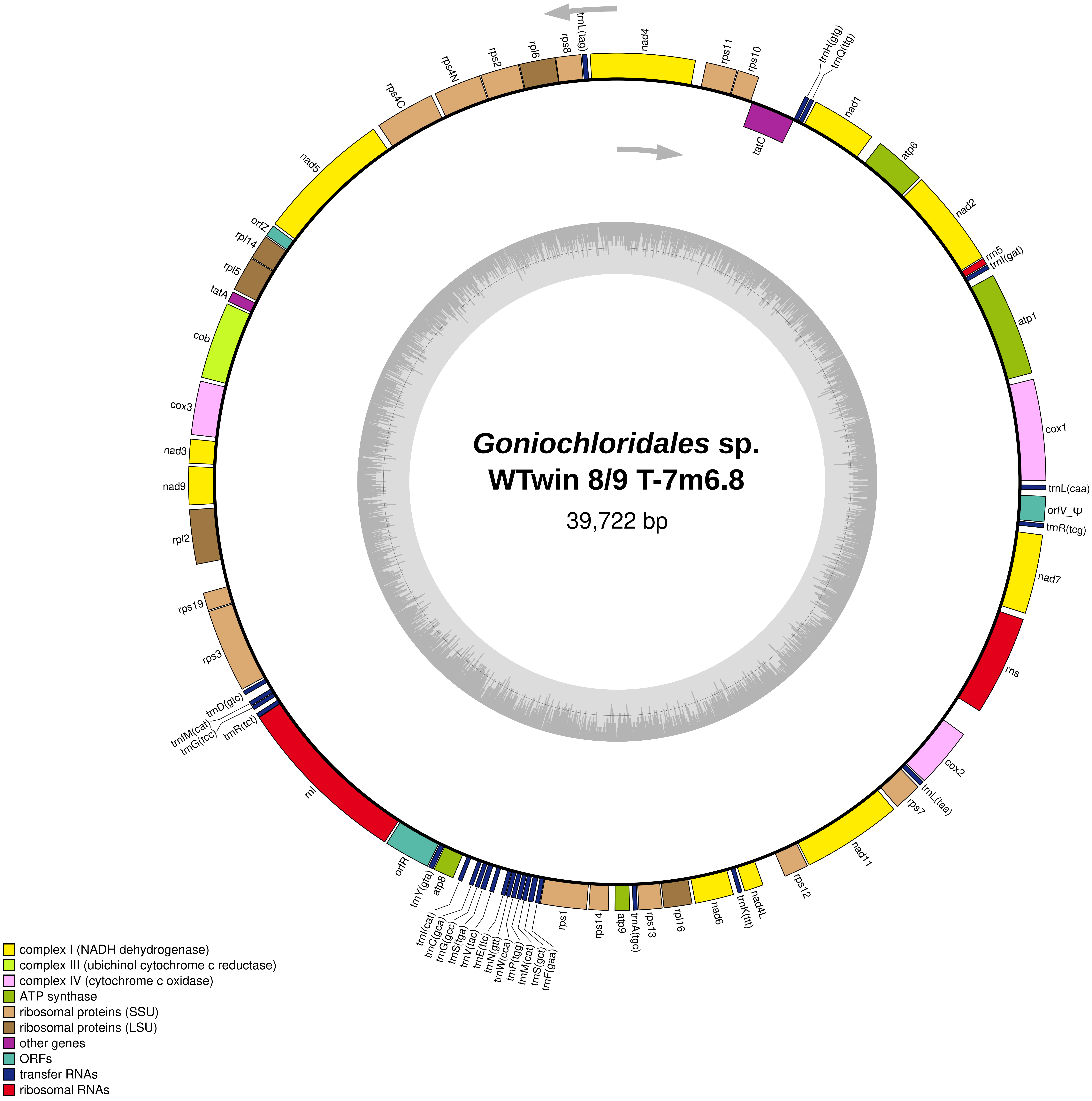
Goniochloridales sp.
WTwin 8/9 T-7m6.8
39,722 bp
trnL(tag)
nad4
rps11
rps10
trnH(gtg)
trnQ(ttg)
nad1
tatC
atp6
nad2
rrn5
trnI(gat)
atp1
cox1
trnL(caa)
orfV_Ψ
trnR(tcg)
nad7
rns
cox2
trnL(taa)
rps7
nad11
rps12
nad4L
trnK(ttt)
nad6
rpl16
rps13
trnA(tgc)
atp9
rps14
rps1
trnF(gaa)
trnS(gct)
trnM(cat)
trnP(tgg)
trnW(cca)
trnN(gtt)
trnE(ttc)
trnV(tac)
trnS(tga)
trnG(gcc)
trnC(gca)
trnI(cat)
atp8
trnY(gta)
orfR
rnl
trnR(tct)
trnG(tcc)
trnfM(cat)
trnD(gtc)
rps3
rps19
rpl2
nad9
nad3
cox3
cob
tatA
rpl5
rpl14
orfZ
nad5
rps4C
rps4N
rps2
rpl6
rps8
complex I (NADH dehydrogenase)
complex III (ubichinol cytochrome c reductase)
complex IV (cytochrome c oxidase)
ATP synthase
ribosomal proteins (SSU)
ribosomal proteins (LSU)
other genes
ORFs
transfer RNAs
ribosomal RNAs

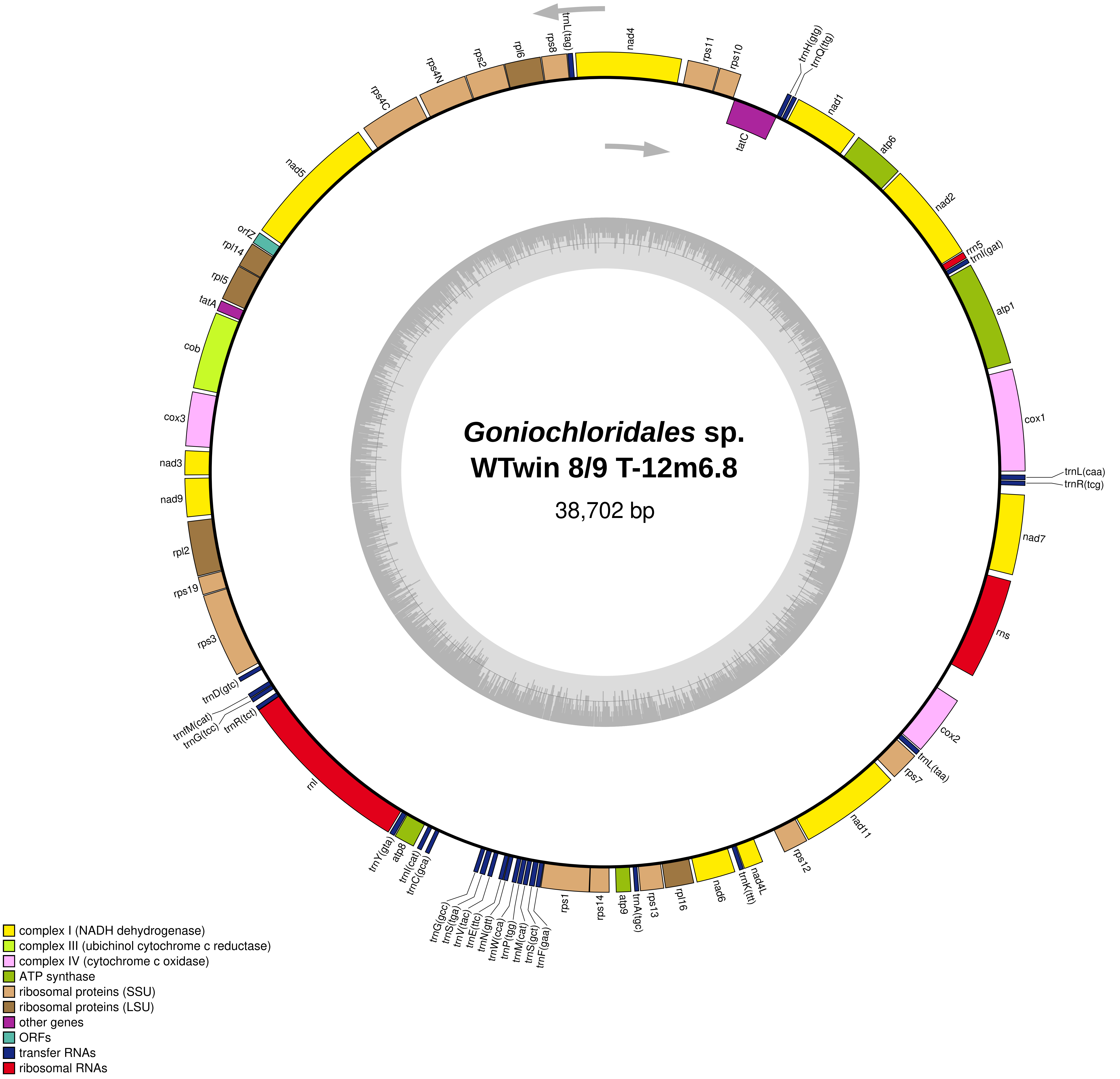
*Goniochloridales* sp.
WTwin 8/9 T-12m6.8
38,702 bp
nad4
rps11
rps10
trnH(gtg)
trnQ(ttg)
tatC
nad1
atp6
nad2
rrn5
trnI(gat)
atp1
cox1
trnL(caa)
trnR(tcg)
nad7
rns
cox2
trnL(taa)
rps7
nad11
rps12
nad4L
trnK(ttt)
nad6
rpl16
rps13
trnA(tgc)
atp9
rps14
rps1
trnG(gcc)
trnS(tga)
trnV(tac)
trnE(ttc)
trnN(gtt)
trnW(cca)
trnP(tgg)
trnM(cat)
trnS(gct)
trnF(gaa)
trnC(gca)
trnI(cat)
atp8
trnY(gta)
rnl
trnR(tct)
trnG(tcc)
trnfM(cat)
trnD(gtc)
rps3
rps19
rpl2
nad9
nad3
cox3
cob
tatA
rpl5
rpl14
orfZ
nad5
rps4C
rps4N
rps2
rpl6
rps8
trnL(tag)
complex I (NADH dehydrogenase)
complex III (ubichinol cytochrome c reductase)
complex IV (cytochrome c oxidase)
ATP synthase
ribosomal proteins (SSU)
ribosomal proteins (LSU)
other genes
ORFs
transfer RNAs
ribosomal RNAs

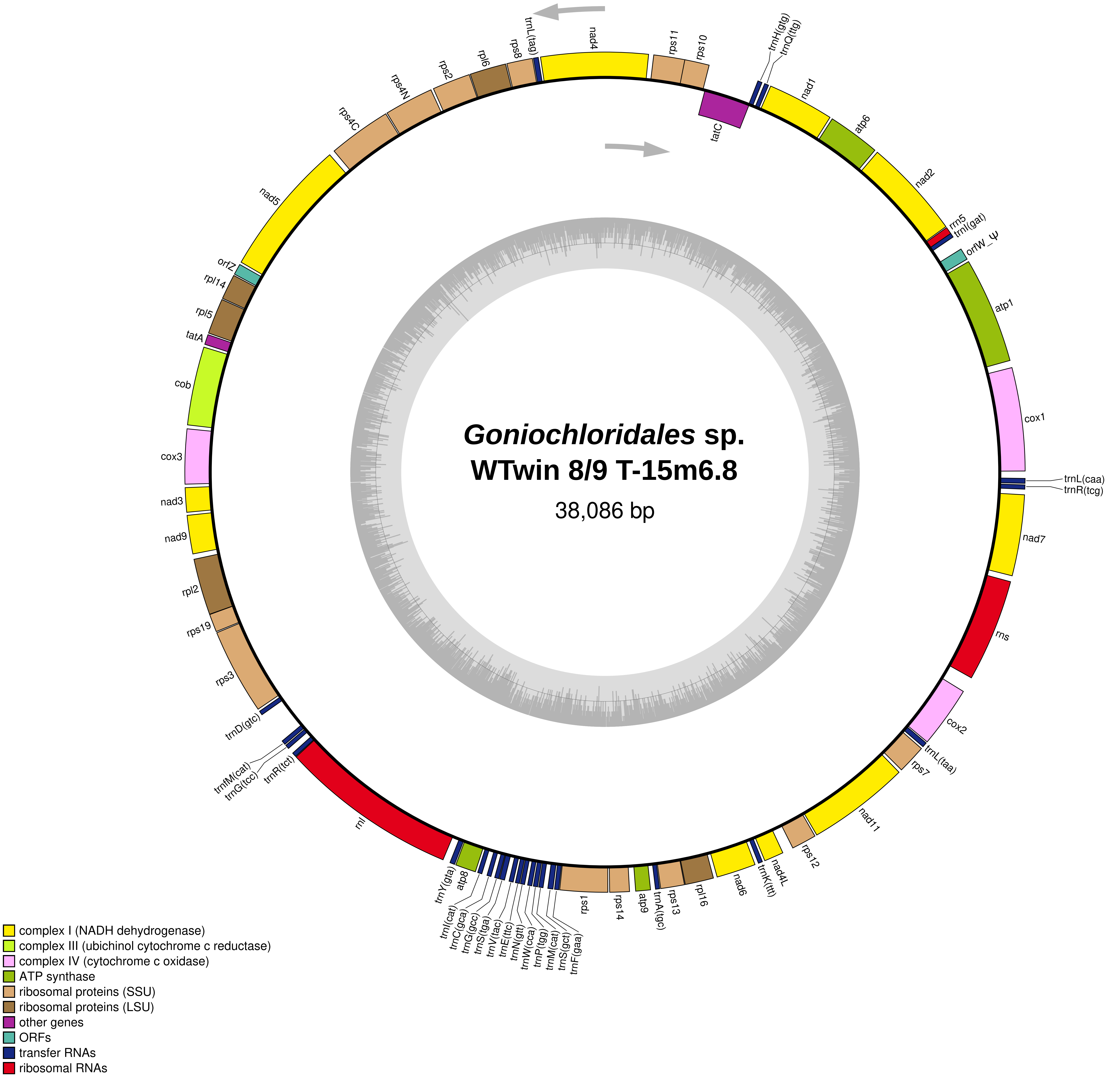
Goniochloridales sp.
WTwin 8/9 T-15m6.8
38,086 bp
complex I (NADH dehydrogenase)
complex III (ubichinol cytochrome c reductase)
complex IV (cytochrome c oxidase)
ATP synthase
ribosomal proteins (SSU)
ribosomal proteins (LSU)
other genes
ORFs
transfer RNAs
ribosomal RNAs

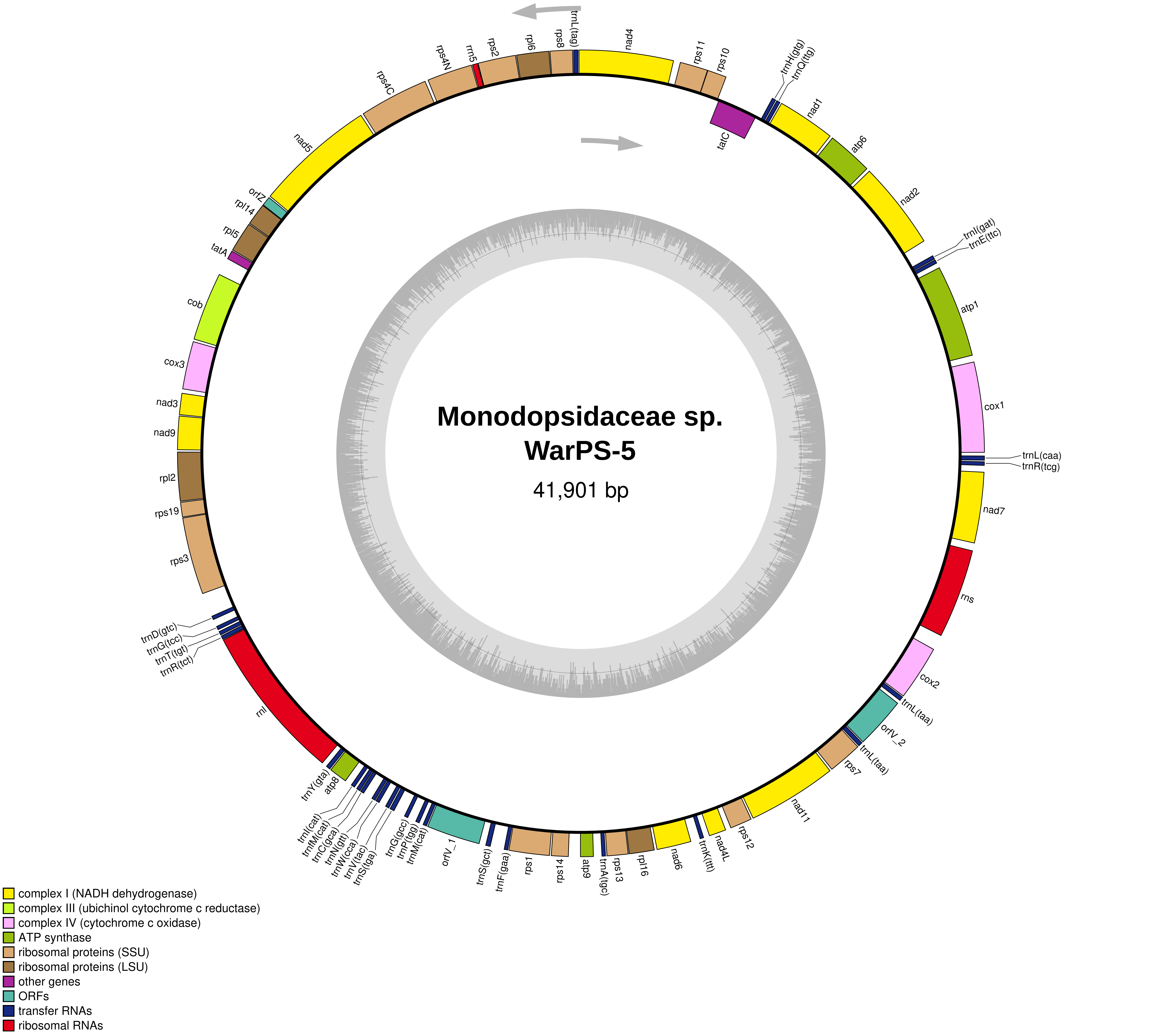
Monodopsidaceae sp.
WarPS-5
41,901 bp
trnL(tag)
nad4
rps11
rps10
trnH(gtg)
trnQ(ttg)
tatC
nad1
atp6
nad2
trnI(gat)
trnE(ttc)
atp1
cox1
trnL(caa)
trnR(tcg)
nad7
rns
cox2
trnL(taa)
orfV_2
trnL(taa)
rps7
nad11
rps12
nad4L
trnK(ttt)
nad6
rpl16
rps13
trnA(tgc)
atp9
rps14
rps1
trnF(gaa)
trnS(gct)
orfV_1
trnM(cat)
trnP(tgg)
trnG(gcc)
trnS(tga)
trnV(tac)
trnW(cca)
trnN(gtt)
trnC(gca)
trnM(cat)
trnI(cat)
atp8
trnY(gta)
rnl
trnR(tct)
trnT(tgt)
trnG(tcc)
trnD(gtc)
rps3
rps19
rpl2
nad9
nad3
cox3
cob
tatA
rpl5
rpl14
orfZ
nad5
rps4C
rps4N
rrn5
rps2
rpl6
rps8
complex I (NADH dehydrogenase)
complex III (ubichinol cytochrome c reductase)
complex IV (cytochrome c oxidase)
ATP synthase
ribosomal proteins (SSU)
ribosomal proteins (LSU)
other genes
ORFs
transfer RNAs
ribosomal RNAs

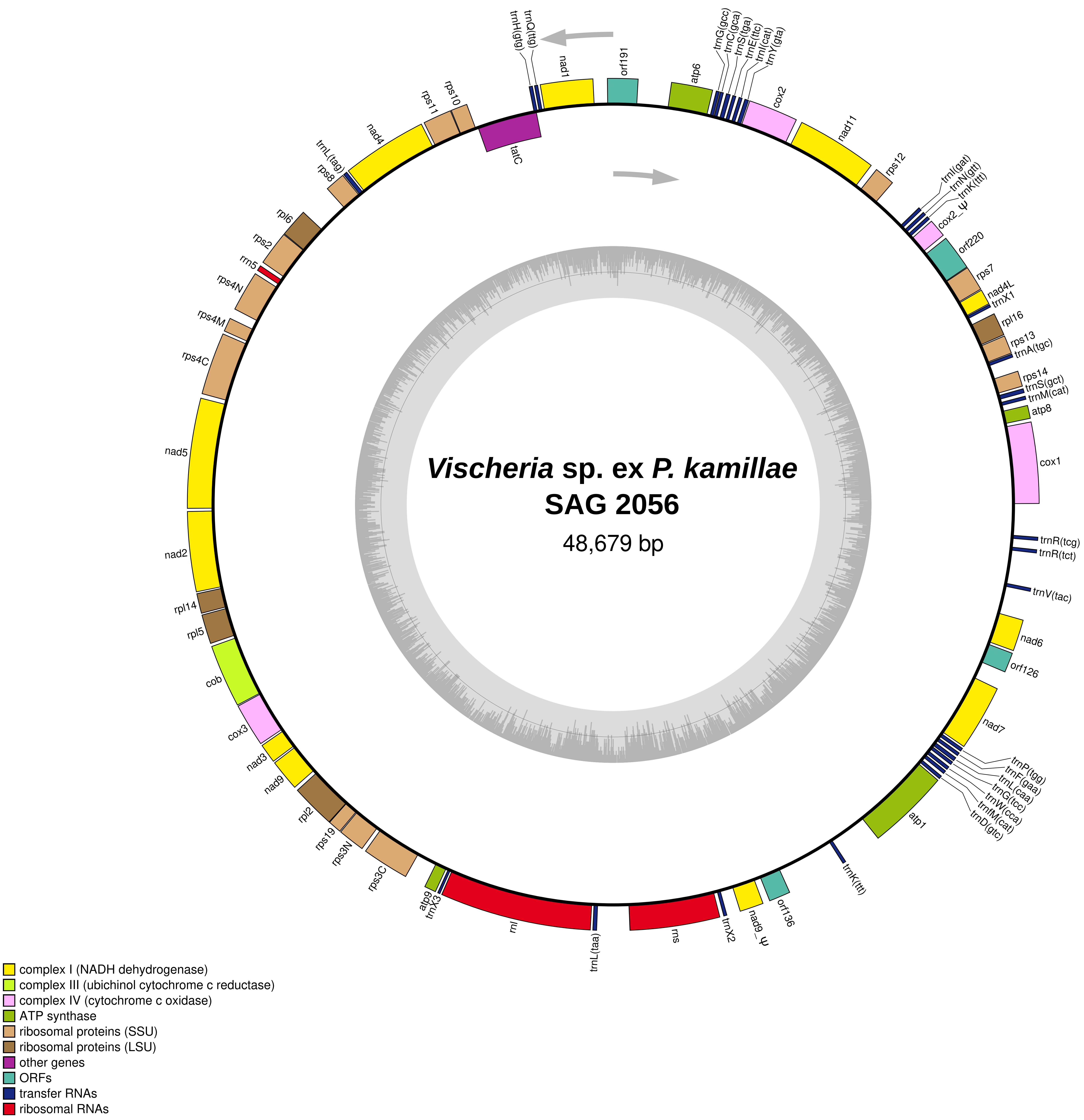

*Vischeria* sp. ex *P. kamillae*
SAG 2056
48,679 bp
trnH(gtg)
trnQ(ttg)
nad1
orf191
atp6
trnG(gcc)
trnC(gca)
trnS(tga)
trnE(ttc)
trnI(cat)
trnY(gta)
cox2
nad11
rps12
trnI(gat)
trnN(gtt)
trnK(ttt)
cox2_ψ
orf220
rps7
nad4L
trnX1
rpl16
rps13
trnA(tgc)
rps14
trnS(gct)
trnM(cat)
atp8
cox1
trnR(tcg)
trnR(tct)
trnV(tac)
nad6
orf126
nad7
trnP(tgg)
trnF(gaa)
trnL(caa)
trnG(tcc)
trnW(cca)
trnM(cat)
trnD(gtc)
atp1
trnK(ttt)
orf136
nad9_ψ
trnX2
rns
trnL(taa)
rnl
trnX3
atp9
rps3C
rps3N
rps19
rpl2
nad9
nad3
cox3
cob
rpl5
rpl14
nad2
nad5
rps4C
rps4M
rps4N
rrn5
rps2
rpl6
rps8
trnL(tag)
nad4
rps11
rps10
tatC
complex I (NADH dehydrogenase)
complex III (ubichinol cytochrome c reductase)
complex IV (cytochrome c oxidase)
ATP synthase
ribosomal proteins (SSU)
ribosomal proteins (LSU)
other genes
ORFs
transfer RNAs
ribosomal RNAs

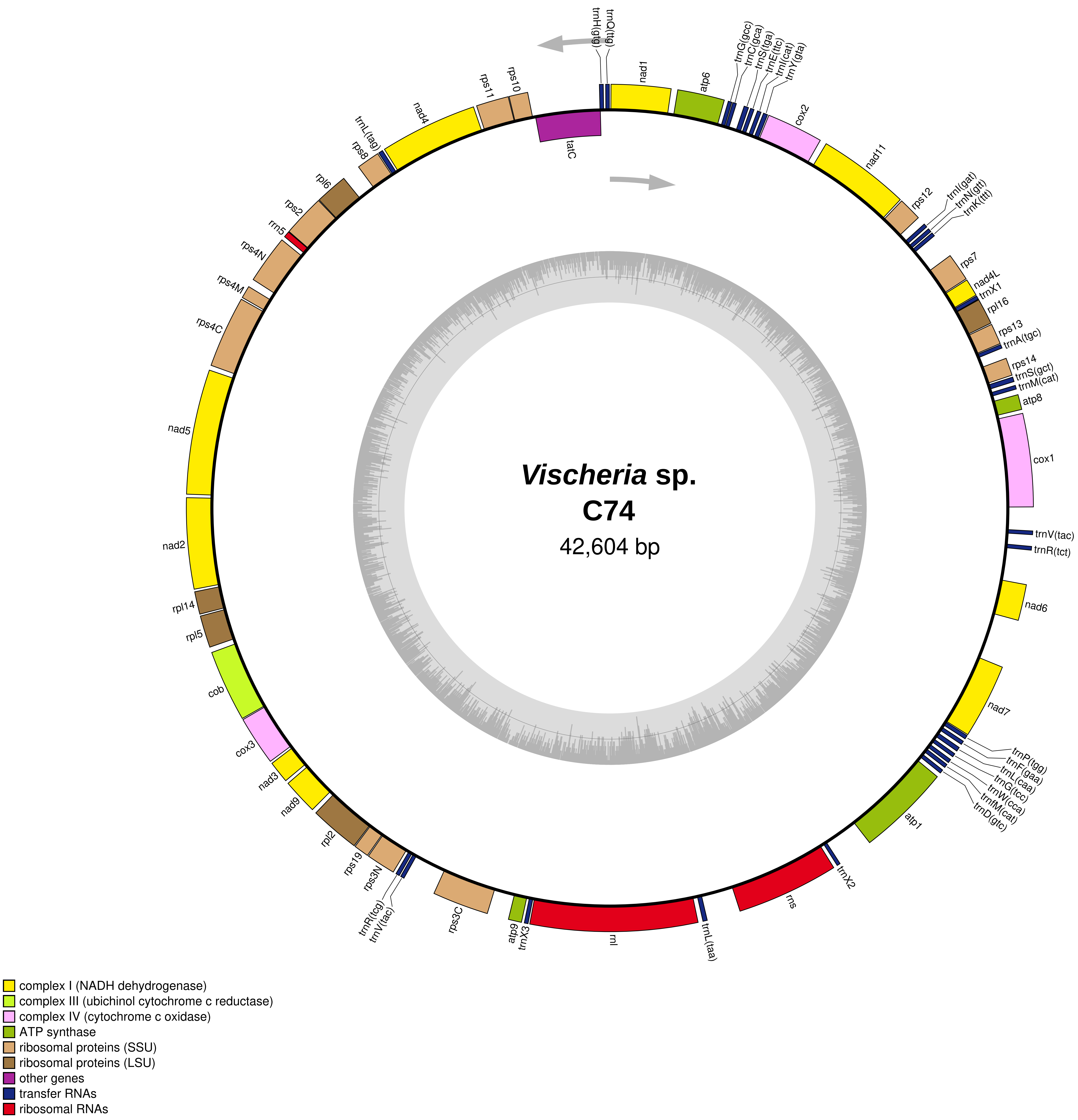
*Vischeria* sp.
C74
42,604 bp
trnH(gtg)
trnQ(ttg)
nad1
atp6
trnG(gcc)
trnC(gca)
trnS(tga)
trnE(ttc)
trnI(cat)
trnY(gta)
cox2
nad11
rps12
trnI(gat)
trnN(gtt)
trnK(ttt)
rps7
nad4L
trnX1
rpl16
rps13
trnA(tgc)
rps14
trnS(gct)
trnM(cat)
atp8
cox1
trnV(tac)
trnR(tct)
nad6
nad7
trnP(tgg)
trnF(gaa)
trnL(caa)
trnG(tcc)
trnW(cca)
trnfM(cat)
trnD(gtc)
atp1
trnX2
rns
trnL(taa)
rnl
trnX3
atp9
rps3C
trnV(tac)
trnR(tcg)
rps3N
rps19
rpl2
nad9
nad3
cox3
cob
rpl5
rpl14
nad2
nad5
rps4C
rps4M
rps4N
rrn5
rps2
rpl6
rps8
trnL(tag)
nad4
rps11
rps10
tatC
complex I (NADH dehydrogenase)
complex III (ubichinol cytochrome c reductase)
complex IV (cytochrome c oxidase)
ATP synthase
ribosomal proteins (SSU)
ribosomal proteins (LSU)
other genes
transfer RNAs
ribosomal RNAs

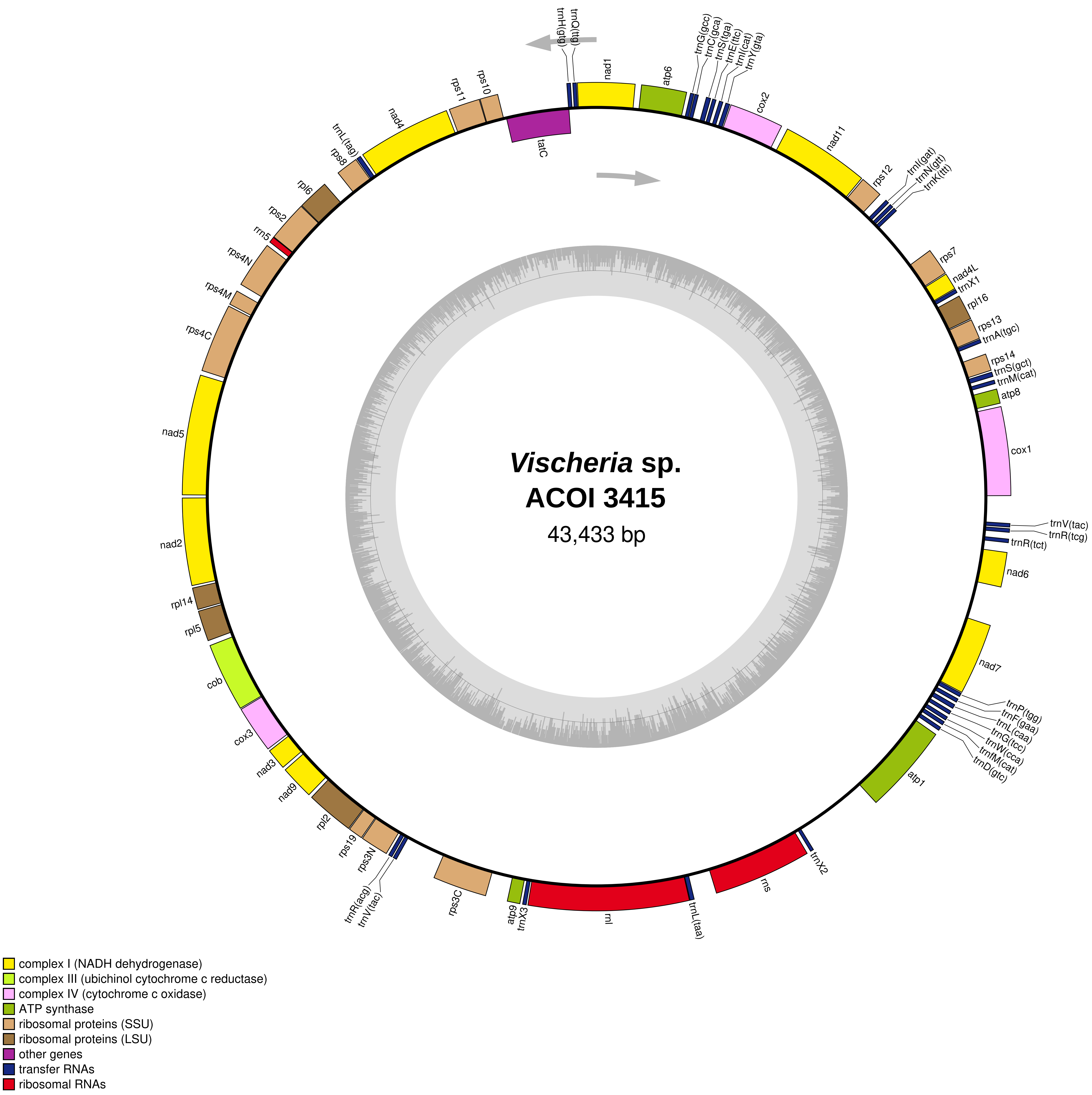

Vischeria sp.
ACOI 3415
43,433 bp
trnH(gtg)
trnQ(ttg)
nad1
atp6
trnG(gcc)
trnC(gca)
trnS(tga)
trnE(ttc)
trnI(cat)
trnY(gta)
cox2
nad11
rps12
trnI(gat)
trnN(gtt)
trnK(ttt)
rps7
nad4L
trnX1
rpl16
rps13
trnA(tgc)
rps14
trnS(gct)
trnM(cat)
atp8
cox1
trnV(tac)
trnR(tcg)
trnR(tct)
nad6
nad7
trnP(tgg)
trnF(gaa)
trnL(caa)
trnG(tcc)
trnW(cca)
trnfM(cat)
trnD(gtc)
atp1
trnX2
rns
trnL(taa)
rnl
trnX3
atp9
rps3C
trnV(tac)
trnR(acg)
rps3N
rps19
rpl2
nad9
nad3
cox3
cob
rpl5
rpl14
nad2
nad5
rps4C
rps4M
rps4N
rrn5
rps2
rpl6
rps8
trnL(tag)
nad4
rps11
rps10
tatC
complex I (NADH dehydrogenase)
complex III (ubichinol cytochrome c reductase)
complex IV (cytochrome c oxidase)
ATP synthase
ribosomal proteins (SSU)
ribosomal proteins (LSU)
other genes
transfer RNAs
ribosomal RNAs

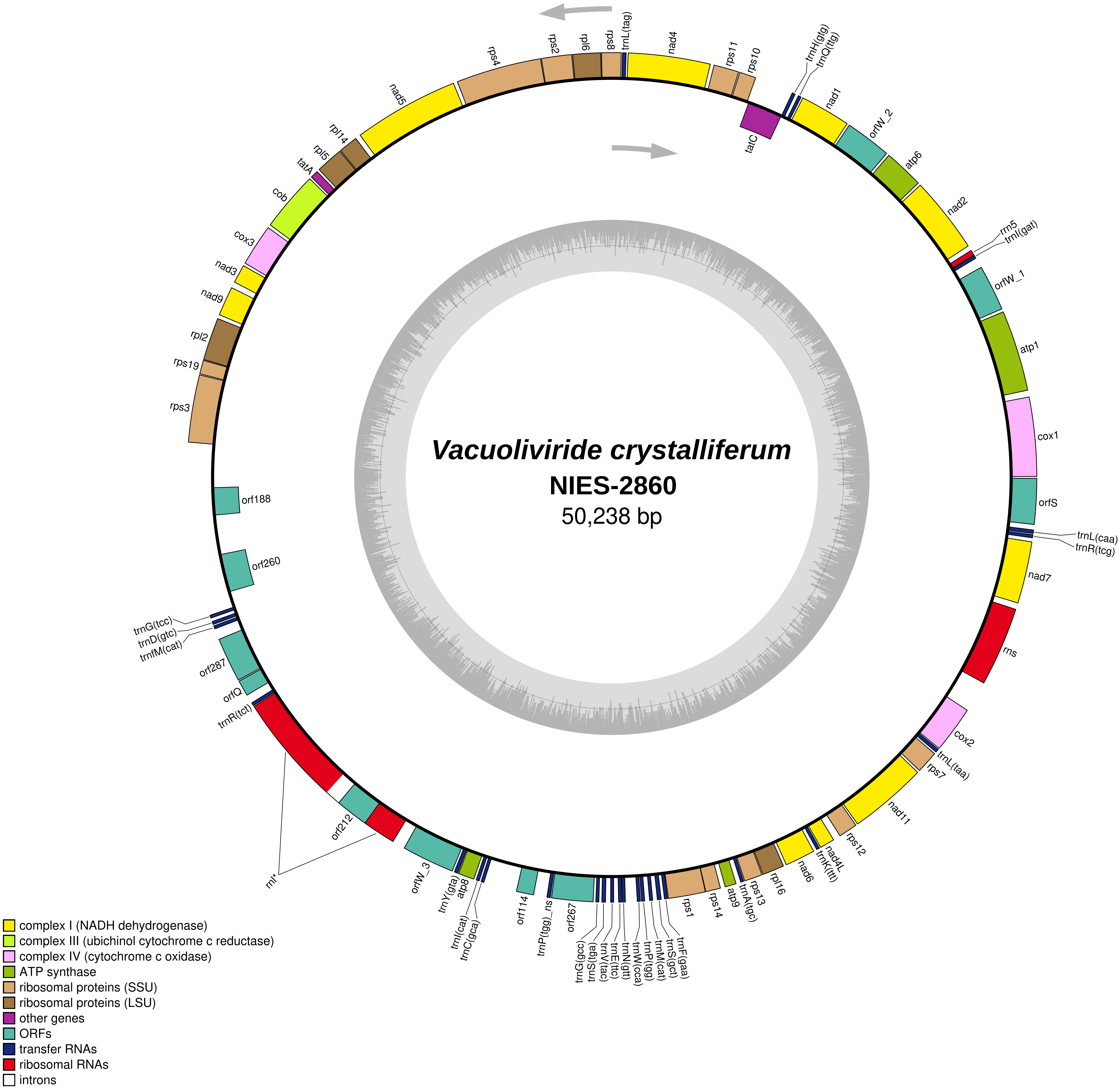

Vacuoliviride crystalliferum
NIES-2860
50,238 bp
trnL(tag)
nad4
rps11
rps10
trnH(gtg)
trnQ(ttg)
tatC
nad1
orfW_2
atp6
nad2
rrn5
trnI(gat)
orfW_1
atp1
cox1
orfS
trnL(caa)
trnR(tcg)
nad7
rns
cox2
trnL(taa)
rps7
nad11
rps12
nad4L
trnK(ttt)
nad6
rpl16
rps13
trnA(tgc)
atp9
rps14
rps1
trnF(gaa)
trnS(gct)
trnM(cat)
trnP(tgg)
trnW(cca)
trnN(gtt)
trnE(ttc)
trnV(tac)
trnS(tga)
trnG(gcc)
orf267
trnP(tgg)_ns
orf114
trnC(gca)
trnI(cat)
atp8
trnY(gta)
orfW_3
rnl*
orf212
trnR(tct)
orfQ
orf287
trnfM(cat)
trnD(gtc)
trnG(tcc)
orf260
orf188
rps3
rps19
rpl2
nad9
nad3
cox3
cob
tatA
rpl5
rpl14
nad5
rps4
rps2
rpl6
rps8
complex I (NADH dehydrogenase)
complex III (ubichinol cytochrome c reductase)
complex IV (cytochrome c oxidase)
ATP synthase
ribosomal proteins (SSU)
ribosomal proteins (LSU)
other genes
ORFs
transfer RNAs
ribosomal RNAs
introns

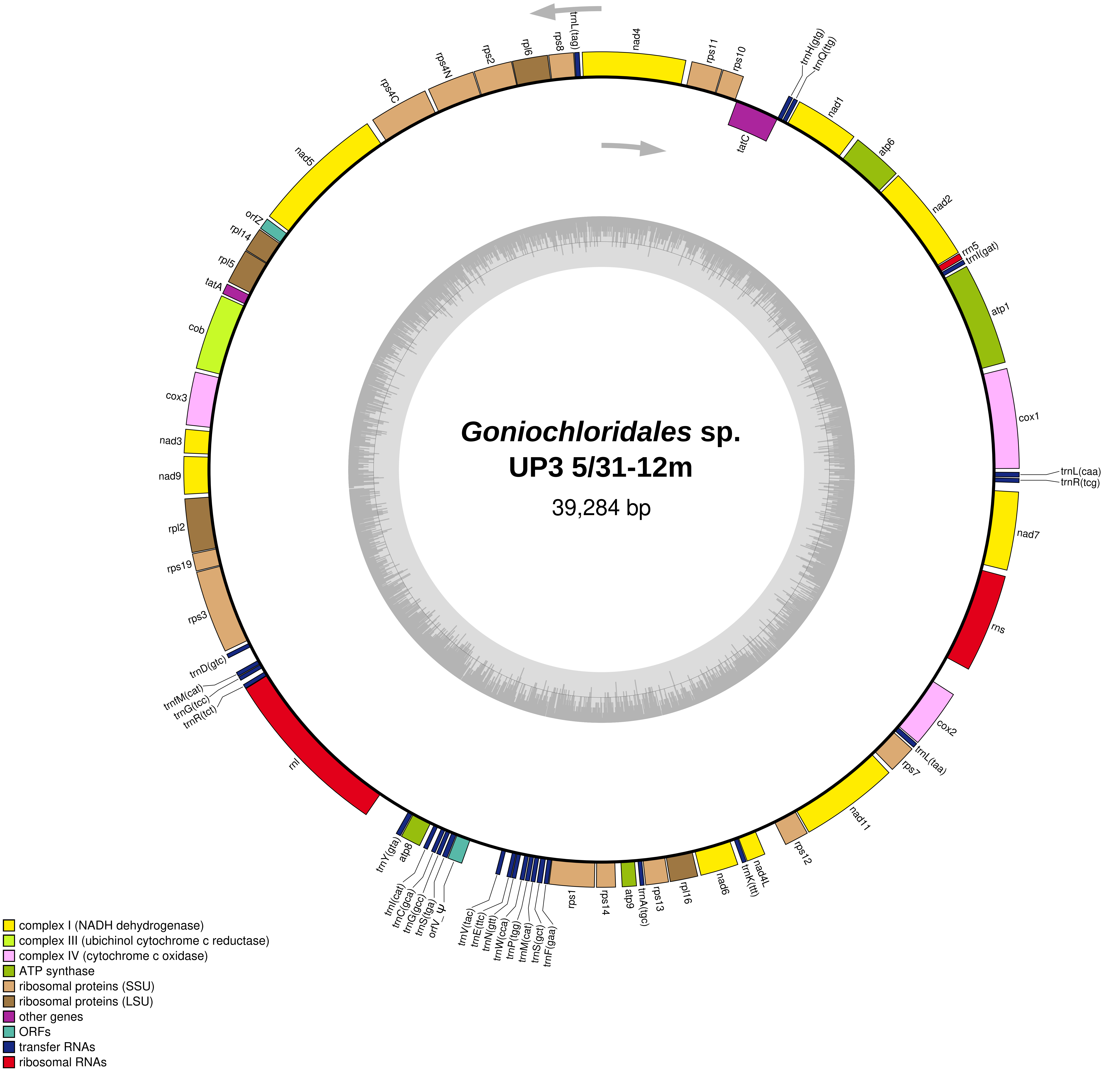
Goniochloridales sp.
UP3 5/31-12m
39,284 bp
trnL(tag)
nad4
rps11
rps10
trnH(gtg)
trnQ(ttg)
tatC
nad1
atp6
nad2
rrn5
trnI(gat)
atp1
cox1
trnL(caa)
trnR(tcg)
nad7
rns
cox2
trnL(taa)
rps7
nad11
rps12
nad4L
trnK(ttt)
nad6
rpl16
rps13
trnA(tgc)
atp9
rps14
rps1
trnF(gaa)
trnS(gct)
trnM(cat)
trnP(tgg)
trnW(cca)
trnN(gtt)
trnE(ttc)
trnV(tac)
orfV_ψ
trnS(tga)
trnG(gcc)
trnC(gca)
trnI(cat)
atp8
trnY(gta)
rnl
trnR(tct)
trnG(tcc)
trnfM(cat)
trnD(gtc)
rps3
rps19
rpl2
nad9
nad3
cox3
cob
tatA
rpl5
rpl14
orfZ
nad5
rps4C
rps4N
rps2
rpl6
rps8
complex I (NADH dehydrogenase)
complex III (ubichinol cytochrome c reductase)
complex IV (cytochrome c oxidase)
ATP synthase
ribosomal proteins (SSU)
ribosomal proteins (LSU)
other genes
ORFs
transfer RNAs
ribosomal RNAs

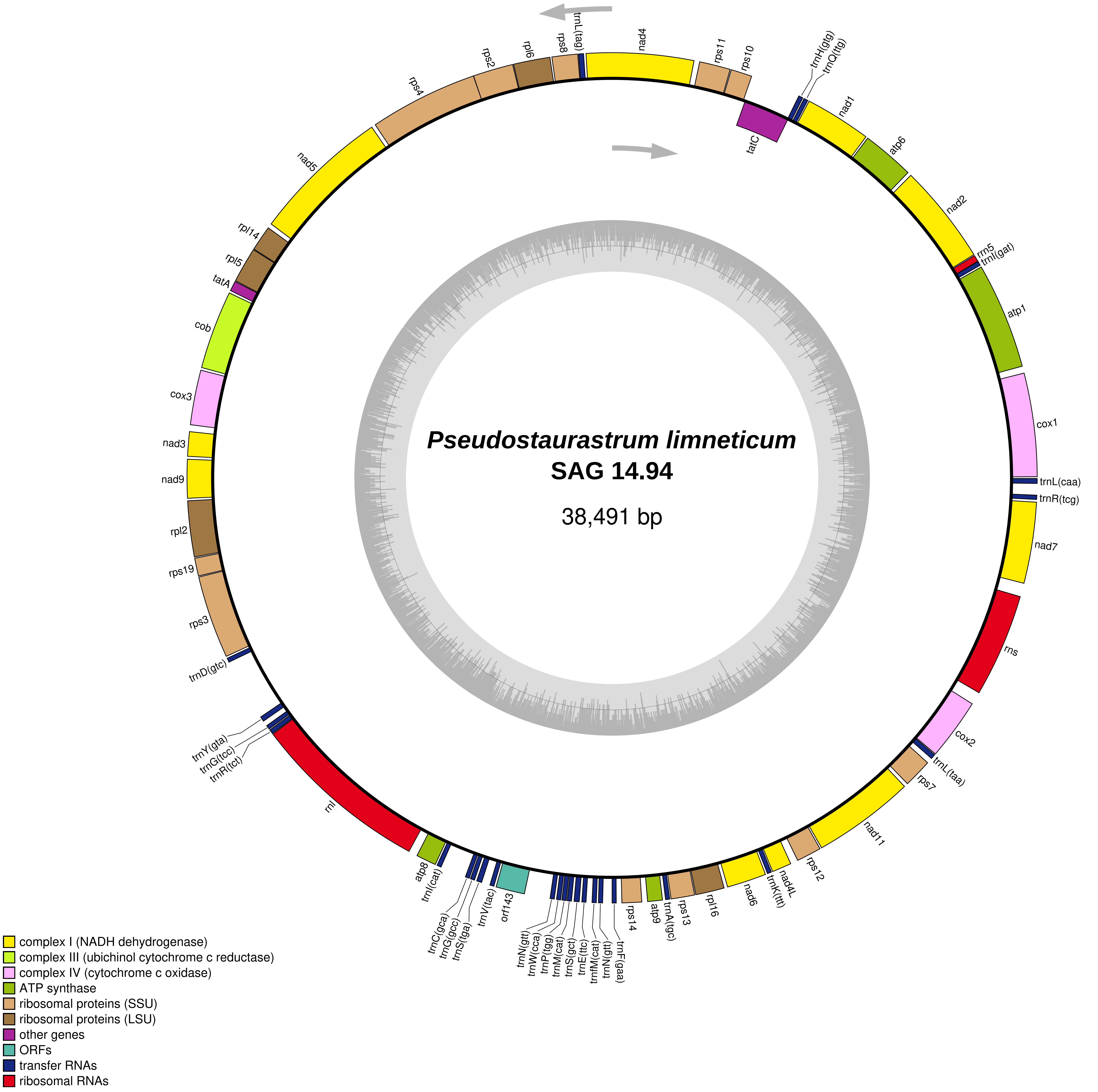

Pseudostaurastrum limneticum
SAG 14.94
38,491 bp
nad4
rps11
rps10
trnH(gtg)
trnQ(ttg)
tatC
nad1
atp6
nad2
rrn5
trnI(gat)
atp1
cox1
trnL(caa)
trnR(tcg)
nad7
rns
cox2
trnL(taa)
rps7
nad11
rps12
nad4L
trnK(ttt)
nad6
rpl16
rps13
trnA(tgc)
atp9
rps14
trnF(gaa)
trnN(gtt)
trnfM(cat)
trnE(ttc)
trnS(gct)
trnM(cat)
trnP(tgg)
trnW(cca)
trnN(gtt)
orf143
trnV(tac)
trnS(tga)
trnG(gcc)
trnC(gca)
trnI(cat)
atp8
rnl
trnR(tct)
trnG(tcc)
trnY(gta)
trnD(gtc)
rps3
rps19
rpl2
nad9
nad3
cox3
cob
tatA
rpl5
rpl14
nad5
rps4
rps2
rpl6
rps8
trnL(tag)
complex I (NADH dehydrogenase)
complex III (ubichinol cytochrome c reductase)
complex IV (cytochrome c oxidase)
ATP synthase
ribosomal proteins (SSU)
ribosomal proteins (LSU)
other genes
ORFs
transfer RNAs
ribosomal RNAs

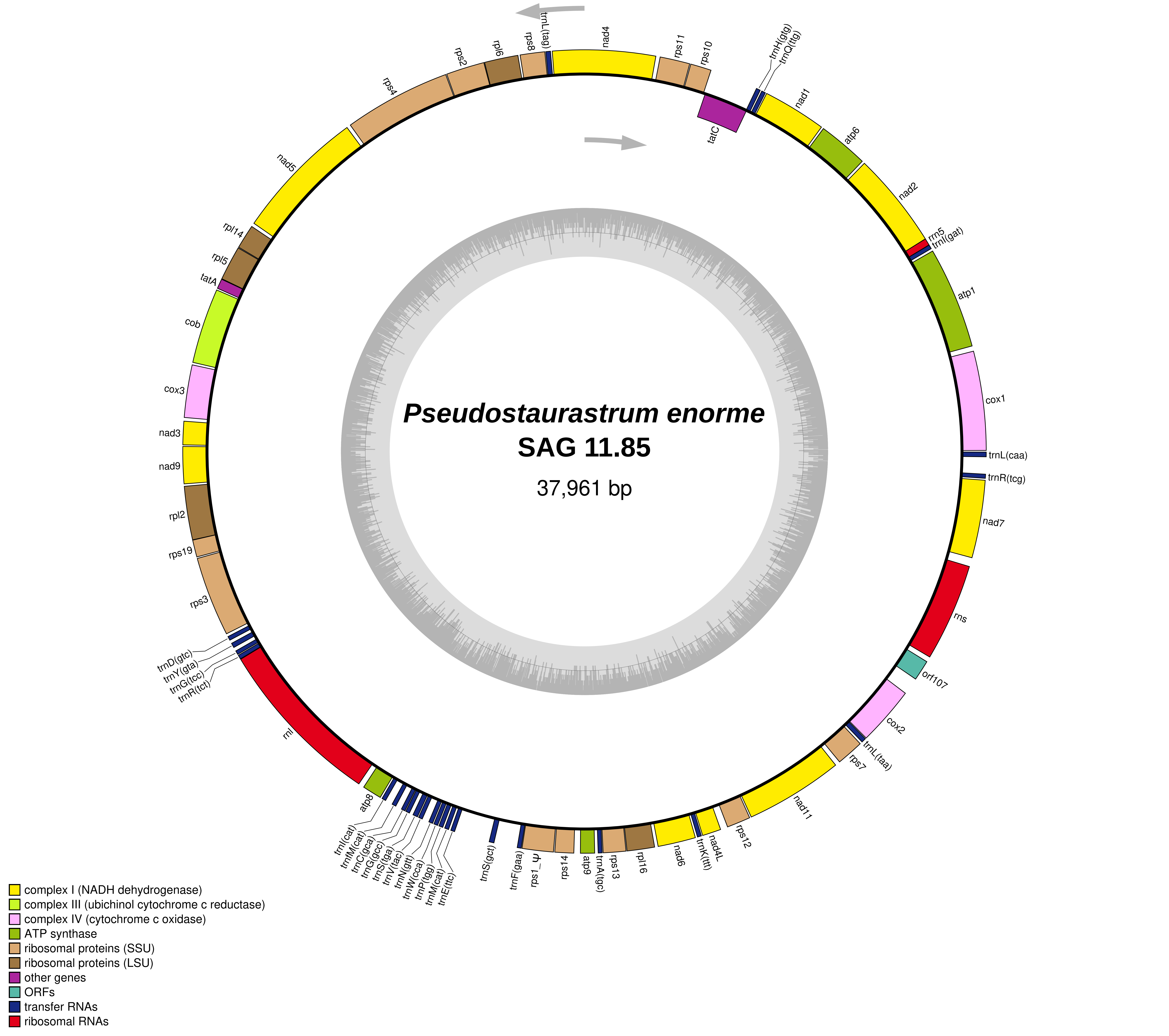
Pseudostaurastrum enorme
SAG 11.85
37,961 bp
nad4
rps11
rps10
trnL(tag)
rps8
rpl6
rps2
rps4
nad5
rpl14
rpl5
tatA
cob
cox3
nad3
nad9
rpl2
rps19
rps3
trnD(gtc)
trnY(gta)
trnG(tcc)
trnR(tct)
rnl
atp8
trnI(cat)
trnfM(cat)
trnC(gca)
trnG(gcc)
trnS(tga)
trnV(tac)
trnN(gtt)
trnW(cca)
trnP(tgg)
trnM(cat)
trnE(ttc)
trnS(gct)
trnF(gaa)
rps1_ψ
rps14
atp9
trnA(tgc)
rps13
rpl16
nad6
trnK(ttt)
nad4L
rps12
nad11
rps7
trnL(taa)
cox2
orf107
rns
nad7
trnR(tcg)
trnL(caa)
cox1
atp1
trnI(gat)
rrn5
nad2
atp6
nad1
trnQ(ttg)
trnH(gtg)
tatC
complex I (NADH dehydrogenase)
complex III (ubichinol cytochrome c reductase)
complex IV (cytochrome c oxidase)
ATP synthase
ribosomal proteins (SSU)
ribosomal proteins (LSU)
other genes
ORFs
transfer RNAs
ribosomal RNAs

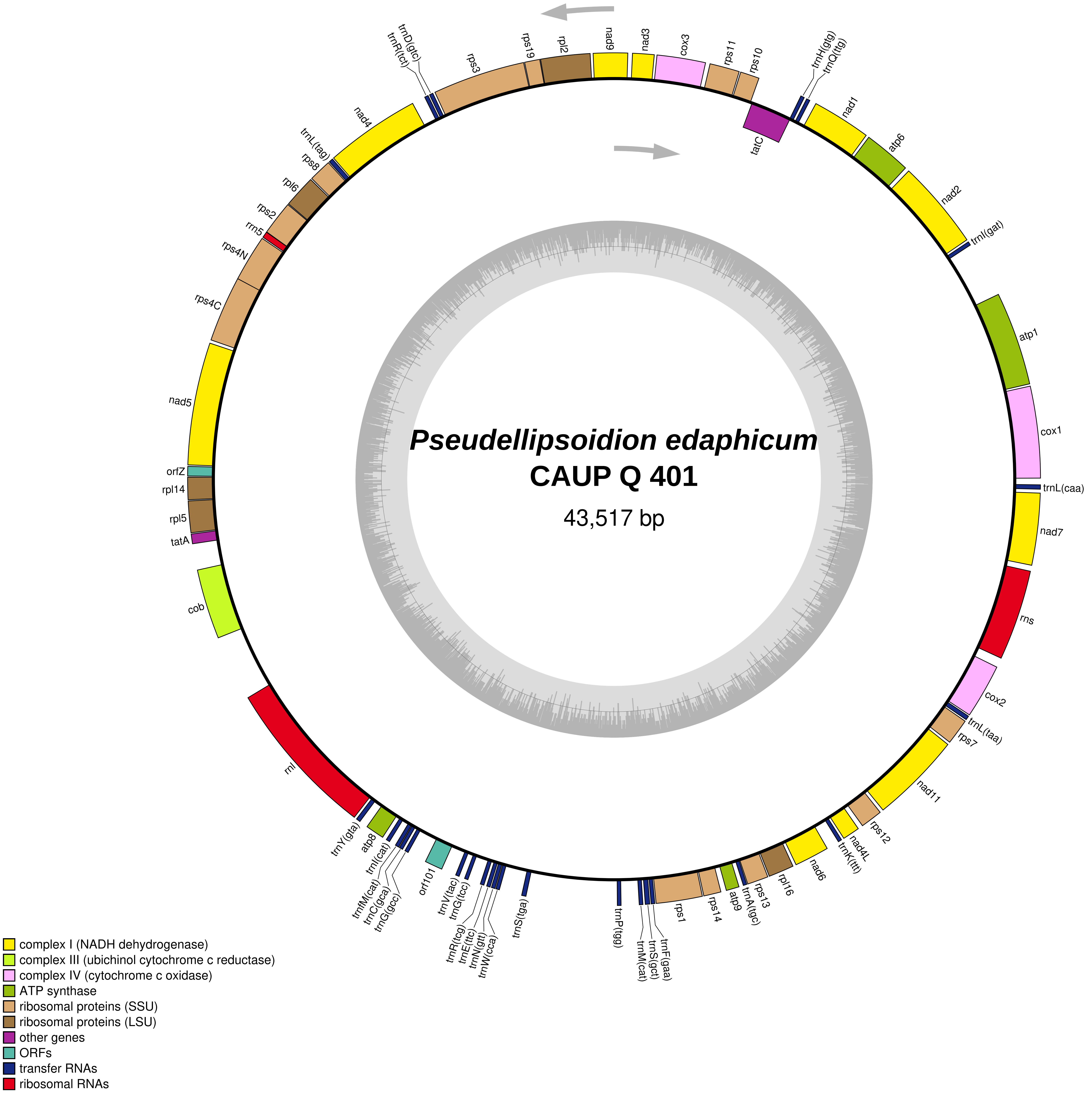

Pseudellipsoidion edaphicum
CAUP Q 401
43,517 bp
complex I (NADH dehydrogenase)
complex III (ubichinol cytochrome c reductase)
complex IV (cytochrome c oxidase)
ATP synthase
ribosomal proteins (SSU)
ribosomal proteins (LSU)
other genes
ORFs
transfer RNAs
ribosomal RNAs
nad9
nad3
cox3
rps11
rps10
rpl2
rps19
rps3
trnD(gtc)
trnR(tct)
trnH(gtg)
trnQ(ttg)
tatC
nad1
atp6
nad2
trnI(gat)
atp1
cox1
trnL(caa)
nad7
rns
cox2
trnL(taa)
rps7
nad11
rps12
nad4L
trnK(ttt)
nad6
rpl16
rps13
trnA(tgc)
atp9
rps14
rps1
trnF(gaa)
trnS(gct)
trnM(cat)
trnP(tgg)
trnS(tga)
trnW(cca)
trnN(gtt)
trnE(ttc)
trnR(tcg)
trnG(tcc)
trnV(tac)
orf101
trnG(gcc)
trnC(gca)
trnfM(cat)
trnI(cat)
atp8
trnY(gta)
rnl
cob
tatA
rpl5
rpl14
orfZ
nad5
rps4C
rps4N
rrn5
rps2
rpl6
rps8
trnL(tag)
nad4

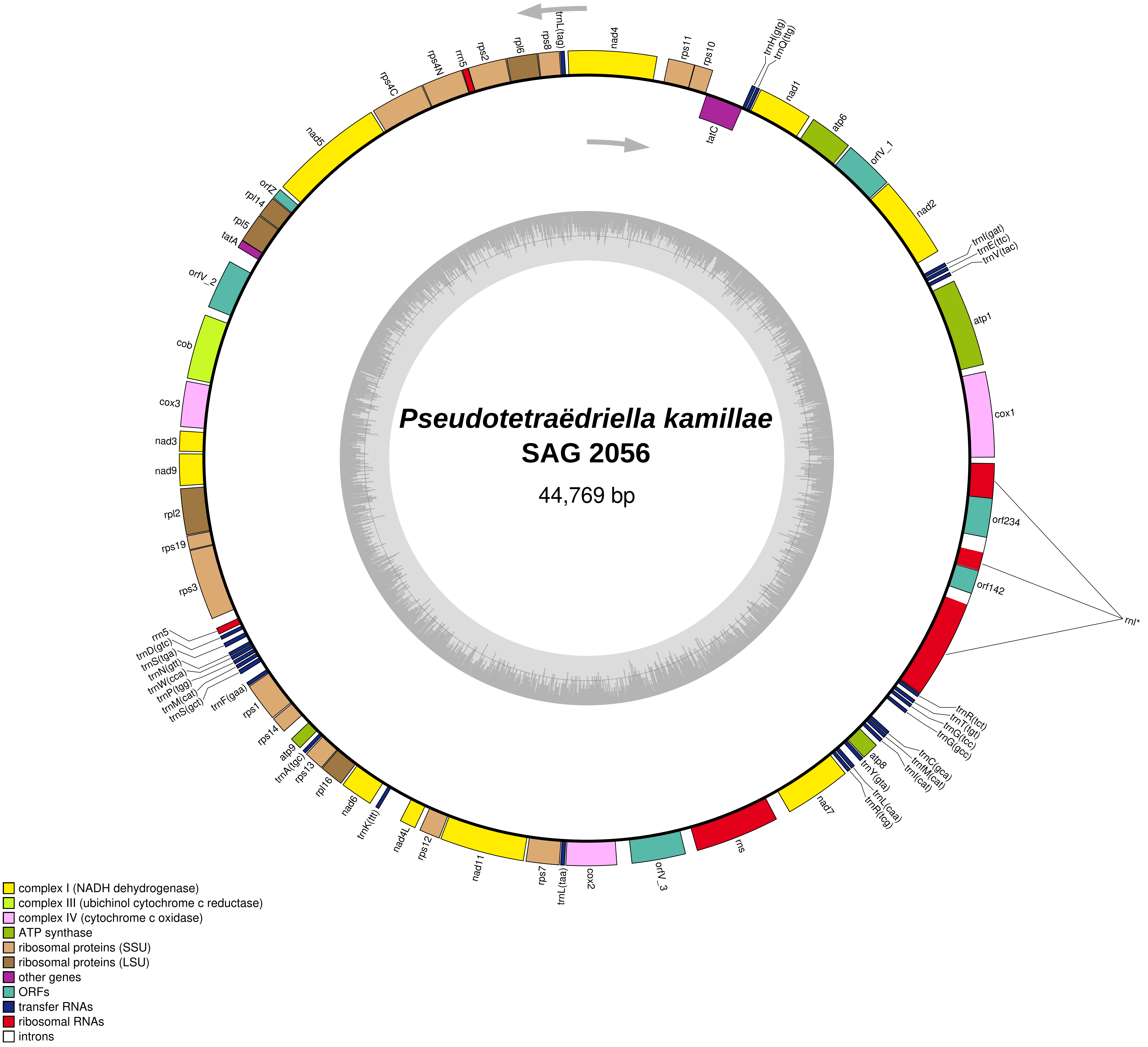
Pseudotetraëdriella kamillae
SAG 2056
44,769 bp
trnL(tag)
nad4
rps11
rps10
trnH(gtg)
trnQ(ttg)
tatC
nad1
atp6
orfV_1
nad2
trnI(gat)
trnE(ttc)
trnV(tac)
atp1
cox1
orf234
orf142
rnl*
trnR(tct)
trnT(tgt)
trnG(tcc)
trnG(gcc)
trnC(gca)
trnfM(cat)
trnI(cat)
atp8
trnY(gta)
trnL(caa)
trnR(tcg)
nad7
rns
orfV_3
cox2
trnL(taa)
rps7
nad11
rps12
nad4L
trnK(ttt)
nad6
rpl16
rps13
trnA(tgc)
atp9
rps14
rps1
trnF(gaa)
trnS(gct)
trnM(cat)
trnP(tgg)
trnW(cca)
trnN(gtt)
trnS(tga)
trnD(gtc)
rrn5
rps3
rps19
rpl2
nad9
nad3
cox3
cob
orfV_2
tatA
rpl5
rpl14
orfZ
nad5
rps4C
rps4N
rrn5
rps2
rpl6
rps8
complex I (NADH dehydrogenase)
complex III (ubichinol cytochrome c reductase)
complex IV (cytochrome c oxidase)
ATP synthase
ribosomal proteins (SSU)
ribosomal proteins (LSU)
other genes
ORFs
transfer RNAs
ribosomal RNAs
introns

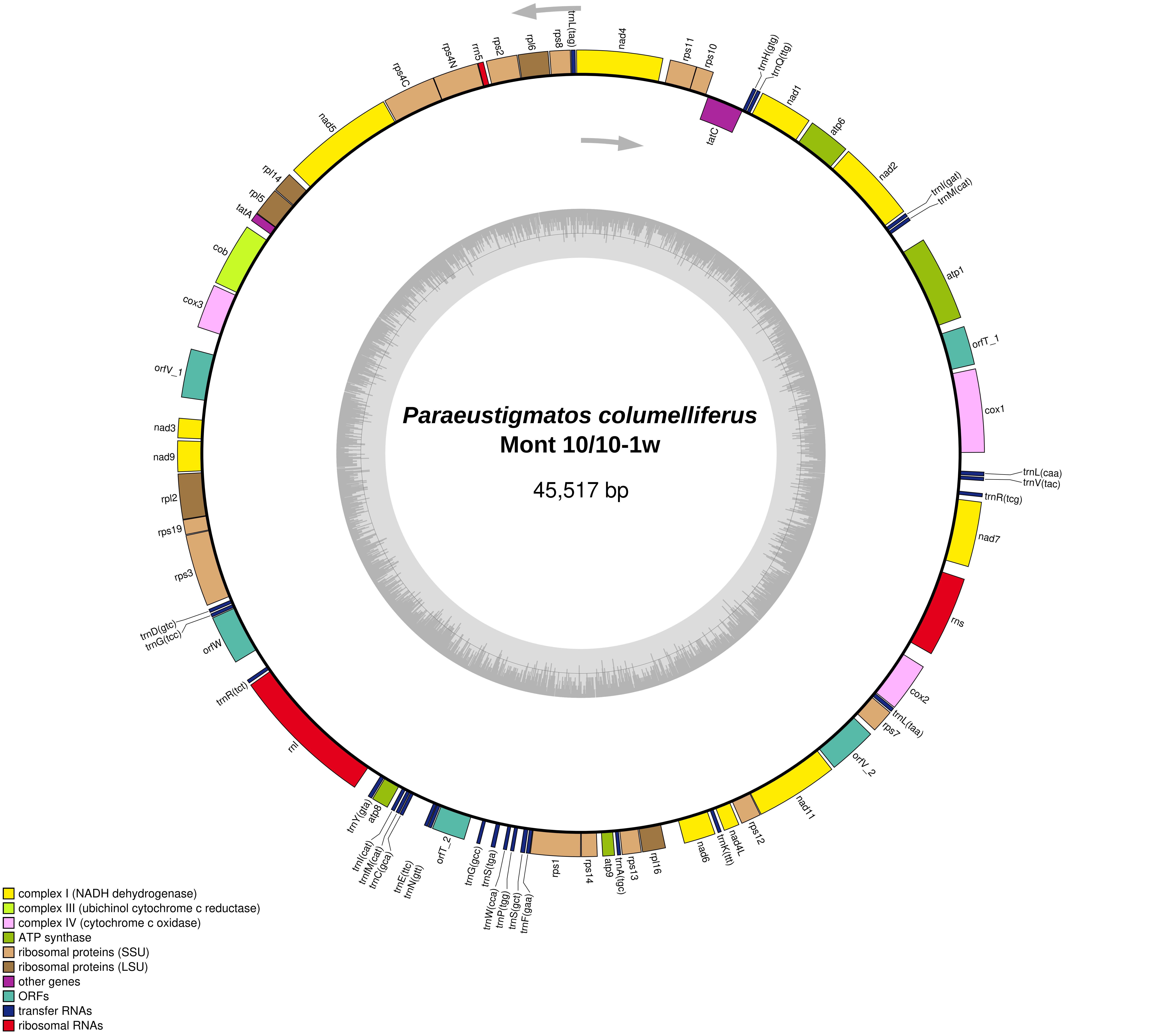
Paraeustigmatos columelliferus
Mont 10/10-1w
45,517 bp
trnL(tag)
nad4
rps11
rps10
trnH(gtg)
trnQ(ttg)
tatC
nad1
atp6
nad2
trnI(gat)
trnM(cat)
atp1
orfT_1
cox1
trnL(caa)
trnV(tac)
trnR(tcg)
nad7
rns
cox2
trnL(taa)
rps7
orfV_2
nad11
rps12
nad4L
trnK(ttt)
nad6
rpl16
rps13
trnA(tgc)
atp9
rps14
rpsl
trnF(gaa)
trnS(gct)
trnP(tgg)
trnW(cca)
trnS(tga)
trnG(gcc)
orfT_2
trnN(gtt)
trnE(ttc)
trnC(gca)
trnfM(cat)
trnI(cat)
atp8
trnY(gta)
rnl
trnR(tct)
orfW
trnG(tcc)
trnD(gtc)
rps3
rps19
rpl2
nad9
nad3
orfV_1
cox3
cob
tatA
rpl5
rpl14
nad5
rps4C
rps4N
rrn5
rps2
rpl6
rps8
complex I (NADH dehydrogenase)
complex III (ubichinol cytochrome c reductase)
complex IV (cytochrome c oxidase)
ATP synthase
ribosomal proteins (SSU)
ribosomal proteins (LSU)
other genes
ORFs
transfer RNAs
ribosomal RNAs

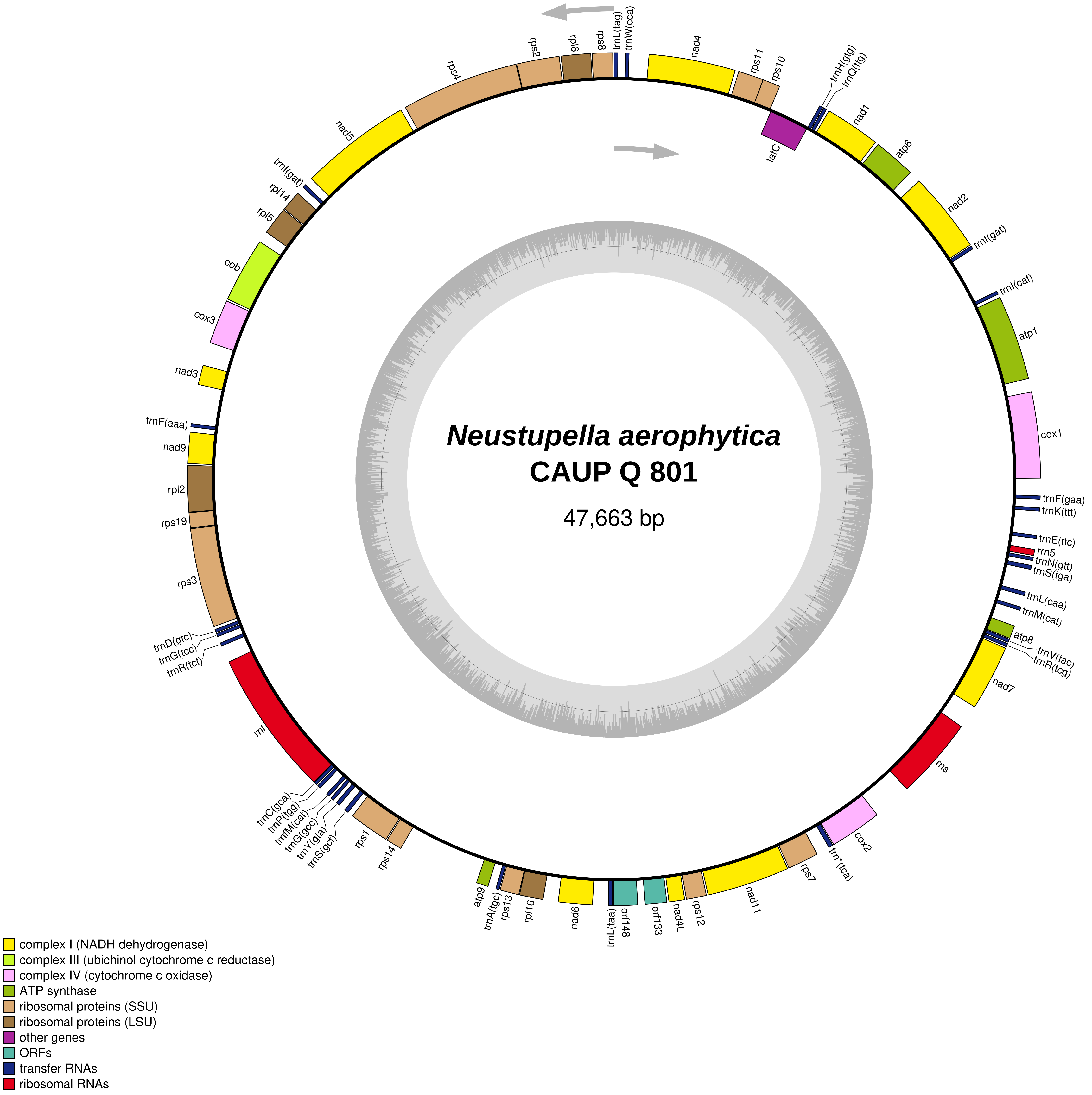
Neustupella aerophytica
CAUP Q 801
47,663 bp
nad4
rps11
rps10
trnH(gtg)
trnQ(ttg)
tatC
nad1
atp6
nad2
trnI(gat)
trnI(cat)
atp1
cox1
trnF(gaa)
trnK(ttt)
trnE(ttc)
rrn5
trnN(gtt)
trnS(tga)
trnL(caa)
trnM(cat)
atp8
trnV(tac)
trnR(tcg)
nad7
rns
cox2
trn*(tca)
rps7
nad11
rps12
nad4L
orf133
orf148
trnL(taa)
nad6
rpl16
rps13
trnA(tgc)
atp9
rps14
rps1
trnS(gct)
trnY(gta)
trnG(gcc)
trnfM(cat)
trnP(tgg)
trnC(gca)
rnl
trnR(tct)
trnG(tcc)
trnD(gtc)
rps3
rps19
rpl2
nad9
trnF(aaa)
nad3
cox3
cob
rpl5
rpl14
trnI(gat)
nad5
rps4
rps2
rpl6
rps8
trnL(tag)
trnW(cca)
complex I (NADH dehydrogenase)
complex III (ubichinol cytochrome c reductase)
complex IV (cytochrome c oxidase)
ATP synthase
ribosomal proteins (SSU)
ribosomal proteins (LSU)
other genes
ORFs
transfer RNAs
ribosomal RNAs

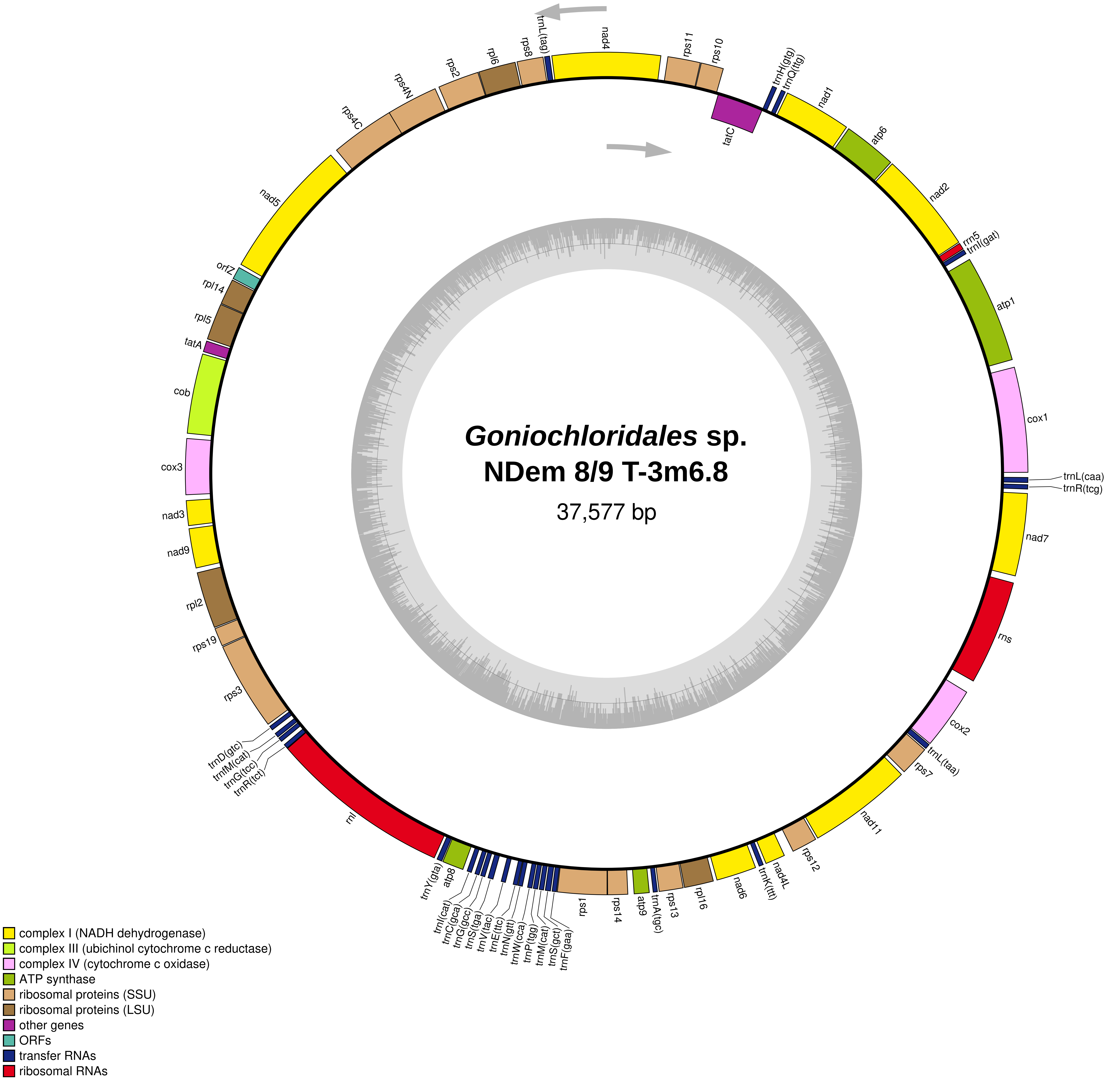

*Goniochloridales* sp.
NDem 8/9 T-3m6.8
37,577 bp
trnL(tag)
nad4
rps11
rps10
tatC
trnH(gtg)
trnQ(ttg)
nad1
atp6
nad2
rrn5
trnI(gat)
atp1
cox1
trnL(caa)
trnR(tcg)
nad7
rns
cox2
trnL(taa)
rps7
nad11
rps12
nad4L
trnK(ttt)
nad6
rpl16
rps13
trnA(tgc)
atp9
rps14
rps1
trnF(gaa)
trnS(gct)
trnM(cat)
trnP(tgg)
trnW(cca)
trnN(gtt)
trnE(ttc)
trnV(tac)
trnS(tga)
trnG(gcc)
trnC(gca)
trnI(cat)
atp8
trnY(gta)
rnl
trnR(tct)
trnG(tcc)
trnfM(cat)
trnD(gtc)
rps3
rps19
rpl2
nad9
nad3
cox3
cob
tatA
rpl5
rpl14
orfZ
nad5
rps4C
rps4N
rps2
rpl6
rps8
complex I (NADH dehydrogenase)
complex III (ubichinol cytochrome c reductase)
complex IV (cytochrome c oxidase)
ATP synthase
ribosomal proteins (SSU)
ribosomal proteins (LSU)
other genes
ORFs
transfer RNAs
ribosomal RNAs

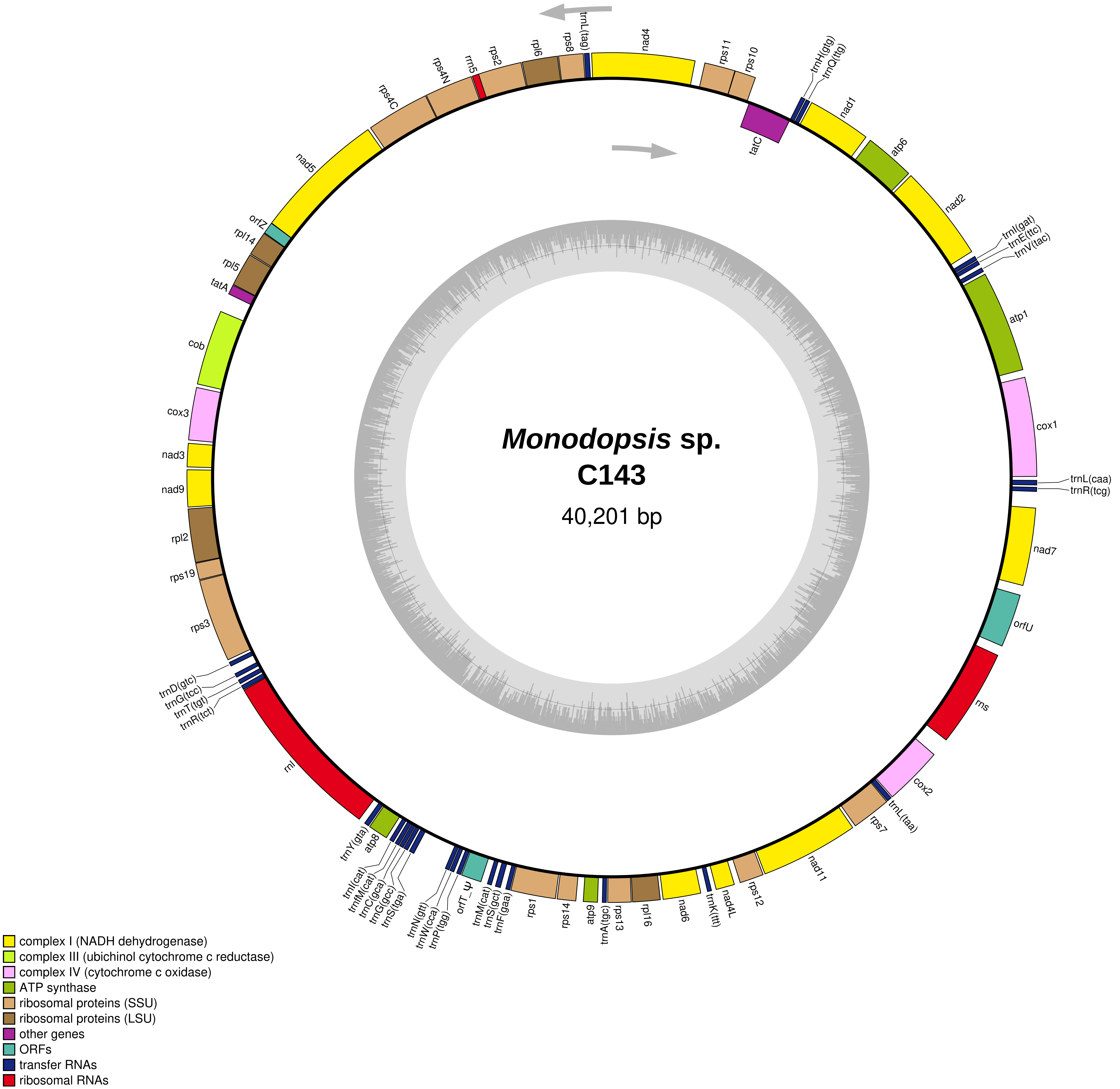
Monodopsis sp.
C143
40,201 bp
trnL(tag)
nad4
rps11
rps10
trnH(gtg)
trnQ(ttg)
tatC
nad1
atp6
nad2
trnI(gat)
trnE(ttc)
trnV(tac)
atp1
cox1
trnL(caa)
trnR(tcg)
nad7
orfU
rns
cox2
trnL(taa)
rps7
nad11
rps12
nad4L
trnK(ttt)
nad6
rpl16
rps13
trnA(tgc)
atp9
rps14
rps1
trnF(gaa)
trnS(gct)
trnM(cat)
orfT_ψ
trnP(tgg)
trnW(cca)
trnN(gtt)
trnS(tga)
trnG(gcc)
trnC(gca)
trnfM(cat)
trnI(cat)
atp8
trnY(gta)
rnl
trnR(tct)
trnT(tgt)
trnG(tcc)
trnD(gtc)
rps3
rps19
rpl2
nad9
nad3
cox3
cob
tatA
rpl5
rpl14
orfZ
nad5
rps4C
rps4N
rrn5
rps2
rpl6
rps8
complex I (NADH dehydrogenase)
complex III (ubichinol cytochrome c reductase)
complex IV (cytochrome c oxidase)
ATP synthase
ribosomal proteins (SSU)
ribosomal proteins (LSU)
other genes
ORFs
transfer RNAs
ribosomal RNAs

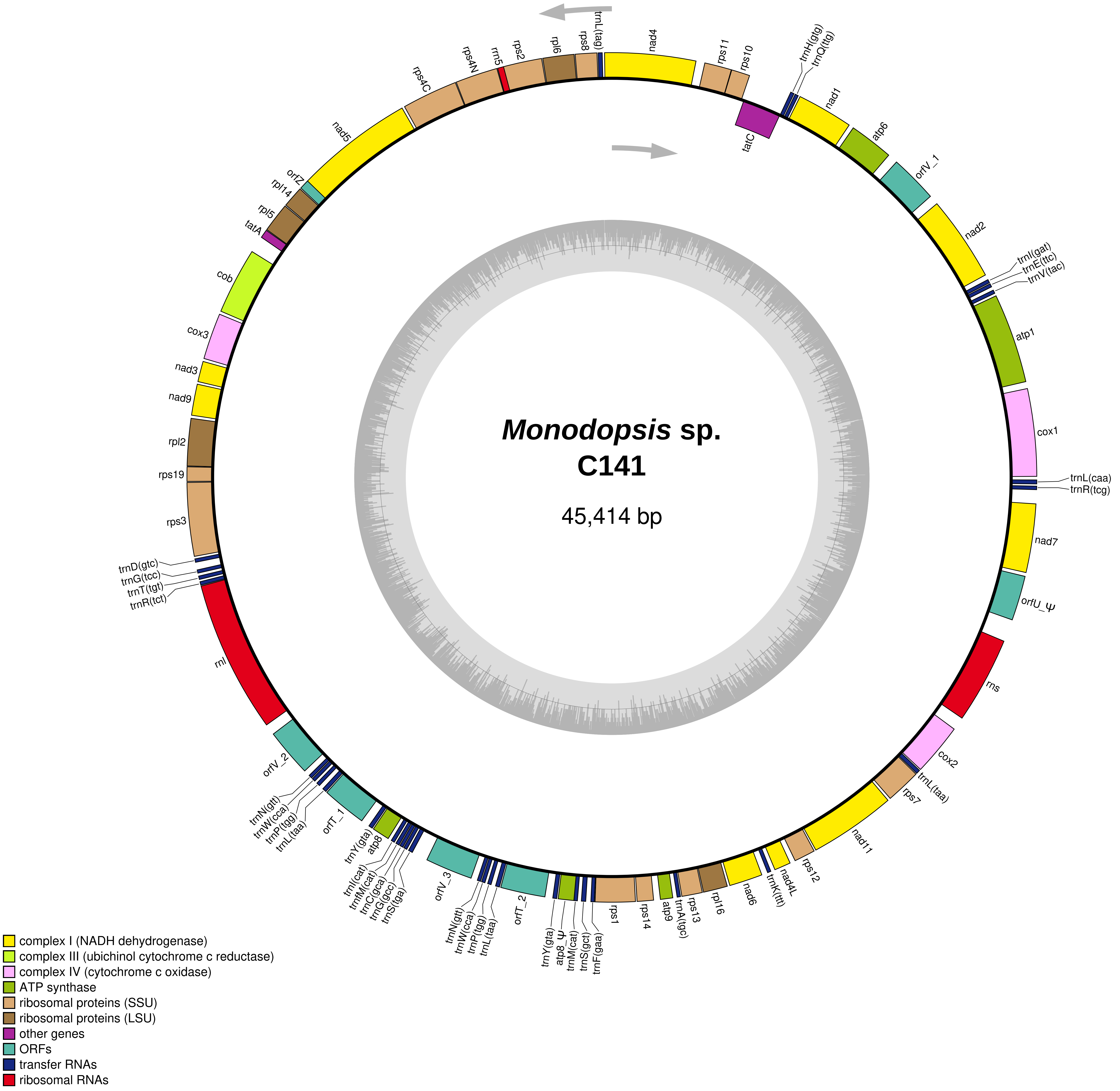
Monodopsis sp.
C141
45,414 bp
trnL(tag)
nad4
rps11
rps10
trnH(gtg)
trnQ(ttg)
tatC
nad1
atp6
orfV_1
nad2
trnI(gat)
trnE(ttc)
trnV(tac)
atp1
cox1
trnL(caa)
trnR(tcg)
nad7
orfU_Ψ
rns
cox2
trnL(taa)
rps7
nad11
rps12
nad4L
trnK(ttt)
nad6
rpl16
rps13
trnA(tgc)
atp9
rps14
rps1
trnF(gaa)
trnS(gct)
trnM(cat)
atp8_Ψ
trnY(gta)
orfT_2
trnL(taa)
trnP(tgg)
trnW(cca)
trnN(gtt)
orfV_3
trnS(tga)
trnG(gcc)
trnC(gca)
trnfM(cat)
trnI(cat)
atp8
trnY(gta)
orfT_1
trnL(taa)
trnP(tgg)
trnW(cca)
trnN(gtt)
orfV_2
rnl
trnR(tct)
trnT(tgt)
trnG(tcc)
trnD(gtc)
rps3
rps19
rpl2
nad9
nad3
cox3
cob
tatA
rpl5
rpl14
orfZ
nad5
rps4C
rps4N
rrn5
rps2
rpl6
rps8
complex I (NADH dehydrogenase)
complex III (ubichinol cytochrome c reductase)
complex IV (cytochrome c oxidase)
ATP synthase
ribosomal proteins (SSU)
ribosomal proteins (LSU)
other genes
ORFs
transfer RNAs
ribosomal RNAs

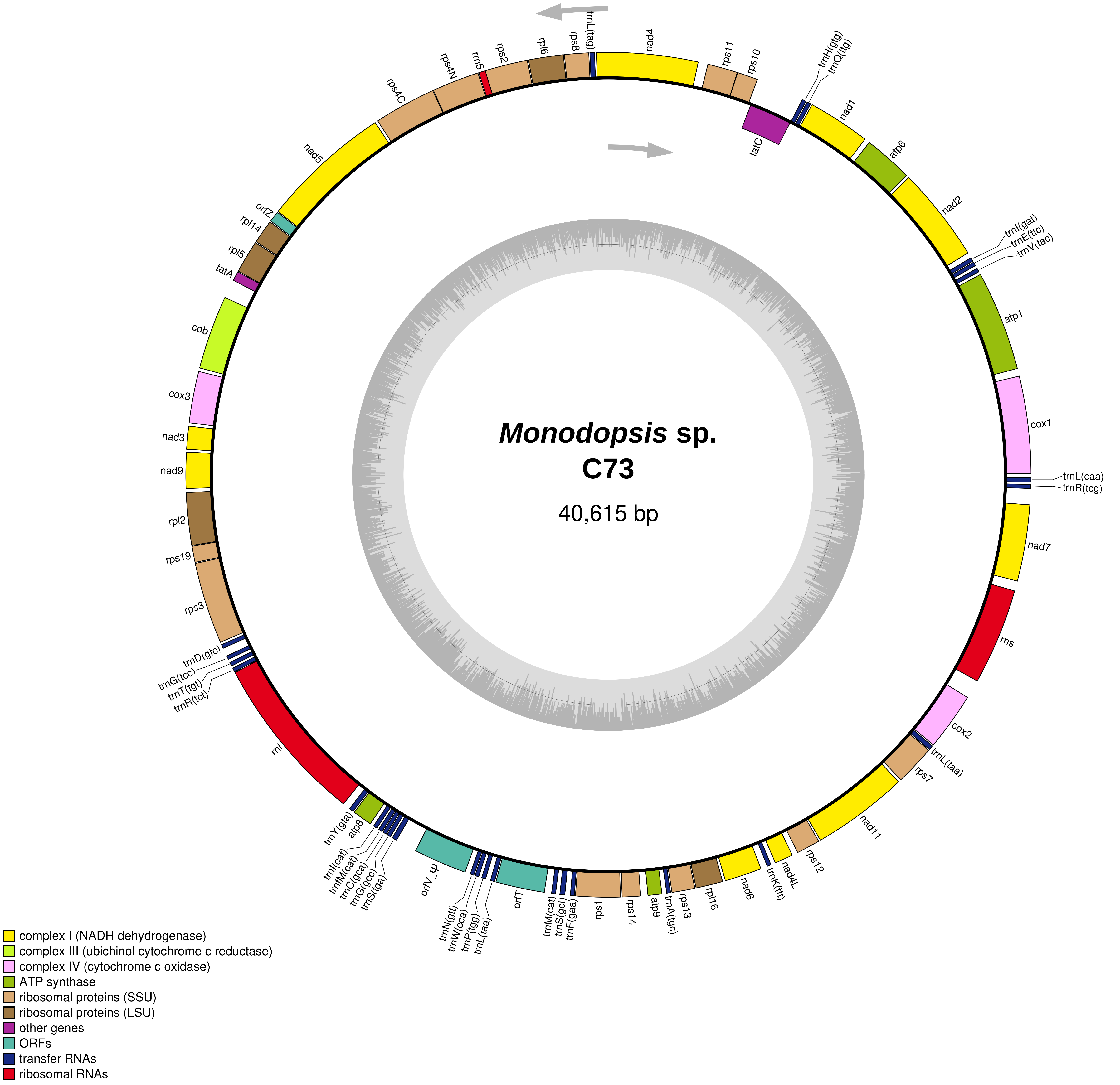
Monodopsis sp.
C73
40,615 bp
nad4
rps11
rps10
trnH(gtg)
trnQ(ttg)
tatC
nad1
atp6
nad2
trnI(gat)
trnE(ttc)
trnV(tac)
atp1
cox1
trnL(caa)
trnR(tcg)
nad7
rns
cox2
trnL(taa)
rps7
nad11
rps12
nad4L
trnK(ttt)
nad6
rpl16
rps13
trnA(tgc)
atp9
rps14
rps1
trnF(gaa)
trnS(gct)
trnM(cat)
orfT
trnL(taa)
trnP(tgg)
trnW(cca)
trnN(gtt)
orfV_ψ
trnS(tga)
trnG(gcc)
trnC(gca)
trnfM(cat)
trnI(cat)
atp8
trnY(gta)
rnl
trnR(tct)
trnT(tgt)
trnG(tcc)
trnD(gtc)
rps3
rps19
rpl2
nad9
nad3
cox3
cob
tatA
rpl5
rpl14
orfZ
nad5
rps4C
rps4N
rrn5
rps2
rpl6
rps8
trnL(tag)
complex I (NADH dehydrogenase)
complex III (ubichinol cytochrome c reductase)
complex IV (cytochrome c oxidase)
ATP synthase
ribosomal proteins (SSU)
ribosomal proteins (LSU)
other genes
ORFs
transfer RNAs
ribosomal RNAs

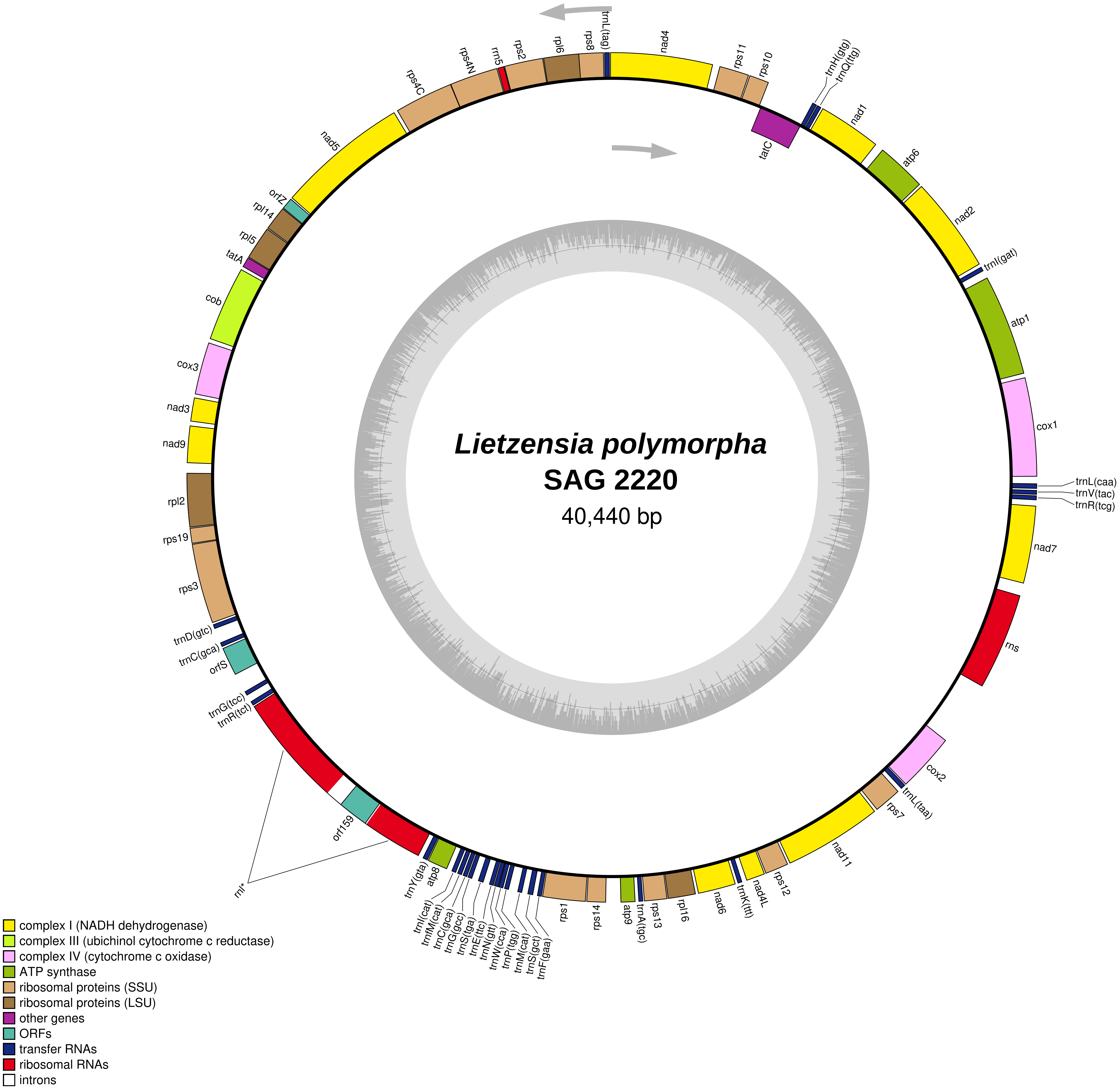

Lietzensia polymorpha
SAG 2220
40,440 bp
trnL(tag)
nad4
rps11
rps10
trnH(gtg)
trnQ(ttg)
tatC
nad1
atp6
nad2
trnI(gat)
atp1
cox1
trnL(caa)
trnV(tac)
trnR(tcg)
nad7
rns
cox2
trnL(taa)
rps7
nad11
rps12
nad4L
trnK(ttt)
nad6
rpl16
rps13
trnA(tgc)
atp9
rps14
rps1
trnF(gaa)
trnS(gct)
trnM(cat)
trnP(tgg)
trnW(cca)
trnN(gtt)
trnE(ttc)
trnS(tga)
trnG(gcc)
trnC(gca)
trnfM(cat)
trnI(cat)
atp8
trnY(gta)
orf159
rnl*
trnR(tct)
trnG(tcc)
orfS
trnC(gca)
trnD(gtc)
rps3
rps19
rpl2
nad9
nad3
cox3
cob
tatA
rpl5
rpl14
orfZ
nad5
rps4C
rps4N
rrn5
rps2
rpl6
rps8
complex I (NADH dehydrogenase)
complex III (ubichinol cytochrome c reductase)
complex IV (cytochrome c oxidase)
ATP synthase
ribosomal proteins (SSU)
ribosomal proteins (LSU)
other genes
ORFs
transfer RNAs
ribosomal RNAs
introns

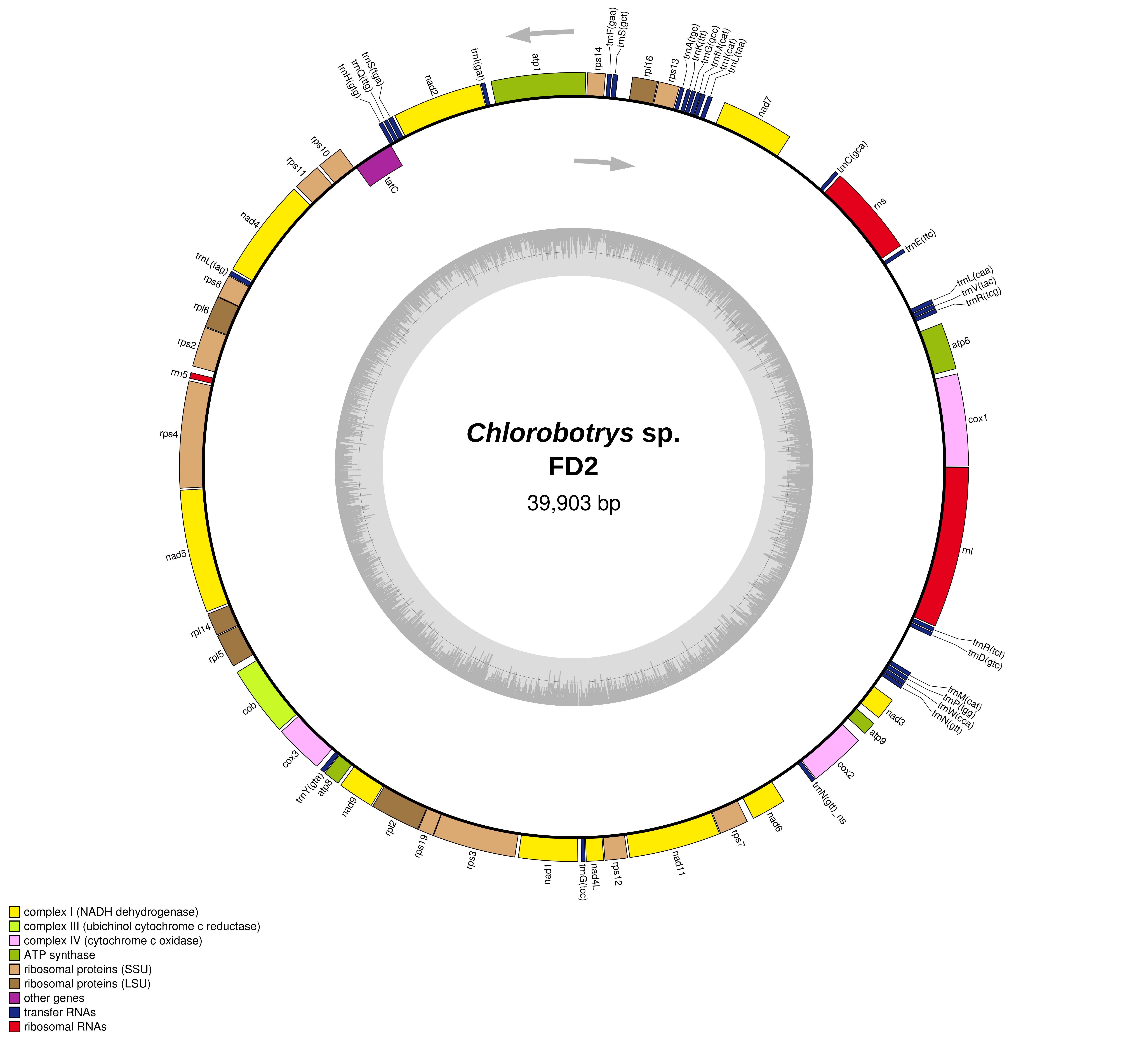
Chlorobotrys sp.
FD2
39,903 bp
atp1
trnI(gat)
nad2
trnS(tga)
trnQ(ttg)
trnH(gtg)
rps10
rps11
tatC
nad4
trnL(tag)
rps8
rpl6
rps2
rrn5
rps4
nad5
rpl14
rpl5
cob
cox3
trnY(gta)
atp8
nad9
rpl2
rps19
rps3
nad1
trnG(tcc)
nad4L
rps12
nad11
rps7
nad6
trnN(gtt)_ns
cox2
atp9
nad3
trnM(cat)
trnP(tgg)
trnW(cca)
trnN(gtt)
trnR(tct)
trnD(gtc)
rnl
cox1
atp6
trnL(caa)
trnV(tac)
trnR(tcg)
trnE(ttc)
rns
trnC(gca)
nad7
trnA(tgc)
trnK(ttt)
trnG(gcc)
trnfM(cat)
trnI(cat)
trnL(taa)
rps13
rpl16
rps14
trnF(gaa)
trnS(gct)
complex I (NADH dehydrogenase)
complex III (ubichinol cytochrome c reductase)
complex IV (cytochrome c oxidase)
ATP synthase
ribosomal proteins (SSU)
ribosomal proteins (LSU)
other genes
transfer RNAs
ribosomal RNAs

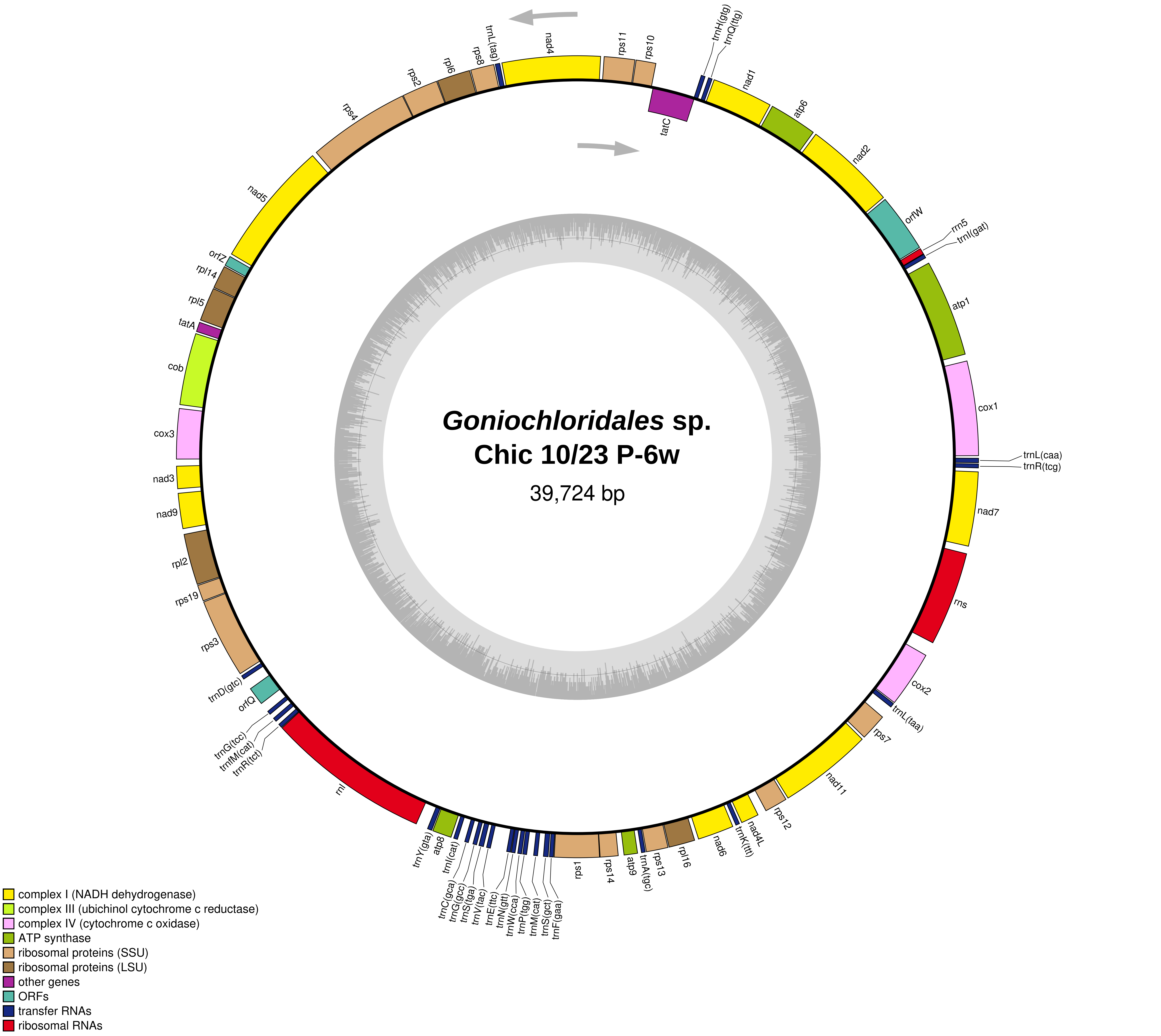

Goniochloridales sp.
Chic 10/23 P-6w
39,724 bp
nad4
rps11
rps10
trnH(gtg)
trnQ(ttg)
nad1
atp6
nad2
orfW
rrn5
trnI(gat)
atp1
cox1
trnL(caa)
trnR(tcg)
nad7
rns
cox2
trnL(taa)
rps7
nad11
rps12
nad4L
trnK(ttt)
nad6
rpl16
rps13
trnA(tgc)
atp9
rps14
rps1
trnF(gaa)
trnS(gct)
trnM(cat)
trnP(tgg)
trnW(cca)
trnN(gtt)
trnE(ttc)
trnV(tac)
trnS(tga)
trnG(gcc)
trnC(gca)
trnI(cat)
atp8
trnY(gta)
rnl
trnR(tct)
trnfM(cat)
trnG(tcc)
orfQ
trnD(gtc)
rps3
rps19
rpl2
nad9
nad3
cox3
cob
tatA
rpl5
rpl14
orfZ
nad5
rps4
rps2
rpl6
rps8
trnL(tag)
tatC
complex I (NADH dehydrogenase)
complex III (ubichinol cytochrome c reductase)
complex IV (cytochrome c oxidase)
ATP synthase
ribosomal proteins (SSU)
ribosomal proteins (LSU)
other genes
ORFs
transfer RNAs
ribosomal RNAs

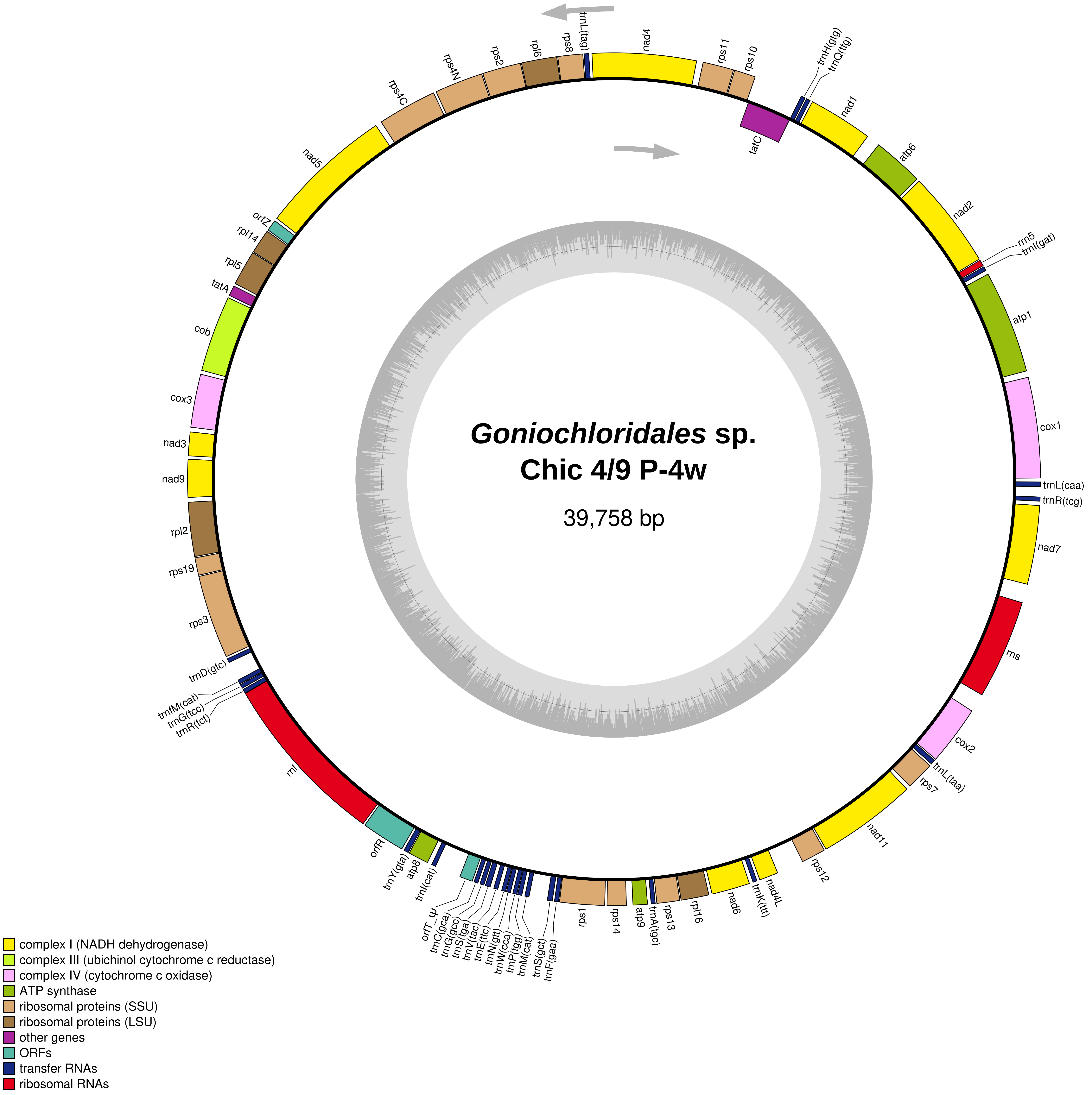
Goniochloridales sp.
Chic 4/9 P-4w
39,758 bp
trnL(tag)
nad4
rps11
rps10
trnH(gtg)
trnQ(ttg)
nad1
tatC
atp6
nad2
rrn5
trnI(gat)
atp1
cox1
trnL(caa)
trnR(tcg)
nad7
rns
cox2
trnL(taa)
rps7
nad11
rps12
nad4L
trnK(ttt)
nad6
rpl16
rps13
trnA(tgc)
atp9
rps14
rps1
trnF(gaa)
trnS(gct)
trnM(cat)
trnP(tgg)
trnW(cca)
trnN(gtt)
trnE(ttc)
trnV(tac)
trnS(tga)
trnG(gcc)
trnC(gca)
ψ
orfT
trnI(cat)
atp8
trnY(gta)
orfR
rnl
trnfM(cat)
trnG(tcc)
trnR(tct)
trnD(gtc)
rps3
rps19
rpl2
nad9
nad3
cox3
cob
tatA
rpl5
rpl14
orfZ
nad5
rps4C
rps4N
rps2
rpl6
rps8
complex I (NADH dehydrogenase)
complex III (ubichinol cytochrome c reductase)
complex IV (cytochrome c oxidase)
ATP synthase
ribosomal proteins (SSU)
ribosomal proteins (LSU)
other genes
ORFs
transfer RNAs
ribosomal RNAs

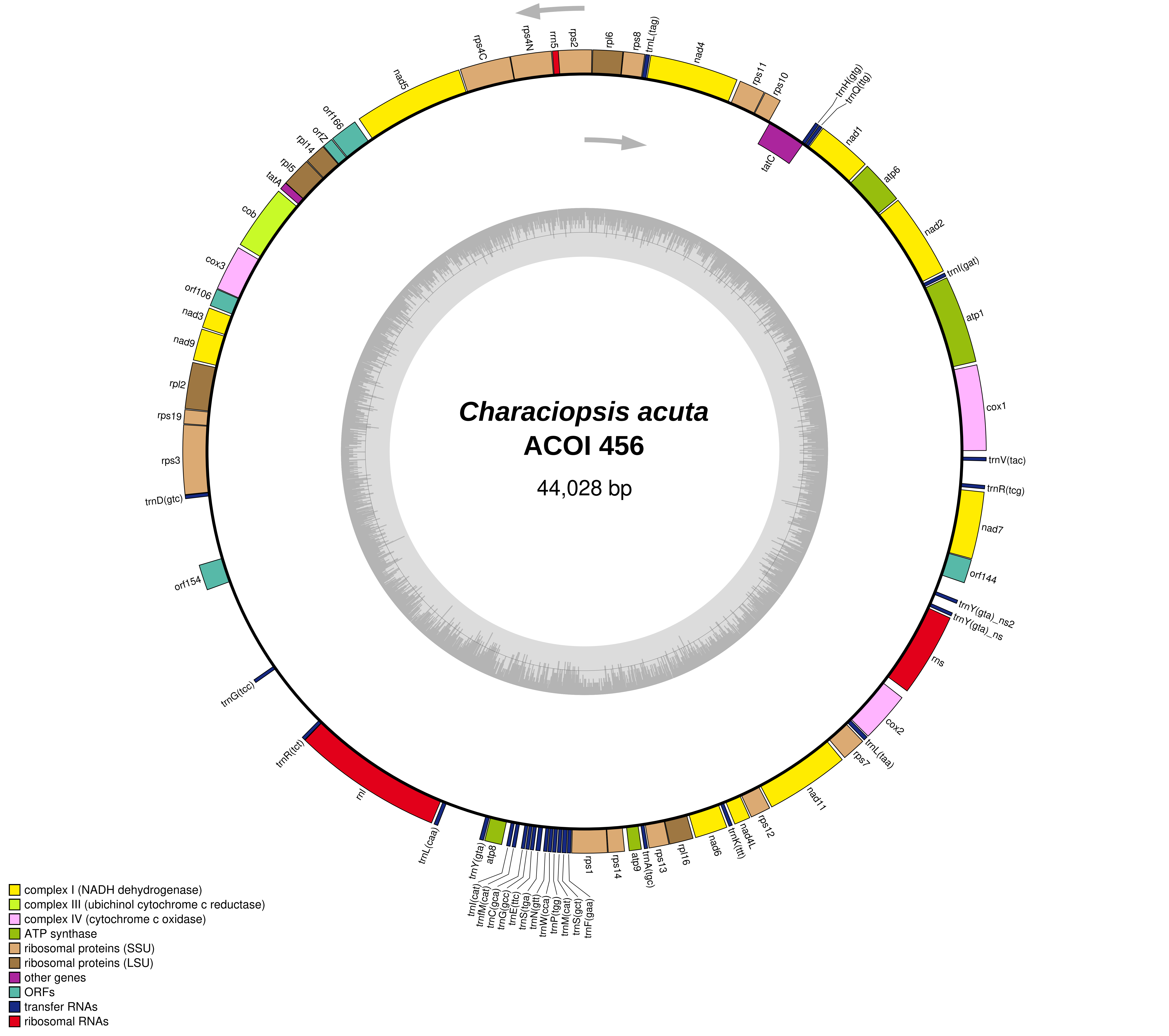

Characiopsis acuta
ACOI 456
44,028 bp
nad5
rps4C
rps4N
rrn5
rps2
rpl6
rps8
trnL(tag)
nad4
rps11
rps10
trnH(gtg)
trnQ(ttg)
tatC
nad1
atp6
nad2
trnI(gat)
atp1
cox1
trnV(tac)
trnR(tcg)
nad7
orf144
trnY(gta)_ns2
trnY(gta)_ns
rns
cox2
trnL(taa)
rps7
nad11
rps12
nad4L
trnK(ttt)
nad6
rpl16
rps13
trnA(tgc)
atp9
rps14
rps1
trnF(gaa)
trnS(gct)
trnM(cat)
trnP(tgg)
trnW(cca)
trnN(gtt)
trnS(tga)
trnE(ttc)
trnG(gcc)
trnC(gca)
trnfM(cat)
trnI(cat)
atp8
trnY(gta)
trnL(caa)
rnl
trnR(tct)
trnG(tcc)
orf154
trnD(gtc)
rps3
rps19
rpl2
nad9
nad3
orf106
cox3
cob
tatA
rpl5
rpl14
orfZ
orf166
complex I (NADH dehydrogenase)
complex III (ubichinol cytochrome c reductase)
complex IV (cytochrome c oxidase)
ATP synthase
ribosomal proteins (SSU)
ribosomal proteins (LSU)
other genes
ORFs
transfer RNAs
ribosomal RNAs

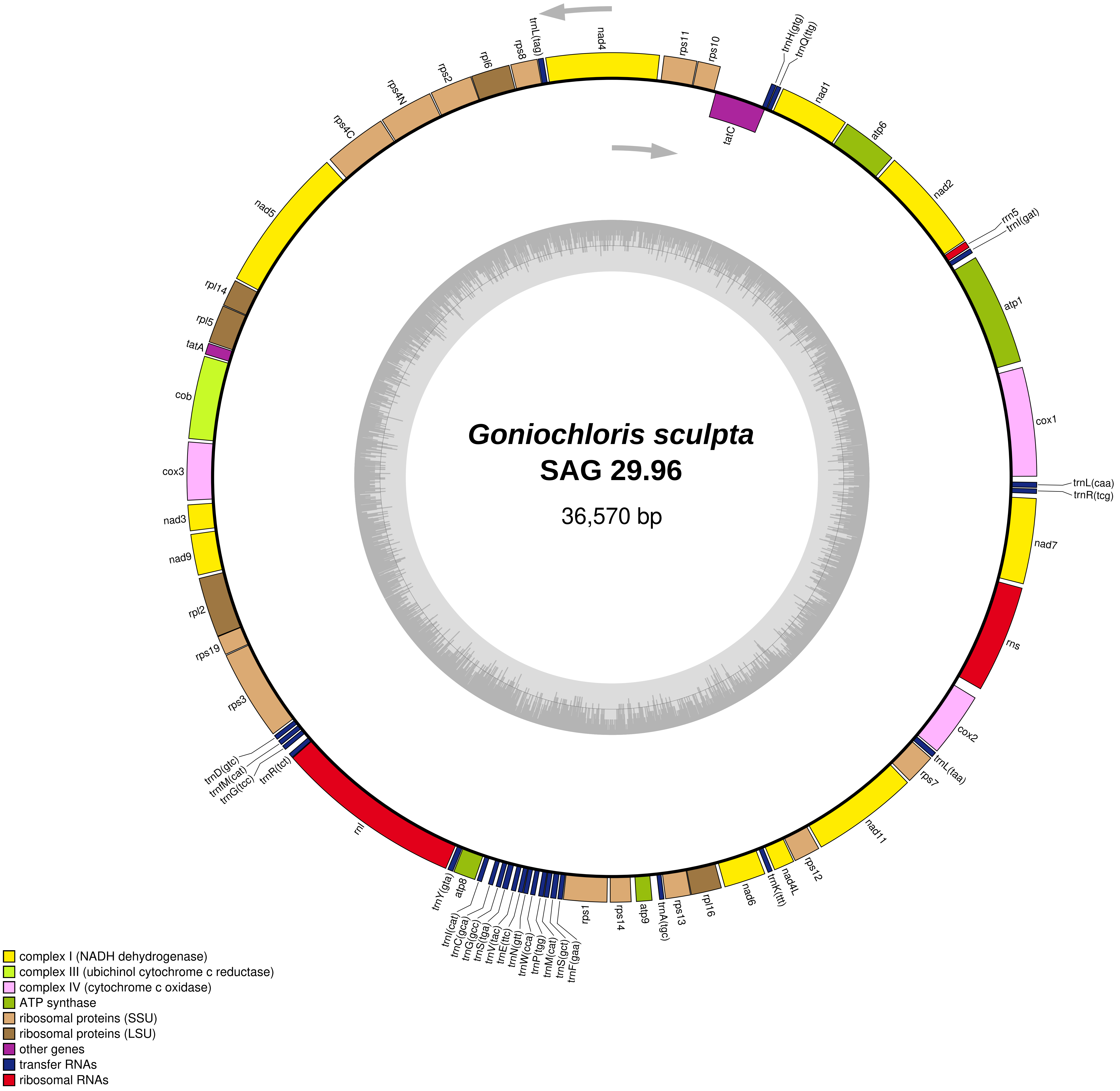

Goniochloris sculpta
SAG 29.96
36,570 bp
trnL(tag)
nad4
rps11
rps10
tatC
trnH(gtg)
trnQ(ttg)
nad1
atp6
nad2
rrn5
trnI(gat)
atp1
cox1
trnL(caa)
trnR(tcg)
nad7
rns
cox2
trnL(taa)
rps7
nad11
rps12
nad4L
trnK(ttt)
nad6
rpl16
rps13
trnA(tgc)
atp9
rps14
rps1
trnF(gaa)
trnS(gct)
trnM(cat)
trnP(tgg)
trnW(cca)
trnN(gtt)
trnE(ttc)
trnV(tac)
trnS(tga)
trnG(gcc)
trnC(gca)
trnI(cat)
atp8
trnY(gta)
rnl
trnR(tct)
trnG(tcc)
trnfM(cat)
trnD(gtc)
rps3
rps19
rpl2
nad9
nad3
cox3
cob
tatA
rpl5
rpl14
nad5
rps4C
rps4N
rps2
rpl6
rps8
complex I (NADH dehydrogenase)
complex III (ubichinol cytochrome c reductase)
complex IV (cytochrome c oxidase)
ATP synthase
ribosomal proteins (SSU)
ribosomal proteins (LSU)
other genes
transfer RNAs
ribosomal RNAs

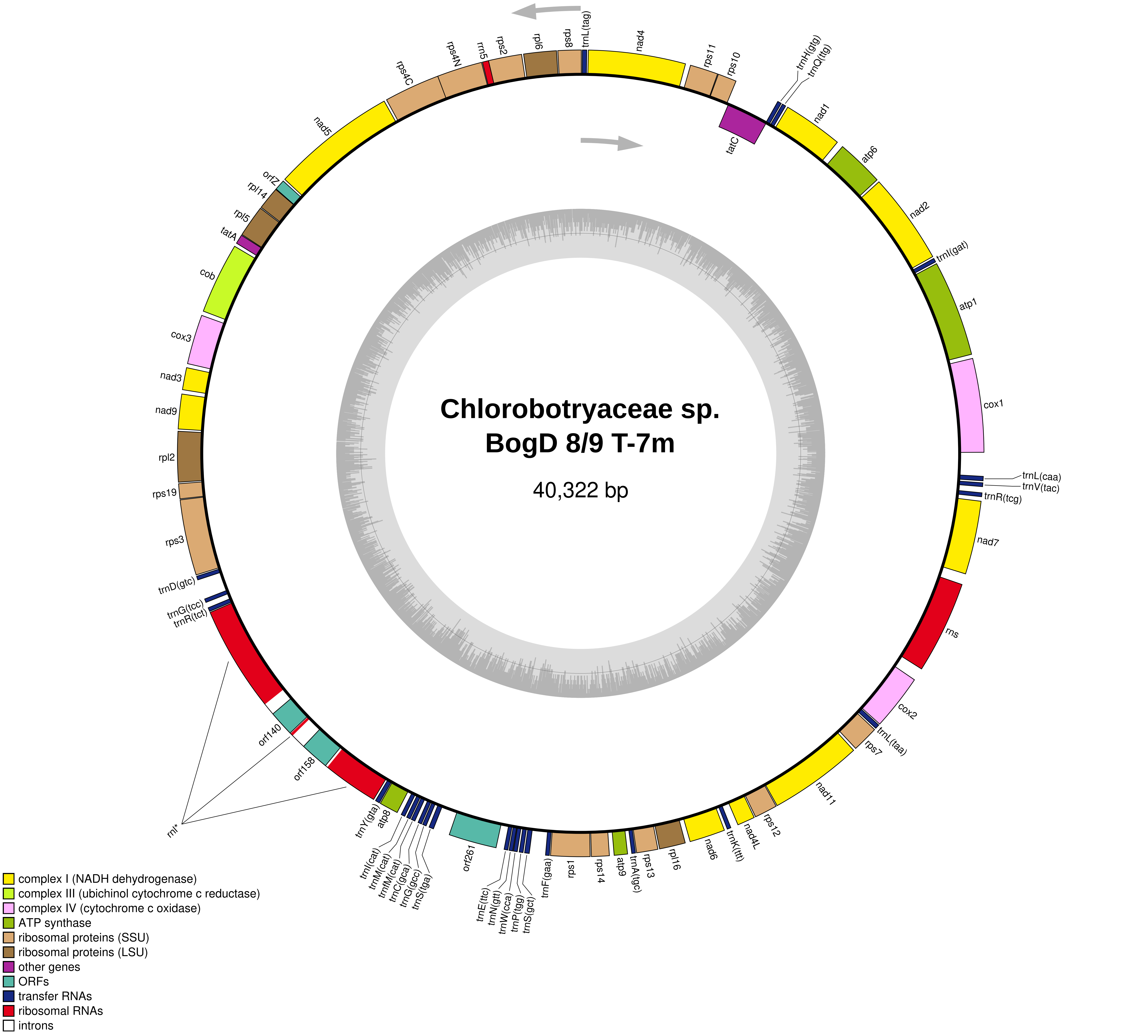
Chlorobotryaceae sp.
BogD 8/9 T-7m
40,322 bp
trnL(tag)
nad4
rps11
rps10
trnH(gtg)
trnQ(ttg)
tatC
nad1
atp6
nad2
trnI(gat)
atp1
cox1
trnL(caa)
trnV(tac)
trnR(tcg)
nad7
rns
cox2
trnL(taa)
rps7
nad11
rps12
nad4L
trnK(ttt)
nad6
rpl16
rps13
trnA(tgc)
atp9
rps14
rps1
trnF(gaa)
trnS(gct)
trnP(tgg)
trnW(cca)
trnN(gtt)
trnE(ttc)
orf261
trnS(tga)
trnG(gcc)
trnC(gca)
trnfM(cat)
trnM(cat)
trnI(cat)
atp8
trnY(gta)
rnl*
orf158
orf140
trnR(tct)
trnG(tcc)
trnD(gtc)
rps3
rps19
rpl2
nad9
nad3
cox3
cob
tatA
rpl5
rpl14
orfZ
nad5
rps4C
rps4N
rrn5
rps2
rpl6
rps8
complex I (NADH dehydrogenase)
complex III (ubichinol cytochrome c reductase)
complex IV (cytochrome c oxidase)
ATP synthase
ribosomal proteins (SSU)
ribosomal proteins (LSU)
other genes
ORFs
transfer RNAs
ribosomal RNAs
introns

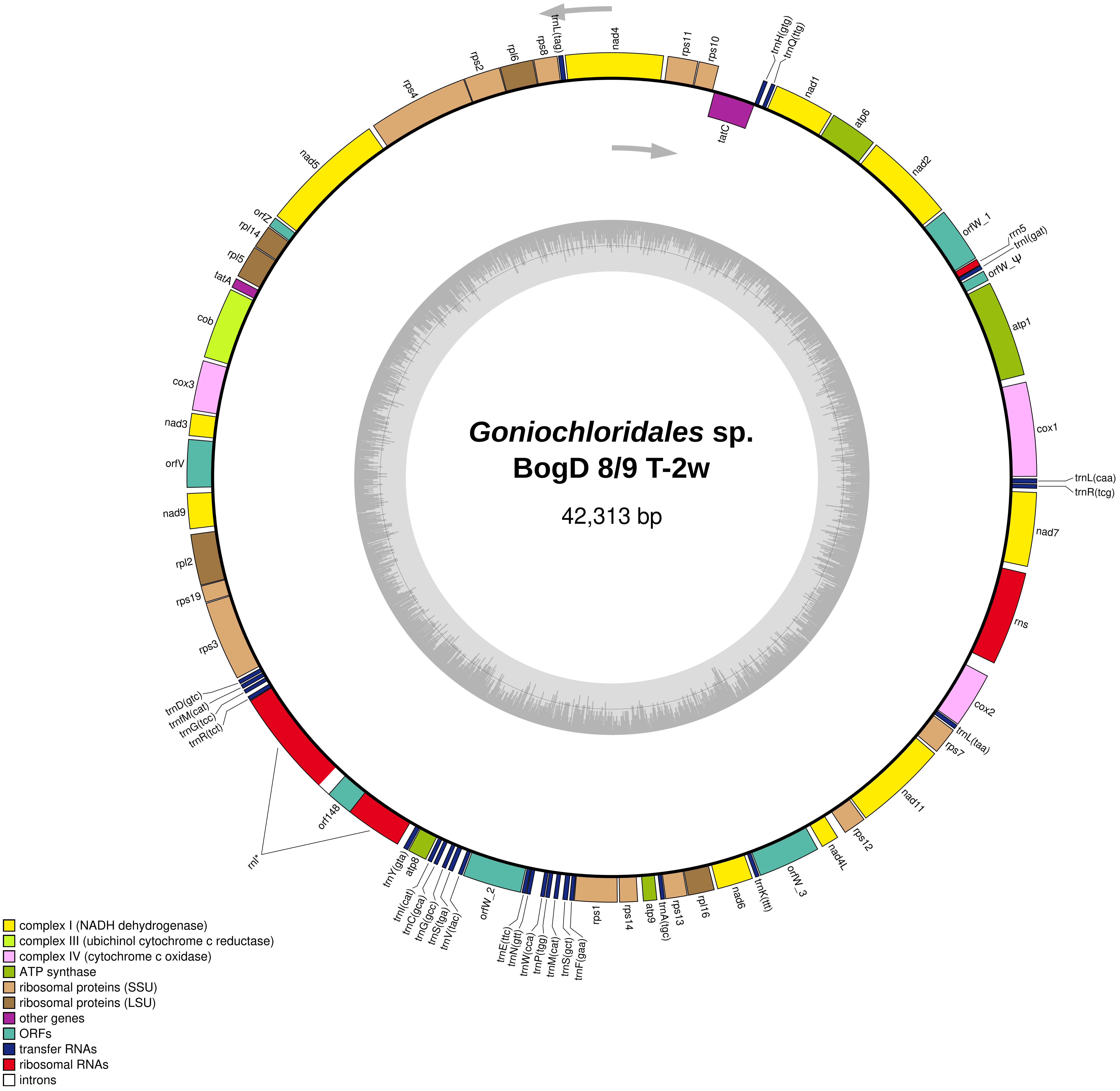

Goniochloridales sp.
BogD 8/9 T-2w
42,313 bp
nad4
rps11
rps10
trnH(gtg)
trnQ(ttg)
nad1
atp6
nad2
orfW_1
rrn5
trnI(gat)
orfW_Ψ
atp1
cox1
trnL(caa)
trnR(tcg)
nad7
rns
cox2
trnL(taa)
rps7
nad11
rps12
nad4L
orfW_3
trnK(ttt)
nad6
rpl16
rps13
trnA(tgc)
atp9
rps14
rps1
trnF(gaa)
trnS(gct)
trnM(cat)
trnP(tgg)
trnW(cca)
trnN(gtt)
trnE(ttc)
orfW_2
trnV(tac)
trnS(tga)
trnG(gcc)
trnC(gca)
trnI(cat)
atp8
trnY(gta)
rnl*
orf148
trnR(tct)
trnG(tcc)
trnfM(cat)
trnD(gtc)
rps3
rps19
rpl2
nad9
orfV
nad3
cox3
cob
tatA
rpl5
rpl14
orfZ
nad5
rps4
rps2
rpl6
rps8
trnL(tag)
tatC
complex I (NADH dehydrogenase)
complex III (ubichinol cytochrome c reductase)
complex IV (cytochrome c oxidase)
ATP synthase
ribosomal proteins (SSU)
ribosomal proteins (LSU)
other genes
ORFs
transfer RNAs
ribosomal RNAs
introns

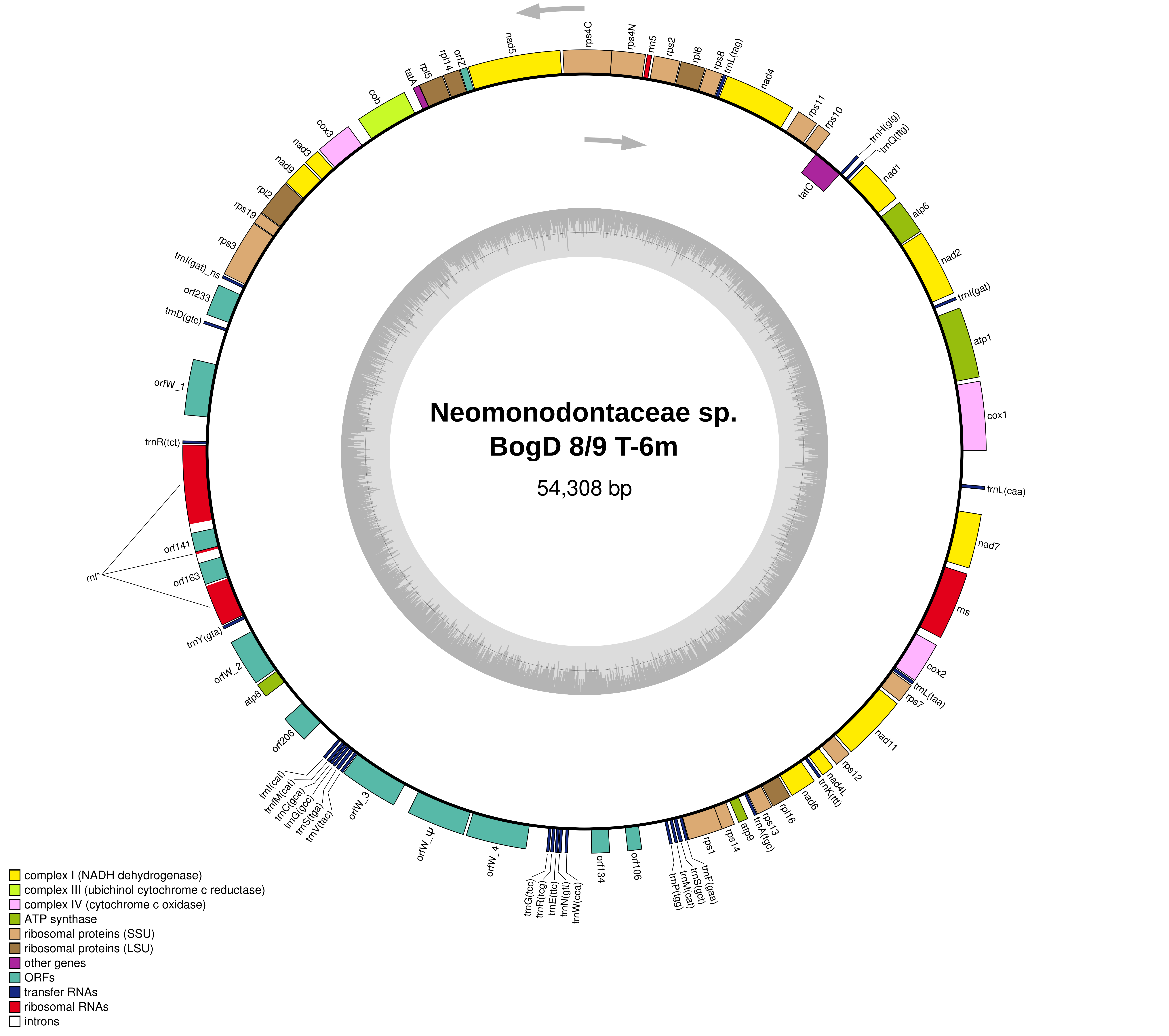

Neomonodontaceae sp.
BogD 8/9 T-6m
54,308 bp
nad5
rps4C
rps4N
rrn5
rps2
rpl6
rps8
trnL(tag)
nad4
rps11
rps10
trnH(gtg)
trnQ(ttg)
tatC
nad1
atp6
nad2
trnI(gat)
atp1
cox1
trnL(caa)
nad7
rns
cox2
trnL(taa)
rps7
nad11
rps12
nad4L
trnK(ttt)
nad6
rpl16
rps13
trnA(tgc)
atp9
rps14
rps1
trnF(gaa)
trnS(gct)
trnM(cat)
trnP(tgg)
orf106
orf134
trnW(cca)
trnN(gtt)
trnE(ttc)
trnR(tcg)
trnG(tcc)
orfW_4
orfW_3
trnV(tac)
trnS(tga)
trnG(gcc)
trnC(gca)
trnfM(cat)
trnI(cat)
orf206
atp8
orfW_2
trnY(gta)
orf163
orf141
rnl*
trnR(tct)
orfW_1
trnD(gtc)
orf233
trnI(gat)_ns
rps3
rps19
rpl2
nad9
nad3
cox3
cob
tatA
rpl5
rpl14
orfZ
complex I (NADH dehydrogenase)
complex III (ubichinol cytochrome c reductase)
complex IV (cytochrome c oxidase)
ATP synthase
ribosomal proteins (SSU)
ribosomal proteins (LSU)
other genes
ORFs
transfer RNAs
ribosomal RNAs
introns

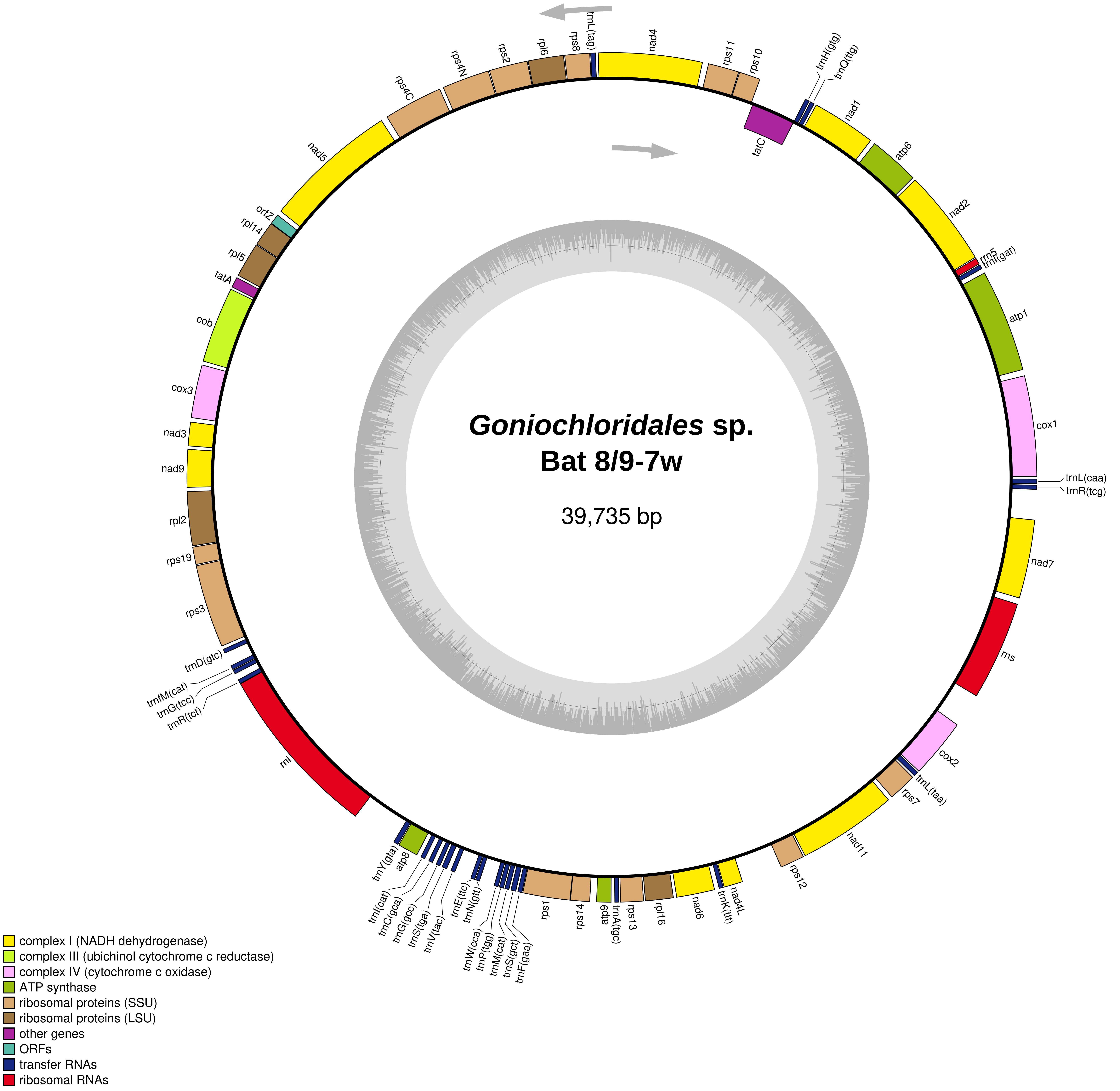
Goniochloridales sp.
Bat 8/9-7w
39,735 bp
trnL(tag)
nad4
rps11
rps10
trnH(gtg)
trnQ(ttg)
tatC
nad1
atp6
nad2
rrn5
trnI(gat)
atp1
cox1
trnL(caa)
trnR(tcg)
nad7
rns
cox2
trnL(taa)
rps7
nad11
rps12
nad4L
trnK(ttt)
nad6
rpl16
rps13
trnA(tgc)
atp9
rps14
rps1
trnF(gaa)
trnS(gct)
trnM(cat)
trnP(tgg)
trnW(cca)
trnN(gtt)
trnE(ttc)
trnV(tac)
trnS(tga)
trnG(gcc)
trnC(gca)
trnI(cat)
atp8
trnY(gta)
rnl
trnR(tct)
trnG(tcc)
trnfM(cat)
trnD(gtc)
rps3
rps19
rpl2
nad9
nad3
cox3
cob
tatA
rpl5
rpl14
orfZ
nad5
rps4C
rps4N
rps2
rpl6
rps8
complex I (NADH dehydrogenase)
complex III (ubichinol cytochrome c reductase)
complex IV (cytochrome c oxidase)
ATP synthase
ribosomal proteins (SSU)
ribosomal proteins (LSU)
other genes
ORFs
transfer RNAs
ribosomal RNAs
