## Supplementary Dataset S3 for "Insights into the phylogeny and enigmatic mitochondrial biology of eustigmatophyte algae from over 50 newly sequenced organellar genomes"

**Supplementary Dataset S3.** Transmembrane helices in the eustigmatophyte mitochondrial proteins encoded by the taxon-specific conserved genes *orfO* to *orfW*. The file presents outputs of TMHMM 2.0 (upper left part of each panel) and DeepTMHMM predictions (upper right and bottom part) for each protein. Sequences deemed to be pseudogenes were excluded from the analysis.

### OrfO

#### OrfO_1 (Orf99) *Microchloropsis gaditana* 3B

```
# WEBSEQUENCE Length: 99
# WEBSEQUENCE Number of predicted TMHs:  2
# WEBSEQUENCE Exp number of AAs in TMHs: 40.26841
# WEBSEQUENCE Exp number, first 60 AAs:  20.62754
# WEBSEQUENCE Total prob of N-in:        0.56293
# WEBSEQUENCE POSSIBLE N-term signal sequence
WEBSEQUENCE     TMHMM2.0        outside      1     3
WEBSEQUENCE     TMHMM2.0        TMhelix      4    26
WEBSEQUENCE     TMHMM2.0        inside      27    68
WEBSEQUENCE     TMHMM2.0        TMhelix     69    88
WEBSEQUENCE     TMHMM2.0        outside     89    99
```

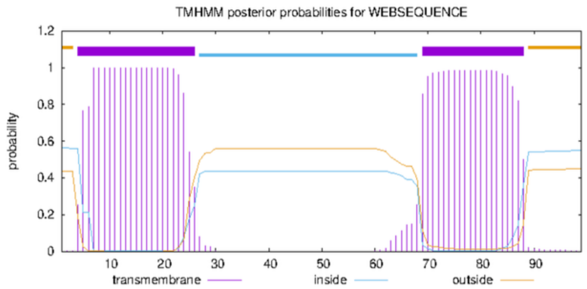


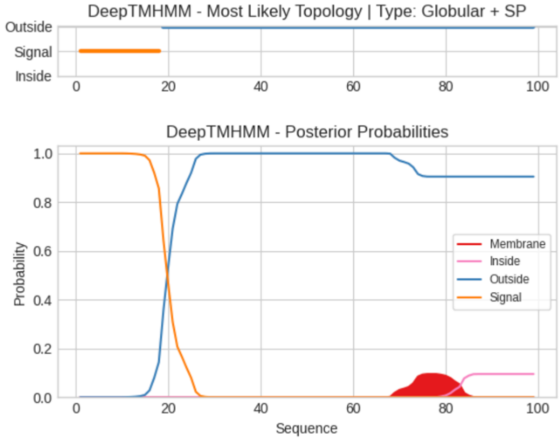


**Predicted Topologies**

```
>Sequence | SP
MKKTYIVIAFLLLLLGEAYYFFVLLHEYHTELLSLIEEVQKNEAEILTRLDQLQQKTHKHAQYWTYKELIGNGAYIVIMVVCLNYSYTKDGNFYFGWFG
SSSSSSSSSSSSSSSSSSOOOOOOOOOOOOOOOOOOOOOOOOOOOOOOOOOOOOOOOOOOOOOOOOOOOOOOOOOOOOOOOOOOOOOOOOOOOOOOOOO

##gff-version 3
# Sequence Length: 99
# Sequence Number of predicted TMRs: 0
Sequence signal 1 18
Sequence outside 19 99
```

#### OrfO_2 (Orf93) *Microchloropsis gaditana* 3B

```
# WEBSEQUENCE Length: 93
# WEBSEQUENCE Number of predicted TMHs:  2
# WEBSEQUENCE Exp number of AAs in TMHs: 44.57682
# WEBSEQUENCE Exp number, first 60 AAs:  22.31544
# WEBSEQUENCE Total prob of N-in:        0.98036
# WEBSEQUENCE POSSIBLE N-term signal sequence
WEBSEQUENCE     TMHMM2.0        inside       1     6
WEBSEQUENCE     TMHMM2.0        TMhelix      7    29
WEBSEQUENCE     TMHMM2.0        outside     30    69
WEBSEQUENCE     TMHMM2.0        TMhelix     70    92
WEBSEQUENCE     TMHMM2.0        inside      93    93
```

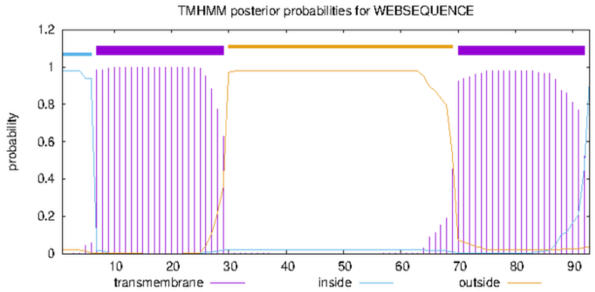


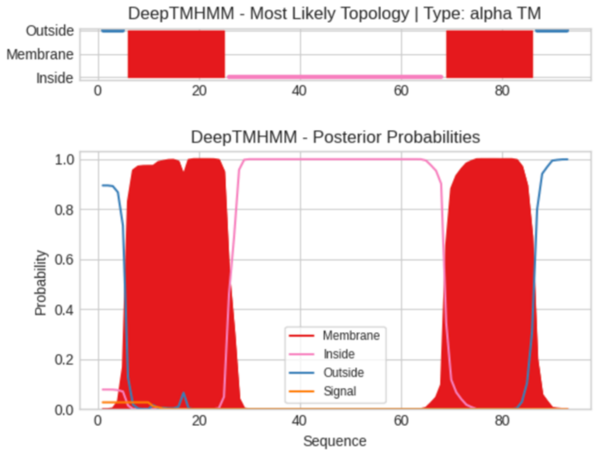


**Predicted Topologies**

```
>Sequence | TM
MRGTDLIIAFLLVWLVESICIFVLLQIYNPEMIGLLVELKNNEAEIAQICKALEKDTAVYTTETTLAQFVTYTSILGAIHGFYTGLASLSIFQ
OOOOOMMMMMMMMMMMMMMMMMMMMIIIIIIIIIIIIIIIIIIIIIIIIIIIIIIIIIIIIIIIIIIIMMMMMMMMMMMMMMMMMMOOOOOOO

##gff-version 3
# Sequence Length: 93
# Sequence Number of predicted TMRs: 2
Sequence outside 1 5
Sequence TMhelix 6 25
Sequence inside 26 68
Sequence TMhelix 69 86
Sequence outside 87 93
```

#### OrfO_1 (Orf99) *Microchloropsis salina* CCMP1776

```
# WEBSEQUENCE Length: 99
# WEBSEQUENCE Number of predicted TMHs:  2
# WEBSEQUENCE Exp number of AAs in TMHs: 40.35676
# WEBSEQUENCE Exp number, first 60 AAs:  20.95247
# WEBSEQUENCE Total prob of N-in:        0.29846
# WEBSEQUENCE POSSIBLE N-term signal sequence
WEBSEQUENCE     TMHMM2.0        outside      1     3
WEBSEQUENCE     TMHMM2.0        TMhelix      4    26
WEBSEQUENCE     TMHMM2.0        inside      27    68
WEBSEQUENCE     TMHMM2.0        TMhelix     69    88
WEBSEQUENCE     TMHMM2.0        outside     89    99
```

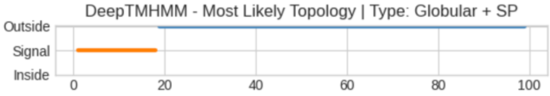


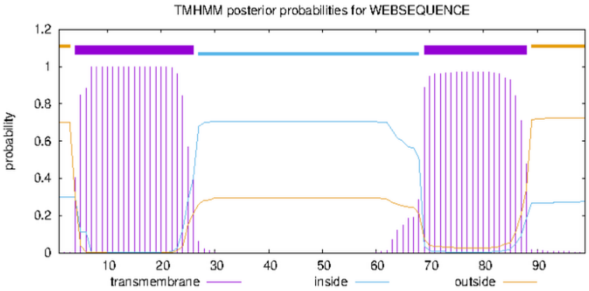


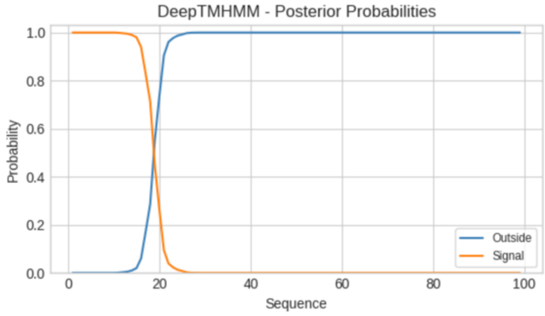


**Predicted Topologies**

```
>Sequence | SP
MKKTYIVIAFLLLLLGEAYYFFVLLHEYHTELLSLIEEVQKNEAEILTRLDQLQQKTHKRAQALTYKELIGNGAYIVIMVVSLNYSYTKDGNFYFGWFG
SSSSSSSSSSSSSSSSSSOOOOOOOOOOOOOOOOOOOOOOOOOOOOOOOOOOOOOOOOOOOOOOOOOOOOOOOOOOOOOOOOOOOOOOOOOOOOOOOOOOO

##gff-version 3
# Sequence Length: 99
# Sequence Number of predicted TMRs: 0
Sequence signal 1 18
Sequence outside 19 99
```

#### OrfO_2 (Orf93) *Microchloropsis salina* CCMP1776

```
# WEBSEQUENCE Length: 93
# WEBSEQUENCE Number of predicted TMHs:  2
# WEBSEQUENCE Exp number of AAs in TMHs: 43.67062
# WEBSEQUENCE Exp number, first 60 AAs:  22.28305
# WEBSEQUENCE Total prob of N-in:        0.96575
# WEBSEQUENCE POSSIBLE N-term signal sequence
WEBSEQUENCE     TMHMM2.0        inside       1     6
WEBSEQUENCE     TMHMM2.0        TMhelix      7    29
WEBSEQUENCE     TMHMM2.0        outside     30    69
WEBSEQUENCE     TMHMM2.0        TMhelix     70    92
WEBSEQUENCE     TMHMM2.0        inside      93    93
```

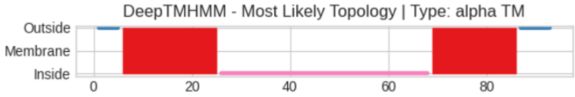


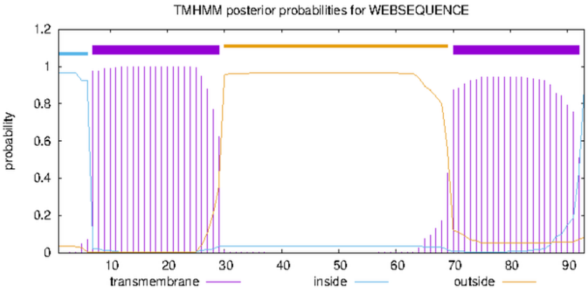


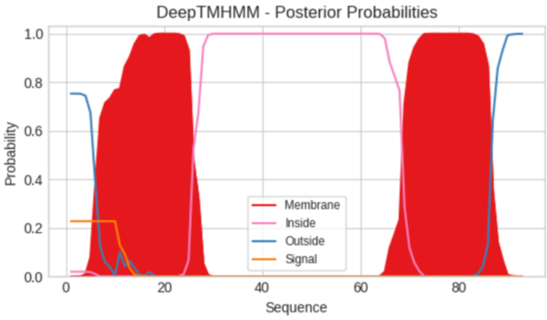


**Predicted Topologies**

```
>Sequence | TM
MRGTDLIITFLLVWLVESICIFVLLQIYNPEMIGLLVELKNNEAEIAQICKALEKDTAVYTTETTLAQFVSYASILGAIHGFSTGLANLSIFQ
OOOOOMMMMMMMMMMMMMMMMMMMMIIIIIIIIIIIIIIIIIIIIIIIIIIIIIIIIIIIIIIIIIIIMMMMMMMMMMMMMMMMMMOOOOOOO

##gff-version 3
# Sequence Length: 93
# Sequence Number of predicted TMRs: 2
Sequence outside 1 5
Sequence TMhelix 6 25
Sequence inside 26 68
Sequence TMhelix 69 86
Sequence outside 87 93
```

### OrfP

#### OrfP (Orf119) *Microchloropsis salina* CCMP1776

```
# WEBSEQUENCE Length: 119
# WEBSEQUENCE Number of predicted TMHs:  3
# WEBSEQUENCE Exp number of AAs in TMHs: 59.15668
# WEBSEQUENCE Exp number, first 60 AAs:  40.83982
# WEBSEQUENCE Total prob of N-in:        0.08482
# WEBSEQUENCE POSSIBLE N-term signal sequence
WEBSEQUENCE     TMHMM2.0        outside      1     3
WEBSEQUENCE     TMHMM2.0        TMhelix      4    23
WEBSEQUENCE     TMHMM2.0        inside      24    34
WEBSEQUENCE     TMHMM2.0        TMhelix     35    57
WEBSEQUENCE     TMHMM2.0        outside     58    99
WEBSEQUENCE     TMHMM2.0        TMhelix    100   117
WEBSEQUENCE     TMHMM2.0        inside     118   119
```

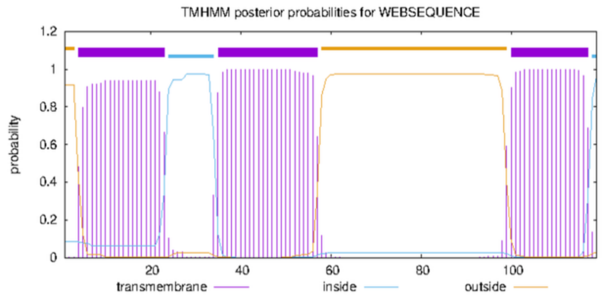


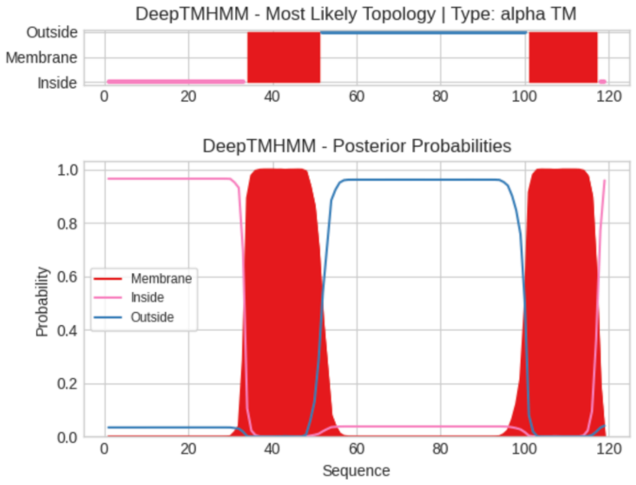


**Predicted Topologies**

```
>Sequence | TM
MKYGCTSFGFCYHFFLYIFKIFINQKIKLKIKDFFIIIFITLLIIVVGYCVDLSVIAHLDLLEQLQMARDNQLELLEAIDELEQSVEADCEEAQARKDTTIGIFLFM
LTAAFFVLGARA
IIIIIIIIIIIIIIIIIIIIIIIIIIIIIIIIIMMMMMMMMMMMMMMMMMMOOOOOOOOOOOOOOOOOOOOOOOOOOOOOOOOOOOOOOOOOOOOOOOOMMMMMMM
MMMMMMMMMMII

##gff-version 3
# Sequence Length: 119
# Sequence Number of predicted TMRs: 2
Sequence inside 1 33
Sequence TMhelix 34 51
Sequence outside 52 100
Sequence TMhelix 101 117
Sequence inside 118 119
```

#### OrfP (Orf53) *Microchloropsis gaditana* 3B

```
# WEBSEQUENCE Length: 53
# WEBSEQUENCE Number of predicted TMHs:  1
# WEBSEQUENCE Exp number of AAs in TMHs: 17.2497
# WEBSEQUENCE Exp number, first 60 AAs:  17.2497
# WEBSEQUENCE Total prob of N-in:        0.78908
# WEBSEQUENCE POSSIBLE N-term signal sequence
WEBSEQUENCE     TMHMM2.0        inside       1    34
WEBSEQUENCE     TMHMM2.0        TMhelix     35    52
WEBSEQUENCE     TMHMM2.0        outside     53    53
```

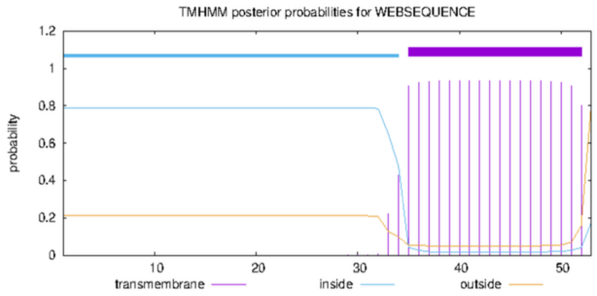


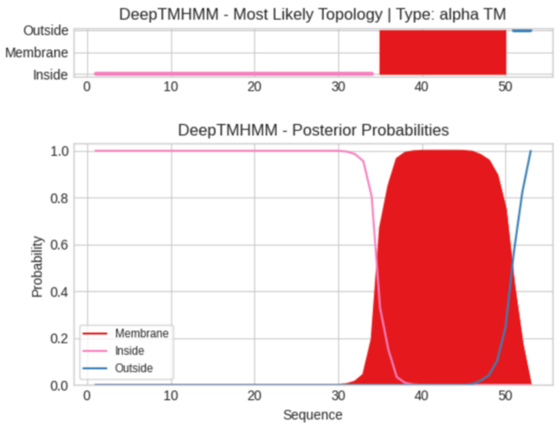


**Predicted Topologies**

```
>Sequence | TM
MARDNQLELLEVIDELEQSIEADCEEAQARKDTTIGIFLFMLTAVLGARAAFF
IIIIIIIIIIIIIIIIIIIIIIIIIIIIIIIIIIMMMMMMMMMMMMMMMMOOO

##gff-version 3
# Sequence Length: 53
# Sequence Number of predicted TMRs: 1
Sequence inside 1 34
Sequence TMhelix 35 50
Sequence outside 51 53
```

### OrfQ

#### OrfQ Goniochloridales sp. Chic 10/23 P-6w

```
# WEBSEQUENCE Length: 100
# WEBSEQUENCE Number of predicted TMHs:  1
# WEBSEQUENCE Exp number of AAs in TMHs: 31.74494
# WEBSEQUENCE Exp number, first 60 AAs:  21.59623
# WEBSEQUENCE Total prob of N-in:        0.92679
# WEBSEQUENCE POSSIBLE N-term signal sequence
WEBSEQUENCE     TMHMM2.0        inside       1     6
WEBSEQUENCE     TMHMM2.0        TMhelix      7    29
WEBSEQUENCE     TMHMM2.0        outside     30   100
```

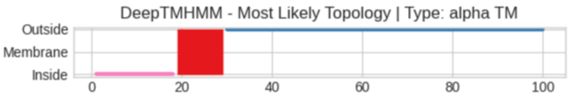


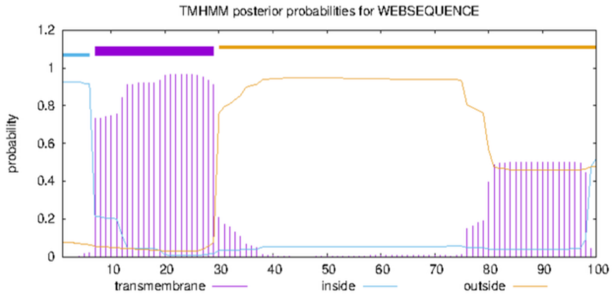


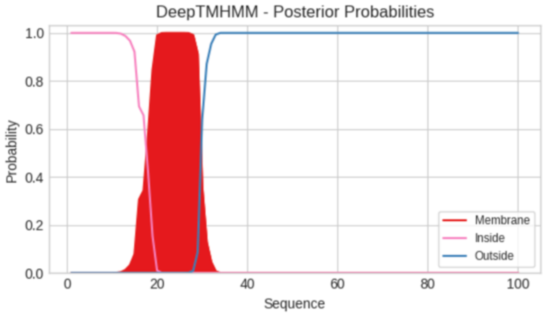


**Predicted Topologies**

```
>Sequence | TM
MRKFFHILLHNYLTNFKTGLAVIVLYFVFQCYYHPVVETYPDIELEKYVYDLLSQHRLNYELAIYSTLIGSSKRRLSLKLYVSFLYKRLLTGTYFAWFRR
IIIIIIIIIIIIIIIIIIMMMMMMMMMMMOOOOOOOOOOOOOOOOOOOOOOOOOOOOOOOOOOOOOOOOOOOOOOOOOOOOOOOOOOOOOOOOOOOOOOOO

##gff-version 3
# Sequence Length: 100
# Sequence Number of predicted TMRs: 1
Sequence inside 1 18
Sequence TMhelix 19 29
Sequence outside 30 100
```

#### OrfQ *Vacuoliviride crystalliferum* NIES-2860

```
# WEBSEQUENCE Length: 105
# WEBSEQUENCE Number of predicted TMHs:  1
# WEBSEQUENCE Exp number of AAs in TMHs: 31.13278
# WEBSEQUENCE Exp number, first 60 AAs:  21.77825
# WEBSEQUENCE Total prob of N-in:        0.98351
# WEBSEQUENCE POSSIBLE N-term signal sequence
WEBSEQUENCE     TMHMM2.0        inside       1    12
WEBSEQUENCE     TMHMM2.0        TMhelix     13    35
WEBSEQUENCE     TMHMM2.0        outside     36   105
```

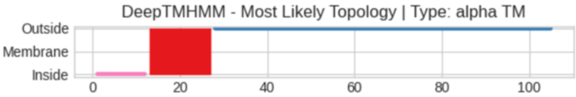


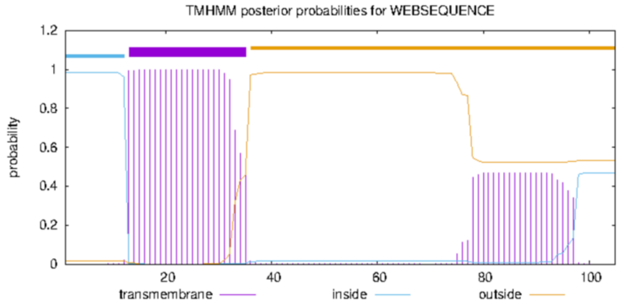


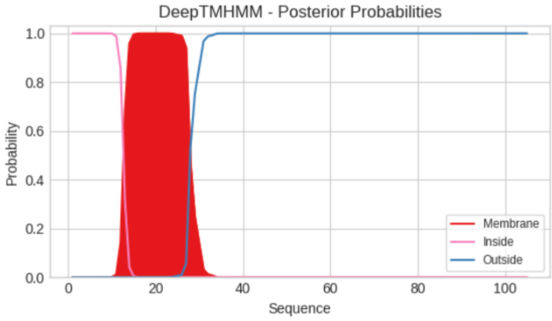


**Predicted Topologies**

```
>Sequence | TM
MKKIYMKNLNKDLLLLIIFIISFIYLFYYGYYINLTISDPLFETQNEVINLLYIKGLDNHIKVYQTIMNKDGRKNIKIFFVFLYKRILTGTLYRTSVKKNTYGNK
IIIIIIIIIIIIMMMMMMMMMMMMMMMOOOOOOOOOOOOOOOOOOOOOOOOOOOOOOOOOOOOOOOOOOOOOOOOOOOOOOOOOOOOOOOOOOOOOOOOOOOOOO

##gff-version 3
# Sequence Length: 105
# Sequence Number of predicted TMRs: 1
Sequence inside 1 12
Sequence TMhelix 13 27
Sequence outside 28 105
```

### OrfR

#### OrfR Goniochloridales sp. Chic 4/9 P-4w

```
# WEBSEQUENCE Length: 221
# WEBSEQUENCE Number of predicted TMHs:  2
# WEBSEQUENCE Exp number of AAs in TMHs: 48.70116
# WEBSEQUENCE Exp number, first 60 AAs:  34.0251
# WEBSEQUENCE Total prob of N-in:        0.59808
# WEBSEQUENCE POSSIBLE N-term signal sequence
WEBSEQUENCE     TMHMM2.0        inside       1     6
WEBSEQUENCE     TMHMM2.0        TMhelix      7    29
WEBSEQUENCE     TMHMM2.0        outside     30    56
WEBSEQUENCE     TMHMM2.0        TMhelix     57    79
WEBSEQUENCE     TMHMM2.0        inside      80   221
```

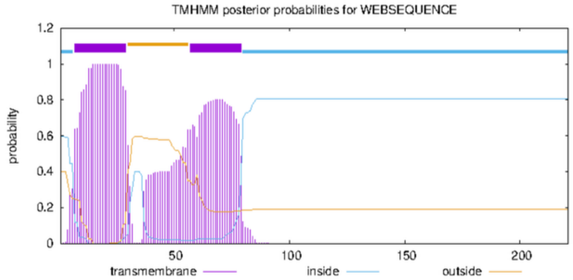


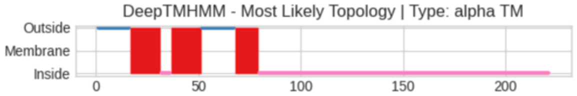


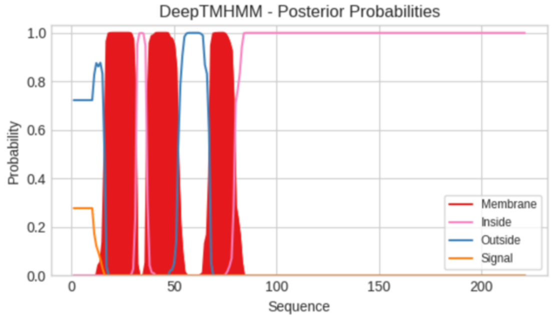


**Predicted Topologies**

```
>Sequence | TM
MEGILIGLFGGSIPPYLLFGIAMAIFLFIFLRKDHEVVFHFSFFWLGKLLFQSWTVSLFEISLSAPQLLGVVTVFYCFLKTAFCTERENGLLPQALDKTTSEIDLEC
SEKDIFEETVVYSDDKTQRLLPPKLGKIANEVLSLKRSIDISSRYYIYDDDVPETKVLSVKKLEVGPLEEQETLSKIPLGVVLGDPLGLSDEMLNIVNKLVNQTVNK
LFKYEKG
OOOOOOOOOOOOOOOOMMMMMMMMMMMMMMMIIIIIMMMMMMMMMMMMMMMOOOOOOOOOOOOOOOOMMMMMMMMMMMMIIIIIIIIIIIIIIIIIIIIIIIIIIII
IIIIIIIIIIIIIIIIIIIIIIIIIIIIIIIIIIIIIIIIIIIIIIIIIIIIIIIIIIIIIIIIIIIIIIIIIIIIIIIIIIIIIIIIIIIIIIIIIIIIIIIIIII
IIIIIII

##gff-version 3
# Sequence Length: 221
# Sequence Number of predicted TMRs: 3
Sequence outside 1 16
Sequence TMhelix 17 31
Sequence inside 32 36
Sequence TMhelix 37 51
Sequence outside 52 67
Sequence TMhelix 68 79
Sequence inside 80 221
```

#### OrfR *Trachydiscus minutus* CCALA 838

```
# WEBSEQUENCE Length: 221
# WEBSEQUENCE Number of predicted TMHs:  3
# WEBSEQUENCE Exp number of AAs in TMHs: 59.3358
# WEBSEQUENCE Exp number, first 60 AAs:  31.6892
# WEBSEQUENCE Total prob of N-in:        0.00286
# WEBSEQUENCE POSSIBLE N-term signal sequence
WEBSEQUENCE     TMHMM2.0        outside      1    23
WEBSEQUENCE     TMHMM2.0        TMhelix     24    43
WEBSEQUENCE     TMHMM2.0        inside      44    49
WEBSEQUENCE     TMHMM2.0        TMhelix     50    72
WEBSEQUENCE     TMHMM2.0        outside     73    81
WEBSEQUENCE     TMHMM2.0        TMhelix     82   104
WEBSEQUENCE     TMHMM2.0        inside     105   221
```

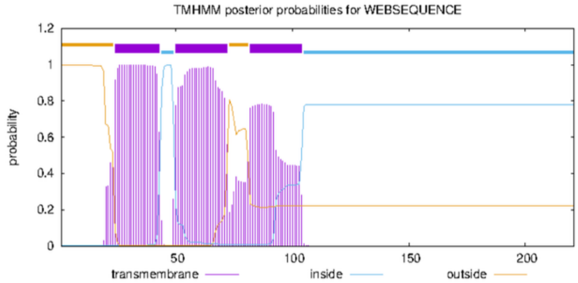


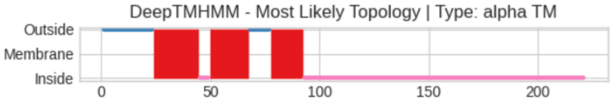


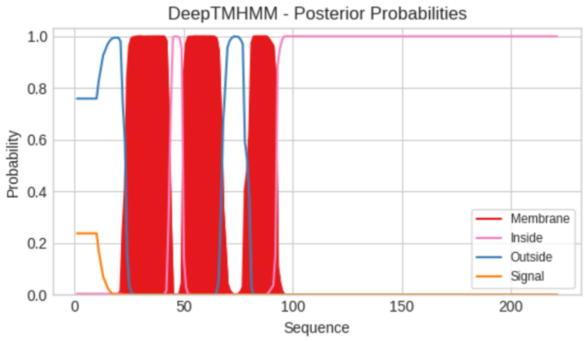


**Predicted Topologies**

```
>Sequence | TM
MLLSPLDQFRILPILPDNIGSFDILFTNSVLILLIALGSFLLCAKKKDELVFHLSFIWLGKLLFLAQSVTFFEVTLSAPQVLTLVVIFYSFLKATSCDPSYGLLPKT
SSRLKRPPSDGLLSDTSSICLMCPSDNSELGDSVPFNWSLGFTEDENLPGLNLDEDFISKSLPSYLAEAVSLPITRENLKTFNESIKRAQIGMVHDNYRILPAEDVA
KQAEATM
OOOOOOOOOOOOOOOOOOOOOOOMMMMMMMMMMMMMMMMMMMMMIIIIIMMMMMMMMMMMMMMMMMMOOOOOOOOOOMMMMMMMMMMMMMMMIIIIIIIIIIIIII
IIIIIIIIIIIIIIIIIIIIIIIIIIIIIIIIIIIIIIIIIIIIIIIIIIIIIIIIIIIIIIIIIIIIIIIIIIIIIIIIIIIIIIIIIIIIIIIIIIIIIIIIIII
IIIIIII

##gff-version 3
# Sequence Length: 221
# Sequence Number of predicted TMRs: 3
Sequence outside 1 23
Sequence TMhelix 24 44
Sequence inside 45 49
Sequence TMhelix 50 67
```

```
Sequence outside 68 77
Sequence TMhelix 78 92
Sequence inside 93 221
```

#### OrfR Goniochloridales sp. WTwin 8/9 T-7m6.8

```
# WEBSEQUENCE Length: 233
# WEBSEQUENCE Number of predicted TMHs:  3
# WEBSEQUENCE Exp number of AAs in TMHs: 59.32875
# WEBSEQUENCE Exp number, first 60 AAs:  38.85326
# WEBSEQUENCE Total prob of N-in:        0.00306
# WEBSEQUENCE POSSIBLE N-term signal sequence
WEBSEQUENCE     TMHMM2.0        outside      1    14
WEBSEQUENCE     TMHMM2.0        TMhelix     15    31
WEBSEQUENCE     TMHMM2.0        inside      32    37
WEBSEQUENCE     TMHMM2.0        TMhelix     38    55
WEBSEQUENCE     TMHMM2.0        outside     56    64
WEBSEQUENCE     TMHMM2.0        TMhelix     65    87
WEBSEQUENCE     TMHMM2.0        inside      88   233
```

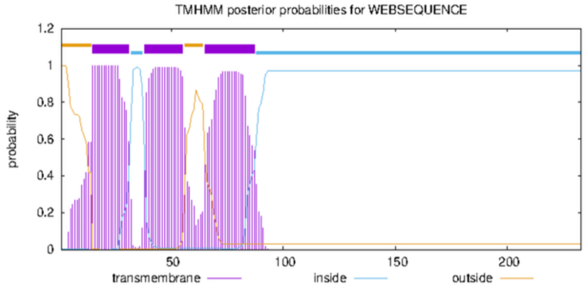


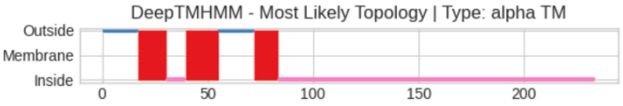


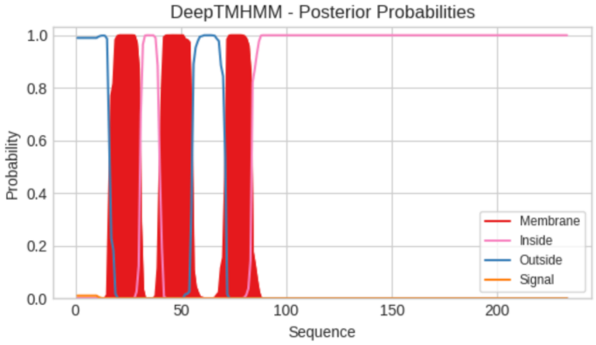


**Predicted Topologies**

```
>Sequence | TM
MDSVLGILFSGSTPHYLPLVIVLVCFLFLFSQRNSRRNYIAIFHLSFIWLGRLLYQSLGFDVFEITLSAPQLLFVVGLLYNFLKVAVYNAGKEKNSEGEKDMDLVYY
NQEDDLSVVDYDDCVCPCCNDFKVYVEEFYLSDEDDDCSCTACDITQHLYDLDSDTDTEEKLKMWNQNDVNESPEFLGLAMVKPVEEFDKQVGFVYKPEETLHDINQ
CILQGCRHCLQCLEILEKM
OOOOOOOOOOOOOOOOMMMMMMMMMMMMMMIIIIIIIIIMMMMMMMMMMMMMMMMOOOOOOOOOOOOOOOOMMMMMMMMMMMMIIIIIIIIIIIIIIIIIIIIIIII
IIIIIIIIIIIIIIIIIIIIIIIIIIIIIIIIIIIIIIIIIIIIIIIIIIIIIIIIIIIIIIIIIIIIIIIIIIIIIIIIIIIIIIIIIIIIIIIIIIIIIIIIIII
IIIIIIIIIIIIIIIIIII

##gff-version 3
# Sequence Length: 233
# Sequence Number of predicted TMRs: 3
Sequence outside 1 16
Sequence TMhelix 17 30
Sequence inside 31 39
Sequence TMhelix 40 55
Sequence outside 56 71
Sequence TMhelix 72 83
Sequence inside 84 233
```

### OrfS

#### OrfS *Liezensia polymorpha* SAG 2220

```
# WEBSEQUENCE Length: 148
# WEBSEQUENCE Number of predicted TMHs:  3
# WEBSEQUENCE Exp number of AAs in TMHs: 69.64024
# WEBSEQUENCE Exp number, first 60 AAs:  26.31666
# WEBSEQUENCE Total prob of N-in:        0.99264
# WEBSEQUENCE POSSIBLE N-term signal sequence
WEBSEQUENCE     TMHMM2.0        inside       1    11
WEBSEQUENCE     TMHMM2.0        TMhelix     12    34
WEBSEQUENCE     TMHMM2.0        outside     35    60
WEBSEQUENCE     TMHMM2.0        TMhelix     61    83
WEBSEQUENCE     TMHMM2.0        inside      84    89
WEBSEQUENCE     TMHMM2.0        TMhelix     90   112
WEBSEQUENCE     TMHMM2.0        outside    113   148
```

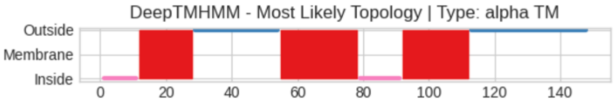


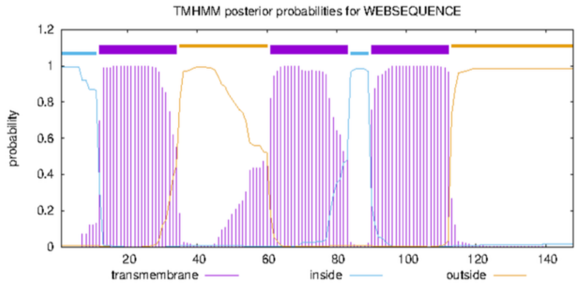


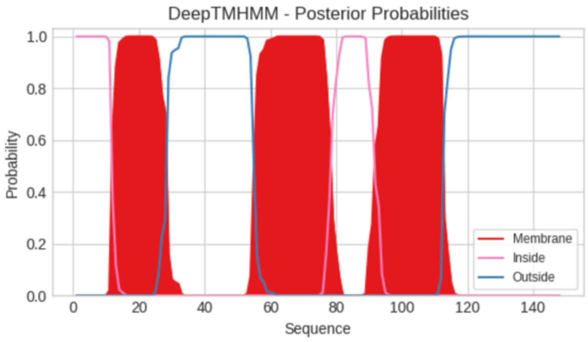


**Predicted Topologies**

```
>Sequence | TM
MKTMKYLQLNRLFLTIFLFLSVILTYYAHSFIGLPHIFSKEKITGIFIQQFYYGILLKICCFMAWYAILSPFLFWLYFYWISFKISITEFVFNIFLGLYWFVFYFIV
LAFLIYTDTMWGLLNILIGQVQENEFNSLNLLKKSLITGAP
IIIIIIIIIIIMMMMMMMMMMMMMMMMMOOOOOOOOOOOOOOOOOOOOOOOOOOMMMMMMMMMMMMMMMMMMMMMMMMIIIIIIIIIIIIIMMMMMMMMMMMMMMMM
MMMMMOOOOOOOOOOOOOOOOOOOOOOOOOOOOOOOOOOOOOOOOOOO

##gff-version 3
# Sequence Length: 148
# Sequence Number of predicted TMRs: 3
Sequence inside 1 11
Sequence TMhelix 12 28
Sequence outside 29 54
Sequence TMhelix 55 78
Sequence inside 79 91
Sequence TMhelix 92 112
Sequence outside 113 148
```

#### OrfS *Vacuoliviride crystalliferum* NIES-2860

```
# WEBSEQUENCE Length: 294
# WEBSEQUENCE Number of predicted TMHs:  3
# WEBSEQUENCE Exp number of AAs in TMHs: 66.5708800000001
# WEBSEQUENCE Exp number, first 60 AAs:  24.12767
# WEBSEQUENCE Total prob of N-in:        0.99803
# WEBSEQUENCE POSSIBLE N-term signal sequence
WEBSEQUENCE     TMHMM2.0        inside       1    12
WEBSEQUENCE     TMHMM2.0        TMhelix     13    33
WEBSEQUENCE     TMHMM2.0        outside     34    59
WEBSEQUENCE     TMHMM2.0        TMhelix     60    82
WEBSEQUENCE     TMHMM2.0        inside      83    94
WEBSEQUENCE     TMHMM2.0        TMhelix     95   117
WEBSEQUENCE     TMHMM2.0        outside    118   294
```

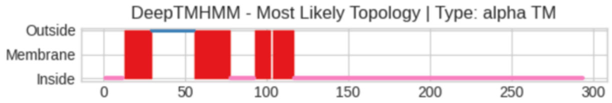


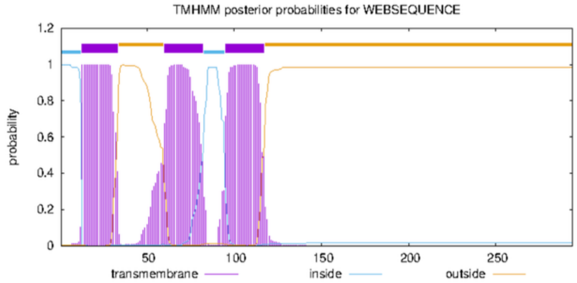


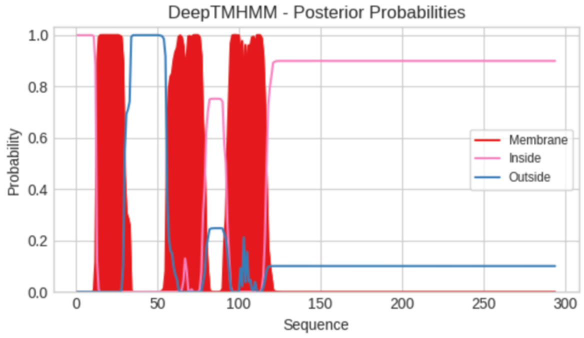


**Predicted Topologies**

```
>Sequence | TM
MTNIKIYYNRNKLFLVITLLMVFLSLYLVHSMIDKPHILSSEKIGTISYQFLLYGILIKFSYFLTFYSLLSSLWFWVYFYWNALKLDLTDVCWDFLIISLWFMFSFI
ILSAILFVDVTLGSLNILVAEINNKPFDILQYFFRSVAQQKHLETPCFKCDDHPPKELGFSGLQDNRLILLMNPAKGDPSNTWGKSELARVVGNDDGSVGACGNILG
GIDHQQKKIQISQDKADQDMFNKTSADCATFADKIKHSLASAARKGAQTGLNTLGGFAQSYKMDTICTNVDSDKIPTTKI
IIIIIIIIIIIIMMMMMMMMMMMMMMMMMOOOOOOOOOOOOOOOOOOOOOOOOOOMMMMMMMMMMMMMMMMMMMMMMIIIIIIIIIIIIIIIMMMMMMMMMMOMMMM
MMMMMMMMMIIIIIIIIIIIIIIIIIIIIIIIIIIIIIIIIIIIIIIIIIIIIIIIIIIIIIIIIIIIIIIIIIIIIIIIIIIIIIIIIIIIIIII
IIIIIIIIIIIIIIIIIIIIIIIIIIIIIIIIIIIIIIIIIIIIIIIIIIIIIIIIIIIIIIIIIIIIIIIIIIIIIIIIIIIIIIIIIIIIIIII

##gff-version 3
# Sequence Length: 294
# Sequence Number of predicted TMRs: 4
Sequence inside 1 12
Sequence TMhelix 13 29
Sequence outside 30 55
Sequence TMhelix 56 77
Sequence inside 78 92
```

```
Sequence TMhelix 93 102
Sequence outside 103 103
Sequence TMhelix 104 116
Sequence inside 117 294
```

### OrfS *Trachydiscus minutus* CCALA 838

```
# WEBSEQUENCE Length: 331
# WEBSEQUENCE Number of predicted TMHs:  3
# WEBSEQUENCE Exp number of AAs in TMHs: 67.0899600000002
# WEBSEQUENCE Exp number, first 60 AAs:  20.18043
# WEBSEQUENCE Total prob of N-in:        0.98468
# WEBSEQUENCE POSSIBLE N-term signal sequence
WEBSEQUENCE     TMHMM2.0        inside       1    27
WEBSEQUENCE     TMHMM2.0        TMhelix     28    47
WEBSEQUENCE     TMHMM2.0        outside     48    72
WEBSEQUENCE     TMHMM2.0        TMhelix     73    95
WEBSEQUENCE     TMHMM2.0        inside      96   107
WEBSEQUENCE     TMHMM2.0        TMhelix    108   130
WEBSEQUENCE     TMHMM2.0        outside    131   331
```

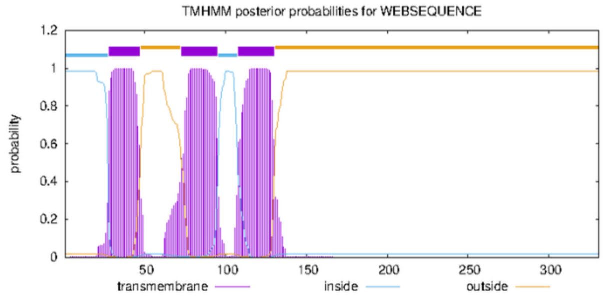


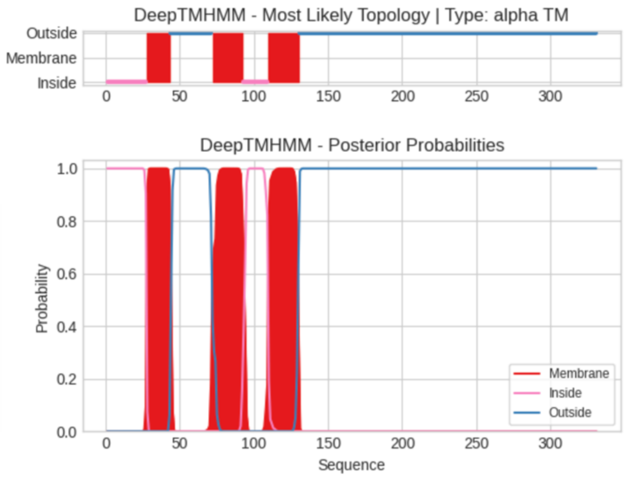


**Predicted Topologies**

```
>Sequence | TM
MSIAHTTNTGFFKRFWLNPLIFYKRNRVFLSLCLLIVALFLWYTHSITGKPFILSRENMESLGIQFFFYSLLIKISWGLTFYSIISSLFFWWLFYFKALKVDAGEYA
IDFILISLWVLFTFVILAAVFFFDLSIVSLKTVLYGLGYNDVDLWQSLTRILAADLHTFSTKDCLTCKDLSASSLDQLCLNKGLMQKEYSQLLFFSLDSPQIKYVVD
DFSNKQSSKFNPTSYPKMESTSEHTTFITAAFNKEGKIAACFDTTAVLEQFKPKPSEVLDNAGSFFSKILKPCTDKMLTQGLSTAQQGLVNAGLDVSCTANVPVDQK
PSSFAAERKG
IIIIIIIIIIIIIIIIIIIIIIIIIIIMMMMMMMMMMMMMMMMOOOOOOOOOOOOOOOOOOOOOOOOOOOOMMMMMMMMMMMMMMMMMMMMMIIIIIIIIIIIIIII
IIMMMMMMMMMMMMMMMMMMMMMOOOOOOOOOOOOOOOOOOOOOOOOOOOOOOOOOOOOOOOOOOOOOOOOOOOOOOOOOOOOOOOOOOOOOOOOOOOOOOOOOOOO
OOOOOOOOOOOOOOOOOOOOOOOOOOOOOOOOOOOOOOOOOOOOOOOOOOOOOOOOOOOOOOOOOOOOOOOOOOOOOOOOOOOOOOOOOOOOOOOOOOOOOOOOOOO
OOOOOOOOOO

##gff-version 3
# Sequence Length: 331
# Sequence Number of predicted TMRs: 3
Sequence inside 1 27
Sequence TMhelix 28 43
Sequence outside 44 71
Sequence TMhelix 72 92
Sequence inside 93 109
Sequence TMhelix 110 130
Sequence outside 131 331
```

### OrfT

#### OrfT (Orf222) *Trachydiscus minutus* CCALA 838

```
# WEBSEQUENCE Length: 222
# WEBSEQUENCE Number of predicted TMHs:  3
# WEBSEQUENCE Exp number of AAs in TMHs: 67.11701
# WEBSEQUENCE Exp number, first 60 AAs:  37.3243
# WEBSEQUENCE Total prob of N-in:        0.99693
# WEBSEQUENCE POSSIBLE N-term signal sequence
WEBSEQUENCE     TMHMM2.0        inside       1     6
WEBSEQUENCE     TMHMM2.0        TMhelix      7    29
WEBSEQUENCE     TMHMM2.0        outside     30    43
WEBSEQUENCE     TMHMM2.0        TMhelix     44    66
WEBSEQUENCE     TMHMM2.0        inside      67    86
WEBSEQUENCE     TMHMM2.0        TMhelix     87   109
WEBSEQUENCE     TMHMM2.0        outside    110   222
```

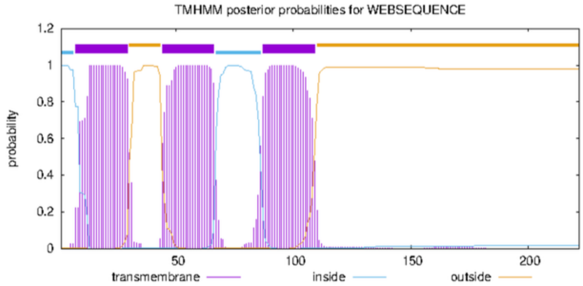


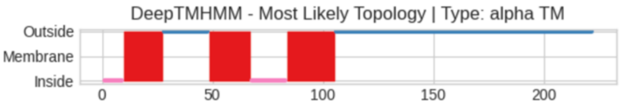


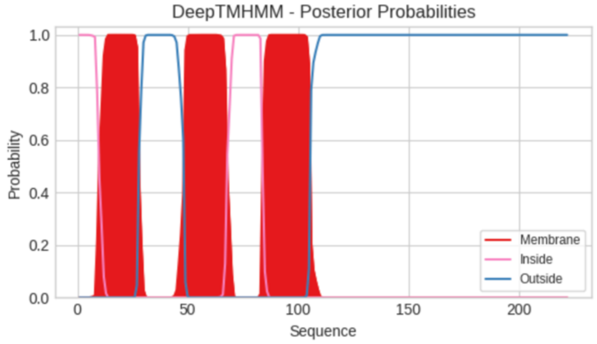


**Predicted Topologies**

```
>Sequence | TM
MYKNLFNKIYGPYFLMLLLLIAVLYILRITTDYNIVSLSSTEWLAIQQLILQGFFFFYAIFFLSFLYTTYKDYYKMIAVYNKGWILFVNFVFFNGFLFILVGILPSY
FFYNSAQLFSNIFLLNYSEFSQVPLAFDDFFQLLMEKKGENDLSTLPKSVGDHTLVIGTTLDGKGYLCINIKKAFNQCIDSQTSTGKMAEKGFNSVQRFFSLKDVVC
KVSTPVEK
IIIIIIIIIMMMMMMMMMMMMMMMMMMOOOOOOOOOOOOOOOOOOOOOMMMMMMMMMMMMMMMMMMMIIIIIIIIIIIIIIIIMMMMMMMMMMMMMMMMMMMMMMOO
OOOOOOOOOOOOOOOOOOOOOOOOOOOOOOOOOOOOOOOOOOOOOOOOOOOOOOOOOOOOOOOOOOOOOOOOOOOOOOOOOOOOOOOOOOOOOOOOOOOOOOOOOOO
OOOOOOOO

##gff-version 3
# Sequence Length: 222
# Sequence Number of predicted TMRs: 3
Sequence inside 1 9
Sequence TMhelix 10 27
Sequence outside 28 48
Sequence TMhelix 49 67
Sequence inside 68 83
Sequence TMhelix 84 105
Sequence outside 106 222
```

#### OrfT_1 *Paraesutigmatos columelliferus* Mont 10/10-1w

```
# WEBSEQUENCE Length: 238
# WEBSEQUENCE Number of predicted TMHs:  4
# WEBSEQUENCE Exp number of AAs in TMHs: 74.06976
# WEBSEQUENCE Exp number, first 60 AAs:  39.95393
# WEBSEQUENCE Total prob of N-in:        0.97550
# WEBSEQUENCE POSSIBLE N-term signal sequence
WEBSEQUENCE     TMHMM2.0        inside       1     4
WEBSEQUENCE     TMHMM2.0        TMhelix      5    23
WEBSEQUENCE     TMHMM2.0        outside     24    37
WEBSEQUENCE     TMHMM2.0        TMhelix     38    60
WEBSEQUENCE     TMHMM2.0        inside      61    66
WEBSEQUENCE     TMHMM2.0        TMhelix     67    89
WEBSEQUENCE     TMHMM2.0        outside     90    93
WEBSEQUENCE     TMHMM2.0        TMhelix     94   116
WEBSEQUENCE     TMHMM2.0        inside     117   238
```

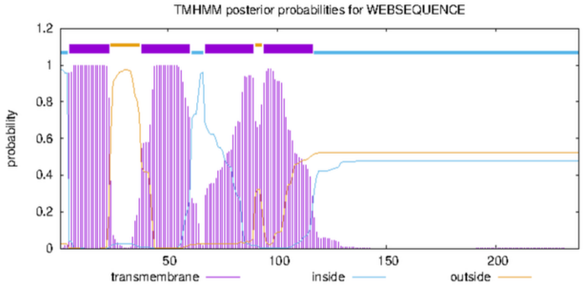


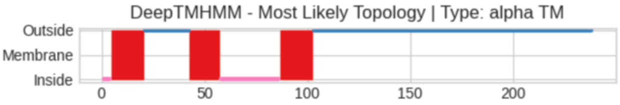


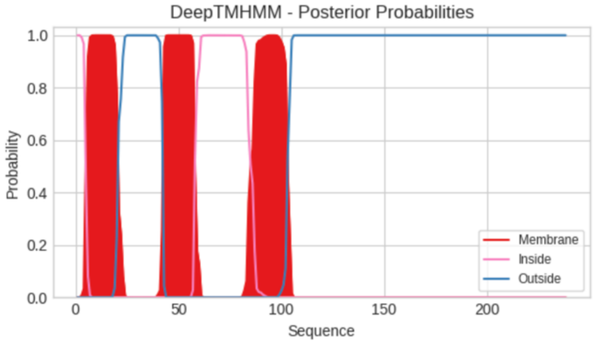


**Predicted Topologies**

```
>Sequence | TM
MKIRVLLIYMMVSVLVLYALTFLGDYKVLGNLENFTLISIKELILISLVLLFGLIALCHLKNLYKQYVFVKSFTYFLYPLNLDFLSLIFSTLLAFLIIPGVIYLNSI
MVFLQVYGYYSQEFLAVNISFEDFMALCMDNAGSVKKVRPSIAPECSSLSSGSEYSVRSASSDSAALPLDDYQVVVAGVHKDKSYVCVKLAKAISMLPSFSLRTSAG
VAKGVLGLPDVICAGYEDEPKEKK
IIIIMMMMMMMMMMMMMMMMOOOOOOOOOOOOOOOOOOOOOOMMMMMMMMMMMMMMMMIIIIIIIIIIIIIIIIIIIIIIIIIIIMMMMMMMMMMMMMMMMMOOOOO
OOOOOOOOOOOOOOOOOOOOOOOOOOOOOOOOOOOOOOOOOOOOOOOOOOOOOOOOOOOOOOOOOOOOOOOOOOOOOOOOOOOOOOOOOOOOOOOOOOOOOOOOOOO
OOOOOOOOOOOOOOOOOOOOOOOO

##gff-version 3
# Sequence Length: 238
# Sequence Number of predicted TMRs: 3
Sequence inside 1 4
Sequence TMhelix 5 20
Sequence outside 21 42
```

```
Sequence TMhelix 43 57
Sequence inside 58 86
Sequence TMhelix 87 102
Sequence outside 103 238
```

#### OrfT_2 *Paraesutigmatos columelliferus* Mont 10/10-1w

```
# WEBSEQUENCE Length: 204
# WEBSEQUENCE Number of predicted TMHs:  3
# WEBSEQUENCE Exp number of AAs in TMHs: 62.50894
# WEBSEQUENCE Exp number, first 60 AAs:  34.59563
# WEBSEQUENCE Total prob of N-in:        0.99762
# WEBSEQUENCE POSSIBLE N-term signal sequence
WEBSEQUENCE     TMHMM2.0        inside       1     6
WEBSEQUENCE     TMHMM2.0        TMhelix      7    25
WEBSEQUENCE     TMHMM2.0        outside     26    44
WEBSEQUENCE     TMHMM2.0        TMhelix     45    65
WEBSEQUENCE     TMHMM2.0        inside      66    77
WEBSEQUENCE     TMHMM2.0        TMhelix     78   100
WEBSEQUENCE     TMHMM2.0        outside    101   204
```

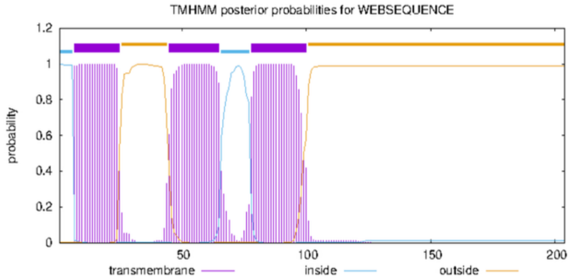


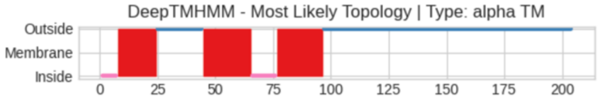


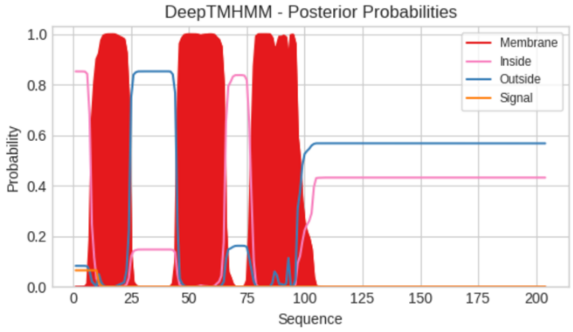


**Predicted Topologies**

```
>Sequence | TM
MGAIKKYIVLFIFILTLISTSIFVSEDPMLFSQACLSVISVKQNVFVLTAFIIALLAFFYMSLLFDGIFSSNNTYYDYFFLVILFVIGFFLFPIFLHWNSFLLISEL
YLSCAADLQSIGVNLSDFWQSCMDSKVFDIEKDELKGHMGFLFEKNGKFFACLDINESINKTSTNWSQKMALQTAKVYKQVPDVLCIEVTDAFKKSL
IIIIIIIMMMMMMMMMMMMMMMMMOOOOOOOOOOOOOOOOOOOOMMMMMMMMMMMMMMMMMMMMMIIIIIIIIIIIMMMMMMMMMMMMMMMMMMMMOOOOOOOOOOO
OOOOOOOOOOOOOOOOOOOOOOOOOOOOOOOOOOOOOOOOOOOOOOOOOOOOOOOOOOOOOOOOOOOOOOOOOOOOOOOOOOOOOOOOOOOOOOOOO

##gff-version 3
# Sequence Length: 204
# Sequence Number of predicted TMRs: 3
Sequence inside 1 7
Sequence TMhelix 8 24
Sequence outside 25 44
Sequence TMhelix 45 65
Sequence inside 66 76
Sequence TMhelix 77 96
Sequence outside 97 204
```

#### OrfT_1 *Monodopsis* sp. C141

```
# WEBSEQUENCE Length: 259
# WEBSEQUENCE Number of predicted TMHs:  4
# WEBSEQUENCE Exp number of AAs in TMHs: 87.44415
# WEBSEQUENCE Exp number, first 60 AAs:  41.36218
# WEBSEQUENCE Total prob of N-in:        0.28202
# WEBSEQUENCE POSSIBLE N-term signal sequence
WEBSEQUENCE     TMHMM2.0        outside      1     4
WEBSEQUENCE     TMHMM2.0        TMhelix      5    24
WEBSEQUENCE     TMHMM2.0        inside      25    30
WEBSEQUENCE     TMHMM2.0        TMhelix     31    53
WEBSEQUENCE     TMHMM2.0        outside     54    67
WEBSEQUENCE     TMHMM2.0        TMhelix     68    90
WEBSEQUENCE     TMHMM2.0        inside      91   110
WEBSEQUENCE     TMHMM2.0        TMhelix    111   133
WEBSEQUENCE     TMHMM2.0        outside    134   259
```

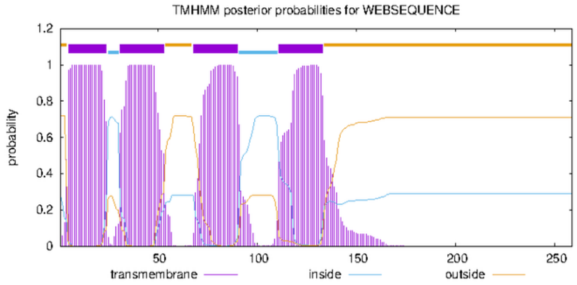


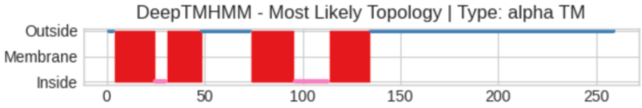


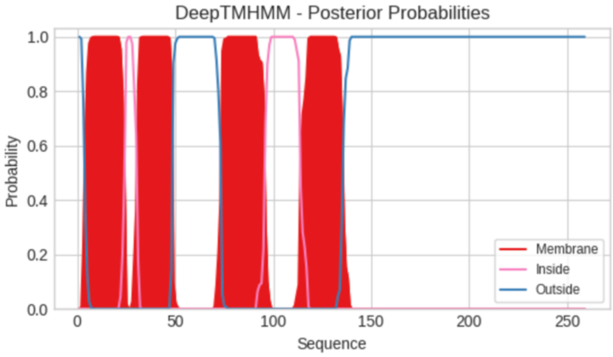


**Predicted Topologies**

```
>Sequence | TM
MKILLFIVLLLIFLFIVSSILFGIKERYLDGITWLFIFATLALIFIFFNSTNIPQLFKFEFLKDLADECCMLYILKIMYTLLFVLFIFNFAKHVFVCWKLEKLRSPN
LTTFVYFFEKVLLICAIATLVYVVIPIATTIQSYLFLAHFFGSHYELFSTLGNFGEFFSTCTEGYPNAEKVDLGDADVMVATIDGHKLLCVNPDKFLINEFDEINDV
EKALVKQRFPGLTFDNSRPLCFDLKPAENIIVESESSLNNTKVKK
OOOMMMMMMMMMMMMMMMMMMMMMIIIIIIMMMMMMMMMMMMMMMMMMOOOOOOOOOOOOOOOOOOOOOOOOOOMMMMMMMMMMMMMMMMMMMMMIIIIIIIIIIII
IIIIIMMMMMMMMMMMMMMMMMMMMMMOOOOOOOOOOOOOOOOOOOOOOOOOOOOOOOOOOOOOOOOOOOOOOOOOOOOOOOOOOOOOOOOOOOOOOOOOOOOOOO
OOOOOOOOOOOOOOOOOOOOOOOOOOOOOOOOOOOOOOOOOOOOO

##gff-version 3
# Sequence Length: 259
# Sequence Number of predicted TMRs: 4
Sequence outside 1 3
Sequence TMhelix 4 24
Sequence inside 25 30
```

Sequence TMhelix 31 48
Sequence outside 49 73
Sequence TMhelix 74 95
Sequence inside 96 113
Sequence TMhelix 114 134
Sequence outside 135 259

#### OrfT_2 *Monodopsis* sp. C141

```
# WEBSEQUENCE Length: 260
# WEBSEQUENCE Number of predicted TMHs:  4
# WEBSEQUENCE Exp number of AAs in TMHs: 87.45757
# WEBSEQUENCE Exp number, first 60 AAs:  41.37709
# WEBSEQUENCE Total prob of N-in:        0.08855
# WEBSEQUENCE POSSIBLE N-term signal sequence
WEBSEQUENCE	TMHMM2.0	outside	     1     4
WEBSEQUENCE	TMHMM2.0	TMhelix	     5    24
WEBSEQUENCE	TMHMM2.0	inside	    25    30
WEBSEQUENCE	TMHMM2.0	TMhelix	    31    53
WEBSEQUENCE	TMHMM2.0	outside	    54    72
WEBSEQUENCE	TMHMM2.0	TMhelix	    73    95
WEBSEQUENCE	TMHMM2.0	inside	    96   118
WEBSEQUENCE	TMHMM2.0	TMhelix	   119   141
WEBSEQUENCE	TMHMM2.0	outside	   142   260
```

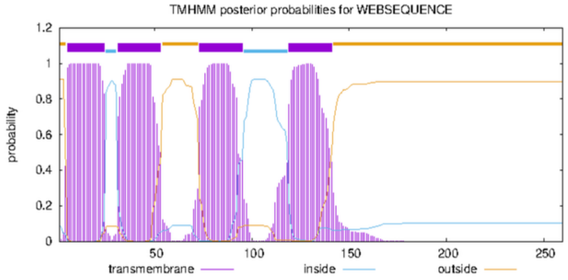


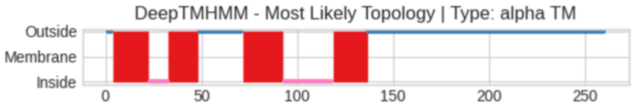


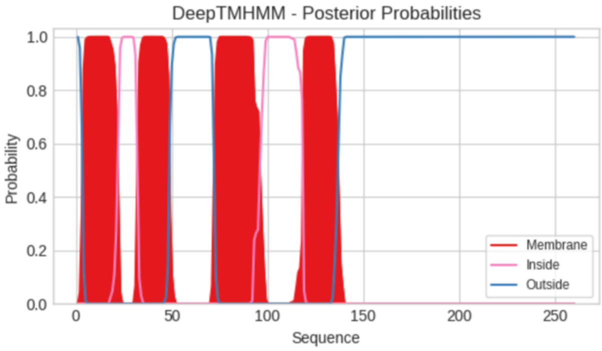


**Predicted Topologies**

```
>Sequence | TM
MKILFFIILFLIFLYIVSIILFGVKELHLDNAVWLFIFVTLAFIFYVLDGVNTPHLLLKFKVLQNLVDQCCMLYILQIMYTLLFALFIFDFIKHVCICWKLEKLRSP
NLITFIYFFKKVLLICAIATLVYLVIPVAMTIQSYLFLAHFFGNHFELFSTLGNFGEFFSMCTEGCPNAVKVDLGDADVMVTTIDGHKLLCVNPDKLLINEFGEVND
VEKALIKQRFPGLTFDNSGPLCFDLKPAENVIVESAPSLNNTEVKK
OOOMMMMMMMMMMMMMMMMMMMIIIIIIIIIIMMMMMMMMMMMMMMMMOOOOOOOOOOOOOOOOOOOOOOOMMMMMMMMMMMMMMMMMMMMMIIIIIIIIIIII
IIIIIIIIIIIIIIMMMMMMMMMMMMMMMMMMOOOOOOOOOOOOOOOOOOOOOOOOOOOOOOOOOOOOOOOOOOOOOOOOOOOOOOOOOOOOOOOOOOOOOOOO
OOOOOOOOOOOOOOOOOOOOOOOOOOOOOOOOOOOOOOOOOOOOOOOOOOOO

##gff-version 3
# Sequence Length: 260
# Sequence Number of predicted TMRs: 4
Sequence outside 1 3
Sequence TMhelix 4 22
Sequence inside 23 32
Sequence TMhelix 33 48
Sequence outside 49 71
Sequence TMhelix 72 92
Sequence inside 93 118
Sequence TMhelix 119 136
Sequence outside 137 260
```

#### OrfT *Monodopsis* sp. C73

```
# WEBSEQUENCE Length: 256
# WEBSEQUENCE Number of predicted TMHs:  4
# WEBSEQUENCE Exp number of AAs in TMHs: 83.6686600000001
# WEBSEQUENCE Exp number, first 60 AAs:  38.48587
# WEBSEQUENCE Total prob of N-in:        0.21215
# WEBSEQUENCE POSSIBLE N-term signal sequence
WEBSEQUENCE	TMHMM2.0	inside	     1     4
WEBSEQUENCE	TMHMM2.0	TMhelix	     5    22
WEBSEQUENCE	TMHMM2.0	outside	    23    31
WEBSEQUENCE	TMHMM2.0	TMhelix	    32    51
WEBSEQUENCE	TMHMM2.0	inside	    52    62
WEBSEQUENCE	TMHMM2.0	TMhelix	    63    85
WEBSEQUENCE	TMHMM2.0	outside	    86   104
WEBSEQUENCE	TMHMM2.0	TMhelix	   105   127
WEBSEQUENCE	TMHMM2.0	inside	   128   256
```

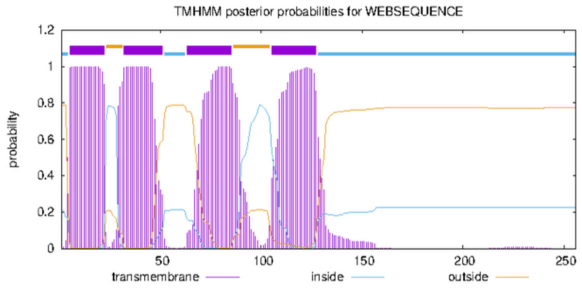


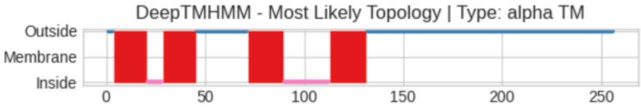


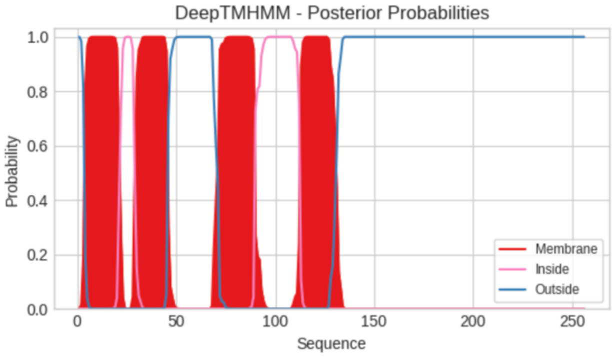

**Predicted Topologies**

```
>Sequence | TM
MKTLLFVSFFLFFIFISMLLGVKKHYHEGAVWLSIFTTLAFIFFIPNGNNIQQALELEFLKNVVNECCMLYILKVTYTFILAFFVLNFVKHVYVCWNLEKLRYSVLI
PFFEKALLICSITTLVYWIIPIAMEVQSYLFLLHFYCTQYDLLVELGSFDEFVIMYTKGDPNSEKINLGDADILLTTSKDGRKLLCVHPERLLTHELGEIGRIKKTL
ITQAFSSVTGFNNLGPVCVVLQPTEGIVVESKPSLNIAEVKK
OOOMMMMMMMMMMMMMMMMMIIIIIIIIMMMMMMMMMMMMMMMMMOOOOOOOOOOOOOOOOOOOOOOOOOOMMMMMMMMMMMMMMMMMMIIIIIIIIIIIIIIIII
IIIIIMMMMMMMMMMMMMMMMMMMOOOOOOOOOOOOOOOOOOOOOOOOOOOOOOOOOOOOOOOOOOOOOOOOOOOOOOOOOOOOOOOOOOOOOOOOOOOOOOOOOOO
OOOOOOOOOOOOOOOOOOOOOOOOOOOOOOOOOOOOOOOOOO

##gff-version 3
# Sequence Length: 256
# Sequence Number of predicted TMRs: 4
Sequence outside 1 3
Sequence TMhelix 4 20
Sequence inside 21 28
Sequence TMhelix 29 45
Sequence outside 46 71
Sequence TMhelix 72 89
Sequence inside 90 112
Sequence TMhelix 113 131
Sequence outside 132 256
```

### OrfT *Monodopsis* sp. MarTras21

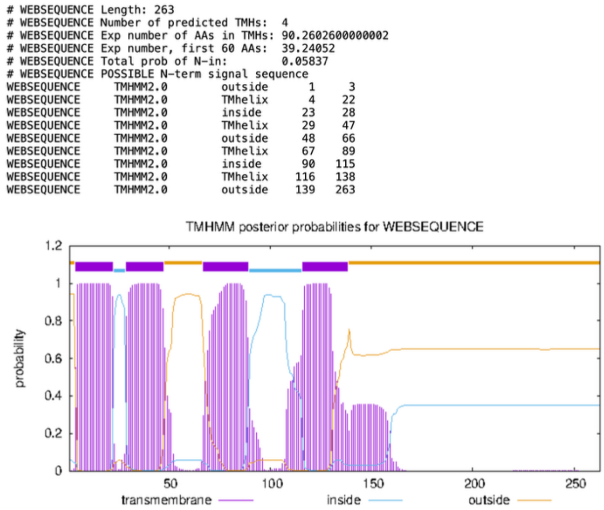

```
# WEBSEQUENCE Length: 263
# WEBSEQUENCE Number of predicted TMHs:  4
# WEBSEQUENCE Exp number of AAs in TMHs: 90.2602600000002
# WEBSEQUENCE Exp number, first 60 AAs:  39.24052
# WEBSEQUENCE Total prob of N-in:        0.05837
# WEBSEQUENCE POSSIBLE N-term signal sequence
WEBSEQUENCE    TMHMM2.0    outside      1     3
WEBSEQUENCE    TMHMM2.0    TMhelix      4    22
WEBSEQUENCE    TMHMM2.0    inside      23    28
WEBSEQUENCE    TMHMM2.0    TMhelix     29    47
WEBSEQUENCE    TMHMM2.0    outside     48    66
WEBSEQUENCE    TMHMM2.0    TMhelix     67    89
WEBSEQUENCE    TMHMM2.0    inside      90   115
WEBSEQUENCE    TMHMM2.0    TMhelix    116   138
WEBSEQUENCE    TMHMM2.0    outside    139   263
```



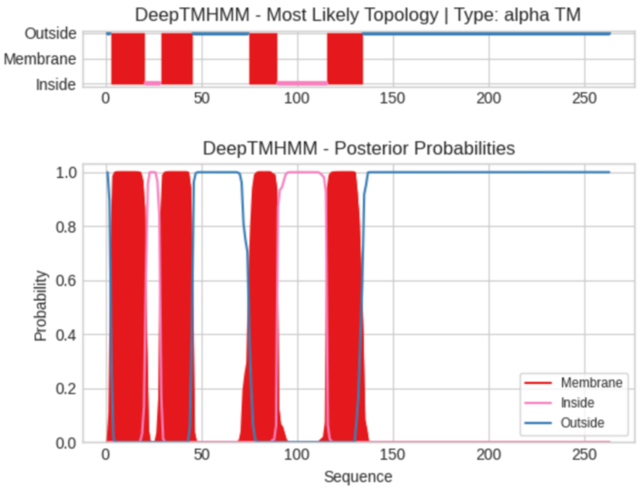


```
>Sequence | TM
MKTLFFIGFFLFFIVISILLGVKGRRLEGVVWLSIFATLAFIFFVPGGNNIQQVLGVEFLKNVVDECCMLYILKITYTAIFAFLIFNFIKNLYVCWKLEKLLWSDLT
IFFDFFEKVLLICSIAILIYWIIPMAMRVHSYLIMLHFFGINYELLSTLGNFEEFVIMCTKGDPNVEKINLGDADVMIATTKSGQKVLCAFPDKIIYNELGEVTGVK
KSLMSKVFTSFTGFKNLGPLCIALKPAEGIVVEATPSLNVSEVITKVKK
OOMMMMMMMMMMMMMMMMMMIIIIIIIIMMMMMMMMMMMMMMMMMOOOOOOOOOOOOOOOOOOOOOOOOOOOOOMMMMMMMMMMMMMMMIIIIIIIIIIIIIIIII
IIIIIIIIMMMMMMMMMMMMMMMMMMMMOOOOOOOOOOOOOOOOOOOOOOOOOOOOOOOOOOOOOOOOOOOOOOOOOOOOOOOOOOOOOOOOOOOOOOOOOOOOOOO
OOOOOOOOOOOOOOOOOOOOOOOOOOOOOOOOOOOOOOOOOOOOOOOOOO

##gff-version 3
# Sequence Length: 263
# Sequence Number of predicted TMRs: 4
Sequence outside 1 2
Sequence TMhelix 3 20
Sequence inside 21 28
Sequence TMhelix 29 45
Sequence outside 46 74
Sequence TMhelix 75 89
Sequence inside 90 115
Sequence TMhelix 116 134
Sequence outside 135 263
```

### OrfT_1 Chlorobotryaceae sp. WTwin 8/9 T-6m6.8

```
# WEBSEQUENCE Length: 219
# WEBSEQUENCE Number of predicted TMHs:  3
# WEBSEQUENCE Exp number of AAs in TMHs: 67.8195300000001
# WEBSEQUENCE Exp number, first 60 AAs:  34.68524
# WEBSEQUENCE Total prob of N-in:        0.88789
# WEBSEQUENCE POSSIBLE N-term signal sequence
WEBSEQUENCE     TMHMM2.0        inside       1     6
WEBSEQUENCE     TMHMM2.0        TMhelix      7    25
WEBSEQUENCE     TMHMM2.0        outside     26    44
WEBSEQUENCE     TMHMM2.0        TMhelix     45    67
WEBSEQUENCE     TMHMM2.0        inside      68    82
WEBSEQUENCE     TMHMM2.0        TMhelix     83   105
WEBSEQUENCE     TMHMM2.0        outside    106   219
```

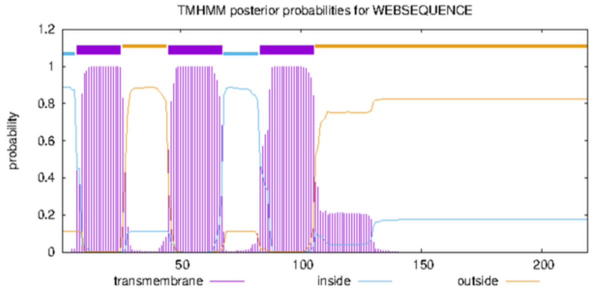


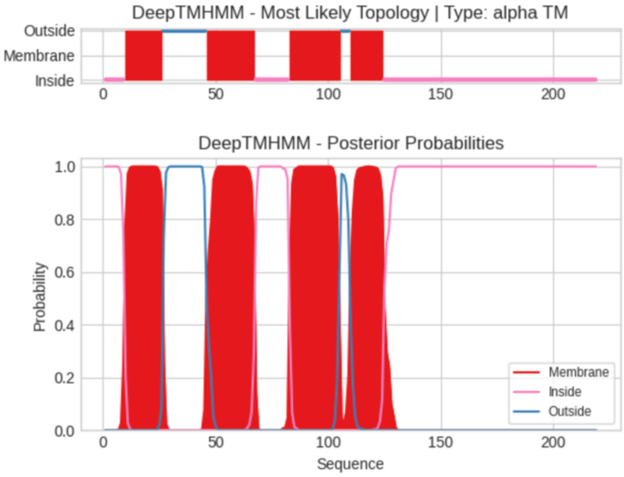


**Predicted Topologies**

```
>Sequence | TM
MSPKSKKYMLFLFYILLFSISCVVFLSEDKNVLGSSFLGLLPAKQFTYFCIVLGCALCIIYYLYILLEEKYAYANLGICIKNLLLELLFVLGFGTISIYLILIFLRW
NSVLLLMHVFIDNILAFQHIGITDYSDFWRLYIDAKEFGIGSKDFGDNLAVIVEKKGKVFGCVDITKTIDTTSASSTEKLVLHTAKSIRRVPDTVCVEMTHLLKGSL
ENPPK
IIIIIIIIIMMMMMMMMMMMMMMMMMOOOOOOOOOOOOOOOOOOOMMMMMMMMMMMMMMMMMMMMMMIIIIIIIIIIIIIIIMMMMMMMMMMMMMMMMMMMMMMMOO
OOMMMMMMMMMMMMMMMIIIIIIIIIIIIIIIIIIIIIIIIIIIIIIIIIIIIIIIIIIIIIIIIIIIIIIIIIIIIIIIIIIIIIIIIIIIIIIIIIIIIIIIIIIIIIII
IIIII

##gff-version 3
# Sequence Length: 219
# Sequence Number of predicted TMRs: 4
Sequence inside 1 9
Sequence TMhelix 10 26
Sequence outside 27 45
Sequence TMhelix 46 67
Sequence inside 68 82
Sequence TMhelix 83 105
Sequence outside 106 109
Sequence TMhelix 110 124
Sequence inside 125 219
```

### OrfT_2 Chlorobotryaceae sp. WTwin 8/9 T-6m6.8

```
# WEBSEQUENCE Length: 216
# WEBSEQUENCE Number of predicted TMHs:  3
# WEBSEQUENCE Exp number of AAs in TMHs: 63.80975
# WEBSEQUENCE Exp number, first 60 AAs:  41.07186
# WEBSEQUENCE Total prob of N-in:        0.96762
# WEBSEQUENCE POSSIBLE N-term signal sequence
WEBSEQUENCE     TMHMM2.0        inside       1     4
WEBSEQUENCE     TMHMM2.0        TMhelix      5    22
WEBSEQUENCE     TMHMM2.0        outside     23    36
WEBSEQUENCE     TMHMM2.0        TMhelix     37    59
WEBSEQUENCE     TMHMM2.0        inside      60    77
WEBSEQUENCE     TMHMM2.0        TMhelix     78   100
WEBSEQUENCE     TMHMM2.0        outside    101   216
```

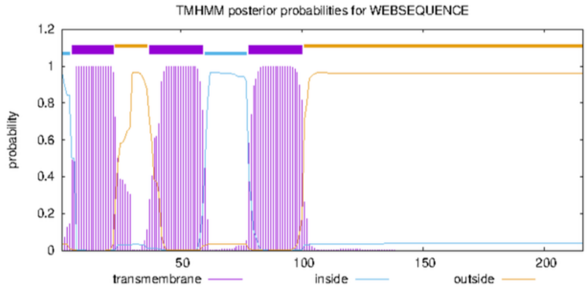


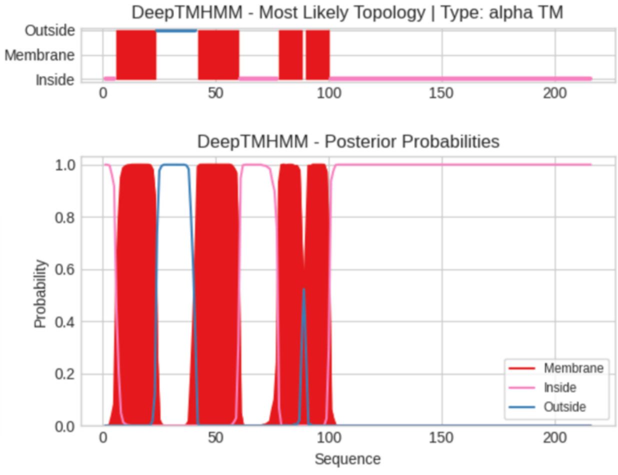


**Predicted Topologies**

```
>Sequence | TM
MPNFNFILIYFCVIGIAFAAFLLDDGTVISSSSLGLLPLKQLLYCLSGVGCALFTIYYLVKKQQSYVYANIGIYVQKLLLESIIVLGFCTITLYIIYLFLRWNSMLL
FAHLYVDNVLDFQSIGLDFGGMWQVLMDSASHQPETPEMGNHIAILTQKDDKVIACLDLAKAIDRTASTSVEKLALKTAKSLYDLPDVTCFEVTDAFKRSMEQIKQV
KK
IIIIIMMMMMMMMMMMMMMMMMMOOOOOOOOOOOOOOOOOOMMMMMMMMMMMMMMMMMMMIIIIIIIIIIIIIIIIIMMMMMMMMMMOMMMMMMMMMMMMIIIIIII
IIIIIIIIIIIIIIIIIIIIIIIIIIIIIIIIIIIIIIIIIIIIIIIIIIIIIIIIIIIIIIIIIIIIIIIIIIIIIIIIIIIIIIIIIIIIIIIIIIIIIIIIIII
II

##gff-version 3
# Sequence Length: 216
# Sequence Number of predicted TMRs: 4
Sequence inside 1 5
Sequence TMhelix 6 23
Sequence outside 24 41
```

```
Sequence TMhelix 42 60
Sequence inside 61 77
Sequence TMhelix 78 88
Sequence outside 89 89
Sequence TMhelix 90 100
Sequence inside 101 216
```

### OrfU

#### OrfU *Trachydiscus minutus* CCALA 838

```
# WEBSEQUENCE Length: 271
# WEBSEQUENCE Number of predicted TMHs:  4
# WEBSEQUENCE Exp number of AAs in TMHs: 102.88635
# WEBSEQUENCE Exp number, first 60 AAs:  42.22372
# WEBSEQUENCE Total prob of N-in:        0.94397
# WEBSEQUENCE POSSIBLE N-term signal sequence
WEBSEQUENCE     TMHMM2.0        inside       1    12
WEBSEQUENCE     TMHMM2.0        TMhelix     13    35
WEBSEQUENCE     TMHMM2.0        outside     36    38
WEBSEQUENCE     TMHMM2.0        TMhelix     39    61
WEBSEQUENCE     TMHMM2.0        inside      62    81
WEBSEQUENCE     TMHMM2.0        TMhelix     82   104
WEBSEQUENCE     TMHMM2.0        outside    105   107
WEBSEQUENCE     TMHMM2.0        TMhelix    108   130
WEBSEQUENCE     TMHMM2.0        inside     131   271
```

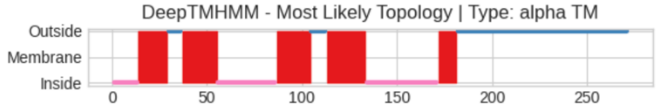


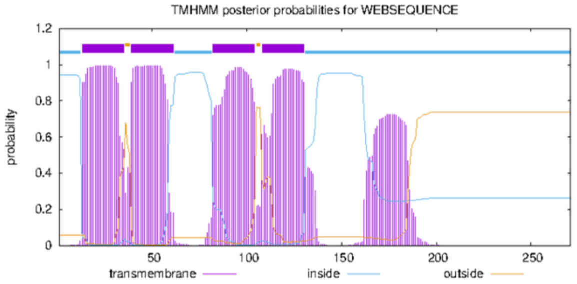


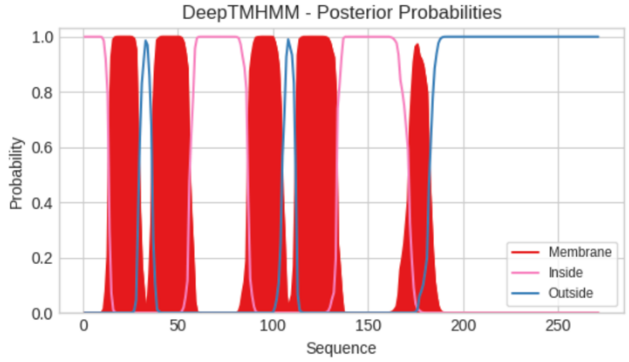


**Predicted Topologies**

```
>Sequence | TM
MVKPKKDFKSVDKGLLFPFFLLITAPLVYSTWGLASAYFYVLFISLGHAYFLLYFPYIQDSNYYQYIIMKKRFYIENLRYKYFFSQNVVTFSRVVVLFSFYLLFNFP
TIYPAIFPIHCIGTVIFMSSVSVDIFNIVSRPRLPDKIKIYHPLLQNRYGPITEIVINKAVPLCASIAGYAASGYLAFAGGFKALNGISEIDPIRNDLLNRYYRFPL
THKWNEMSLSAWSTFRNKPRVCELSYDRQIFVFLEREGQFVETVAKAAANLCDIKKD
IIIIIIIIIIIIIMMMMMMMMMMMMMMMMOOOOOOOMMMMMMMMMMMMMMMMMMMIIIIIIIIIIIIIIIIIIIIIIIIIIIIIIIMMMMMMMMMMMMMMMMMMOOO
OOOOOMMMMMMMMMMMMMMMMMMMMMIIIIIIIIIIIIIIIIIIIIIIIIIIIIIIIIIIIIIIMMMMMMMMMMOOOOOOOOOOOOOOOOOOOOOOOOOOOOOOOOO
OOOOOOOOOOOOOOOOOOOOOOOOOOOOOOOOOOOOOOOOOOOOOOOOOOOOOOOOO

##gff-version 3
# Sequence Length: 271
# Sequence Number of predicted TMRs: 5
Sequence inside 1 13
Sequence TMhelix 14 29
Sequence outside 30 36
Sequence TMhelix 37 55
Sequence inside 56 86
Sequence TMhelix 87 104
Sequence outside 105 112
Sequence TMhelix 113 133
Sequence inside 134 171
Sequence TMhelix 172 181
Sequence outside 182 271
```

#### OrfU *Monodopsis* sp. C143

```
# WEBSEQUENCE Length: 274
# WEBSEQUENCE Number of predicted TMHs:  5
# WEBSEQUENCE Exp number of AAs in TMHs: 103.92029
# WEBSEQUENCE Exp number, first 60 AAs:  36.78949
# WEBSEQUENCE Total prob of N-in:        0.99938
# WEBSEQUENCE POSSIBLE N-term signal sequence
WEBSEQUENCE     TMHMM2.0        inside       1    20
WEBSEQUENCE     TMHMM2.0        TMhelix     21    40
WEBSEQUENCE     TMHMM2.0        outside     41    43
WEBSEQUENCE     TMHMM2.0        TMhelix     44    66
WEBSEQUENCE     TMHMM2.0        inside      67    86
WEBSEQUENCE     TMHMM2.0        TMhelix     87   109
WEBSEQUENCE     TMHMM2.0        outside    110   113
WEBSEQUENCE     TMHMM2.0        TMhelix    114   133
WEBSEQUENCE     TMHMM2.0        inside     134   165
WEBSEQUENCE     TMHMM2.0        TMhelix    166   188
WEBSEQUENCE     TMHMM2.0        outside    189   274
```

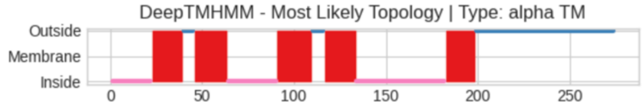


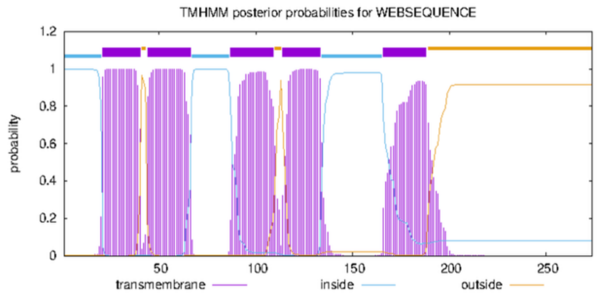


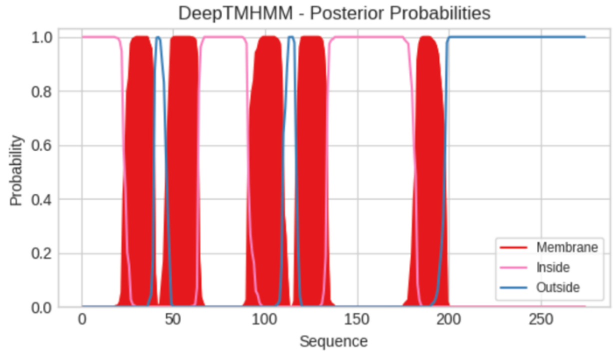

**Predicted Topologies**

```
>Sequence | TM
VFKKTKLSKRIKSPIAAKINTYFYFPIFMLVSVSIIYLLYDWSITVLYIMLLSLLHIVFSFVFPYTRESKYIFYSKQKENFFKQHEKYGQIFTEFQVTLAYAFMIYS
FFQMFKGSPFMVLLLLFQFISLVVIFSHDVTNIRNIIARPSLPSDIKSLRPFYYSTRRFGAFFDIVYPICKTCIKVILAAYVGYYGGWMFWNGEQVYEPLHSKIISW
HYNFSQDYEWTKTRHSDWVDFSTNLTFSNLKHDRQVELFDLWQKSKKTKIFADLNILLNK
IIIIIIIIIIIIIIIIIIIIIIMMMMMMMMMMMMMMMMMOOOOOOMMMMMMMMMMMMMMMMMMIIIIIIIIIIIIIIIIIIIIIIIIIIIMMMMMMMMMMMMMMMMM
MMOOOOOOOMMMMMMMMMMMMMMMMMIIIIIIIIIIIIIIIIIIIIIIIIIIIIIIIIIIIIIIIIIIIIIIIIIIMMMMMMMMMMMMMMMMOOOOOOOOOOOOOOOO
OOOOOOOOOOOOOOOOOOOOOOOOOOOOOOOOOOOOOOOOOOOOOOOOOOOOOOOOOOOO

##gff-version 3
# Sequence Length: 274
# Sequence Number of predicted TMRs: 5
Sequence inside 1 22
Sequence TMhelix 23 39
Sequence outside 40 45
Sequence TMhelix 46 63
Sequence inside 64 90
Sequence TMhelix 91 109
Sequence outside 110 116
Sequence TMhelix 117 133
Sequence inside 134 182
Sequence TMhelix 183 198
Sequence outside 199 274
```

### OrfU (Orf254) *Monodopsis* sp. MarTras21

```
# WEBSEQUENCE Length: 283
# WEBSEQUENCE Number of predicted TMHs:  4
# WEBSEQUENCE Exp number of AAs in TMHs: 88.5345000000001
# WEBSEQUENCE Exp number, first 60 AAs:  36.20429
# WEBSEQUENCE Total prob of N-in:        0.99366
# WEBSEQUENCE POSSIBLE N-term signal sequence
WEBSEQUENCE     TMHMM2.0        inside       1    19
WEBSEQUENCE     TMHMM2.0        TMhelix     20    42
WEBSEQUENCE     TMHMM2.0        outside     43    45
WEBSEQUENCE     TMHMM2.0        TMhelix     46    68
WEBSEQUENCE     TMHMM2.0        inside      69    88
WEBSEQUENCE     TMHMM2.0        TMhelix     89   111
WEBSEQUENCE     TMHMM2.0        outside    112   115
WEBSEQUENCE     TMHMM2.0        TMhelix    116   138
WEBSEQUENCE     TMHMM2.0        inside     139   283
```

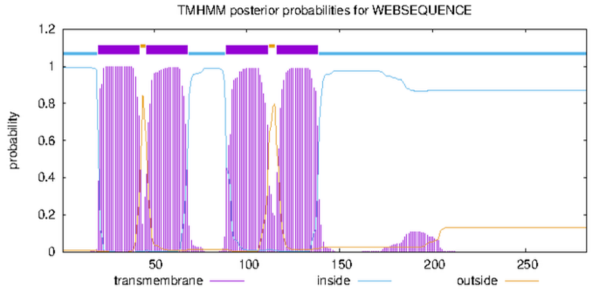


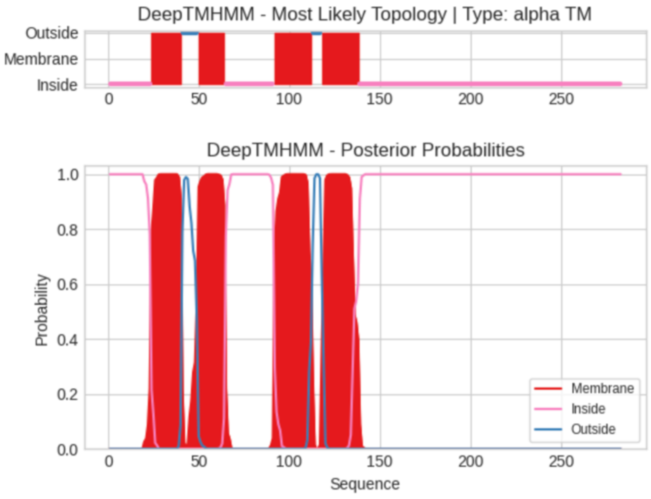


**Predicted Topologies**

```
>Sequence | TM
IVLKETKLSKRIKSPIAAKINTYFYFPIFMLVSVSIIFLLYGWSITAPYIMLSSLLHLVFSFVFPYTHESKYAFYLKQKENFFKQHEKYGQIFTEFQVTLAFAFMIY
SFFQMLSGSSFMALFVSFQFISLVIIFSHGVTNIRNIIARPSLPSDIKLLRPYYYSTRRFGAFSDAFFETVYPVCKTCIKISLAAYTGYYGGWMFWNGDQVYEPIHC
KITSWHYNFPKDYEWTKTRHSDWVDFSTDPELSNLKHNCQVELFDLWQESKKSKLRADICILLDKQLVT
IIIIIIIIIIIIIIIIIIIIIIIMMMMMMMMMMMMMMMMMOOOOOOOOOMMMMMMMMMMMMMMMIIIIIIIIIIIIIIIIIIIIIIIIIIIMMMMMMMMMMMMMMMMM
MMMMOOOOOMMMMMMMMMMMMMMMMMMMMMIIIIIIIIIIIIIIIIIIIIIIIIIIIIIIIIIIIIIIIIIIIIIIIIIIIIIIIIIIIIIIIIIIIIIIIIIIIII
IIIIIIIIIIIIIIIIIIIIIIIIIIIIIIIIIIIIIIIIIIIIIIIIIIIIIIIIIIIIIIIIIIII

##gff-version 3
# Sequence Length: 283
# Sequence Number of predicted TMRs: 4
Sequence inside 1 23
Sequence TMhelix 24 40
Sequence outside 41 49
Sequence TMhelix 50 64
Sequence inside 65 91
Sequence TMhelix 92 112
Sequence outside 113 117
Sequence TMhelix 118 138
Sequence inside 139 283
```

### OrfV

#### OrfV Goniochloridales sp. BogD 8/9 T-2w

```
# WEBSEQUENCE Length: 258
# WEBSEQUENCE Number of predicted TMHs:  4
# WEBSEQUENCE Exp number of AAs in TMHs: 84.9429
# WEBSEQUENCE Exp number, first 60 AAs:  37.82259
# WEBSEQUENCE Total prob of N-in:        0.43278
# WEBSEQUENCE POSSIBLE N-term signal sequence
WEBSEQUENCE     TMHMM2.0        outside      1     3
WEBSEQUENCE     TMHMM2.0        TMhelix      4    23
WEBSEQUENCE     TMHMM2.0        inside      24    29
WEBSEQUENCE     TMHMM2.0        TMhelix     30    52
WEBSEQUENCE     TMHMM2.0        outside     53    66
WEBSEQUENCE     TMHMM2.0        TMhelix     67    89
WEBSEQUENCE     TMHMM2.0        inside      90    95
WEBSEQUENCE     TMHMM2.0        TMhelix     96   118
WEBSEQUENCE     TMHMM2.0        outside    119   258
```

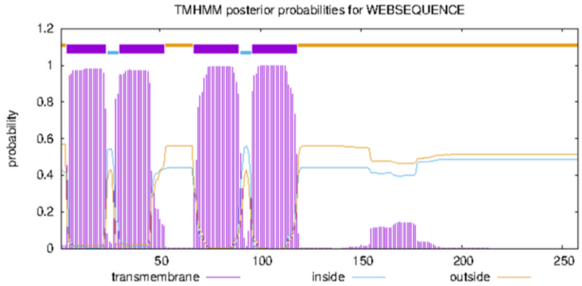


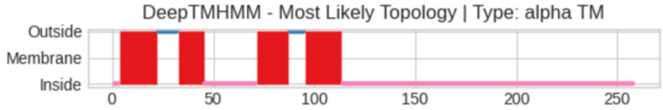


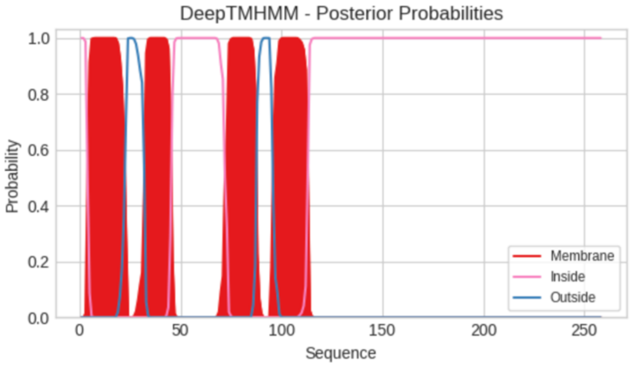


**Predicted Topologies**

```
>Sequence | TM
MSRFFIPYFSLSCINLIISYLLFDLDVFFIRQFFLTTGLLFFVVFEQDLTIFSKDSTYINSILFKKYPILRFEFPFTLLCLGSIFVISSFTTEQYLLNIVGGILLIT
VLLIFYRYYCIWETPELAIYFKIPLHLGKNGKRNFSTWPTWAMAAKPVVIKVCVECTKLSGFAFMAAYGVPKAIYGANYRSPVVQWVGYQFTGLKSNNEFFYSLAVD
FVNQYPDLKTEIVEDDSCTIVPQKLMQQISKKNMPVPPLKLKDF
IIIMMMMMMMMMMMMMMMMMMMOOOOOOOOOOMMMMMMMMMMMMMIIIIIIIIIIIIIIIIIIIIIIIIIIMMMMMMMMMMMMMMMMOOOOOOOOMMMMMMMMMMMM
MMMMMMIIIIIIIIIIIIIIIIIIIIIIIIIIIIIIIIIIIIIIIIIIIIIIIIIIIIIIIIIIIIIIIIIIIIIIIIIIIIIIIIIIIIIIIIIIIIIIIIIIIII
IIIIIIIIIIIIIIIIIIIIIIIIIIIIIIIIIIIIIIIIIIII

##gff-version 3
# Sequence Length: 258
# Sequence Number of predicted TMRs: 4
Sequence inside 1 3
Sequence TMhelix 4 22
Sequence outside 23 32
Sequence TMhelix 33 45
Sequence inside 46 71
Sequence TMhelix 72 87
Sequence outside 88 95
Sequence TMhelix 96 113
Sequence inside 114 258
```

#### OrfV_1 *Paraeustigmatos columelliferus* Mont 10/10-1w

```
# WEBSEQUENCE Length: 297
# WEBSEQUENCE Number of predicted TMHs:  4
# WEBSEQUENCE Exp number of AAs in TMHs: 87.72743
# WEBSEQUENCE Exp number, first 60 AAs:  32.37255
# WEBSEQUENCE Total prob of N-in:        0.66354
# WEBSEQUENCE POSSIBLE N-term signal sequence
WEBSEQUENCE     TMHMM2.0        inside       1    20
WEBSEQUENCE     TMHMM2.0        TMhelix     21    39
WEBSEQUENCE     TMHMM2.0        outside     40    42
WEBSEQUENCE     TMHMM2.0        TMhelix     43    60
WEBSEQUENCE     TMHMM2.0        inside      61    86
WEBSEQUENCE     TMHMM2.0        TMhelix     87   106
WEBSEQUENCE     TMHMM2.0        outside    107   115
WEBSEQUENCE     TMHMM2.0        TMhelix    116   138
WEBSEQUENCE     TMHMM2.0        inside     139   297
```

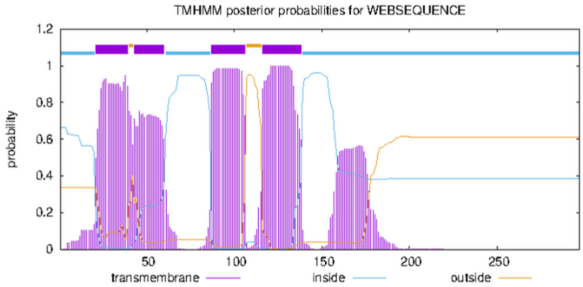


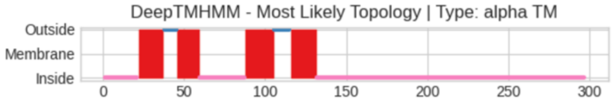


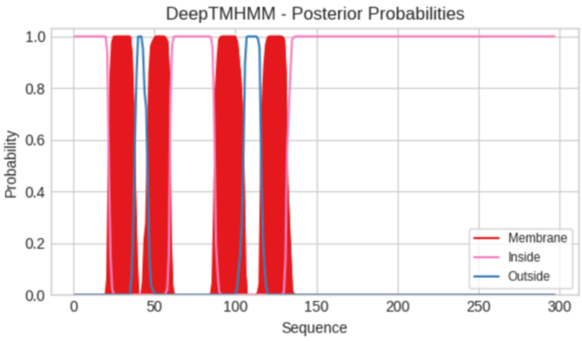


**Predicted Topologies**

```
>Sequence | TM
MKIKLYLSQIINIFLSMLKKHPYLIVGFTLSNIIVCWFFNFSLVTLAVRQLLVLLFLFFYTQKSEYFQEYLKECTEIRKRLAQESPLLDYLPVMYFLSLMGYYWLLD
QPIDTPFGVFPLLVLNLWLLTFFLTFYRASLIMHISELRDKHNIYQASRVYGVASFATIKTTAHTFSMGGRMVIALIFGLETVPRFRDGDLDARGIPSNLLSFYQYG
LTARTNDEMKSALYLLKRYPELKSVIIDPSDGHTIDPRPLAQQLLEKNLVVKPFFRLVTEPSDEALGKIQDLAKSSEVKVKWR
IIIIIIIIIIIIIIIIIIIIIMMMMMMMMMMMMMMMMOOOOOOOOMMMMMMMMMMMMMMIIIIIIIIIIIIIIIIIIIIIIIIIIIMMMMMMMMMMMMMMMMMMMOO
OOOOOOOOMMMMMMMMMMMMMMMMIIIIIIIIIIIIIIIIIIIIIIIIIIIIIIIIIIIIIIIIIIIIIIIIIIIIIIIIIIIIIIIIIIIIIIIIIIIIIIIIIII
IIIIIIIIIIIIIIIIIIIIIIIIIIIIIIIIIIIIIIIIIIIIIIIIIIIIIIIIIIIIIIIIIIIIIIIIIIIIIIIIIII
```

```
##gff-version 3
# Sequence Length: 297
# Sequence Number of predicted TMRs: 4
Sequence inside 1 21
Sequence TMhelix 22 37
Sequence outside 38 45
Sequence TMhelix 46 59
Sequence inside 60 87
Sequence TMhelix 88 105
Sequence outside 106 115
Sequence TMhelix 116 131
Sequence inside 132 297
```

#### OrfV_2 *Paraeustigmatos columelliferus* Mont 10/10-1w

```
# WEBSEQUENCE Length: 299
# WEBSEQUENCE Number of predicted TMHs:  4
# WEBSEQUENCE Exp number of AAs in TMHs: 87.4803900000001
# WEBSEQUENCE Exp number, first 60 AAs:  36.91347
# WEBSEQUENCE Total prob of N-in:        0.98795
# WEBSEQUENCE POSSIBLE N-term signal sequence
WEBSEQUENCE     TMHMM2.0        inside       1    20
WEBSEQUENCE     TMHMM2.0        TMhelix     21    40
WEBSEQUENCE     TMHMM2.0        outside     41    43
WEBSEQUENCE     TMHMM2.0        TMhelix     44    66
WEBSEQUENCE     TMHMM2.0        inside      67    85
WEBSEQUENCE     TMHMM2.0        TMhelix     86   105
WEBSEQUENCE     TMHMM2.0        outside    106   114
WEBSEQUENCE     TMHMM2.0        TMhelix    115   137
WEBSEQUENCE     TMHMM2.0        inside     138   299
```

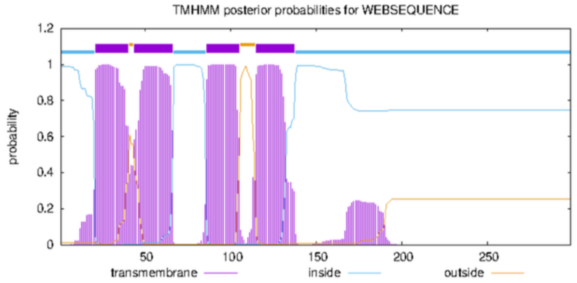


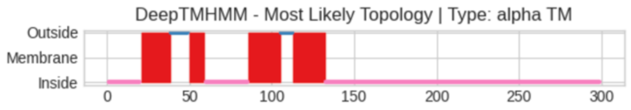


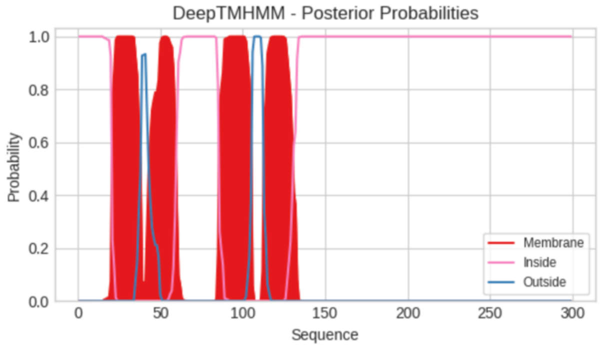


**Predicted Topologies**

```
>Sequence | TM
MFMKKIKNFFISFVKFVGEHLYVLLGLTLVNILLCWVLKFGILVFFLRQIVIFGGYFCYTKYSWFFKECRNETLFINTRLLEESSILKFSPVVSFISFLGIFWFLDQ
PENSDFGFTALIICNIWFINVMMNIQKLSIIRRQPELKQKFSLLKSIKLYGIASYHTAKTLYHGGYTAVKFVASLLFGLEGIARLGEGDIDARGLPSNLYSYYKHGL
TARTNSQLRNGVHLVDLYPELRSQIVYQKDGHTIDPRLLAQKLLEKKAENKDFLKFKLEHEHKLNLPQPQLPNWGEWLDWYRGKK
IIIIIIIIIIIIIIIIIIIIMMMMMMMMMMMMMMMMMMOOOOOOOOOOOMMMMMMMMMMIIIIIIIIIIIIIIIIIIIIIIIIIIMMMMMMMMMMMMMMMMMMMMOO
OOOOOMMMMMMMMMMMMMMMMMMMMIIIIIIIIIIIIIIIIIIIIIIIIIIIIIIIIIIIIIIIIIIIIIIIIIIIIIIIIIIIIIIIIIIIIIIIIIIIIIIIIIII
IIIIIIIIIIIIIIIIIIIIIIIIIIIIIIIIIIIIIIIIIIIIIIIIIIIIIIIIIIIIIIIIIIIIIIIIIIIIIIIIIIIIIII

##gff-version 3
# Sequence Length: 299
# Sequence Number of predicted TMRs: 4
Sequence inside 1 20
Sequence TMhelix 21 38
Sequence outside 39 49
Sequence TMhelix 50 59
Sequence inside 60 85
Sequence TMhelix 86 105
Sequence outside 106 112
Sequence TMhelix 113 132
Sequence inside 133 299
```

#### OrfV_1 *Monodopsis* sp. C141

```
# WEBSEQUENCE Length: 278
# WEBSEQUENCE Number of predicted TMHs:  4
# WEBSEQUENCE Exp number of AAs in TMHs: 86.8886
# WEBSEQUENCE Exp number, first 60 AAs:  36.61783
# WEBSEQUENCE Total prob of N-in:        0.87531
# WEBSEQUENCE POSSIBLE N-term signal sequence
WEBSEQUENCE     TMHMM2.0        inside       1    12
WEBSEQUENCE     TMHMM2.0        TMhelix     13    33
WEBSEQUENCE     TMHMM2.0        outside     34    37
WEBSEQUENCE     TMHMM2.0        TMhelix     38    57
WEBSEQUENCE     TMHMM2.0        inside      58    77
WEBSEQUENCE     TMHMM2.0        TMhelix     78   100
WEBSEQUENCE     TMHMM2.0        outside    101   109
WEBSEQUENCE     TMHMM2.0        TMhelix    110   132
WEBSEQUENCE     TMHMM2.0        inside     133   278
```

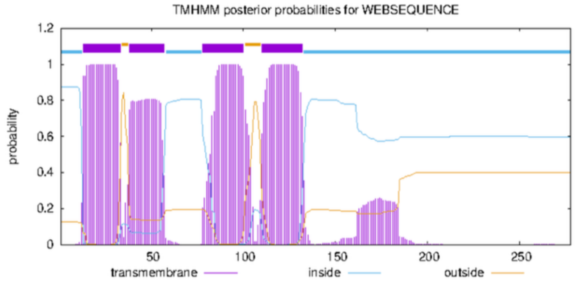


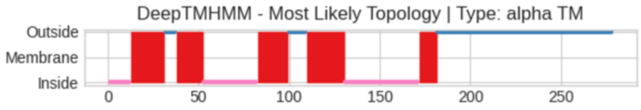


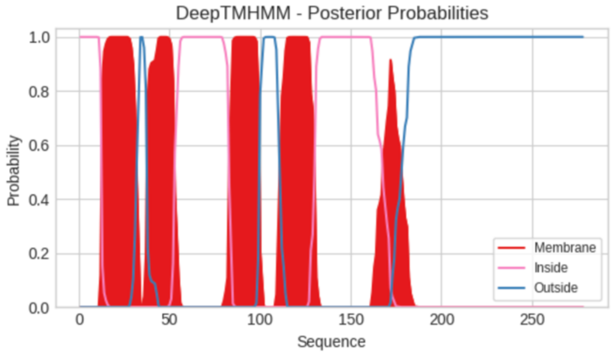

**Predicted Topologies**

```
>Sequence | TM
MFKFDKNQHNKNFLYTLIFLSFSLMNIFLAFFIKDIYIFLTRETVMCFLLLLHYIYSDELAALSKEDEQISLILNKKYPVLRFEFPLTLAGGLSIFVFFSLSIYPDC
TTICMLSLIIFCVYSVLAFYSVFCVLRTPETRDFYNIKISISQHGIWSFYTFSKVFHVCKLCAKVGGLGLVCTWAMPKVFHGDIMYRGDLLNFIAYPYTGKKANSDW
DMMKANELLNSYPEDRILILKKDGCTIDPSLLKKQMQARDVEFPLAGMEHLYRLNFWPFSREGK
IIIIIIIIIIIIMMMMMMMMMMMMMMMMMMMOOOOOOMMMMMMMMMMMMMMMIIIIIIIIIIIIIIIIIIIIIIIIIIIIIIMMMMMMMMMMMMMMMMMOOOOOOOO
OOMMMMMMMMMMMMMMMMMMMMMIIIIIIIIIIIIIIIIIIIIIIIIIIIIIIIIIIIIIIIIIMMMMMMMMMMOOOOOOOOOOOOOOOOOOOOOOOOOOOOOOOOO
OOOOOOOOOOOOOOOOOOOOOOOOOOOOOOOOOOOOOOOOOOOOOOOOOOOOOOOOOOOOOOOO

##gff-version 3
# Sequence Length: 278
# Sequence Number of predicted TMRs: 5
Sequence inside 1 12
Sequence TMhelix 13 31
Sequence outside 32 37
Sequence TMhelix 38 52
Sequence inside 53 82
Sequence TMhelix 83 99
Sequence outside 100 109
Sequence TMhelix 110 130
Sequence inside 131 171
Sequence TMhelix 172 181
Sequence outside 182 278
```

#### OrfV_2 *Monodopsis* sp. C141

```
# WEBSEQUENCE Length: 279
# WEBSEQUENCE Number of predicted TMHs:  4
# WEBSEQUENCE Exp number of AAs in TMHs: 89.5661900000001
# WEBSEQUENCE Exp number, first 60 AAs:  39.31078
# WEBSEQUENCE Total prob of N-in:        0.79560
# WEBSEQUENCE POSSIBLE N-term signal sequence
WEBSEQUENCE     TMHMM2.0        inside       1    12
WEBSEQUENCE     TMHMM2.0        TMhelix     13    32
WEBSEQUENCE     TMHMM2.0        outside     33    35
WEBSEQUENCE     TMHMM2.0        TMhelix     36    58
WEBSEQUENCE     TMHMM2.0        inside      59    84
WEBSEQUENCE     TMHMM2.0        TMhelix     85   104
WEBSEQUENCE     TMHMM2.0        outside    105   107
WEBSEQUENCE     TMHMM2.0        TMhelix    108   130
WEBSEQUENCE     TMHMM2.0        inside     131   279
```

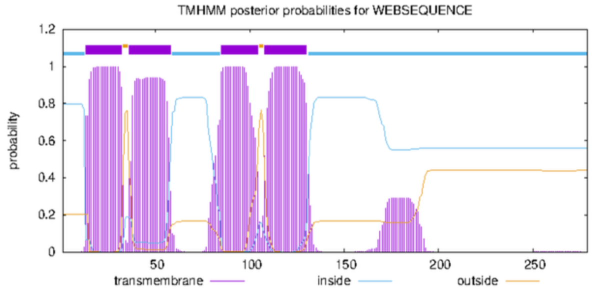


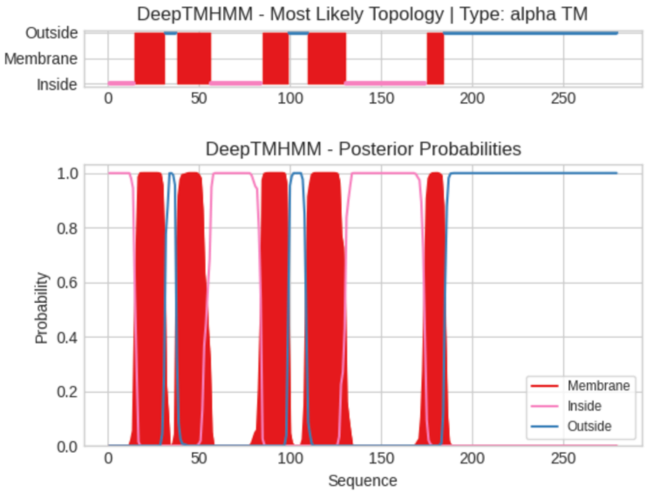


**Predicted Topologies**

```
>Sequence | TM
MFNFNELRNNKKLKYTFIFFFLSLLNIFVAFYFKDIYVFLIRETIMCFFLLFNYFFSNELEALSQENNNVHYILKNKYPILKLEFFLTVGFTLGVIVLANLSIYPEY
AIICMIASFICCTYIVLIIYSAFCILRTPEARDSFNIKKSISRHGIRSFSTYSKIFNTCKSCGKAAGVGLFCTWLIPAFTNWDVTHRSDLFNSLSAPLTGGMKARSG
RDISTANYLFDLYPEDRPLVMQKDGCTIDSSLLKKRMAARELEVPPLVNINITVIPDLWGKSKGK
IIIIIIIIIIIIIIMMMMMMMMMMMMMMMMMOOOOOOMMMMMMMMMMMMMMMMMMMIIIIIIIIIIIIIIIIIIIIIIIIIIIIMMMMMMMMMMMMMMMOOOOOOOO
OOMMMMMMMMMMMMMMMMMMMMMIIIIIIIIIIIIIIIIIIIIIIIIIIIIIIIIIIIIIIIIIIIIMMMMMMMMMMOOOOOOOOOOOOOOOOOOOOOOOOOOOOOO
OOOOOOOOOOOOOOOOOOOOOOOOOOOOOOOOOOOOOOOOOOOOOOOOOOOOOOOOOOOOOOOOO

##gff-version 3
# Sequence Length: 279
# Sequence Number of predicted TMRs: 5
Sequence inside 1 14
Sequence TMhelix 15 31
Sequence outside 32 37
Sequence TMhelix 38 56
Sequence inside 57 84
Sequence TMhelix 85 99
Sequence outside 100 109
Sequence TMhelix 110 130
Sequence inside 131 174
Sequence TMhelix 175 184
Sequence outside 185 279
```

#### OrfV_3 *Monodopsis* sp. C141

```
# WEBSEQUENCE Length: 275
# WEBSEQUENCE Number of predicted TMHs:  3
# WEBSEQUENCE Exp number of AAs in TMHs: 86.3095200000001
# WEBSEQUENCE Exp number, first 60 AAs:  32.44734
# WEBSEQUENCE Total prob of N-in:        0.63189
# WEBSEQUENCE POSSIBLE N-term signal sequence
WEBSEQUENCE     TMHMM2.0        outside      1    14
WEBSEQUENCE     TMHMM2.0        TMhelix     15    33
WEBSEQUENCE     TMHMM2.0        inside      34    78
WEBSEQUENCE     TMHMM2.0        TMhelix     79   101
WEBSEQUENCE     TMHMM2.0        outside    102   110
WEBSEQUENCE     TMHMM2.0        TMhelix    111   133
WEBSEQUENCE     TMHMM2.0        inside     134   275
```

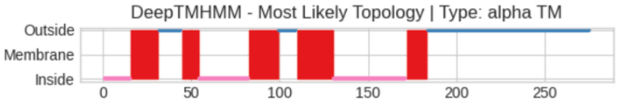


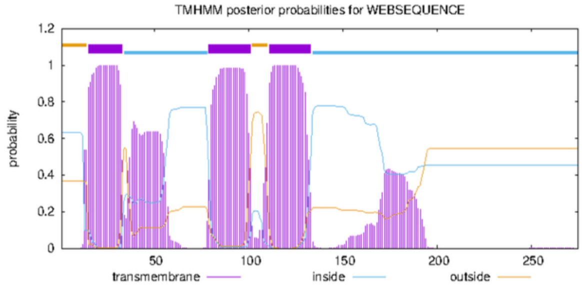


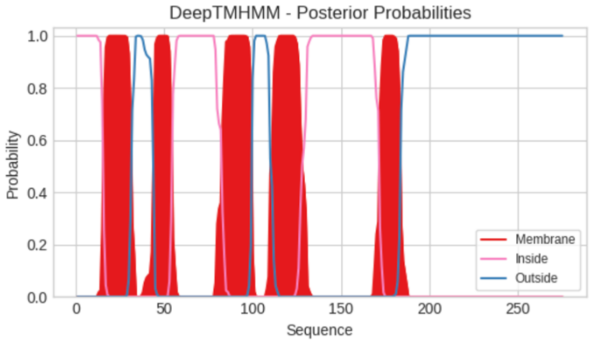


**Predicted Topologies**

```
>Sequence | TM
MFNFNELRNNKKLKYTLIFFSLSLLNAFIAFYLKDIYVFLIRETIMCFFLLFNYSYSDELVIWSQEEDYIRLILKEKHPILRLEFPLSMCFILGLTIFFNYRSYPGC
AGICMITLLTCCVYIALAFYSVFCIFKTPEVRDCYNIKRSISHHGIWSFSTFIKVLHICKPCLKVVALIAISAALIPKVLSGDITYRGDLMNTLSFPWTEMKSHNDW
EIMKANYLFDQYPEDRLLVMQKDGCTIDSLLLDKQMQARSVNFPIQYPPSVNFWSFLHKDK
IIIIIIIIIIIIIIIMMMMMMMMMMMMMMMMOOOOOOOOOOOOOMMMMMMMMMMIIIIIIIIIIIIIIIIIIIIIIIIIIIIMMMMMMMMMMMMMMMMMOOOOOOOO
OOMMMMMMMMMMMMMMMMMMMMMIIIIIIIIIIIIIIIIIIIIIIIIIIIIIIIIIIIIIIIIIMMMMMMMMMMMMOOOOOOOOOOOOOOOOOOOOOOOOOOOOOOO
OOOOOOOOOOOOOOOOOOOOOOOOOOOOOOOOOOOOOOOOOOOOOOOOOOOOOOOOOOOOO
```

```
##gff-version 3
# Sequence Length: 275
# Sequence Number of predicted TMRs: 5
Sequence inside 1 15
Sequence TMhelix 16 31
Sequence outside 32 44
Sequence TMhelix 45 54
Sequence inside 55 82
Sequence TMhelix 83 99
Sequence outside 100 109
Sequence TMhelix 110 130
Sequence inside 131 171
Sequence TMhelix 172 183
Sequence outside 184 275
```

#### OrfV_1 *Pseudotetraëdriella kamillae* SAG 2056

```
# WEBSEQUENCE Length: 297
# WEBSEQUENCE Number of predicted TMHs:  5
# WEBSEQUENCE Exp number of AAs in TMHs: 97.1195600000002
# WEBSEQUENCE Exp number, first 60 AAs:  35.4863
# WEBSEQUENCE Total prob of N-in:        0.98929
# WEBSEQUENCE POSSIBLE N-term signal sequence
WEBSEQUENCE     TMHMM2.0        inside       1     6
WEBSEQUENCE     TMHMM2.0        TMhelix      7    26
WEBSEQUENCE     TMHMM2.0        outside     27    29
WEBSEQUENCE     TMHMM2.0        TMhelix     30    47
WEBSEQUENCE     TMHMM2.0        inside      48    74
WEBSEQUENCE     TMHMM2.0        TMhelix     75    97
WEBSEQUENCE     TMHMM2.0        outside     98   100
WEBSEQUENCE     TMHMM2.0        TMhelix    101   123
WEBSEQUENCE     TMHMM2.0        inside     124   153
WEBSEQUENCE     TMHMM2.0        TMhelix    154   176
WEBSEQUENCE     TMHMM2.0        outside    177   297
```

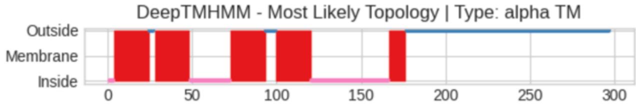


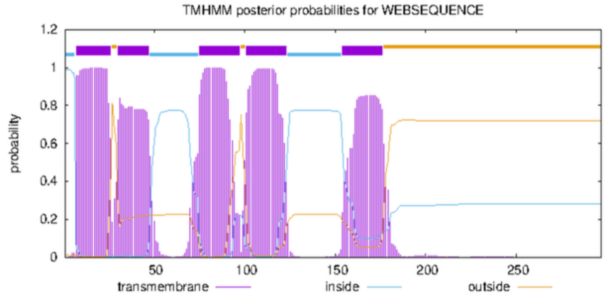


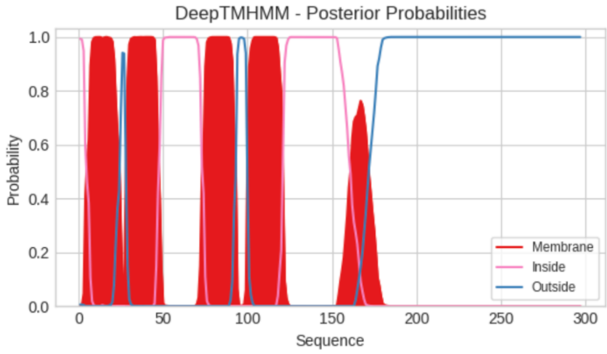


**Predicted Topologies**

```
>Sequence | TM
MKKFAKLILIFLLSTIMNGVTSYFLGDVFIFLIRELLLTLSVFLLYNYETEILDYIMESRRLDSILISKYPYLKYGGYVFFVFSFSFFFVVCFPDMVDTFIYRVAAF
LVGSSFFFSLYKLYALVQTPEAHFTPQISLKSVGHKNYRGFSSSSQWAAYFKTYFGVSTSTLFSAAAMTYPLSKIGFGMDYRYPPASAFGYVLTGLKSDSDFAFNAA
NRVVNMYGEDAIPFITETDGCTVSFSKTLKYVIFKGDDPRLLFGNLYDSSAVFEGAKAQPPKGVFDNSAEFETQKVLPPKKDG
IIIMMMMMMMMMMMMMMMMMMMMMOOOMMMMMMMMMMMMMMMMMMMMMIIIIIIIIIIIIIIIIIIIIIIIIMMMMMMMMMMMMMMMMMMMMMOOOOOOMMMMMMMM
MMMMMMMMMMMMMIIIIIIIIIIIIIIIIIIIIIIIIIIIIIIIIIIIIIIIIIIIIIIMMMMMMMMMMOOOOOOOOOOOOOOOOOOOOOOOOOOOOOOOOOOOOOO
OOOOOOOOOOOOOOOOOOOOOOOOOOOOOOOOOOOOOOOOOOOOOOOOOOOOOOOOOOOOOOOOOOOOOOOOOOOOOOOOOOO
```

```
##gff-version 3
# Sequence Length: 297
# Sequence Number of predicted TMRs: 5
Sequence inside 1 3
Sequence TMhelix 4 24
Sequence outside 25 27
Sequence TMhelix 28 48
Sequence inside 49 72
Sequence TMhelix 73 93
Sequence outside 94 99
Sequence TMhelix 100 120
Sequence inside 121 166
Sequence TMhelix 167 176
Sequence outside 177 297
```

### OrfV_2 *Pseudotetraëdriella kamillae* SAG 2056

```
# WEBSEQUENCE Length: 294
# WEBSEQUENCE Number of predicted TMHs:  4
# WEBSEQUENCE Exp number of AAs in TMHs: 96.9766200000001
# WEBSEQUENCE Exp number, first 60 AAs:  40.10888
# WEBSEQUENCE Total prob of N-in:        0.68365
# WEBSEQUENCE POSSIBLE N-term signal sequence
WEBSEQUENCE     TMHMM2.0        inside       1    11
WEBSEQUENCE     TMHMM2.0        TMhelix     12    34
WEBSEQUENCE     TMHMM2.0        outside     35    39
WEBSEQUENCE     TMHMM2.0        TMhelix     40    59
WEBSEQUENCE     TMHMM2.0        inside      60    85
WEBSEQUENCE     TMHMM2.0        TMhelix     86   108
WEBSEQUENCE     TMHMM2.0        outside    109   117
WEBSEQUENCE     TMHMM2.0        TMhelix    118   140
WEBSEQUENCE     TMHMM2.0        inside     141   294
```

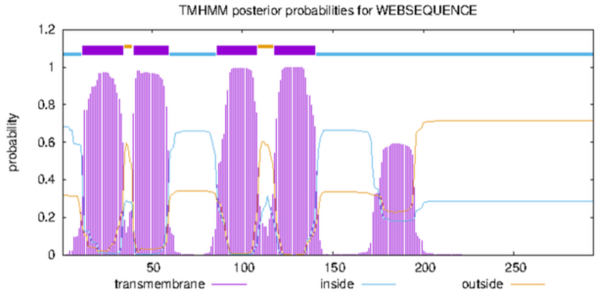


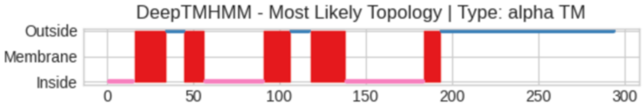


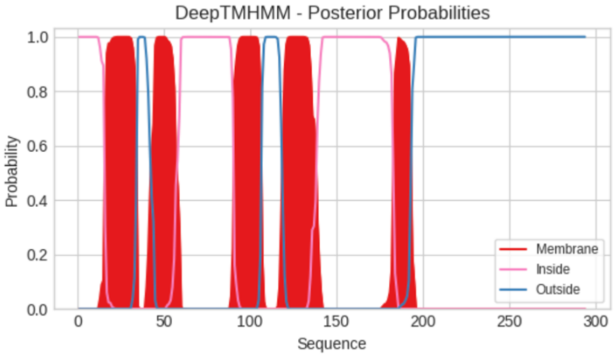


**Predicted Topologies**

```
>Sequence | TM
MARILNFFNSSMVAKALIKSLPFLAFSYLFLFVFKGSIELFFLRFFLILLFSLFTVSMGDNLKQVWTMAVIEIEASDRKLLQLYPFFKWEGPLFVFSFFGCYIVMWC
AANPGSEWGALCLVILVAQIFLAFLVILKVLAIVTTPDIREHYNVIRSIKLHGFRSFGTISKVAALCKNCLTTTAGVFVGGVAAPILIDGNLDARGFAVNIFANLVH
DTGASTHEEMQRFLELKKKSPEEAAKVIGKDWTGWIKNVDSVLLDEASKKAGILKGRPLVSVDFDHDVKPGAFYKSFWRP
IIIIIIIIIIIIIIIMMMMMMMMMMMMMMMMMMMOOOOOOOOOOMMMMMMMMMMMMIIIIIIIIIIIIIIIIIIIIIIIIIIIIIIIIIIMMMMMMMMMMMMMMMMO
OOOOOOOOOOMMMMMMMMMMMMMMMMMMMMMIIIIIIIIIIIIIIIIIIIIIIIIIIIIIIIIIIIIIIIIIIIIIIIIMMMMMMMMMMOOOOOOOOOOOOOOOOOOOOO
OOOOOOOOOOOOOOOOOOOOOOOOOOOOOOOOOOOOOOOOOOOOOOOOOOOOOOOOOOOOOOOOOOOOOOOOOOOOOOOO
```

```
##gff-version 3
# Sequence Length: 294
# Sequence Number of predicted TMRs: 5
Sequence inside 1 15
Sequence TMhelix 16 34
Sequence outside 35 44
Sequence TMhelix 45 56
Sequence inside 57 90
Sequence TMhelix 91 106
Sequence outside 107 117
Sequence TMhelix 118 138
Sequence inside 139 183
Sequence TMhelix 184 193
Sequence outside 194 294
```

### OrfV_3 *Pseudotetraëdriella kamillae* SAG 2056

```
# WEBSEQUENCE Length: 321
# WEBSEQUENCE Number of predicted TMHs:  5
# WEBSEQUENCE Exp number of AAs in TMHs: 111.63504
# WEBSEQUENCE Exp number, first 60 AAs:  42.57983
# WEBSEQUENCE Total prob of N-in:        0.40935
# WEBSEQUENCE POSSIBLE N-term signal sequence
WEBSEQUENCE     TMHMM2.0        inside       1     4
WEBSEQUENCE     TMHMM2.0        TMhelix      5    22
WEBSEQUENCE     TMHMM2.0        outside     23    36
WEBSEQUENCE     TMHMM2.0        TMhelix     37    59
WEBSEQUENCE     TMHMM2.0        inside      60    63
WEBSEQUENCE     TMHMM2.0        TMhelix     64    86
WEBSEQUENCE     TMHMM2.0        outside     87   100
WEBSEQUENCE     TMHMM2.0        TMhelix    101   123
WEBSEQUENCE     TMHMM2.0        inside     124   135
WEBSEQUENCE     TMHMM2.0        TMhelix    136   158
WEBSEQUENCE     TMHMM2.0        outside    159   321
```

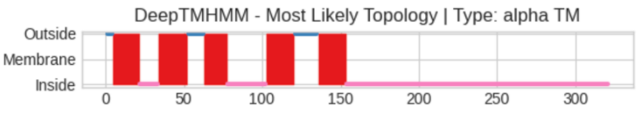


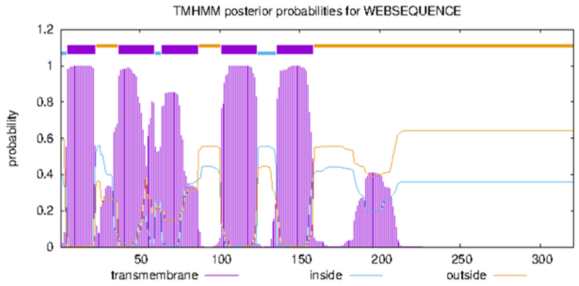


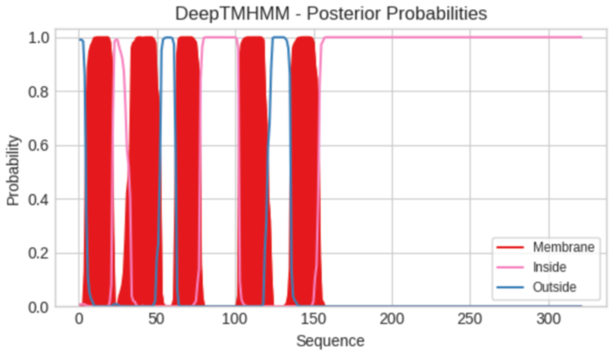


**Predicted Topologies**

```
>Sequence | TM
MEHYLILSTLLFFIGLFGIVLNRKNILIVIFKEISPVLIFWTVFNFLCLYIFEDIFFFLYRQIFLVVGGFTVGYIKIQYKELFLLFLSENRHIEDVFKLQYPILNLY
IPLLVLSVVGLYFNILKGEDPEYDVYCRIFVYLQVSFLVISFLKIFYSVTTPEIASYYNLPKAIKTGGVRSFGTAKVLLHYCKFCGYGIAGCFVGDGAYAYLSTGDP
DSMGVIPTQAALRSQKFSSRTFSGMRNALICKDRLPPEEFNKLLDWDLTNKRSVANAARIEKVMRDKLKIEPFGPSSFRPRVSTEHKIGWDFSVLKAFKNMFGKKPD
OOOOMMMMMMMMMMMMMMMMMIIIIIIIIIIIIMMMMMMMMMMMMMMMMMMMOOOOOOOOOOMMMMMMMMMMMMMMMIIIIIIIIIIIIIIIIIIIIIIIIIMMMMM
MMMMMMMMMMMMMOOOOOOOOOOOOOOOMMMMMMMMMMMMMMMMMMIIIIIIIIIIIIIIIIIIIIIIIIIIIIIIIIIIIIIIIIIIIIIIIIIIIIIIIIIIII
IIIIIIIIIIIIIIIIIIIIIIIIIIIIIIIIIIIIIIIIIIIIIIIIIIIIIIIIIIIIIIIIIIIIIIIIIIIIIIIIIIIIIIIIIIIIIIIIIIIIIIIIIII

##gff-version 3
# Sequence Length: 321
# Sequence Number of predicted TMRs: 5
Sequence outside 1 4
Sequence TMhelix 5 21
Sequence inside 22 33
Sequence TMhelix 34 52
Sequence outside 53 62
Sequence TMhelix 63 77
Sequence inside 78 102
Sequence TMhelix 103 120
Sequence outside 121 135
Sequence TMhelix 136 153
Sequence inside 154 321
```

### OrfV_1 Monodopsidaceae sp. WarPS-5

```
# WEBSEQUENCE Length: 302
# WEBSEQUENCE Number of predicted TMHs:  4
# WEBSEQUENCE Exp number of AAs in TMHs: 98.3350900000001
# WEBSEQUENCE Exp number, first 60 AAs:  37.27716
# WEBSEQUENCE Total prob of N-in:        0.88070
# WEBSEQUENCE POSSIBLE N-term signal sequence
WEBSEQUENCE     TMHMM2.0        inside       1    12
WEBSEQUENCE     TMHMM2.0        TMhelix     13    32
WEBSEQUENCE     TMHMM2.0        outside     33    35
WEBSEQUENCE     TMHMM2.0        TMhelix     36    55
WEBSEQUENCE     TMHMM2.0        inside      56    75
WEBSEQUENCE     TMHMM2.0        TMhelix     76    98
WEBSEQUENCE     TMHMM2.0        outside     99   107
WEBSEQUENCE     TMHMM2.0        TMhelix    108   130
WEBSEQUENCE     TMHMM2.0        inside     131   302
```

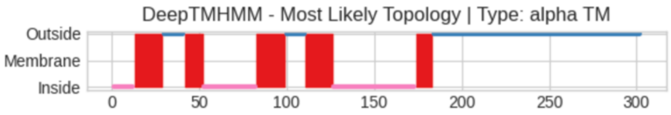


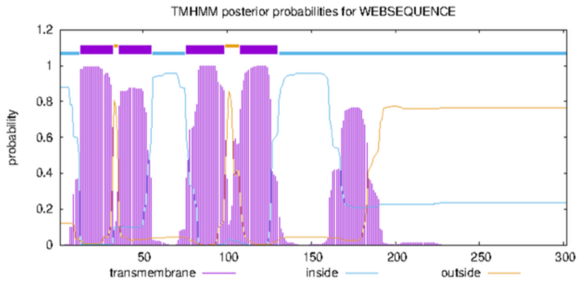


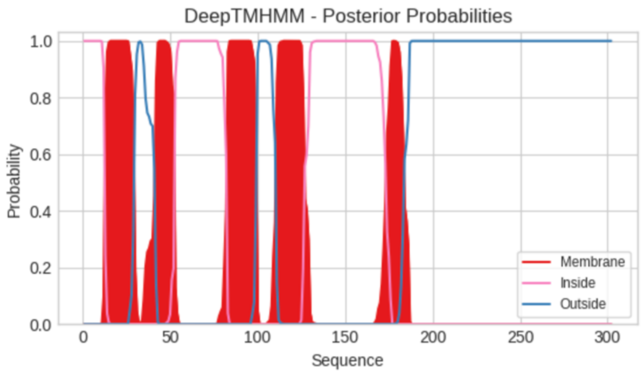


**Predicted Topologies**

```
>Sequence | TM
MSNLNKNTIFNPIFLFAFLSGLNMLFCFFVLYDPIIFTIREIFIFLFVVLRFKYSKEFSLIVHDIKEMDNILISKYPILKYQFFFFFAGFILFLGLSFVPDYSLYLP
GLYFLSYFSLFVAGLLLIYRMFVSFLTPELAHFSDIRTSVKLHGFRSFSTTSKLAVICKNCFYGSMGAGLILLGAPKLLCLNLDIRSPFSNFMSKHLLYGGIVSTSE
YTWDLAFEIYQIDPNYNVYNKNGFADRELCRKFFIDRGMDVPKPKAFLEVTHDVRHSLTVPSLPKLLKKESLSEWIEYWTWQSPPKKP
IIIIIIIIIIIIMMMMMMMMMMMMMMMMMOOOOOOOOOOOOMMMMMMMMMMMIIIIIIIIIIIIIIIIIIIIIIIIIIIIIMMMMMMMMMMMMMMMMMMOOOOOOOO
OOOMMMMMMMMMMMMMMMMIIIIIIIIIIIIIIIIIIIIIIIIIIIIIIIIIIIIIIIIIIIIIIIMMMMMMMMMMOOOOOOOOOOOOOOOOOOOOOOOOOOOOOOO
OOOOOOOOOOOOOOOOOOOOOOOOOOOOOOOOOOOOOOOOOOOOOOOOOOOOOOOOOOOOOOOOOOOOOOOOOOOOOOOOOOOOOOOO
```

```
##gff-version 3
# Sequence Length: 302
# Sequence Number of predicted TMRs: 5
Sequence inside 1 12
Sequence TMhelix 13 29
Sequence outside 30 41
Sequence TMhelix 42 52
Sequence inside 53 82
Sequence TMhelix 83 99
Sequence outside 100 110
Sequence TMhelix 111 126
Sequence inside 127 173
Sequence TMhelix 174 183
Sequence outside 184 302
```

### OrfV_2 Monodopsidaceae sp. WarPS-5

```
# WEBSEQUENCE Length: 290
# WEBSEQUENCE Number of predicted TMHs:  4
# WEBSEQUENCE Exp number of AAs in TMHs: 86.4644799999999
# WEBSEQUENCE Exp number, first 60 AAs:  31.65751
# WEBSEQUENCE Total prob of N-in:        0.98124
# WEBSEQUENCE POSSIBLE N-term signal sequence
WEBSEQUENCE     TMHMM2.0        inside       1    16
WEBSEQUENCE     TMHMM2.0        TMhelix     17    36
WEBSEQUENCE     TMHMM2.0        outside     37    50
WEBSEQUENCE     TMHMM2.0        TMhelix     51    70
WEBSEQUENCE     TMHMM2.0        inside      71    89
WEBSEQUENCE     TMHMM2.0        TMhelix     90   108
WEBSEQUENCE     TMHMM2.0        outside    109   122
WEBSEQUENCE     TMHMM2.0        TMhelix    123   145
WEBSEQUENCE     TMHMM2.0        inside     146   290
```

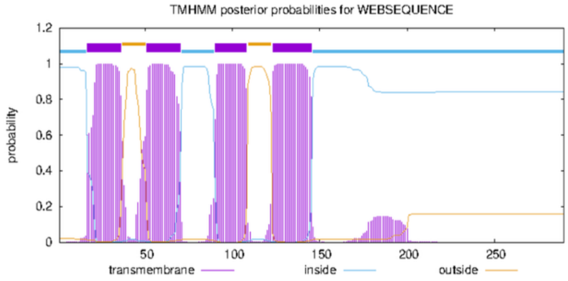


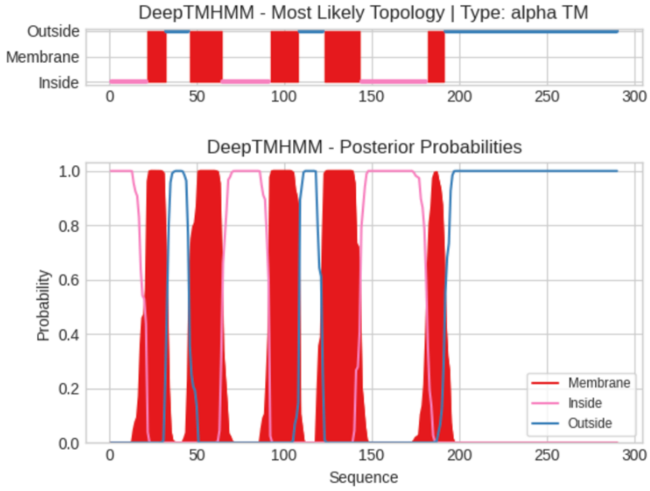


**Predicted Topologies**

```
>Sequence | TM
MRISLFIGSIIRKFADFILRKLLYLLIFTFFNYMYCIEFKNDIQLFLIRELFVIVVLAIADFFSDYLSILSKDTQYIEETIYRDFPILKYRLLVAGCILFILFAIPG
IKLDRNSEFSEFIPFIISLLGLLTVLFIVMMLIHLLAILSNPQISKYYDVIGAIRHHGVRSFATAKALVQVCTYCVKGAGALILCDFTYPIVVDSNLTARSSIANYI
SLHHRPYHMMSRLGTDLNDAIHIVERNPELRSQIVDSNNTVIPGKVDEIYRTPSKYGLYRPGASLQADVNFKWWPF
IIIIIIIIIIIIIIIIIIIIIMMMMMMMMMMMOOOOOOOOOOOOOMMMMMMMMMMMMMMMMMMMIIIIIIIIIIIIIIIIIIIIIIIIIIIMMMMMMMMMMMMMMMM
MOOOOOOOOOOOOOOMMMMMMMMMMMMMMMMMMMMMIIIIIIIIIIIIIIIIIIIIIIIIIIIIIIIIIIIIIIMMMMMMMMMMOOOOOOOOOOOOOOOOOOOOOOOO
OOOOOOOOOOOOOOOOOOOOOOOOOOOOOOOOOOOOOOOOOOOOOOOOOOOOOOOOOOOOOOOOOOOOOOOOOOOO

##gff-version 3
# Sequence Length: 290
# Sequence Number of predicted TMRs: 5
Sequence inside 1 21
Sequence TMhelix 22 32
Sequence outside 33 45
Sequence TMhelix 46 64
Sequence inside 65 91
Sequence TMhelix 92 108
Sequence outside 109 122
Sequence TMhelix 123 143
Sequence inside 144 181
Sequence TMhelix 182 191
Sequence outside 192 290
```

### OrfV_2 Chlorobotryaceae sp. Wtwin 8/9 T-6m6.8

```
# WEBSEQUENCE Length: 283
# WEBSEQUENCE Number of predicted TMHs:  3
# WEBSEQUENCE Exp number of AAs in TMHs: 86.22408
# WEBSEQUENCE Exp number, first 60 AAs:  21.18155
# WEBSEQUENCE Total prob of N-in:        0.21146
# WEBSEQUENCE POSSIBLE N-term signal sequence
WEBSEQUENCE     TMHMM2.0        outside      1    43
WEBSEQUENCE     TMHMM2.0        TMhelix     44    66
WEBSEQUENCE     TMHMM2.0        inside      67    92
WEBSEQUENCE     TMHMM2.0        TMhelix     93   112
WEBSEQUENCE     TMHMM2.0        outside    113   121
WEBSEQUENCE     TMHMM2.0        TMhelix    122   144
WEBSEQUENCE     TMHMM2.0        inside     145   283
```

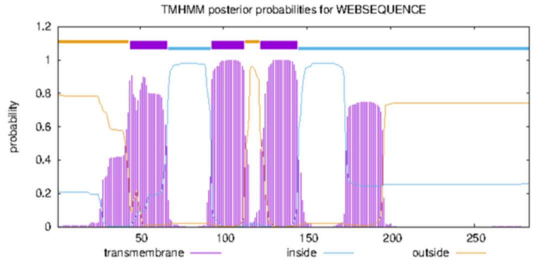


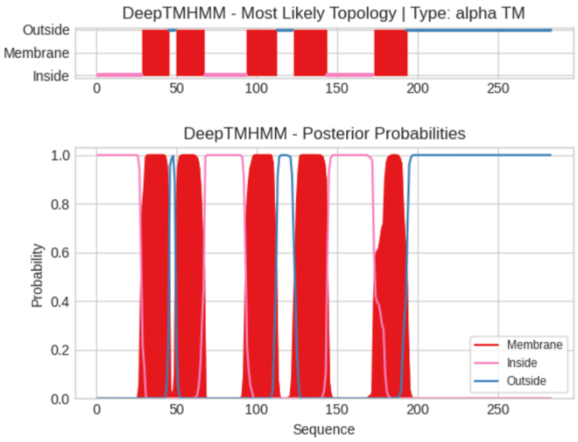

**Predicted Topologies**

```
>Sequence | TM
MILLNLQLLFLLKPWFSKNWVIFKQTSAYRIVVGLTLLNSFFFFFHLEPVVFFLRQLALLVWVGFYMKNQQLFSDINSEDAEVRLLLKRHYPIFTYEPYTMVLAFLG
MFVQITYTDSAFLQLVSPFLVFVYGVLLLVKVLFILGTLEIREVYKKALISPANMGKRRFSSFSKAAVHGCKLCLQAAVGFGFSAWIGPKFYMGDYDYRGDAWNSIT
GPLAGYKSRSEQLCREANYLCLYDKDAIAKVTDSNGMLDKALIDQQYQVKGIAKPVTIVVEHRFPWSRK
IIIIIIIIIIIIIIIIIIIIIIIIIIIIMMMMMMMMMMMMMMMMMOOOOMMMMMMMMMMMMMMMMMMIIIIIIIIIIIIIIIIIIIIIIIIIIMMMMMMMMMMMMMM
MMMMMOOOOOOOOOOMMMMMMMMMMMMMMMMMMMMMIIIIIIIIIIIIIIIIIIIIIIIIIIIIIMMMMMMMMMMMMMMMMMMMMMOOOOOOOOOOOOOOOOOOOOO
OOOOOOOOOOOOOOOOOOOOOOOOOOOOOOOOOOOOOOOOOOOOOOOOOOOOOOOOOOOOOOOOOOOOO

##gff-version 3
# Sequence Length: 283
# Sequence Number of predicted TMRs: 5
Sequence inside 1 28
Sequence TMhelix 29 45
Sequence outside 46 49
Sequence TMhelix 50 67
Sequence inside 68 93
Sequence TMhelix 94 112
Sequence outside 113 122
Sequence TMhelix 123 143
Sequence inside 144 172
Sequence TMhelix 173 193
Sequence outside 194 283
```

### OrfW

#### OrfW_1 Goniochloridales sp. BogD 8/9 T-2w

```
# WEBSEQUENCE Length: 299
# WEBSEQUENCE Number of predicted TMHs:  5
# WEBSEQUENCE Exp number of AAs in TMHs: 102.51518
# WEBSEQUENCE Exp number, first 60 AAs:  45.83893
# WEBSEQUENCE Total prob of N-in:        0.88637
# WEBSEQUENCE POSSIBLE N-term signal sequence
WEBSEQUENCE     TMHMM2.0        inside       1     2
WEBSEQUENCE     TMHMM2.0        TMhelix      3    25
WEBSEQUENCE     TMHMM2.0        outside     26    29
WEBSEQUENCE     TMHMM2.0        TMhelix     30    47
WEBSEQUENCE     TMHMM2.0        inside      48    51
WEBSEQUENCE     TMHMM2.0        TMhelix     52    71
WEBSEQUENCE     TMHMM2.0        outside     72    80
WEBSEQUENCE     TMHMM2.0        TMhelix     81    98
WEBSEQUENCE     TMHMM2.0        inside      99   104
WEBSEQUENCE     TMHMM2.0        TMhelix    105   122
WEBSEQUENCE     TMHMM2.0        outside    123   299
```

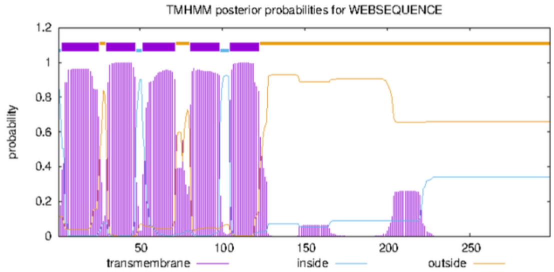


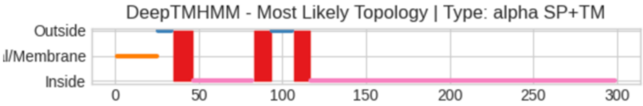


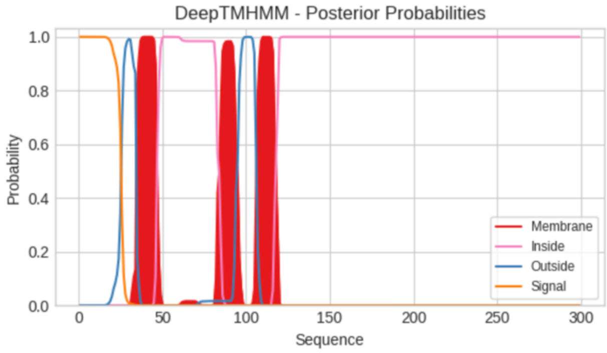


**Predicted Topologies**

```
>Sequence | SP+TM
MNFSLLFGTLPFSLLFGLIPYRILGVFHSDYFLRLIIGIFFFFLFFFQMKRLLVKILNIPCVQLLFFYLLTFENFIYTMQPWVRYLFLFSFFFNLGAWPNQTEDPLF
WYLLVLFFFTFYRQMRFIFLNPEFDGSSYTTQTICKEDDLTWNNIVLNMSSICLSFFGKPIQNQLGQVLSQRGFGQLSKRYVFTSRLFGSVKTHPELATTIAGTLGA
GLGFGISEYRNSIRHSELLNETKRTNLIQEQAVQVQKQLADEAKRANDLKEKELMLQFSQTGKKVTDLSTSDFLPPVFLKDAFFF
SSSSSSSSSSSSSSSSSSSSSSSSSOOOOOOOOOMMMMMMMMMMMMIIIIIIIIIIIIIIIIIIIIIIIIIIIIIIIIIIIIMMMMMMMMMMMOOOOOOOOOOOOOM
MMMMMMMMMIIIIIIIIIIIIIIIIIIIIIIIIIIIIIIIIIIIIIIIIIIIIIIIIIIIIIIIIIIIIIIIIIIIIIIIIIIIIIIIIIIIIIIIIIIIIII
IIIIIIIIIIIIIIIIIIIIIIIIIIIIIIIIIIIIIIIIIIIIIIIIIIIIIIIIIIIIIIIIIIIIIIIIIIIIIIIIIIIII

##gff-version 3
# Sequence Length: 299
# Sequence Number of predicted TMRs: 3
Sequence signal 1 25
Sequence outside 26 34
Sequence TMhelix 35 46
Sequence inside 47 82
Sequence TMhelix 83 93
Sequence outside 94 106
Sequence TMhelix 107 116
Sequence inside 117 299
```

#### OrfW_2 Goniochloridales sp. BogD 8/9 T-2w

```
# WEBSEQUENCE Length: 318
# WEBSEQUENCE Number of predicted TMHs:  4
# WEBSEQUENCE Exp number of AAs in TMHs: 91.66464
# WEBSEQUENCE Exp number, first 60 AAs:  41.37924
# WEBSEQUENCE Total prob of N-in:        0.50444
# WEBSEQUENCE POSSIBLE N-term signal sequence
WEBSEQUENCE     TMHMM2.0        inside       1     6
WEBSEQUENCE     TMHMM2.0        TMhelix      7    26
WEBSEQUENCE     TMHMM2.0        outside     27    35
WEBSEQUENCE     TMHMM2.0        TMhelix     36    58
WEBSEQUENCE     TMHMM2.0        inside      59    64
WEBSEQUENCE     TMHMM2.0        TMhelix     65    87
WEBSEQUENCE     TMHMM2.0        outside     88    96
WEBSEQUENCE     TMHMM2.0        TMhelix     97   119
WEBSEQUENCE     TMHMM2.0        inside     120   318
```

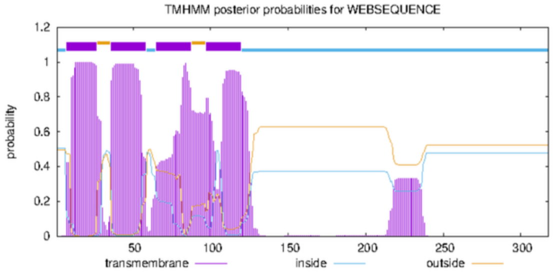


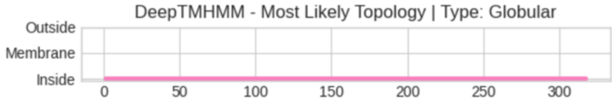


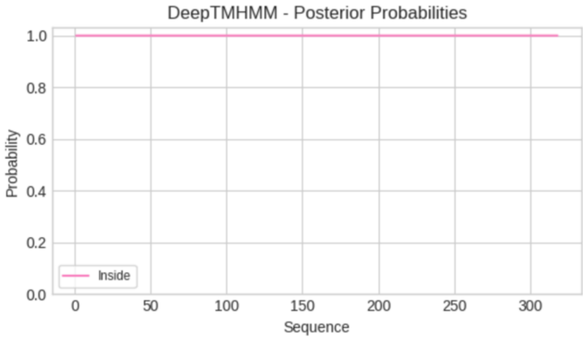


**Predicted Topologies**

```
>Sequence | GLOB
MEQIKKLPPLSFFLLLFILFFRLSLLFGLYDIESRIFFVVFNLLISCYFWKPLLLRTLNSRLFQLWFLFCFSIENWLMCSPLLFLYSCFTWTLATVWFSFLIFDDLN
LTLNVAISILLFHHLYFSLQKMFLDAQLDFLTVEIRSPQDLTWGNILKAISEFSTHYCGFPIHNRLNEKHLSRLFPNTTKRFMFKSAGMVFRQNMGKLQGFASEHPD
ALKVVSGVITATAGGLGFFASLDIKKQELDIKKQSLELEKYKIDKESETRCYVANKQVEVEQLKLQILKVAKDQTDGVKSQIEVKAPSLCNRLEDISLSNFFFF
IIIIIIIIIIIIIIIIIIIIIIIIIIIIIIIIIIIIIIIIIIIIIIIIIIIIIIIIIIIIIIIIIIIIIIIIIIIIIIIIIIIIIIIIIIIIIIIIIIIIIIIIIII
IIIIIIIIIIIIIIIIIIIIIIIIIIIIIIIIIIIIIIIIIIIIIIIIIIIIIIIIIIIIIIIIIIIIIIIIIIIIIIIIIIIIIIIIIIIIIIIIIIIIIIIIIII
IIIIIIIIIIIIIIIIIIIIIIIIIIIIIIIIIIIIIIIIIIIIIIIIIIIIIIIIIIIIIIIIIIIIIIIIIIIIIIIIIIIIIIIIIIIIIIIIIIIIIIII

##gff-version 3
# Sequence Length: 318
# Sequence Number of predicted TMRs: 0
Sequence inside 1 318
```

### OrfW_3 Goniochloridales sp. BogD 8/9 T-2w

```
# WEBSEQUENCE Length: 336
# WEBSEQUENCE Number of predicted TMHs:  4
# WEBSEQUENCE Exp number of AAs in TMHs: 91.3108400000002
# WEBSEQUENCE Exp number, first 60 AAs:  42.34521
# WEBSEQUENCE Total prob of N-in:        0.00353
# WEBSEQUENCE POSSIBLE N-term signal sequence
WEBSEQUENCE     TMHMM2.0        outside      1     9
WEBSEQUENCE     TMHMM2.0        TMhelix     10    29
WEBSEQUENCE     TMHMM2.0        inside      30    35
WEBSEQUENCE     TMHMM2.0        TMhelix     36    58
WEBSEQUENCE     TMHMM2.0        outside     59    80
WEBSEQUENCE     TMHMM2.0        TMhelix     81   103
WEBSEQUENCE     TMHMM2.0        inside     104   109
WEBSEQUENCE     TMHMM2.0        TMhelix    110   132
WEBSEQUENCE     TMHMM2.0        outside    133   336
```

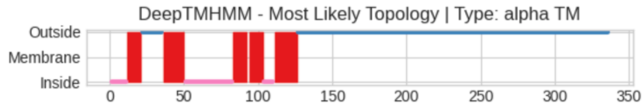


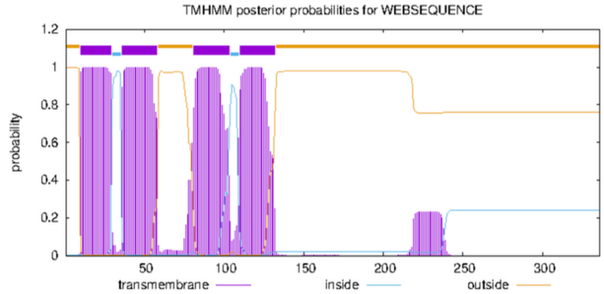


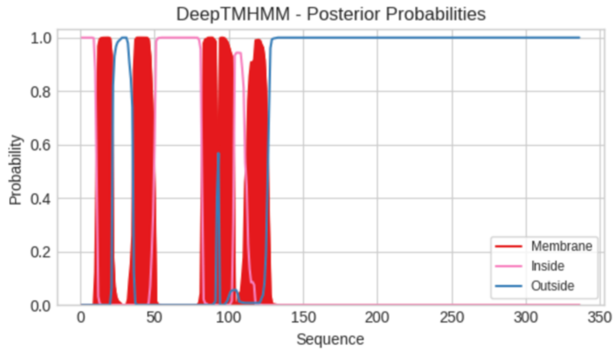


**Predicted Topologies**

```
>Sequence | TM
MEFFGNKPPLAFWLLLSLFFFRLSLVVGFYDIIHRIFFVFVSLCLFFYFFFKWIFYRVLNSTLFQYWFFFGLTLENFLIRMPFLVSLLLFLVWPFVIVCWTYLLSNNI
QLTLNVAISLVLFYHLYFGIQNMFLNPQMVYSTFEVESPNDITWENILKAISGVSNNYCGLPIANKLKQQDLSRCFPYIAKRFMFKNAAKAMDANMGRLQSFVVEHP
DACKIVGAALTTIVAGGASLMTLNLEKEKMASSIEVAKITAKAAESAAKATESAAKATESAANISLQAAKVSADAIIYKADKDFKAQQLKFQASSSDSSVEVKAPSL
FADKADWVWYDFLYP
IIIIIIIIIIIMMMMMMMMMMOOOOOOOOOOOOOOMMMMMMMMMMMMMMMIIIIIIIIIIIIIIIIIIIIIIIIIIIIIIIIMMMMMMMMMMOMMMMMMMMMMIIII
IIIMMMMMMMMMMMMMMMMOOOOOOOOOOOOOOOOOOOOOOOOOOOOOOOOOOOOOOOOOOOOOOOOOOOOOOOOOOOOOOOOOOOOOOOOOOOOOOOOOOOOOO
OOOOOOOOOOOOOOOOOOOOOOOOOOOOOOOOOOOOOOOOOOOOOOOOOOOOOOOOOOOOOOOOOOOOOOOOOOOOOOOOOOOOOOOOOOOOOOOOOOOOOOOOOOO
OOOOOOOOOOOOOOO

##gff-version 3
# Sequence Length: 336
# Sequence Number of predicted TMRs: 5
Sequence inside 1 11
Sequence TMhelix 12 21
Sequence outside 22 35
Sequence TMhelix 36 50
Sequence inside 51 82
Sequence TMhelix 83 92
Sequence outside 93 93
Sequence TMhelix 94 103
Sequence inside 104 110
Sequence TMhelix 111 126
Sequence outside 127 336
```

### OrfW_1 Neomonodontaceae sp. BogD 8/9 T-6m

```
# WEBSEQUENCE Length: 408
# WEBSEQUENCE Number of predicted TMHs:  3
# WEBSEQUENCE Exp number of AAs in TMHs: 117.03181
# WEBSEQUENCE Exp number, first 60 AAs:  21.93132
# WEBSEQUENCE Total prob of N-in:        0.49882
# WEBSEQUENCE POSSIBLE N-term signal sequence
WEBSEQUENCE     TMHMM2.0        inside       1    35
WEBSEQUENCE     TMHMM2.0        TMhelix     36    58
WEBSEQUENCE     TMHMM2.0        outside     59    61
WEBSEQUENCE     TMHMM2.0        TMhelix     62    84
WEBSEQUENCE     TMHMM2.0        inside      85   121
WEBSEQUENCE     TMHMM2.0        TMhelix    122   144
WEBSEQUENCE     TMHMM2.0        outside    145   408
```

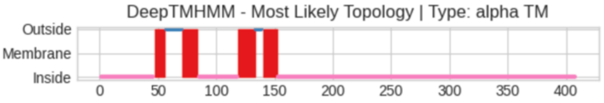


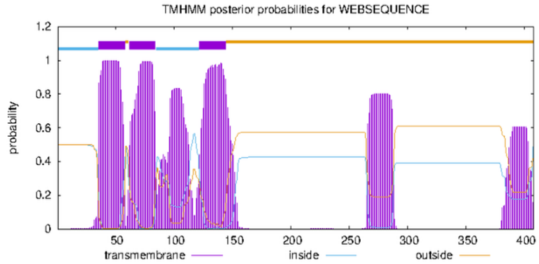


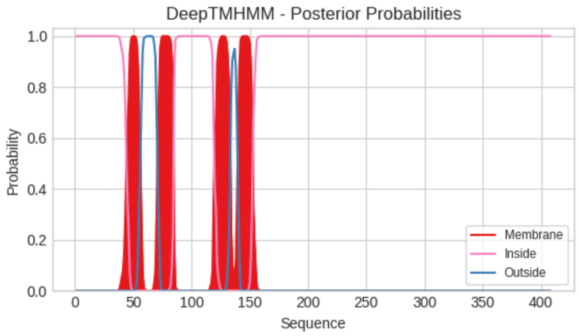


**Predicted Topologies**

```
>Sequence | TM
MLAFLFKKTMRWSWKNHFLTGLSEIKSLFFRLLKTIGPFFSIIMVEMCFILFLVHIPYKLLSAINYTMPVWSWGVLSFLLLFLFRKTILVASLNSRVIQAFVFFFLS
LENYLYRSIHNIIRFLSGFFIIVCLVDGLDGMVGFYTLISFLFLFNLHQQFCRLVLNPESIPGENREDSDLKVRNREHFTWAHVVKMMSRYSESLTGKKITSRFPGR
SDTSFPFAVKKFMIHSQAWLIAKEVTKEATKKLKEGGIEGLLDKVTVYNKPLSYIGTGLVSAGAGSGAVIFYWQHQQDLNIKEKDLALREKIVEEDLKIKNRVADAE
AALKDAQKDEIRERIRRANEDHIRSLEAPPTLGDVIWGRKRLPEPSSVKAPSLIKEEESFWDVFEFMLWELLEFLGYFYIWVKFFFL
IIIIIIIIIIIIIIIIIIIIIIIIIIIIIIIIIIIIIIIIIIIIIMMMMMMMMMMMOOOOOOOOOOOOOOMMMMMMMMMMMMMMIIIIIIIIIIIIIIIIIIIIIII
IIIIIIIIIIMMMMMMMMMMMMMMMMOOOOOOMMMMMMMMMMMMMIIIIIIIIIIIIIIIIIIIIIIIIIIIIIIIIIIIIIIIIIIIIIIIIIIIIIIIIIIIIII
IIIIIIIIIIIIIIIIIIIIIIIIIIIIIIIIIIIIIIIIIIIIIIIIIIIIIIIIIIIIIIIIIIIIIIIIIIIIIIIIIIIIIIIIIIIIIIIIIIIIIIIIIII
IIIIIIIIIIIIIIIIIIIIIIIIIIIIIIIIIIIIIIIIIIIIIIIIIIIIIIIIIIIIIIIIIIIIIIIIIIIIIIIIIIIIIII
```

```
##gff-version 3
# Sequence Length: 408
# Sequence Number of predicted TMRs: 4
Sequence inside 1 46
Sequence TMhelix 47 56
Sequence outside 57 70
Sequence TMhelix 71 84
Sequence inside 85 118
Sequence TMhelix 119 133
Sequence outside 134 139
Sequence TMhelix 140 152
Sequence inside 153 408
```

### OrfW_2 Neomonodontaceae sp. BogD 8/9 T-6m

```
# WEBSEQUENCE Length: 345
# WEBSEQUENCE Number of predicted TMHs:  3
# WEBSEQUENCE Exp number of AAs in TMHs: 95.9934500000002
# WEBSEQUENCE Exp number, first 60 AAs:  31.15922
# WEBSEQUENCE Total prob of N-in:        0.73291
# WEBSEQUENCE POSSIBLE N-term signal sequence
WEBSEQUENCE     TMHMM2.0        inside       1    38
WEBSEQUENCE     TMHMM2.0        TMhelix     39    61
WEBSEQUENCE     TMHMM2.0        outside     62    73
WEBSEQUENCE     TMHMM2.0        TMhelix     74    91
WEBSEQUENCE     TMHMM2.0        inside      92    95
WEBSEQUENCE     TMHMM2.0        TMhelix     96   118
WEBSEQUENCE     TMHMM2.0        outside    119   345
```

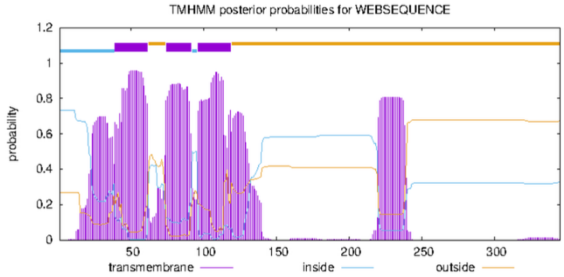


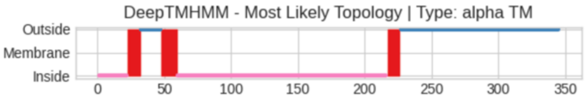


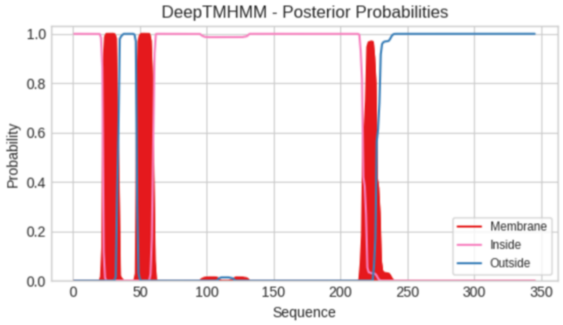


**Predicted Topologies**

```
>Sequence | TM
MRSFLQTIFRDLKTFFKTYKTELLFSFFIGLIPFKILDLCGIEVPTKFVLVILVVLILFLFRKKIAPFLLDNVVFQWWFFSVCFFENFVFSTYSRLFLFYITLAFVV
IYADILDFDLLDSFVVFLLFYGFYRKLSFFTLNPDSVFPKKSGNLGLTKDEDLTWEYMISMFSIFSERVSGRAVKNRLPSHYSPLKFPFIVKRYMIRPIFRELWENR
HSIKEFISWIGGGALAVGAIGIAAQKNLNKAEQLRVQEESNRLERERLDHDRERLRFDKDKYAGKPSDSLPSTKILPVYEGPDGKITRTPHFPPLETKAPSMESSSF
SFSDFLSFCEAFFSLNNLSSIVNF
IIIIIIIIIIIIIIIIIIIIIIMMMMMMMMMMOOOOOOOOOOOOOOOMMMMMMMMMMMMIIIIIIIIIIIIIIIIIIIIIIIIIIIIIIIIIIIIIIIIIIIIIIIII
IIIIIIIIIIIIIIIIIIIIIIIIIIIIIIIIIIIIIIIIIIIIIIIIIIIIIIIIIIIIIIIIIIIIIIIIIIIIIIIIIIIIIIIIIIIIIIIIIIIIIIIIIIII
IIMMMMMMMMMMOOOOOOOOOOOOOOOOOOOOOOOOOOOOOOOOOOOOOOOOOOOOOOOOOOOOOOOOOOOOOOOOOOOOOOOOOOOOOOOOOOOOOOOOOOOOOOO
OOOOOOOOOOOOOOOOOOOOOOOO

##gff-version 3
# Sequence Length: 345
# Sequence Number of predicted TMRs: 3
Sequence inside 1 22
Sequence TMhelix 23 32
Sequence outside 33 47
Sequence TMhelix 48 59
Sequence inside 60 216
Sequence TMhelix 217 226
Sequence outside 227 345
```

### OrfW_3 Neomonodontaceae sp. BogD 8/9 T-6m

```
# WEBSEQUENCE Length: 431
# WEBSEQUENCE Number of predicted TMHs:  5
# WEBSEQUENCE Exp number of AAs in TMHs: 110.95866
# WEBSEQUENCE Exp number, first 60 AAs:  42.00789
# WEBSEQUENCE Total prob of N-in:        0.91023
# WEBSEQUENCE POSSIBLE N-term signal sequence
WEBSEQUENCE     TMHMM2.0        inside       1     1
WEBSEQUENCE     TMHMM2.0        TMhelix      2    21
WEBSEQUENCE     TMHMM2.0        outside     22    25
WEBSEQUENCE     TMHMM2.0        TMhelix     26    45
WEBSEQUENCE     TMHMM2.0        inside      46    57
WEBSEQUENCE     TMHMM2.0        TMhelix     58    80
WEBSEQUENCE     TMHMM2.0        outside     81    99
WEBSEQUENCE     TMHMM2.0        TMhelix    100   122
WEBSEQUENCE     TMHMM2.0        inside     123   217
WEBSEQUENCE     TMHMM2.0        TMhelix    218   240
WEBSEQUENCE     TMHMM2.0        outside    241   431
```

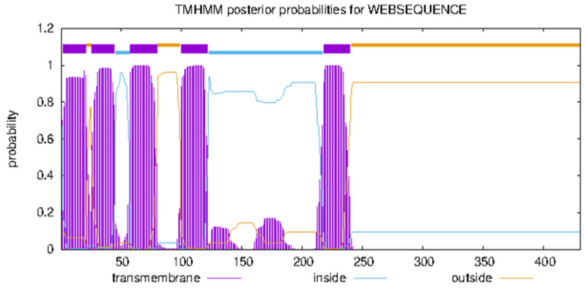


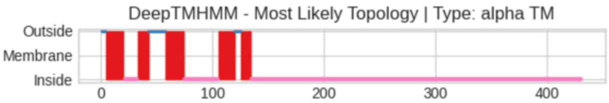


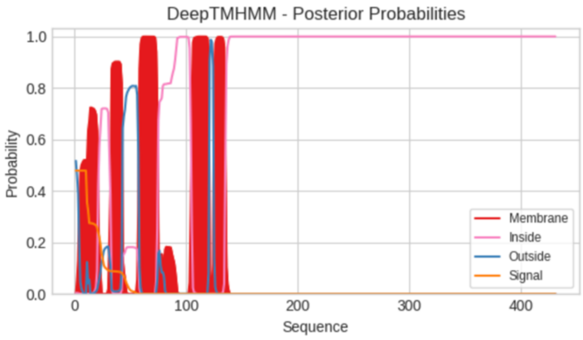

**Predicted Topologies**

```
>Sequence | TM
MLYLFLVFLISATFGTVNLFYWARGVTPPNVSLFMFFIFFLLRCLDGLSQKQLLENCLIVLAFLLILALVFIFFSRILNSFVWQSFVFFCFRLEESFFKIYPYGVIF
TSMGFAILVALLLHGDRFGYLLASFFIGTYFHTKNQFLQPELVAEYEEQEKKKIYLFSWEDFITLVNFVGLKVYGTGVQNKLSAKSVKRTFPFLQQRYLTAGRQTLR
NVSGIVFGAGGIAGFIVALANHYGLITTKKMELDHVTYTQRHDTARKFETQIKDLEISLVKCIKNRTVCDEYTLLTRETVDSIAKQAVYDLCTRKHNLDPQEVVESW
MACERRLEIIRFMQRYPTSEDALAQINQAVRRQVRLFDASDISISRMQNVPLTPVSDQGAVVTSPFSDQLGESEGDTTTDSVSNSSGILTITQGRANSMVQPSFFDY
FWS
OOOMMMMMMMMMMMMMMMMMIIIIIIIIIIIIMMMMMMMMMMOOOOOOOOOOOOOOOMMMMMMMMMMMMMMMMMIIIIIIIIIIIIIIIIIIIIIIIIIIIIIIMMM
MMMMMMMMMMMMMOOOOMMMMMMMMMMIIIIIIIIIIIIIIIIIIIIIIIIIIIIIIIIIIIIIIIIIIIIIIIIIIIIIIIIIIIIIIIIIIIIIIIIIIIIII
IIIIIIIIIIIIIIIIIIIIIIIIIIIIIIIIIIIIIIIIIIIIIIIIIIIIIIIIIIIIIIIIIIIIIIIIIIIIIIIIIIIIIIIIIIIIIIIIIIIIIIIIIII
IIIIIIIIIIIIIIIIIIIIIIIIIIIIIIIIIIIIIIIIIIIIIIIIIIIIIIIIIIIIIIIIIIIIIIIIIIIIIIIIIIIIIIIIIIIIIIIIIIIIIIIIIII
III

##gff-version 3
# Sequence Length: 431
# Sequence Number of predicted TMRs: 5
Sequence outside 1 3
Sequence TMhelix 4 20
Sequence inside 21 32
Sequence TMhelix 33 42
Sequence outside 43 57
Sequence TMhelix 58 74
Sequence inside 75 104
Sequence TMhelix 105 120
Sequence outside 121 124
Sequence TMhelix 125 134
Sequence inside 135 431
```

### OrfW_4 Neomonodontaceae sp. BogD 8/9 T-6m

```
# WEBSEQUENCE Length: 443
# WEBSEQUENCE Number of predicted TMHs:  4
# WEBSEQUENCE Exp number of AAs in TMHs: 115.40856
# WEBSEQUENCE Exp number, first 60 AAs:  36.1687
# WEBSEQUENCE Total prob of N-in:        0.79057
# WEBSEQUENCE POSSIBLE N-term signal sequence
WEBSEQUENCE     TMHMM2.0        inside       1     4
WEBSEQUENCE     TMHMM2.0        TMhelix      5    27
WEBSEQUENCE     TMHMM2.0        outside     28    57
WEBSEQUENCE     TMHMM2.0        TMhelix     58    80
WEBSEQUENCE     TMHMM2.0        inside      81   100
WEBSEQUENCE     TMHMM2.0        TMhelix    101   120
WEBSEQUENCE     TMHMM2.0        outside    121   214
WEBSEQUENCE     TMHMM2.0        TMhelix    215   237
WEBSEQUENCE     TMHMM2.0        inside     238   443
```

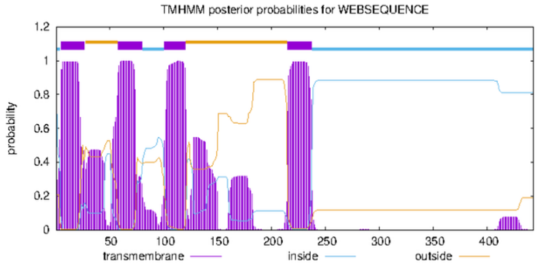


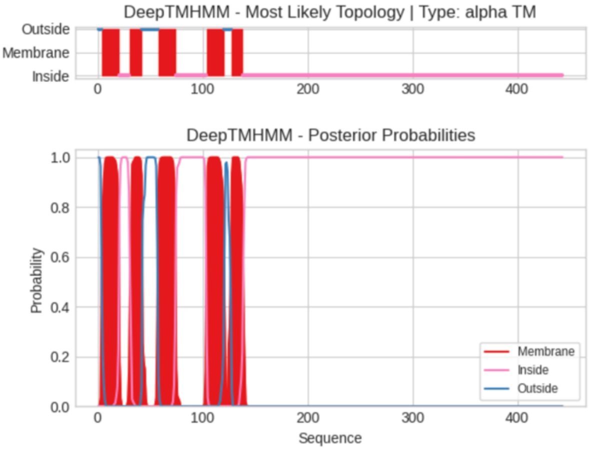


**Predicted Topologies**

```
>Sequence | TM
MYSLILLLVVGVIVGTINLFYWARGVTPPNASLSMFLMFFFLRSLNRGLQEYPLGEGIIILVFLLILAVLYVFMSKLLNSYLWQCFVFFCFWLEDEFFRRYPYSLVV
TSIFFGILTWCLIFEDDFPYLLLSFLIGMYLNTKNQFLQPILLKEYEKMEKKNLNVVCWDDFLILVNFVALKVYGTKVENRLSPRLSRYPFSFIQQRYLGAGRESMP
TILTAVFGSGGIAGVLLMISNYFTSVSTKKMELDHSTITDYRRRIQDLQKDIVDIDGVIRKCLSKNNVCASYSLFTRLKLDMLAEIEAYRHAMDHLSVNPIDLFAEW
KVCPRRFEAINYIKKHPTSEEALAQIQLLIRQRIRSFDSSLTEGAYVSKSLVDVDSEYYRELLAKSPIANGVPSIDYDDEGIQVDLKDPFSGIAGGVILTLKGPAIA
PSLKKWNIFDFFPFL
OOOMMMMMMMMMMMMMMMMMIIIIIIIIIIMMMMMMMMMMMMOOOOOOOOOOOOOOOMMMMMMMMMMMMMMMMMIIIIIIIIIIIIIIIIIIIIIIIIIIIIIMMMM
MMMMMMMMMMMMMOOOOOOOMMMMMMMMMMMIIIIIIIIIIIIIIIIIIIIIIIIIIIIIIIIIIIIIIIIIIIIIIIIIIIIIIIIIIIIIIIIIIIIIIIIII
IIIIIIIIIIIIIIIIIIIIIIIIIIIIIIIIIIIIIIIIIIIIIIIIIIIIIIIIIIIIIIIIIIIIIIIIIIIIIIIIIIIIIIIIIIIIIIIIIIIIIIIIIII
IIIIIIIIIIIIIIIIIIIIIIIIIIIIIIIIIIIIIIIIIIIIIIIIIIIIIIIIIIIIIIIIIIIIIIIIIIIIIIIIIIIIIIIIIIIIIIIIIIIIIIIIIII
IIIIIIIIIIIIIII

##gff-version 3
# Sequence Length: 443
# Sequence Number of predicted TMRs: 5
Sequence outside 1 3
Sequence TMhelix 4 20
Sequence inside 21 30
Sequence TMhelix 31 42
Sequence outside 43 57
Sequence TMhelix 58 74
Sequence inside 75 103
Sequence TMhelix 104 120
Sequence outside 121 127
Sequence TMhelix 128 138
Sequence inside 139 443
```

### OrfW_1 *Vacuoliviride crystalliferum* NIES-2860

```
# WEBSEQUENCE Length: 296
# WEBSEQUENCE Number of predicted TMHs:  4
# WEBSEQUENCE Exp number of AAs in TMHs: 110.42294
# WEBSEQUENCE Exp number, first 60 AAs:  42.32419
# WEBSEQUENCE Total prob of N-in:        0.99415
# WEBSEQUENCE POSSIBLE N-term signal sequence
WEBSEQUENCE     TMHMM2.0        inside        1     6
WEBSEQUENCE     TMHMM2.0        TMhelix       7    26
WEBSEQUENCE     TMHMM2.0        outside      27    35
WEBSEQUENCE     TMHMM2.0        TMhelix      36    58
WEBSEQUENCE     TMHMM2.0        inside       59    64
WEBSEQUENCE     TMHMM2.0        TMhelix      65    87
WEBSEQUENCE     TMHMM2.0        outside      88   106
WEBSEQUENCE     TMHMM2.0        TMhelix     107   129
WEBSEQUENCE     TMHMM2.0        inside      130   296
```

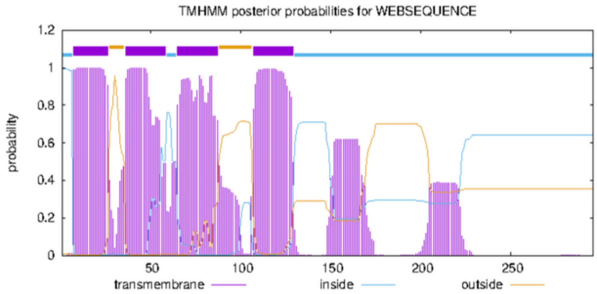


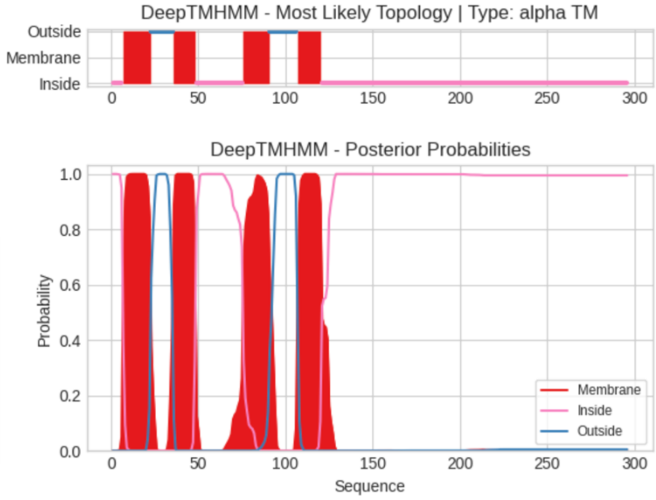


**Predicted Topologies**

```
>Sequence | TM
MQKFKNFFVYLPFSLLFGLIPYRILGIFQADYSTRLIIGILFFILFFYQMKNLVVKILNNPVIQLLFFYLLTFENFIYLMSPFFRYLFLFSFFFNIGAWPNQTEDPL
FWYVFAIFLFTFYRQMLAIFVNPEFDGSLYSIKTVRKEDDLKWDFIVINMSSICLLFFGKPIQNQLGQILSQRSFGQLSKRYVFTSRFFASVKTHPELATAIISSLG
AGVGFGISEYRNSIRHSELLEQQRQANQIQNQLADEAKRANDLKERELLLQLPKPLNENIKNVSGMDSLFVPMISTKDIFFL
IIIIIIMMMMMMMMMMMMMMMMOOOOOOOOOOOOOMMMMMMMMMMMMMIIIIIIIIIIIIIIIIIIIIIIIIIIIMMMMMMMMMMMMMMMOOOOOOOOOOOOOOOOM
MMMMMMMMMMMMMIIIIIIIIIIIIIIIIIIIIIIIIIIIIIIIIIIIIIIIIIIIIIIIIIIIIIIIIIIIIIIIIIIIIIIIIIIIIIIIIIIIIIIIIIII
IIIIIIIIIIIIIIIIIIIIIIIIIIIIIIIIIIIIIIIIIIIIIIIIIIIIIIIIIIIIIIIIIIIIIIIIIIIIIIIIII

##gff-version 3
# Sequence Length: 296
# Sequence Number of predicted TMRs: 4
Sequence inside 1 6
Sequence TMhelix 7 22
Sequence outside 23 35
Sequence TMhelix 36 48
Sequence inside 49 75
Sequence TMhelix 76 90
Sequence outside 91 106
Sequence TMhelix 107 120
Sequence inside 121 296
```

### OrfW_2 *Vacuoliviride crystalliferum* NIES-2860

```
# WEBSEQUENCE Length: 298
# WEBSEQUENCE Number of predicted TMHs:  4
# WEBSEQUENCE Exp number of AAs in TMHs: 89.73892
# WEBSEQUENCE Exp number, first 60 AAs:  38.24003
# WEBSEQUENCE Total prob of N-in:        0.94849
# WEBSEQUENCE POSSIBLE N-term signal sequence
WEBSEQUENCE     TMHMM2.0        inside        1    12
WEBSEQUENCE     TMHMM2.0        TMhelix      13    32
WEBSEQUENCE     TMHMM2.0        outside      33    41
WEBSEQUENCE     TMHMM2.0        TMhelix      42    64
WEBSEQUENCE     TMHMM2.0        inside       65    70
WEBSEQUENCE     TMHMM2.0        TMhelix      71    93
WEBSEQUENCE     TMHMM2.0        outside      94   107
WEBSEQUENCE     TMHMM2.0        TMhelix     108   125
WEBSEQUENCE     TMHMM2.0        inside      126   298
```

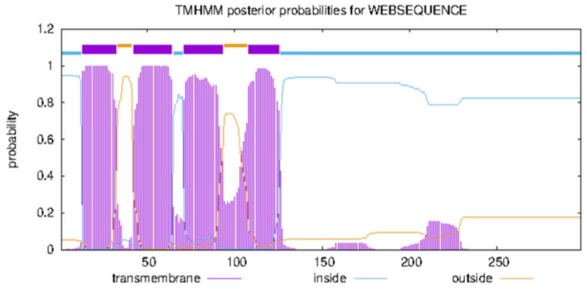


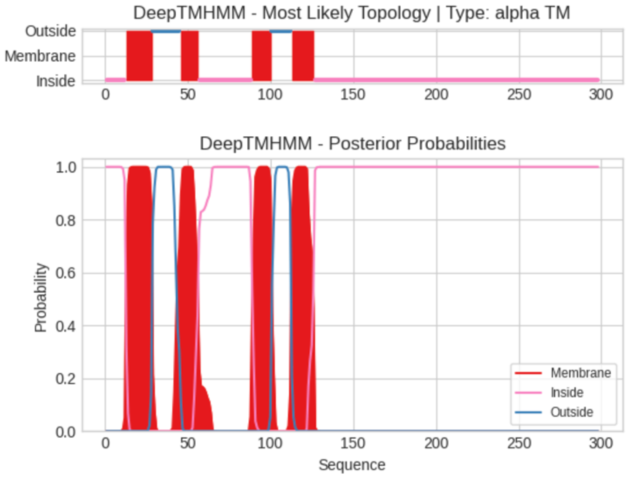


**Predicted Topologies**

```
>Sequence | TM
MQKLRIDFSANKFLIYLPFALLFGLIPYRLCGIFQMDYNYRLYLGLFFFVLFLCTMGELVILLLNSIAVQLLFYYLLLFENFVLGMRPFYLYSFLVSFFFCLGGGPS
LTADSMFWYILILFLYSFYRQMRHIFVNPQFENSSYVSSHIKKVEDLTWDLIIQNMSTICLSLFGQPVQNQLGKPMTQRSFNQISKRYVWGRIYGFAKAHPEAVTFV
STIASATGVTWGLYNQKTAEQNAVFRHKEQMDFEKLRHKEVMDLERDKLAFEINKLKLGGAKVKDLSSKEDFYRIIVDEPFFFL
IIIIIIIIIIIIMMMMMMMMMMMMMMMMOOOOOOOOOOOOOOOOOMMMMMMMMMMMIIIIIIIIIIIIIIIIIIIIIIIIIIIIIIIIMMMMMMMMMMMMOOOOOOO
OOOOOMMMMMMMMMMMMMMIIIIIIIIIIIIIIIIIIIIIIIIIIIIIIIIIIIIIIIIIIIIIIIIIIIIIIIIIIIIIIIIIIIIIIIIIIIIIII
IIIIIIIIIIIIIIIIIIIIIIIIIIIIIIIIIIIIIIIIIIIIIIIIIIIIIIIIIIIIIIIIIIIIIIIIIIIIIIIIIIII

##gff-version 3
# Sequence Length: 298
# Sequence Number of predicted TMRs: 4
```

```
Sequence inside 1 12
Sequence TMhelix 13 28
Sequence outside 29 45
Sequence TMhelix 46 56
Sequence inside 57 88
Sequence TMhelix 89 100
Sequence outside 101 112
Sequence TMhelix 113 126
Sequence inside 127 298
```

#### OrfW_3 *Vacuoliviride crystalliferum* NIES-2860

```
# WEBSEQUENCE Length: 338
# WEBSEQUENCE Number of predicted TMHs:  4
# WEBSEQUENCE Exp number of AAs in TMHs: 85.12756
# WEBSEQUENCE Exp number, first 60 AAs:  42.71718
# WEBSEQUENCE Total prob of N-in:        0.08122
# WEBSEQUENCE POSSIBLE N-term signal sequence
WEBSEQUENCE	TMHMM2.0	outside	     1	  14
WEBSEQUENCE	TMHMM2.0	TMhelix	    15	  37
WEBSEQUENCE	TMHMM2.0	inside	    38	  38
WEBSEQUENCE	TMHMM2.0	TMhelix	    39	  61
WEBSEQUENCE	TMHMM2.0	outside	    62	  80
WEBSEQUENCE	TMHMM2.0	TMhelix	    81	 103
WEBSEQUENCE	TMHMM2.0	inside	   104	 109
WEBSEQUENCE	TMHMM2.0	TMhelix	   110	 124
WEBSEQUENCE	TMHMM2.0	outside	   125	 338
```

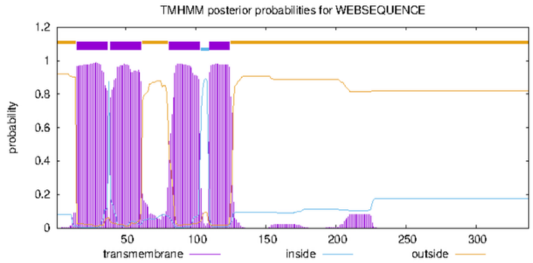


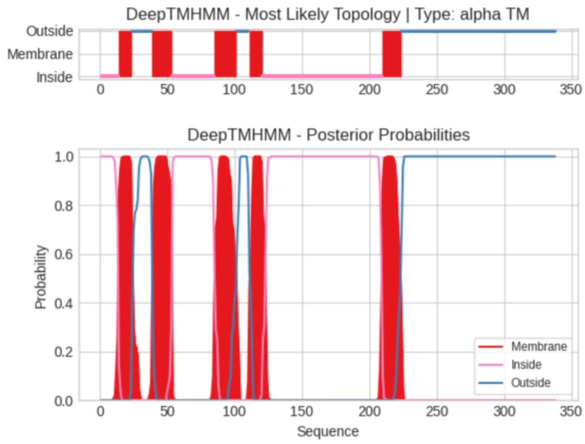


**Predicted Topologies**

```
>Sequence | TM
MNFFSLSKEFKVKFAFVLLFGLIPYRILGIVGISGLWKVVVGLGFLALFYVLMKEITPLLLNSHFLQLFFFFCLRVEEYLCTTSLFYALFFVSCIPLELTILAGNTQ
DPYFWYFLLLFLVNLYYRLKEVFLNPQFEGSVYPELEAKVEHNENKFTWNFIVLNMSAISKGLFKVSVENQLSKVMSQRSFPYIAKRYVFTSVLRAIKGHPEIISAA
GVTASIVGVVYSEQKNAQRHQESLQVTRETSEATVKAQLEVASSQDRLALAQDRLALAQEKVARSQDRLALVQEKVAQEMALARTNPVGALQLKQSLKEGEVKAPSL
SKDHPEELFSFLKFFFF
IIIIIIIIIIIIIMMMMMMMMMMOOOOOOOOOOOOOOOMMMMMMMMMMMMMMMIIIIIIIIIIIIIIIIIIIIIIIIIIIIIIIMMMMMMMMMMMMMMMMMOOOOOO
OOOMMMMMMMMMMIIIIIIIIIIIIIIIIIIIIIIIIIIIIIIIIIIIIIIIIIIIIIIIIIIIIIIIIIIIIIIIIIIIIIIIIIIIIIIIIIIIIIIIIIIIIMMMMM
MMMMMMMMMOOOOOOOOOOOOOOOOOOOOOOOOOOOOOOOOOOOOOOOOOOOOOOOOOOOOOOOOOOOOOOOOOOOOOOOOOOOOOOOOOOOOOOOOOOOOO
OOOOOOOOOOOOOOOOO

##gff-version 3
# Sequence Length: 338
# Sequence Number of predicted TMRs: 5
Sequence inside 1 13
Sequence TMhelix 14 23
Sequence outside 24 38
Sequence TMhelix 39 53
Sequence inside 54 84
Sequence TMhelix 85 101
Sequence outside 102 110
Sequence TMhelix 111 120
Sequence inside 121 209
Sequence TMhelix 210 223
Sequence outside 224 338
```

#### OrfW (Orf322) *Trachydiscus minutus* CCALA 838

```
# WEBSEQUENCE Length: 322
# WEBSEQUENCE Number of predicted TMHs:  4
# WEBSEQUENCE Exp number of AAs in TMHs: 111.09867
# WEBSEQUENCE Exp number, first 60 AAs:  33.99447
# WEBSEQUENCE Total prob of N-in:        0.97679
# WEBSEQUENCE POSSIBLE N-term signal sequence
WEBSEQUENCE	TMHMM2.0	inside	     1	  20
WEBSEQUENCE	TMHMM2.0	TMhelix	    21	  40
WEBSEQUENCE	TMHMM2.0	outside	    41	  49
WEBSEQUENCE	TMHMM2.0	TMhelix	    50	  72
WEBSEQUENCE	TMHMM2.0	inside	    73	  78
WEBSEQUENCE	TMHMM2.0	TMhelix	    79	  96
WEBSEQUENCE	TMHMM2.0	outside	    97	 110
WEBSEQUENCE	TMHMM2.0	TMhelix	   111	 133
WEBSEQUENCE	TMHMM2.0	inside	   134	 322
```

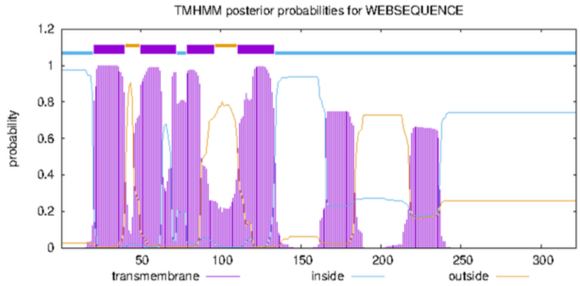


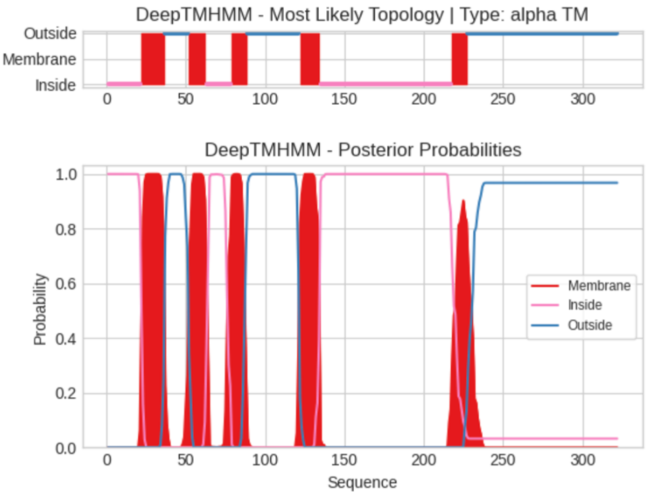

**Predicted Topologies**

```
>Sequence | TM
MTQLTFISRFYSLIKDFFRRNMFYFPFALLFGLIPYRFFGVFHLDYTYRLTIGLFFFLLFLFTMQELVTILLNNIVVQLLFFYLLAFEDFVRGMAPFYQYSFLFSFS
FSLGGGPALTEDSMFWYILILFFFSFYRQMRDIFVNPQFEASSYFPLHVKKVEDLTWDLIVINMSSICVALFGQPIQNQLGKRMSQRSFNQISKRYVWGRVIGFAKT
HPEAVTIVSSIAGGVGFTWGLYNQKISEANAMLRHKEQMHFQKIAEENLMLRHKEQMHFQKIAEENLMLRHKEHMDLEKLKLTKGRIKDLSYRKDLGDMDCDEYFFF
F
IIIIIIIIIIIIIIIIIIIIIMMMMMMMMMMMMMMMOOOOOOOOOOOOOOOMMMMMMMMMMMIIIIIIIIIIIIIIIIMMMMMMMMMMOOOOOOOOOOOOOOOOOOOO
OOOOOOOOOOOOOMMMMMMMMMMMMMIIIIIIIIIIIIIIIIIIIIIIIIIIIIIIIIIIIIIIIIIIIIIIIIIIIIIIIIIIIIIIIIIIIIIIIIIIIIIII
IIIMMMMMMMMMMOOOOOOOOOOOOOOOOOOOOOOOOOOOOOOOOOOOOOOOOOOOOOOOOOOOOOOOOOOOOOOOOOOOOOOOOOOOOOOOOOOOOOOOOOOOO
O

##gff-version 3
# Sequence Length: 322
# Sequence Number of predicted TMRs: 5
Sequence inside 1 21
Sequence TMhelix 22 36
Sequence outside 37 51
Sequence TMhelix 52 62
Sequence inside 63 78
Sequence TMhelix 79 88
Sequence outside 89 121
Sequence TMhelix 122 134
Sequence inside 135 217
Sequence TMhelix 218 227
Sequence outside 228 322
```

### OrfW Goniochloridales sp. Chic 10/23 P-6w

```
# WEBSEQUENCE Length: 314
# WEBSEQUENCE Number of predicted TMHs:  5
# WEBSEQUENCE Exp number of AAs in TMHs: 93.4047800000001
# WEBSEQUENCE Exp number, first 60 AAs:  45.87445
# WEBSEQUENCE Total prob of N-in:        0.88727
# WEBSEQUENCE POSSIBLE N-term signal sequence
WEBSEQUENCE     TMHMM2.0        inside       1     4
WEBSEQUENCE     TMHMM2.0        TMhelix      5    27
WEBSEQUENCE     TMHMM2.0        outside     28    30
WEBSEQUENCE     TMHMM2.0        TMhelix     31    48
WEBSEQUENCE     TMHMM2.0        inside      49    52
WEBSEQUENCE     TMHMM2.0        TMhelix     53    72
WEBSEQUENCE     TMHMM2.0        outside     73    75
WEBSEQUENCE     TMHMM2.0        TMhelix     76    98
WEBSEQUENCE     TMHMM2.0        inside      99   104
WEBSEQUENCE     TMHMM2.0        TMhelix    105   122
WEBSEQUENCE     TMHMM2.0        outside    123   314
```

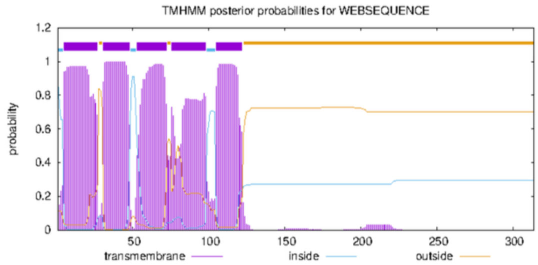


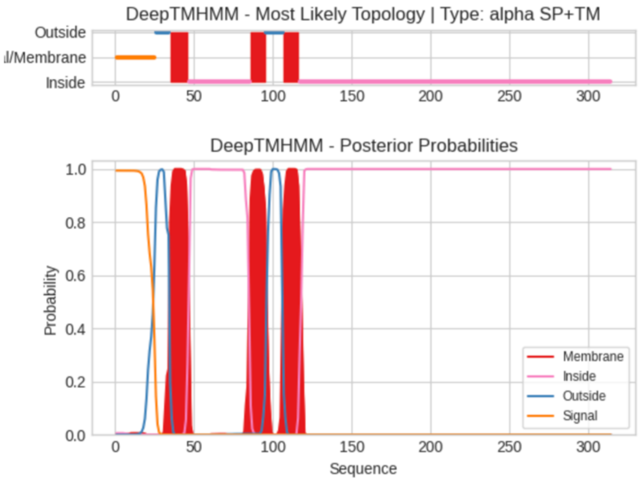


**Predicted Topologies**

```
>Sequence | SP+TM
MNFSLLFATLPFSFLFGLIPYKILGVFHTDYIFRLIVGFVFFFFFFFHRKKLVVRILNLPFTQLLFFYFLTFENFIYTMTPKVRYFFLFTFFFNLGAWPNQMQDPLF
WYLLAIFFLTFYRQMRSMFLNPEFNGSCYSTQTIQKDEDLRWETVVVNMSSICLSFFGKPIQNQLGSTLSQRTFAQLTKRYVFTSRLFGTIKTHPELTTTIAGSLGA
GLGFGVSEYRNSIRHSELLEETKRSNNLQEQAIQVQKDVAYETKQANEIQRQLAEQAKRANDLKERELLLQLPKSEKTILKDISCSNIAFRPSFHDIFFF
SSSSSSSSSSSSSSSSSSSSSSSSSOOOOOOOOOMMMMMMMMMMMMIIIIIIIIIIIIIIIIIIIIIIIIIIIIIIIIIIIIIIIMMMMMMMMMMOOOOOOOOOOOM
MMMMMMMMMIIIIIIIIIIIIIIIIIIIIIIIIIIIIIIIIIIIIIIIIIIIIIIIIIIIIIIIIIIIIIIIIIIIIIIIIIIIIIIIIIIIIIIIIIIIIIIII
IIIIIIIIIIIIIIIIIIIIIIIIIIIIIIIIIIIIIIIIIIIIIIIIIIIIIIIIIIIIIIIIIIIIIIIIIIIIIIIIIIIIIIIIIIIIIIIIIIIII

##gff-version 3
# Sequence Length: 314
# Sequence Number of predicted TMRs: 3
Sequence signal 1 25
Sequence outside 26 34
Sequence TMhelix 35 46
Sequence inside 47 85
Sequence TMhelix 86 95
Sequence outside 96 106
Sequence TMhelix 107 116
Sequence inside 117 314
```

### OrfW *Paraeustigmatos columelliferus* Mont 10/10-1w

```
# WEBSEQUENCE Length: 302
# WEBSEQUENCE Number of predicted TMHs:  3
# WEBSEQUENCE Exp number of AAs in TMHs: 71.65233
# WEBSEQUENCE Exp number, first 60 AAs:  40.85827
# WEBSEQUENCE Total prob of N-in:        0.90623
# WEBSEQUENCE POSSIBLE N-term signal sequence
WEBSEQUENCE     TMHMM2.0        inside       1     6
WEBSEQUENCE     TMHMM2.0        TMhelix      7    29
WEBSEQUENCE     TMHMM2.0        outside     30    32
WEBSEQUENCE     TMHMM2.0        TMhelix     33    55
WEBSEQUENCE     TMHMM2.0        inside      56    82
WEBSEQUENCE     TMHMM2.0        TMhelix     83   115
WEBSEQUENCE     TMHMM2.0        outside    116   302
```

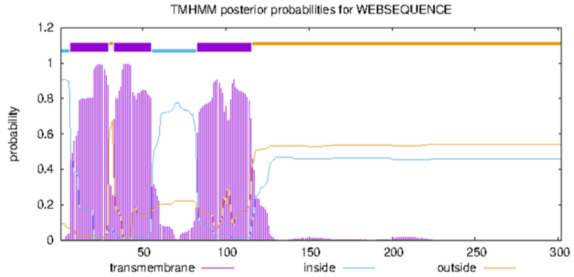


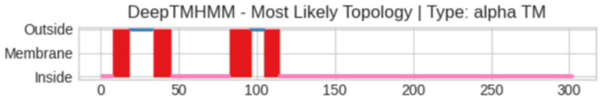


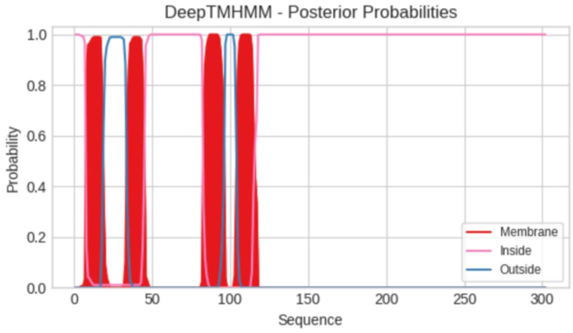


**Predicted Topologies**

```
>Sequence | TM
MKLFFRQFVQYFFIGVIPYKLLSIFGFSIEVKLVLGLIFFLIFLFKLPNIVCSLLNSVLVQKILFWVLERENFFLAKSVKKQFLLAFLTGLSIGLWSTKAPDPFFVI
FLIGFLFYCYKNFKSLFLNPILEGNYSDQNSYDISWNKIIDAVSATSIVLFKLPVQNKLALSIGSKTFPNVIKRYMFARGAKSAGEAIKQNSEVAGVLGTVLGFGIS
EFFQQSINAEIERNSQRRHQEILELQKKIDANVERRHQENLAEQRRASDIAESKVNLLKNTGEDLPSSGIPCSYEPTFTISEILEKIF
IIIIIIMMMMMMMMMMMOOOOOOOOOOOOOOOMMMMMMMMMMMMIIIIIIIIIIIIIIIIIIIIIIIIIIIIIIIIIIIIIMMMMMMMMMMMMMMOOOOOOOOMMM
MMMMMMMIIIIIIIIIIIIIIIIIIIIIIIIIIIIIIIIIIIIIIIIIIIIIIIIIIIIIIIIIIIIIIIIIIIIIIIIIIIIIIIIIIIIIIIIIIIIIIIIIII
IIIIIIIIIIIIIIIIIIIIIIIIIIIIIIIIIIIIIIIIIIIIIIIIIIIIIIIIIIIIIIIIIIIIIIIIIIIIIIIIIIIIIIIII

##gff-version 3
# Sequence Length: 302
# Sequence Number of predicted TMRs: 4
Sequence inside 1 7
Sequence TMhelix 8 18
Sequence outside 19 33
Sequence TMhelix 34 45
Sequence inside 46 82
Sequence TMhelix 83 96
Sequence outside 97 104
Sequence TMhelix 105 114
Sequence inside 115 302
```
